# *De novo* sequencing of complex glycans by ion mobility-mass spectrometry using a self-expanding reference database

**DOI:** 10.1101/2025.05.20.655041

**Authors:** Javier Sastre Toraño, Julia Vreugdenhil, Gaël M. Vos, Kevin Hooijschuur, Shannon Vogelaar, Christian Klein, John Fjeldsted, Bernd Stahl, Geert-Jan Boons

**Author notes:** These authors contributed equally (Javier Sastre Toraño, Julia Vreugdenhil).

## Abstract

It is essential to determine exact structures of glycans in complex biological samples to understand their biology and exploit their diagnostics, therapeutics and nutraceuticals potential. An unresolved analytical challenge is the identification of isomeric glycan structures in complex biological samples. Ion mobility (IM) combined with MS enables separation of isomeric glycans and identification by comparing their intrinsic collision cross section (CCS) values with similar data of synthetic standards. To identify glycans without the need to synthesize all biologically occurring glycans, we describe here an IM-MS *de novo* sequencing method based on fragment identification and sequence assembly. CCS values of additional fragments from glycans in biological samples resulted in a self-expanding reference database, gradually facilitating the sequencing of glycans of increasing complexity and expanding the database from an initial 20 standards to 332 unique entries. The methodology was employed to determine exact structures of human milk oligosaccharides and *N*-glycans of biotherapeutics.

## Introduction

All eukaryotic cells are covered by a dense layer of glycans that are mediators of many biological processes such as protein folding, cell proliferation and differentiation, modulation of cell signaling, fertilization and immune regulation. Specific changes in cell surface glycosylation are associated with diseases such as inflammatory bowel syndrome, septic shock and neurological disorders such as Alzheimer’s and Parkinson’s disease. Aberrant glycosylation is also a determinant of cancer cell proliferative, invasiveness and immune suppressive properties^1, 2, 3, 4, 5^. Furthermore, many pathogens employ glycans of host cells for attachment and/or entry and are factors of host range and tissue tropism^6^. Also, many biologicals such as monoclonal antibodies are modified by complex glycans, which determine biological activity and pharmacokinetic properties^7, 8, 9^.

To understand their functions at a molecular level, it is essential to determine the exact structures of glycans in complex biological samples. An understanding of glycan structures will make it possible to develop the next generation glycan-based diagnostics, therapeutics and nutraceuticals and will advance the development of therapeutic glycoproteins. Glycans are assembled from monosaccharides that can be linked at different sites and in different configurations resulting in many possible isomeric structures that have the same molecular formula but differ in stereochemistry of monosaccharides, anomeric configurations, linkage positions and branching^9, 10^. Current analytical approaches to analyze complex mixtures of glycans are primarily based on mass spectrometry (MS)^11, 12^. These methods provide in general compositions but not exact molecular structures because MS alone cannot readily differentiate between isomeric structures. Isomeric glycans usually have very different biological properties and for example α2,3-linked sialosides are employed as receptors by avian influenza A viruses (IAV) whereas human IAVs use α2,6-linked sialosides for cell attachment and entry^6^. Thus, it is critical to develop analytical methods that can determine exact structures of glycans.

The hyphenation of MS to ion mobility spectrometry (IMS) adds a separation dimension and offers the prospect of isomeric glycan identification^13, 14^. In IMS, gas phase ions are separated based on their mobility through a drift cell filled with a static inert buffer gas under the influence of an electric field. Higher charge state ions migrate with a higher mobility in this field, while ions with larger surface areas experience more ion-neutral collisions with buffer gas molecules and therefore their mobility is more reduced. IMS makes it possible to calculate intrinsic rotationally averaged ion-neutral collision cross section (CCS) values from drift or arrival times^15^. Isomeric intact glycan or fragment ions can have different surface areas^16, 17, 18, 19, 20^ and therefore may exhibit distinct CCS values and arrival time distributions (ATDs) that can be exploited for glycan identification.

To make ion mobility (IM)-MS suitable for exact glycan structure determination, a database is needed that can correlate mass-to-charge ratios (*m/z*), ATDs and CCS values to isomeric glycan structures. Advances in chemoenzymatic synthesis make it possible to prepare collections of isomeric glycans^21, 22, 23, 24, 25^ that can be subjected to IM-MS to establish CCS values of intact and fragment ions to populate the required database. For example, we subjected a library of synthetic *O*-acetylated sialosides^26^ to drift tube IM-MS and established intrinsic CCS values of diagnostic fragment ions which could be used to determine *O*-acetylation patterns and anomeric linkages of released *O*- and *N*-linked glycans^27^. Despite the attraction of this approach, it is not possible to prepare all possible naturally occurring glycans due to the enormous structural diversity^28^. To address this limitation, we describe here an IM-MS methodology for high-throughput exact structure identification and *de novo* sequencing of glycans using a limited number of synthetic glycan standards having common glyco-epitopes. In this approach, the standards are subjected to in-source collisional activated dissociation (CAD) to produce informative intact and fragment ions for an initial IM-MS reference database. Next, glycans obtained from a biological source can be subjected to IM-MS and rapid identification can be made for those compounds that have an entry (*m/z* and ATD or CCS values) of the intact ion in the reference database (steps 1-3, Fig. 1). Compounds that do not have an entry will be subjected to CAD and *de novo* sequencing by identifying structures of fragment ions that can be puzzled together, leveraging the established biosynthesis pathways, to elucidate the exact glycan structure (steps 4-6, Fig. 1). The premise of the approach is that although the intact ion may be absent in the database, the presence of fragment ions will facilitate *de novo* sequencing. The *m/z*, ATD, and CCS values of the elucidated intact and fragment ions can be added to the reference database for rapid identification in other samples or for the *de novo* sequencing of larger and more complex structures (Fig. 1). Thus, the IM-MS methodology continuously exploits data from previous analyses of biological samples to elucidate new glycan structures while at the same time expanding the reference database. We demonstrate here that by applying data of 12 glyco-epitope fragment ions, the reference database could be expanded to 332 entries. The IM-MS methodology with a self-expanding reference database will make it possible to identify exact structures rapidly and unambiguously in biological samples without a need for a large collection of synthetic standards.

**Fig. 1.**
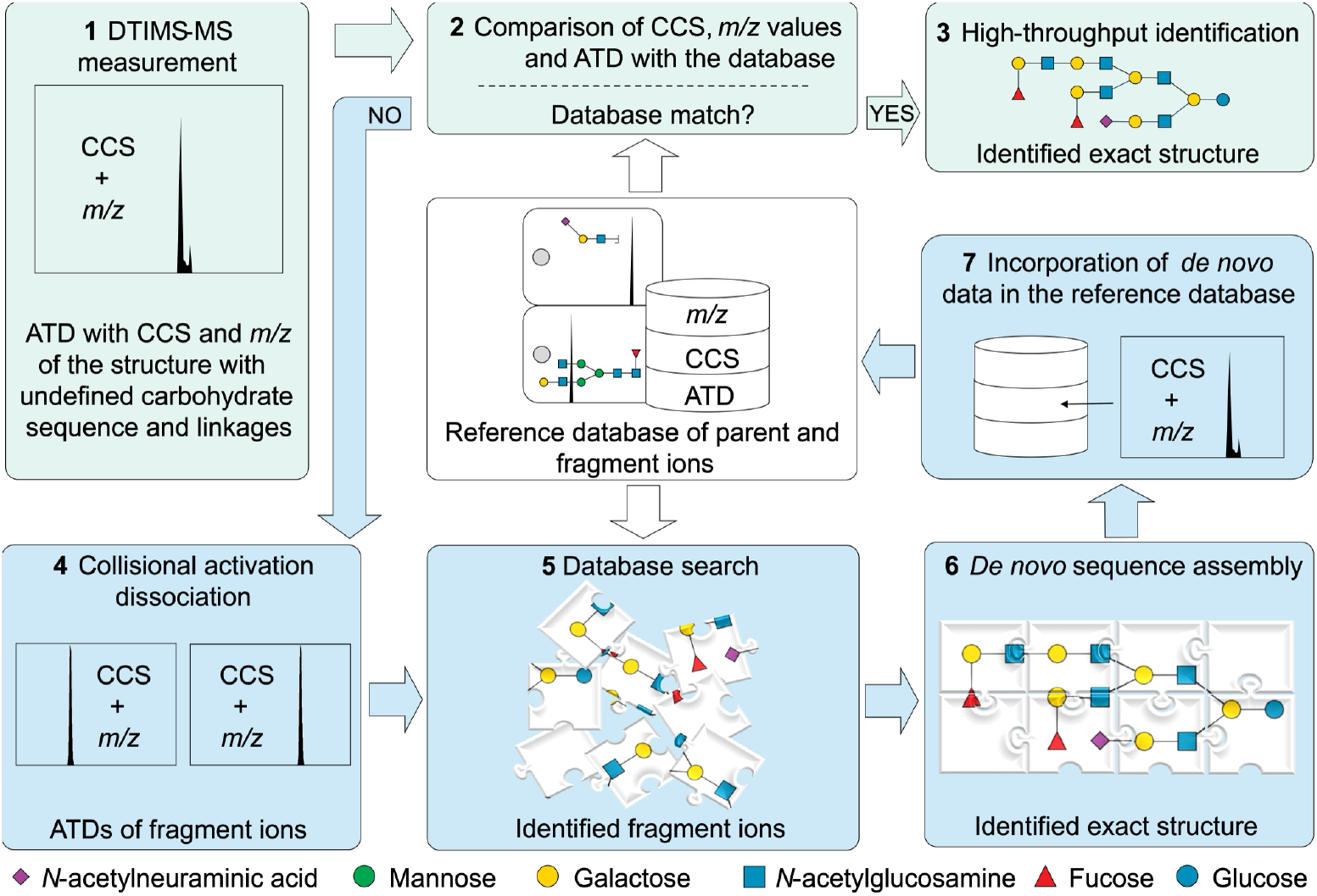
Schematic overview of the integrated IM-MS high-throughput identification and *de novo* sequencing methods for exact glycan structure determination. The green route (1-3) uses arrival time distributions (ATDs) or collision cross section (CCS) and *m/z* values of intact glycans from the reference database for high-throughput identification of glycans from biological samples with undefined sequence and linkage-type. For glycans without an entry in the reference database, the blue route is followed: Glycans are first subjected to in-source collisional activation dissociation to produce fragment ions (4). These ions are then identified using fragment ion entries from the reference database (5), which provides insights into the types of glycosidic linkages between the monosaccharides units and the overall architecture of the glycan. The assembly of the fragments (6) reconstructs the full glycan structure, enabling the accurate identification of complex compounds. ATDs and CCS values of the elucidated intact structure and of newly identified fragment ions are included in the reference database (7) for future high-throughput identification in other samples and *de novo* sequencing of other unknown structures. Each step in the process builds on the previous one, progressively leading to the identification of structures with increasing complexity.

## Results

### Identification of exact structures of HMOs

To develop the *de-novo* sequencing methodology, we first prepared and analyzed human milk oligosaccharides (HMOs), which contain epitopes commonly found in other glycan classes. HMOs possess probiotic properties by serving as metabolic substrates for beneficial bacteria, thereby shaping the intestinal microbiota of breast-fed infants^29, 30, 31^. In addition, they protect from infections by acting as decoys for receptors of pathogens^32, 33^. HMOs can also modulate epithelial and immune cell responses and reduce excessive mucosal leukocyte infiltration and activation. The health benefits of HMOs correlate to specific structures and their abundance,^34, 35, 36^ which may vary between milk donors and changes during lactation. Given the health benefits of HMOs, this class of compounds is receiving increasing attention for the development of functional foods.

All HMOs have a reducing lactose moiety that can be extended by *N*-acetyl lactosamine (LacNAc; galactose-β(1,4)-*N*-acetylglucosamine) to give a type II chain (compound **1**, Fig. 2a) or Lacto-*N*-biose (LNB; galactose-β(1,3)*-N*-acetylglucosamine) to give a type I chain (compound **2**, Fig. 2a). The resulting tetrasaccharides can be functionalized with fucosides or sialosides or be converted into an I-branched structure by the addition of *N*-acetylglucosamine (GlcNAc) to the penultimate galactose. Iterative additions of LacNAc or LNB structures, followed by fucosylation and sialylation results in structurally very diverse compounds.

**Fig. 2.**
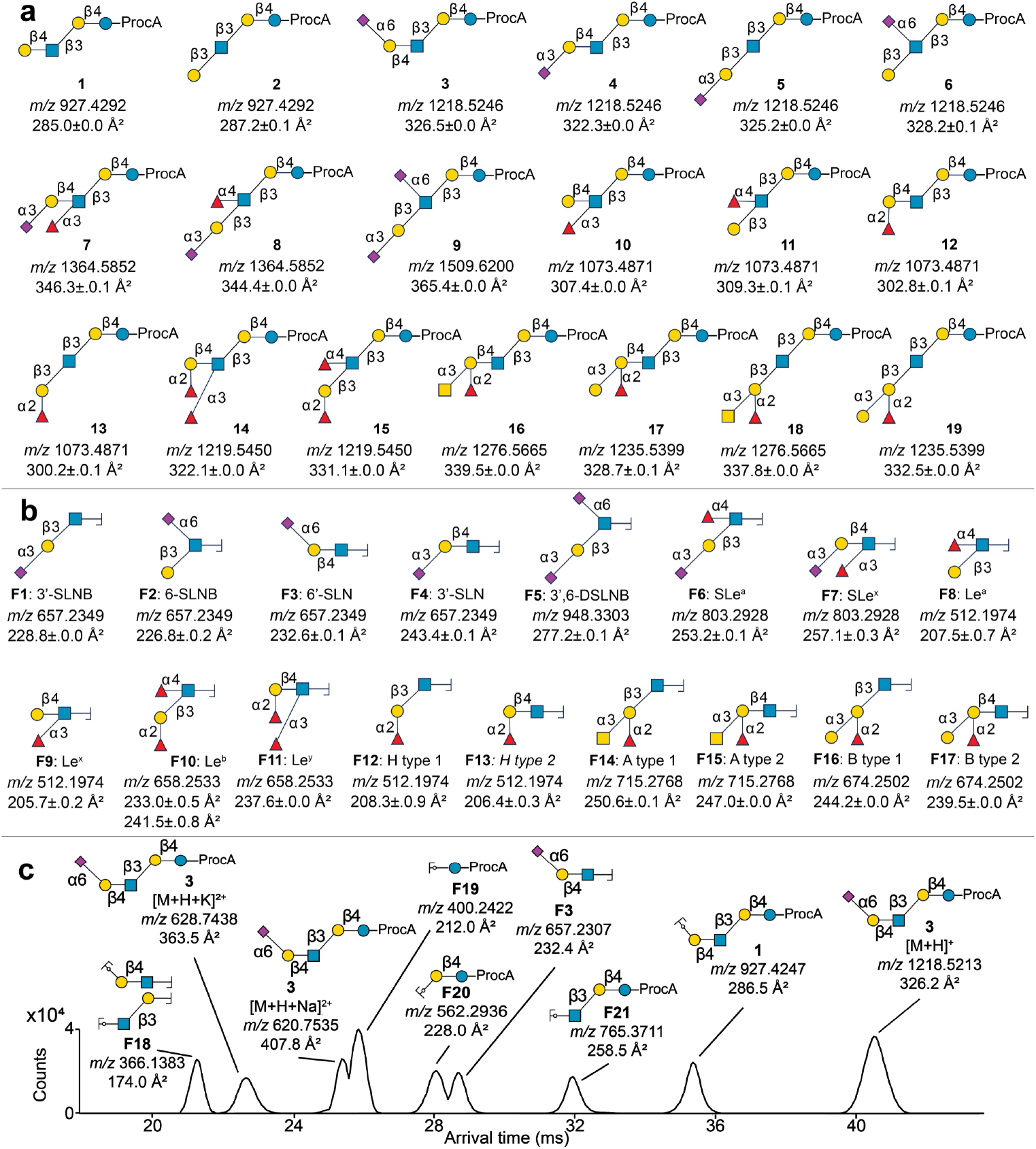
HMO and glyco-epitope library for the development of high-throughput identification and *de novo* sequencing of glycans. **a** Structures, theoretical *m/z* values (as M+H]^+^ ions) and measured ^DT^CCS_N2_ values (n=3 measurements) of synthetic HMO standards, derivatized with procainamide (ProcA) at their reducing end by reductive amination. The library includes structures of the sialyl-lacto-*N*-tetraose (LST) series (**3**-**6**), disialyllacto-*N*-tetraose (DSNLT; **9**) and structures with Lewis (**7, 8, 10, 11, 14, 15**) and ABO blood group antigen motifs (**12, 13, 16**-**19**). **b** B-type glyco-epitope fragments (as oxonium ions) of the standards, *m/z* and ^DT^CCS_N2_ values (n=3 measurements). **c** ATD of the linear HMO α6-sialyllacto-N-tetraose c (obtained from human milk donor 1; n=1 measurement) and its fragment ions with their measured *m/z* and CCS values (as [M+H]^+^ or oxonium ions). Accurate *m/z* values of double-charged ions were used to determine the structural composition of the full structure and *m/z* and ^DT^CCS_N2_ values of fragment ions were used to identify the fragments. The identified fragments were used for *de novo* assembly of the full structure. The other *de novo* identified fragments (as protonated or oxonium ions) were used as new entries for the reference database.

To develop an initial reference database of *m/z* and CCS values, a collection of HMOs having common epitopes was prepared. Lacto-*N*-neotetraose (compound **1**, Fig. 2a) and lacto-*N*-tetraose (compound **2**, Fig. 2a) were used as starting materials, and a panel of glycosyltransferases was employed to install galactosides, *N*-acetylglucosaminosides, sialosides and fucosides to give Lewis and ABO blood group epitopes (Fig. 2a and Supplementary Fig. 1). The standards were derivatized at the reducing end with procainamide (ProcA)^37^ and analyzed in triplicate with drift tube (DT)IM-MS in positive ion mode. The MS spectra of intact ions showed protonated, sodiated and potassiated ions and their CCS values were calculated from their IMS arrival times using single field CCS calibration employing calibration standards with known *m/z* and CCS values (Supplementary Dataset 1)^38^. Most structures exhibited distinct *m/z* and CCS combinations for the different ion adducts, allowing the use of multiple ions to enhance identification accuracy in biological samples. This approach is particularly useful when CCS values for isomers with specific adduct ions differ by less than 1%^38^, as seen with protonated ions of **3** and **5** (Fig. 2a). The standards were also subjected to in-source CAD in positive ion mode, which provided B-type fragment ions^39^, which are oxonium ions that retain the non-reducing end, originating from glycosidic bond fragmentation only^39^ (Supplementary Data 1). The fragments showed unique combinations of *m/z* and CCS values for isomeric structures, such as for **F1-F4** (Fig. 2b) that differ in sialic acid linkage, and **F10**-**F11** (Fig. 2b) that differ in fucose linkages.

HMOs from five different human milk samples were extracted and separated into acidic or neutral HMO fractions. The HMOs were derivatized with ProcA and analyzed with liquid chromatography (LC)-IM-MS. Most of the extracted HMOs were resolved chromatographically and 14 structures could be rapidly identified by matching their CCS and *m/z* values with reference data (Supplementary Datasets 1 and 2). To identify structures without an entry in the database, the HMOs were subjected to in-source CAD in positive ion mode, and *de novo* identified. Parent ions were detected as double-charged protonated, sodiated and potassiated ions in the fragmentation spectra, while fragment ions were mostly observed as single-charged protonated or oxonium ions. The accurate masses of the intact double-charged ions revealed the monosaccharide compositions and by matching CCS and *m/z* values of fragment ions to entries in the reference database, the glycosidic linkages between monosaccharides could be determined. Full HMO structures could then be reconstructed by assembling the fragments taking into consideration well-established biosynthesis pathways^31^. For example, a compound detected in all milk samples having an *m/z* of 1218.5 is composed of glucose (Glc), *N*-acetylneuraminic acid (Neu5Ac), GlcNAc, galactose (Gal), and the ProcA label (Fig. 2c). The CCS value of the fragment ion at *m/z* 657 corresponds to fragment ion **F3** (Fig. 2b) in the reference database, revealing the presence of Neu5Ac α6-linked to a LacNAc moiety. This fragment can only be attached to lactose (**F20**) in a β3-linked fashion according to the biosynthetic rules, and therefore the full structure was identified as α6-sialyllacto-*N*-tetraose c with a ProcA label at the reducing end. The correct structural assignment was validated by matching the CCS value of the protonated parent ion with the CCS value of standard **3** (Fig. 2a). The data of the other fragment ions were added to the reference database to progressively elucidate more complex structures having the same fragment ions.

The identification of fragments in fucosylated HMOs can be challenging using only MS, as fucosyl residues can rearrange to nucleophilic moieties within the structure upon activation, which leads to incorrect fucose linkage assignment^40^. We determined the CCS value of a 3-fucosyllactose standard, with native α3-linked fucose on glucose (Supplementary Dataset 1)^41^, and compared it to isomeric fragment ions with rearranged fucose from all fucosylated standards (Fig. 2a). The 3-fucosyllactose standard exhibited a CCS of 254.6±0.1 Å^2^ (n=3), while fragments with a rearranged fucoside showed a CCS of 252.4±0.5 Å^2^ (n=36, Supplementary Dataset 1), enabling the differentiation of isomeric fragments with native and rearranged fucose and highlighting the potential of IM-MS for accurate identification of fucosylated structures.

In addition to functionalization with sialosides and fucosides, HMOs can be elongated by LNB and LacNAc moieties that determine type I (**F22**) and type II chains (**F23**, Fig. 3a), respectively. Differentiation between type I and type II chains can be achieved through the identification of terminal (**F3**, Fig. 2c) or elongated B-type fragment ions. For example, sialyl Lewis^x^ (sLe^x^) which is a sialylated and fucosylated type II structure has a CCS of 257.1 Å^2^ (Fig. 1b) whereas sialyl Lewis^a^ (sLe^a^), which is an isomeric type I structure, gives a CCS value of 244.2 Å^2^ (Fig. 1b) and thus can readily be differentiated. Other isomeric type I and II structures such as Lewis^x^ (Le^x^) *vs*. Lewis^a^ (Le^a^) also poses different CCS values. Furthermore, the CCS values of linear B-type fragment ions having two or more LacNAc moieties or LacNAc moieties extended with one terminal LNB moiety can also differentiate chain type. For example, CAD of compound **20** having a terminal LNB moiety gives B-fragment ions **F22** (*m/z* = 731.3, CCS = 262.0 Å^2^) and Y-type fragment ion **F20** (Fig. 3a). The later moiety confirms that the B-ion is linked to the reducing lactose moiety. A similar fragmentation of compound **21**, which has a terminal LacNAc moiety, gives fragment **F23** (*m/z* = 731.3, CCS = 252.2 Å^2^) and **F20**. The CCS values of **F22** and **F23** are substantially different and can differentiate between a type I or II terminal unit.

**Fig. 3.**
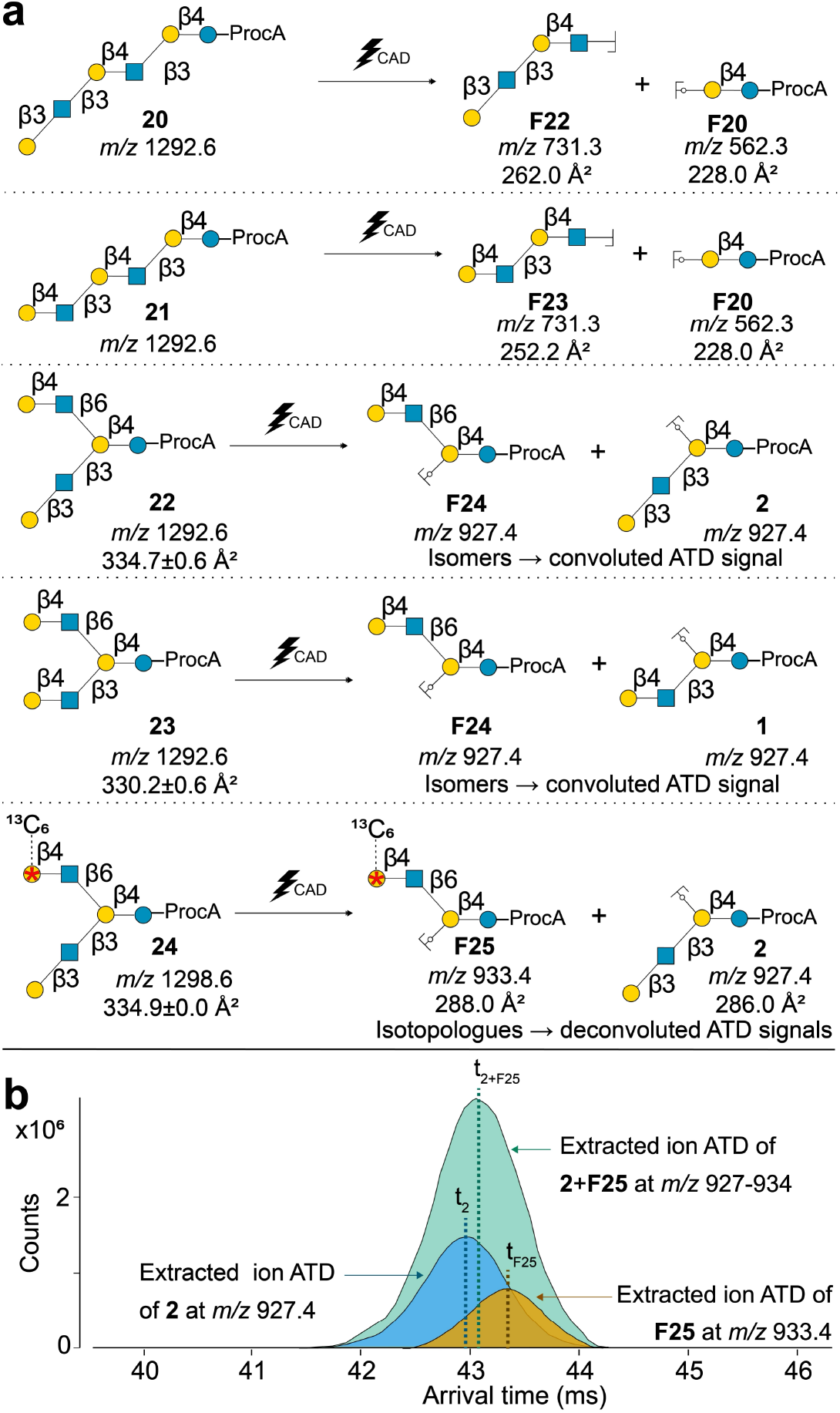
Fragmentation of linear and I-branched structures into diagnostic chain type fragment ions. **a** Linear extended structures (**20**-**21** as [M+H]^+^ ions) give complementary Y-type fragment **F20** (as [M+H]^+^ ion) and B-type fragments (**F22** and **F23** as oxonium ions) upon CAD, that can be employed to determine the chain type. When the complementary fragment ions are not present in the mass spectrum, the structure is I-branched (**22** and **23** as [M+H+]^+^ ion). I-branched structures produce isomeric fragment ions **F24** and **2** for type I (**22**), and **F24** and **1** for type II structures (**23**). The isomeric fragments have close CCS values, which results in a convoluted signal for these ions in the ATD when IMS resolution is not sufficient for their separation. The CCS values of **22** (n=5 independent measurements) and **23** (n=7 independent measurements) were determined from different HMOs extracted from milk. The isotopically labeled type II structure **24** (as [M+H]^+^ ion, n=3 measurements) produces isotopologue s **2** and **F25** (as [M+H]^+^ ions. **b** The isotopologues **2** and **F25** enable the generation of extracted ion ATDs at their respective *m/z* values, revealing minimal separation in arrival time between the isotopologue fragment ions **2** (t_2_, in blue) and **F25** (t_F25_, in orange). The isotopologues also facilitate the simulation of the signal expected if **2** and **F25** were isomers, by extracting an ATD based on the combined *m/z* values of both ions (in green). This ATD exhibits a single peak, with convoluted signals representing both **2** and **F25**, showing a weighted average arrival time (t_2+F25_) and CCS, depending on the abundances of the single fragment ions.

I-branching provides further structural diversity to HMOs and constitutes the addition of GlcNAc to the C-6 hydroxyl of an internal Gal and can occur on a type I and II chain as in compounds **22** and **23**, respectively. I-branching can generally be identified by the presence of several fragment ions derived from terminal glyco-epitopes and by the absence of linear complementary B- and Y-type fragment ions. I-branched type I chain **22** produces isomeric fragments **F24** and **2**, while type II chain **23** gives **F24** and **1** (Fig. 3a) and were expected to determine the chain type. However, HMOs from milk samples having backbone structures **22** and **23** produced a single signal in the ATD for the isomeric fragments. This is likely due to the similar arrival times and CCS of fragment ions **F24** to **1** and **2** (Fig. 3a). To study the fragmentation preferences of I-branched structures and possible co-migration of these isomeric fragments in IMS, we enzymatically synthesized an I-branched type I chain HMO having a ^13^C_6_-labeled Gal on the β6-arm and analyzed this standard by IM-MS (**24**, Fig. 3a; Supplementarry Data 1). This isotopically labeled structure indeed produced similar fragments as for **23**, but with one being heavier (**F25**, Fig. 3a). This facilitated the generation of extracted ion ATDs at their respective *m/z* values, revealing minimal separation in arrival times between the isotopologue fragments **2** (blue) and **F25** (orange, Fig. 3b). Additionally, it allowed the simulation of how **2** and **F25** would appear in the ATD if they were isomers, by extracting an ATD based on the combined *m/z* value of both ions (green, Fig. 3b). This ATD exhibited a convoluted signal representing both **2** and **F25** and showed a weighted average arrival time (t_2+F25_, Fig 3b) and thus CCS, which depended on the abundances of the individual signals. The convoluted signal of **1** and **2** (Fig. 3a) hindered chain type determination and therefore CCS values of terminal epitope and backbone fragments from I-branched HMOs (**22**-**23**, Fig. 3a) were used accordingly. Distinct CCS values were observed for backbone fragments with type I (**22**, Fig. 3a; 334.7±0.6 Å^2^, n=5) and type II chains (**23**, Fig. 3a; 330.2±0.6 Å^2^, n=7 independent measurements) in I-branched structures from milk samples, with the type I chain values validated by standard **24** (Fig 3a; 334.9±0.0 Å^2^, n=3 independent measurments).

Based on the gathered information, a structure with degree of polymerization of 9 (DP9) was *de novo* sequenced as shown in Fig. 4. The composition was determined from the accurate mass of potassium and sodium adducted ions containing three Gal, one Glc, two GlcNAc, two Fuc, and one Neu5Ac moiety and a ProcA label (step 1). A Glc, Gal and GlcNAc moiety are linked according to the well-established HMO biosynthesis forming a core trisaccharide (GlcNAc-β(1,3-)Galβ(1,4)-Glc) and the ProcA label is attached to the reducing end (step 1). In step 2 and 3, the fragments at *m/z* 657 and 658 were identified as **F3** and **F11** (Fig. 2b), respectively, based on their CCS values. These fragments are terminal epitopes and therefore should occur on different arms indicating an I-branched structure. I-branching was further confirmed by the absence of complementary extended B-type fragment ions along with the Y-type lactose fragment ion (step 4). The previously identified compound **3** (Fig. 2c) is a fragment ion in this structure (step 5) revealing a type II chain having a β3-linkage of **F3** to Gal of lactose. The GlcNAc moiety, which is part of **F11**, is β6-linked to Gal of lactose based on biosynthetic rules of I-branched structures, and thus **F11** must be attached to the β6-arm revealing compound **25** by *de novo* sequencing (Fig. 4). The identification of the structure of the β6-arm as fragment ion **F26** (step 6), provided a new entry for the reference database facilitating the identification of other I-branched structures having a Le^y^ motif at the β6-arm.

**Fig. 4.**
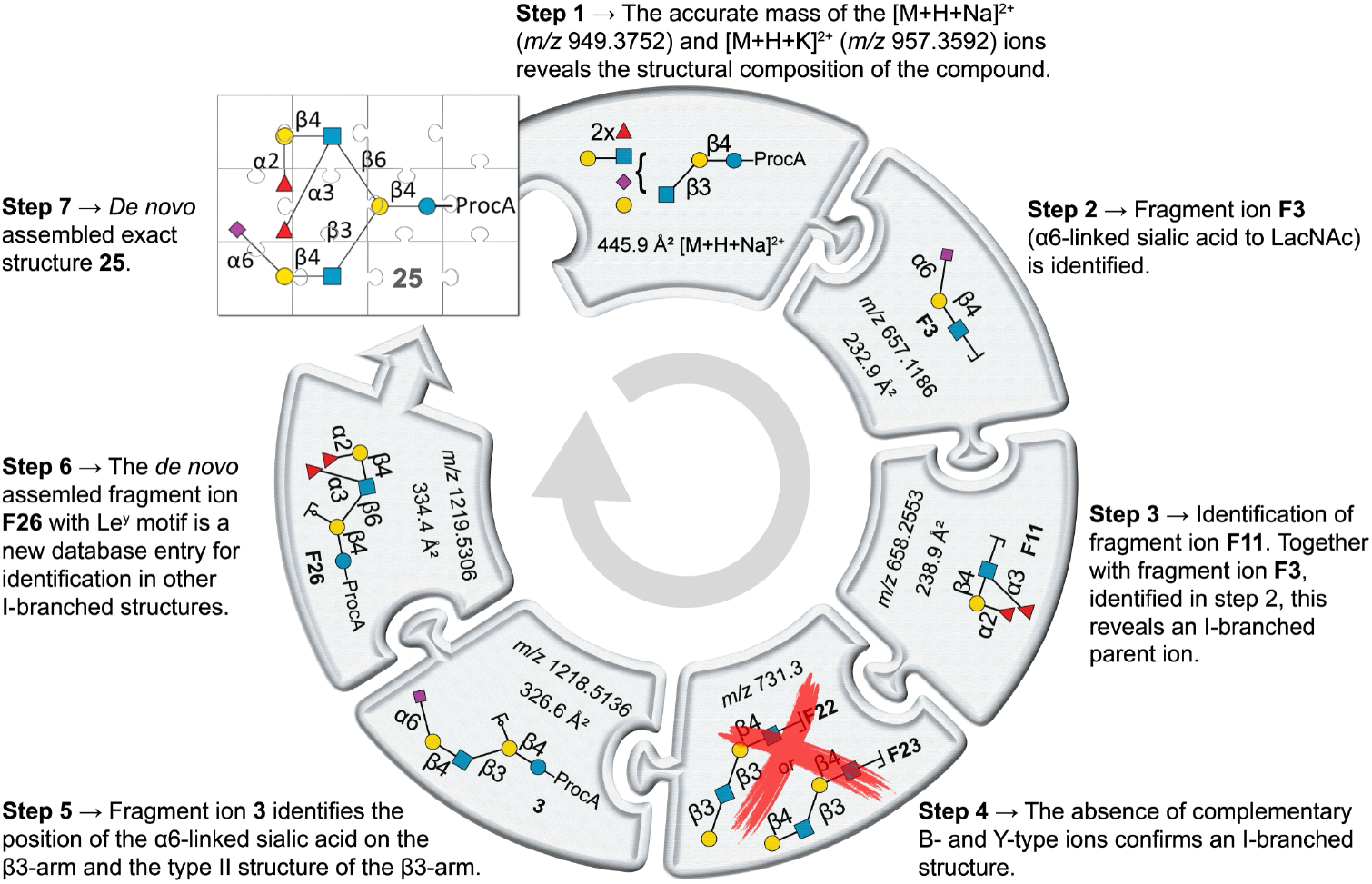
*De novo* sequencing cycle of an I-branched structure derived from human milk. (n=1 measurement). Double-charged ions were used to determine the structural composition (step 1). The identification of two terminal epitope fragment ions (**F3** and **F11** as oxonium ions) revealed a branched structure (step 2 and 3), which was confirmed by the absence of complementary extended B-type ions along with the Y-type lactose core fragment ion (step 4). The linkage of **F3** to the lactose core was determined by the identification of **3** as fragment ion (as [M+H]^+^ ion; step 5) and by *de novo* assembly of the fragments, compound **25** was identified (step 7). The *de novo* identified fragment ion **F26** (as [M+H]^+^ ion) was added to the reference database for identification in other samples (step 6).

By analyzing the five donor milk samples (Supplementary Fig. 2-141), 186 new entries for the reference database were obtained (Supplementary Data 3). Exact structures of the 43 most abundant HMOs, having a complexity up to DP9 and encompassing both linear and I-branched structures with various terminal epitope motifs, were assigned. (Table 1). The assignment of several HMOs with type II structures was validated by the analysis of synthetic standards^41^ (Supplementary Fig. 142-145) and comparison of their CCS values. Interestingly, for one of the identified compositions, comprising of Lacto-*N*-triose, two Gal, one *N*-GlcNAc, one Fuc and one Neu5Ac, in principle 36 different isomers are possible (Extended Data Fig. 1). However, only one of these structures was detected in all donor samples and only three other isomers in one donor sample (Table 1). This observation indicates that the glycosyltransferases involved in the biosynthesis of HMOs have more intricate specificities and fine structural details determine acceptor capabilities.

**Table 1.**
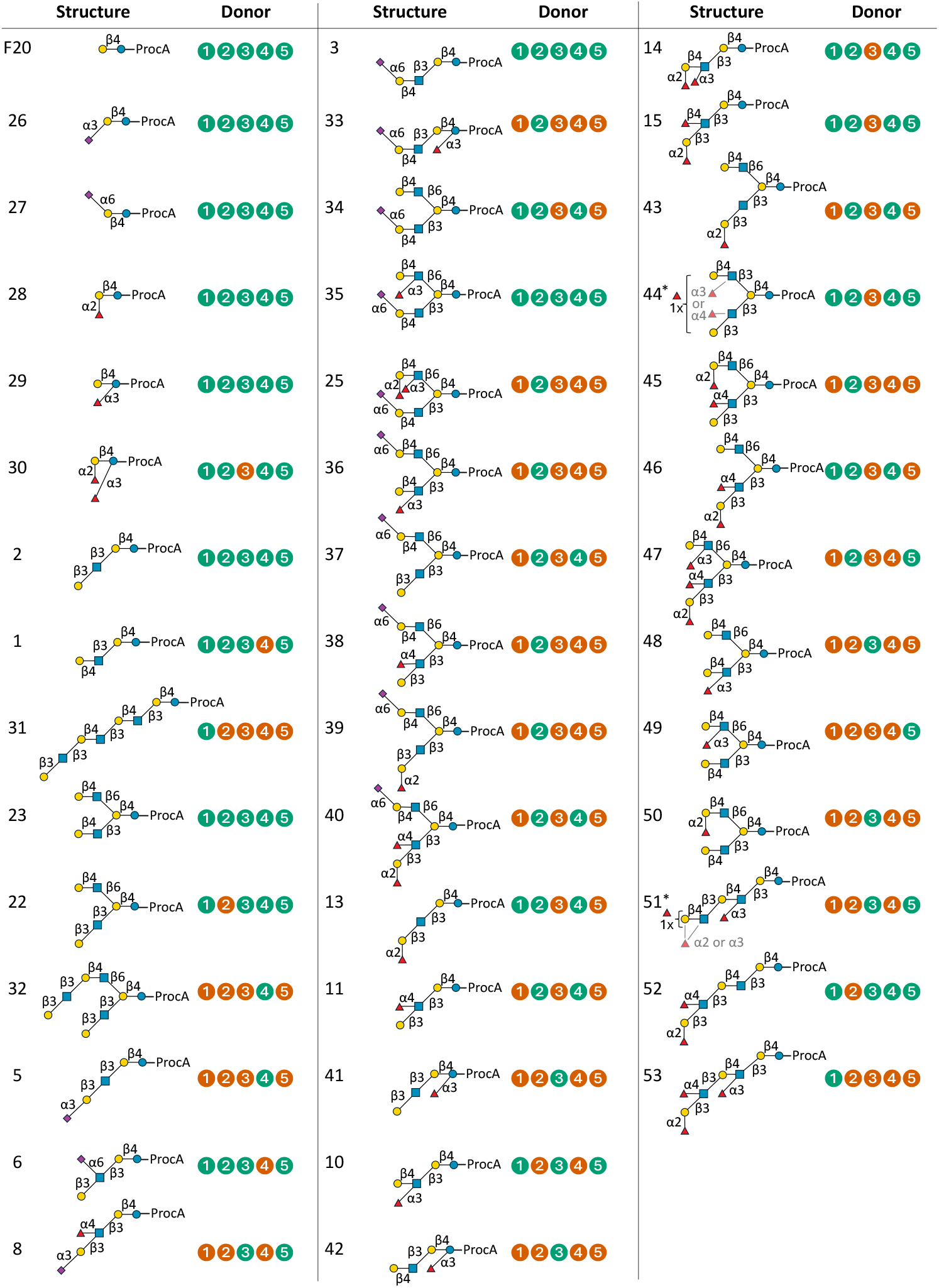
*De novo* identified HMO structures in the milk samples of donors 1-5. The HMO structures identified in a donor sample are marked in green, while a structure that was not detected in the respective donor sample is marked in red.

Individual genetics of mothers play an important role in the variability of HMO structures, specifically by the expression of the fucosyltransferases FUT2 and FUT3^42^. Milk from secretor mothers (Se+), for example, is characterized by high levels of α2-fucosylated HMOs, such as 2’-fucosyllactose and LNFP I. Non-Secretor mothers (Se-) lack a functional FUT2 enzyme and therefore have less α2-fucosylated HMOs in their milk. Additionally, mothers classified as Le positive (Le+) express the FUT3 enzyme, which facilitates the addition of α4-linked Fuc to a GlcNAc residue. Conversely, milk from Le negative mothers (Le-) contains less α4-fucosylated HMOs, such as LNFP II^31^. Based on the elucidated structures, the five donors could be classified according to their Se and Le status into one of the four milk groups: Se+/Le-, Se-/Le-, Se+/Le+, and Se-/Le+^43^. Donor three was classified in milk group four (Se-/Le+), while the other donors were classified in milk group three (Se+/Le+). These classifications can provide insights into the diverse composition of human milk that impacts infant growth, development and overall health^31, 42^.

### Identification of exact structures of N-glycans

Next, attention was focused on the identification of *N*-glycans, which have a common pentasaccharide core consisting of Man(α1-3)[Man(β1-6)]Man(α1-4)*N*-GlcNAc(β1-4)β-*N*-GlcNAc that can be further branched and extended with additional LacNAc moieties^44^. The resulting multi-antennae structures can be sialylated and fucosylated, generating terminal epitopes similar to those found on HMOs. Recently, we demonstrated that *N*-glycans can adopt several stable conformations in the gas phase, which can be separated in the IMS dimension and used for exact structure identification by comparing ATD fingerprints with reference database entries^18^. Here, a further collection of 49 chemoenzymatically synthesized complex *N*-glycan standards, including many isomeric structures^18, 45, 46, 47^, was subjected to IM-MS to expand the reference database with high resolution (HR-)ATDs (Extended Data Fig. 2). The standards include bi- and tri-antennary glycans having various numbers of LacNAc extensions modified with different linkage types of Neu5Ac and Fuc. The standards were derivatized with 2-aminobenzoic acid (2AA) by reductive amination and analyzed in triplicate by IM-MS in positive and negative ion mode with nitrogen as buffer gas. The HR-ATDs of the glycan standards showed unique fingerprints for each glycan (**54, 55, 57** and **58**, Fig. 5a; Supplementary Fig. 147-152). The distributions are predominantly multimodal, having on average around three conformers in the ATDs for deprotonated, protonated, protonated-sodium and protonated-potassium adduct ions and were unique for all isomeric structures. The distributions were reproducible and not affected by eluent composition in LC-IM-MS experiments and employed electrospray source (standard electrospray ionization source or the Jet Stream interface with superheated nitrogen; Supplementary Fig. 153-159), other than minor changes in peak intensities^18^.

**Fig. 5.**
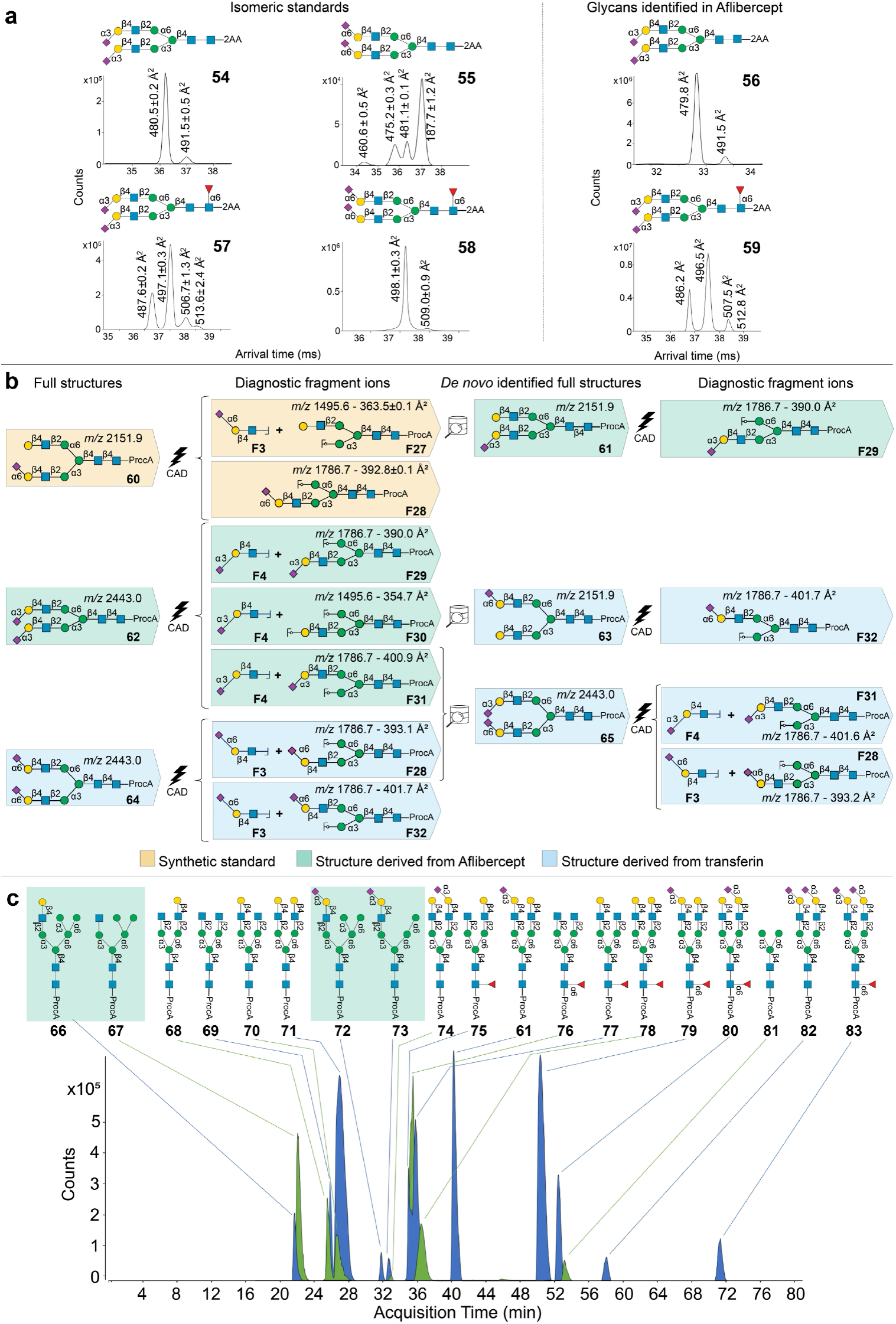
ATD fingerprinting identification and *de novo* sequencing of glycans derived from biological samples. **a** HR-ATDs with ^DT^CCS_N2_ values (n=3) of disialylated *N*-glycan standards **54-55, 57-58** with different sialic acid linkages and core fucosylation, measured as [M-2H]^2−^ ions. HR-ATDs with ^DT^CCS_N2_ values (n=1) of identified *N*-glycans from Aflibercept measured as [M-2H]^2−^ ions, with α2,3-linked sialic acids and core fucosylation (**56** and **59**). **b** *De novo* identification route of *N*-glycans (as [M+H]^+^ ions) after CAD. Standard **60** produced the diagnostic primary fragment ions **F27** and **F28**, which allowed the identification of structure **61** from Aflibercept. Structure **62** from Aflibercept produced additional diagnostic primary fragment ions **F29-F31**. Structure **64** from transferrin produced diagnostic primary fragment ions **F28**-**F32**. The diagnostic fragment ions **F27**-**F32** can be used to either directly identify all non-, mono- and disialylated structures of biantennary glycans, such as **65**, or reveal new pathways to identify further elongated and branched *N*-glycans.

The ATDs, CCS and *m/z* values of the standards were employed for the identification of *N*-glycans released from Aflibercept, which is an Fc fusion protein of the extracellular domain of the human VEGF receptor^48^, which is used for the treatment of macular degeneration^49^ and metastatic colorectal cancer. The *N*-glycans were released with PNGase F, derivatized with 2AA and analyzed by LC-IM-MS. By matching *m/z* and ATD fingerprints with those in the reference database, six structures could be identified (Supplementary Fig. 160). This includes glycans having a core fucoside and α2,3-linked Neu5Ac (compounds **56** and **59**, Fig. 5a) which is consistent with production in Chinese hamster ovary cells that lack α2,6-sialyltransferase expression^50^.

*De novo* structure identification was performed for glycans without an entry in the reference database. For this purpose, PNGase F-released glycans were derivatized at the reducing end with ProcA, which offers greater sensitivity in positive ion mode compared to aromatic amines such as 2AA^51^, while providing glycosidic bond fragmentation only. Compositions of 19 *N*-glycans from Aflibercept could be determined by accurate mass measurement of the sodiated and potassiated doubly charged ions, which did not fragment at the applied activation energy. From the compositions, it could be established that all structures have a biantennary architecture, including hybrid and complex type glycans. Hybrid type structures have mono-, di- or tri-mannosyl residues at the α6-linked arm and a complex epitope at the α3-arm^52^. Therefore, exact structures of these compounds (**66, 67, 72**, and **73** Fig. 5c), could be identified from *m/z* and CCS values of previously identified fragment ions (Supplementary Fig. 163-167). For example, fragment ion **F3** (Fig. 5b) demonstrated the presence of an α2,3-sialyl LacNAc moiety at the α3-mannosyl arm in compounds **72** and **73** (Fig. 5c).

To distinguish complex isomeric *N*-glycans having a unique epitope at each antenna, we analyzed standard **60** (Fig. 5b). It has an α2,6-linked Neu5Ac at the α3-mannosyl antenna which upon activation produces the diagnostic primary fragment ions **F27** and **F28** (Fig. 5b and Supplementary Fig. 161). The structure of disialylated *N*-glycan **62** released from Aflibercept could readily be determined from its composition and the presence of fragment ion **F4** (Fig. 5b and Supplementary Fig. 180). Upon activation, it produced multiple pairs of isomeric diagnostic fragment ions, including secondary fragment ion **F30** (Fig. 5b). This fragment is isobaric to **F27** but has a substantially different CCS value (354.7 Å^2^ *vs*. 364.8 Å^2^) and therefore its structure could readily be identified as only one alternative isomer is possible. In a similar way, the primary fragment ions **F29** and **F31** were identified by using fragment ions of **61** as a reference (Fig. 5b). The structure of monosialylated **61** and **74** (Fig. 5c) could be determined from previously determined diagnostic fragment ions **F27** and **F30**, respectively. The linkage type of Neu5Ac was determined by matching the corresponding fragment ion with **F4** (Fig. 2b, Fig. 5b and Supplementary Fig. 176-177). No isomers are possible for *N*-glycans **69** and **81** (Fig. 5c), allowing structural identification based on their fragment ions (Supplementary Figs. 163 and 168). The isomeric structures **68** and **70-71** (Fig. 5c) were identified by the presence of diagnostic fragment ions **F27** and **F30** (Fig. 5b). Following a similar approach, diagnostic fragment ions for core-fucosylated *N*-glycans **75**–**80** and **83** (Fig. 5c) were obtained through the identification of **F27, F30** and **F4** (Fig. 5b) in their fragmentation spectra (Supplementary Figs. 172–173, 175, 178–179, and 181).

The new entries enable the identification of glycans with longer antennae and having various terminal epitopes. The database of CCS and *m/z* values was, for example, applied for the identification of isomeric glycans released from transferrin^53^. The glycans of transferrin are employed as biomarkers for various diseases and more accurate assignment of structures will increase prognostic and diagnostic value^54^. It made it possible to identify isomeric bis-sialylated *N*-glycans having different combinations of 2,3- and 2,6-linked sialosides. In this respect, compound **64** (Fig. 5b) could easily be identified using fragment ion **F3** (Fig. 2b). Upon activation, this structure produced diagnostic fragments **F28** and **F32** (Fig. 5b), which were identified using fragments of **60** (Fig. 5b) as reference for comparative analysis (Supplementary Fig. 175-178). Fragment ions **F3** and **F4** (Fig. 2b) demonstrated that di-sialylated glycan **65** is modified by an α3- and an α6-linked Neu5Ac. Fragments **F28** and **F31** (Fig. 5b) established that the bottom arm of **65** is modified by α2,6-linked Neu5Ac revealing the structure of this glycan. The structural assignment of this compound was validated by comparing the ATDs and CCS values of the released glycan (Supplementary Fig. 223), labeled with 2AA at the reducing end, with those of a previously reported standard^18^. The isomeric fragments **F27**-**F32** and their core-fucosylated glycoforms can be used to either directly identify all non-, mono- and disialylated structures of biantennary glycans or to uncover new pathways for identifying more extended *N*-glycans in biological samples.

Many common fragment ions were identified in the mass fragmentation spectra of the different *N*-glycans, which facilitated validation of assigned fragment ions and full structures. The assignments were also validated by matching ATD fingerprints and CCS values of identified structures from proteins with a library of ProcA-derivatized standards. Additionally, these standards were also used to demonstrate that ATDs of *N*-glycans are minimally affected by variations in the energy applied during CAD (Supplementary Figs. 188-222).

## Discussion

The identification of exact structures of glycans in complex biological samples is fundamentally a problem of isomerism that cannot readily be addressed by MS alone^55^. The hyphenation of MS with IMS provides a promising means to distinguish isomeric glycans. Glycans form stable gas-phase conformations during electrospray ionization, giving an intrinsic CCS value for small structures, while larger structures can have multiple stable conformations providing an intrinsic ATD fingerprint. Exact structures of glycans in biological samples can be determined by comparing *m/z* and CCS values or ATD patterns with those in a reference database.^27^ Such a database can be established by subjecting synthetic and well-defined glycan standards to IM-MS. This approach has, for example, been used to determine *O*-acetylation patterns and anomeric linkages of sialosides of *O*- and *N*-linked glycans released from tissues and mucins^27^. Despite the attraction of this approach, it is not possible to prepare all possible naturally occurring glycans due to the enormous structural diversity^28^. To address this challenge, we have developed an IM-MS method for rapid glycan identification and *de novo* sequencing based on a self-expanding reference database of CCS values and ATD patterns. It is based on the use of a limited number of synthetic standards that is subjected to fragmentation to create an initial database of CCS or ATDs and corresponding *m/z* values of molecule or fragment ions. Compounds that do not have an entry are subjected to *de novo* sequencing by identifying structures of fragment ions that can be puzzled together leveraging knowledge of glycan biosynthetic pathways. The identification of CCS and ATDs of additional fragments result in a self-expanding reference database, gradually facilitating the sequencing of HMOs and *N*-glycans of increasing complexity. In this study, we established a reference database from an initial 20 glycan standards to 332 unique entries. It is the expectation that the methodology can be extended to other classes of glycans and glycoconjugates. For example, there are 8 common *O*-linked glycoprotein cores, some of which are isomeric, and it is the expectation that the use of a limited number of synthetic standards can provide CCS values of diagnostic fragment ions to identify core type. Furthermore, the different classes of glycosphingolipids have isomeric core structures that are expected to be distinguishable based on CCS values of fragment ions. *O*-glycans and glycosphingolipids have extensions like those of HMOs, and thus the current database should make it possible to establish structural elements of these compound classes.

Several types of IMS, including travelling wave (TWIMS), trapped (TIMS) and DTIMS, can separate isomeric glycans^56, 57, 58, 59^ and it is the expectation that the *de-novo* sequencing approach can be employed across different instrument setups. HR-IMS instruments may separate isomeric ions that remain unresolved in conventional DTIMS, further broadening the scope of the *de novo* sequencing methodology. In addition, combination with selective fragmentation of precursor ions prior to IMS separation will enable targeted structural analysis of glycans.

The IM-MS methodology will empower the glyco-research field with the ability to rapidly and precisely identify complex glycans in complex biological samples. This knowledge is critical to correlate glycan structures with biological functions and will for example make it possible to more accurately characterize biopharmaceuticals that usually are glycosylated. It will accelerate the development of diagnostics with greater prognostic or diagnostic value. Also, it will make it possible to develop next generation nutraceuticals such as infant formula with tailor made compositions. Accurate characterization of glycan structures is expected to uncover new insight in the biosynthesis of these compounds. Here, we show that of the 43 possible isomers of a DP9 HMO, only four are observed, indicating that ensuing glycosyl transferases have more complex acceptor requirements than is currently known.

## Methods

### Materials and reagents

Sodium acetate, sodium cyanoborohydride, tris(hydroxymethyl)aminomethane (TRIS), acetic acid (LC-MS grade), ammonium hydroxide (LC-MS grade), formic acid (LC-MS grade), sodium acetate, DEAE Sepharose Fast Flow ion exchange resin, ammonium bicarbonate, sulfuric acid, ethanol, trifluoroacetic acid (TFA), dimethylsulfoxide (DMSO), dichloromethane (DCM), 2-aminobenzoic acid (2AA), procainamide HCl (ProcA) and ammonium acetate were acquired from Merck (Darmstadt HE, Germany). Hypercarb 25 and 200 mg porous graphitized carbon (PGC) solid phase extraction (SPE) cartridges were obtained from Thermo Fisher Scientific (Waltham MA). Acetonitrile (LC-MS grade) was purchased from Biosolve B.V. (Valkenswaard, The Netherlands). Acetone (LC-MS grade) was obtained from Boom (Meppel, The Netherlands). Ultrapure water was produced by a Synergy UV water purification system from Merck-Millipore (Burlington MA). Peptide:N-glycosidase F (PNGaseF) and glycosyltransferases were produced in-house. UDP-Gal, UDP-*N*-acetylglucosamine, UDP-*N*-acetylgalactosamine and CMP-Neu5Ac were purchased from Roche Diagnostics (Basel, Switserland). Calf intestine alkaline phosphatase (CIAP) was obtained from Thermo Fisher Scientific (Waltham MA). GDP-fucose was prepared using L-fucokinase/GDP-fucose pyrophosphorylase. Lacto-*N*-tetraose was a gift from Friesland Campina (Amersfoort, The Netherlands) and purified by preparative LC before use. Endoglycosidase F2 (Endo-F2) was purchased from New England Biolabs (Ipswich MA). D_2_O (99.9%) was purchased from Buchem (Apeldoorn, The Netherlands). Avantor C_18_ 1ml SPE cartridges (100 mg) were acquired from Baker (Radnor PA). Minitrap G-10 cartridges were obtained from GE Healthcare (Chicago IL) and Vivaspin 20 10kD filters were acquired from Sartorius (Götingen NI, Germany). Bio-Gel P2 gel polyacrylamide beads were acquired from Bio-Rad (Hercules CA). Human milk was purchased from Medix Biochemica (Epoo, Finland).

### ProcA and 2AA labelling

Labels were attached to free reducing end glycans via reductive amination. A 120 μL volume of glycan standards or released glycans in water was mixed with 40 μL labelling solution (30 mg/mL ProcA or 2AA and 75 mg/mL sodium cyanoborohydride in DMSO) and 20 μL acetic acid. The mixture was vortexed and incubated at 68 °C for 2 hours, unless specified otherwise. The reaction mix was evaporated under a nitrogen flow and the residue was dissolved in 1 mL water for PGC SPE purification.

### PGC SPE purification

Solutions of released or derivatized glycans were purified by PGC SPE with a 25 mg cartridge. The cartridge was conditioned with 1 mL ACN followed by 1 mL water. The sample was loaded on the cartridge and washed with 1 mL 0.05% TFA, followed by 1 mL ACN/water/TFA (5%/94.95%/0.05%).

Glycans were eluted with 1 mL ACN/0.1% TFA (50%/50%). The eluent was evaporated under a nitrogen stream and the sample was redissolved in water for LC-DTIMS-MS analysis.

### LC-DTIMS-MS

The LC-DTIMS-MS system consisted of an Agilent Technologies (Santa Clara, CA) Infinity LC system with a 1290 Binary pump, a 1290 sampler and a 1260 thermostatted column compartment. The LC system was coupled via a jet stream interface to an Agilent Technologies 6560 Ion Mobility Q-TOF LC/MS system. The DTIMS-MS was used in both negative and positive ion mode. The MS source parameters used were: capillary voltage of 3500 V, nozzle voltage of 1000 V, nebulizer pressure of 40 psi, N_2_ drying gas heated to 300 °C at 8 L/min and sheath gas heated to 350 °C at 11 L/min. The ion mobility measurements were conducted using an DTIMS transient rate of 16 transients/frame, a trap fill time of 3900 μs, a trap release time of 250 μs, a drift tube entrance voltage of 1700 V for positive mode and −1700 V for negative ion mode (unless specified otherwise), a drift tube exit voltage of 250 V for positive and −250 V for negative ion mode, nitrogen as drift tube gas type, a drift tube pressure of 3.95 Torr, a trap funnel pressure of 3.80 Torr and a multiplexing pulsing sequence length of 4 bit. For in-source activation of HMO structures a fragmentor voltage of 500 V was used. For *N*-glycans, a fragmentor voltage of 550 V was applied for fragmentation and 360 V for the analysis of intact structures.

### Analysis of *N*-glycan reference standards derivatized with 2AA

Glycan standards were labelled with 2AA at the reducing end and analyzed with hydrophilic interaction liquid chromatography (HILIC) with a 20×2.1 mm (5μm) Merck SeQuant ZIC HILIC column (Burlington, MA) using 1 μL injection volumes of 1 μg/μL solutions with an eluent composition of ACN/water with 0.1% formic acid for positive ion mode and ACN/10 mM ammonium acetate (pH 4.4) for negative ion mode MS, at a flow rate of 0.2 mL/min. The eluent compositions were adjusted for each compound to ensure proper retention on the chromatographic column.

### Analysis of *N*-glycans from Aflibercept

Aflibercept solution of 1 mg/mL was loaded on a 10 kDa spinfilter and centrifuged till near dryness at 4500 RPM. A 4x 1-ml volume of 100 mM sodium acetate (pH 4.5) was added, and the sample was centrifuged again (4500 RPM leaving at least 100 μl solution). *N*-glycans were released enzymatically from the biological Aflibercept by PNGaseF, derivatized with 2AA or ProcA and further purified by PGC SPE before LC-DTIMS-MS analysis.

2AA-derivatized *N*-glycans were analyzed on a 150 × 2.1 mm (3 μm) Merck Sequant ZIC-HILIC column using a gradient with 10 mM ammonium acetate (pH 4.4; A) and ACN (B, starting at 20% A for 5 min and increasing to 50 % A in 25 min at a flow rate of 1 mL/min. The column was maintained at 40 °C during analysis and 15 μL of sample was injected for analysis. A mass range of *m/z* 100-3200 was used for DTIMS-MS measurements.

For the analysis of ProcA-derivatized glycans, 15 μL sample was injected onto a 150 × 4.6 mm (3 μm) Thermo Fischer Scientific Hypercarb PGC column, which was maintained at 60 °C during analysis, and separated with a gradient using 0.1% formic acid (A) and ACN (B) at a flow rate of 1 mL/min. The gradient started at 25% B for 5 min and increased to 35% in 100 min. DTIMS-MS in positive ion mode used a fragmentor voltage of 550 V and a mass range of *m/z* 100-3200.

### NMR characterization of HMO structures

Standards were dissolved in D_2_O and their ^1^H NMR spectra were recorded on a Bruker 600 MHz Advance Neo NMR spectrometer (Billerica MA). NMR Chemical shifts were recorded in parts per million (ppm) relative to the residual H_2_O (δ 4.790) peak. NMR signals were represented as: chemical shift and multiplicity (d = doublet, t = triplet, q = quadruplet, dd = doublet of doublets, app. = apparent) and J coupling if applicable. Assignment of ^1^H NMR signals was conducted based on ^1^H NMR, 2D COSY (COSYGPSW), 2D TOCSY (MLEVPHSW), 2D NOESY (NOESYPHSW) and 2D HSQC (HSQCEDETGPSISP). ^13^C NMR signal assignment was extracted from 2D HSQC spectra.

### Analysis of acidic and neutral HMOs in human milk

Human breast milk samples from 5 individuals were analyzed. A volume of 1 mL milk from each individual was centrifuged at 4824 x *g* for 10 minutes. The solution was frozen to solidify the fat layer and the supernatant was transferred to a clean tube. This process was repeated 3 times to ensure the removal of fat. Subsequently, the samples were loaded on 10 kDa spinfilters and centrifuged at 2415 x *g* for 20 minutes to remove high molecular weight compounds. The filtrate was freeze dried and the residue was redissolved in 2 mL water for purification by SPE using a 200 mg PGC cartridge. The cartridge was conditioned with 2 mL 90% ACN/10% water, followed by 2 mL 0.05% TFA in water. The sample was loaded on the cartridge and washed with 2 mL 0.1% TFA in water, followed by 2 mL ACN/0.05% TFA in water (5%/95%, v/v). The HMOs were eluted with 2 mL ACN/0.05% TFA in water (50%/50%, v/v). The eluate was dried under a nitrogen flow and the residue was redissolved in water. The HMO sample solution was loaded on a DEAE Sepharose Fast Flow ion exchange column with a 3 mL bed volume. The neutral HMOs were eluted with 6 ml water and subsequently the acidic HMOs were eluted with 1M ammonium bicarbonate. Alle fractions were freeze dried. Subsequently, the acidic HMO fractions were dissolved in 3 mL water and the neutral HMO fractions in 1 ml water. The HMOs in these solutions were derivatized via reductive amination with ProcA by taking 500 μL of each sample solution and adding 100 μL acetic acid and 300 μL labeling solution, containing 30 mg/mL procainamide and 75 mg/mL cyanoborohydrate in DMSO. The mixture was incubated at room temperature overnight and then evaporated under a nitrogen flow. The residue was redissolved in 1 ml water and purified using 200 mg PGC SPE cartridges as described above. The eluate was freeze dried and reconstituted in 300 μL water for PGC LC-DTIMS-MS.

A sample of 5 μL was injected onto a 150 × 4.6 mm Thermo Scientific Hypercarb PGC column with 3 μm particles, which was maintained at 60 °C during analysis, and separated with a gradient using 0.1% formic acid (A) and ACN (B) at a flow rate of 0.5 mL/min. The gradient started at 20% B for 5 min and increased to 40% in 90 min and then to 100% in 10 min and maintained at 100 % for 15 min. DTIMS-MS was used in positive ion mode and used a drift tube entrance voltage of 1700 V, a fragmentor voltage of 500 V and a mass range of m/z 100-3200.

### DTIMS-MS data processing

Masses of raw 4 bit multiplexed DTIMS-MS data were recalibrated on reference masses with *m/z* 121 and *m/z* 922 using the DTIMS-MS Data File Reprocessing Utility in the Agilent Technologies MassHunter software (v10.0). Reprocessed data was demultiplexed using the PNNL Preprocessor software v4.0 (Pacific Northwest National Laboratory, Richland WA) using an interpolation of 3 drift bins and a 5 point moving average smoothing in the DTIMS dimension^60^. Features were identified with the Agilent Technologies MassHunter IMS Browser software v10.0 using an unbiased isotope model. High-resolution ATDs were obtained using Agilent Technologies HRdm v2.0 software with an *m/z* width multiplier of 12, saturation check of 0.40 and an IF multiplier of 1.125 with SSS and Post QC enabled^61^.

## Data availability

The authors declare that the data supporting the findings of this study are available within the paper and its supplementary information files. The ion mobility-mass spectrometric source data that support the findings of this study are available in MassIVE with the identifier MSV000097548 (doi:10.25345/C5MS3KD50; https://massive.ucsd.edu/ProteoSAFe/dataset.jsp?task=9b8b516b30ab449a9c7d12bffacc64c6).

## Acknowledgments

This research was supported by the Agilent Technologies Applications and Core Technology University Research grant ACT-UR-4725 awarded to J.S.T., the Dutch Top Sector Life Sciences & Health (LSH-TKI) grant LSHM21030 awarded to G.J.B., and is part of the project Technology for Dissecting Carbohydrates in Food, Biopharma, and Biomedicine (DISC) awarded to G.J.B with file number KICH1.ST01.20.014 of the research program KIC Key Technologies, which is (partly) financed by the Dutch Research Council (NWO).

## Author contributions

J.S.T., J.V., G.M.V., K.H. and S.V. performed the experiments. J.S.T., J.V., K.H., S.V., J.F. and C.K. analyzed the data and J.S.T., J.V., K.H., S.V. and B.S. interpreted the results. J.S.T., J.V. and G.J.B. wrote the manuscript. J.S.T. and G.J.B supervised the research project. All authors have given approval to the final version of the manuscript.

## Author identification

