## Extended Data Figs 1 & 2 for "*De novo* sequencing of complex glycans by ion mobility-mass spectrometry using a self-expanding reference database"

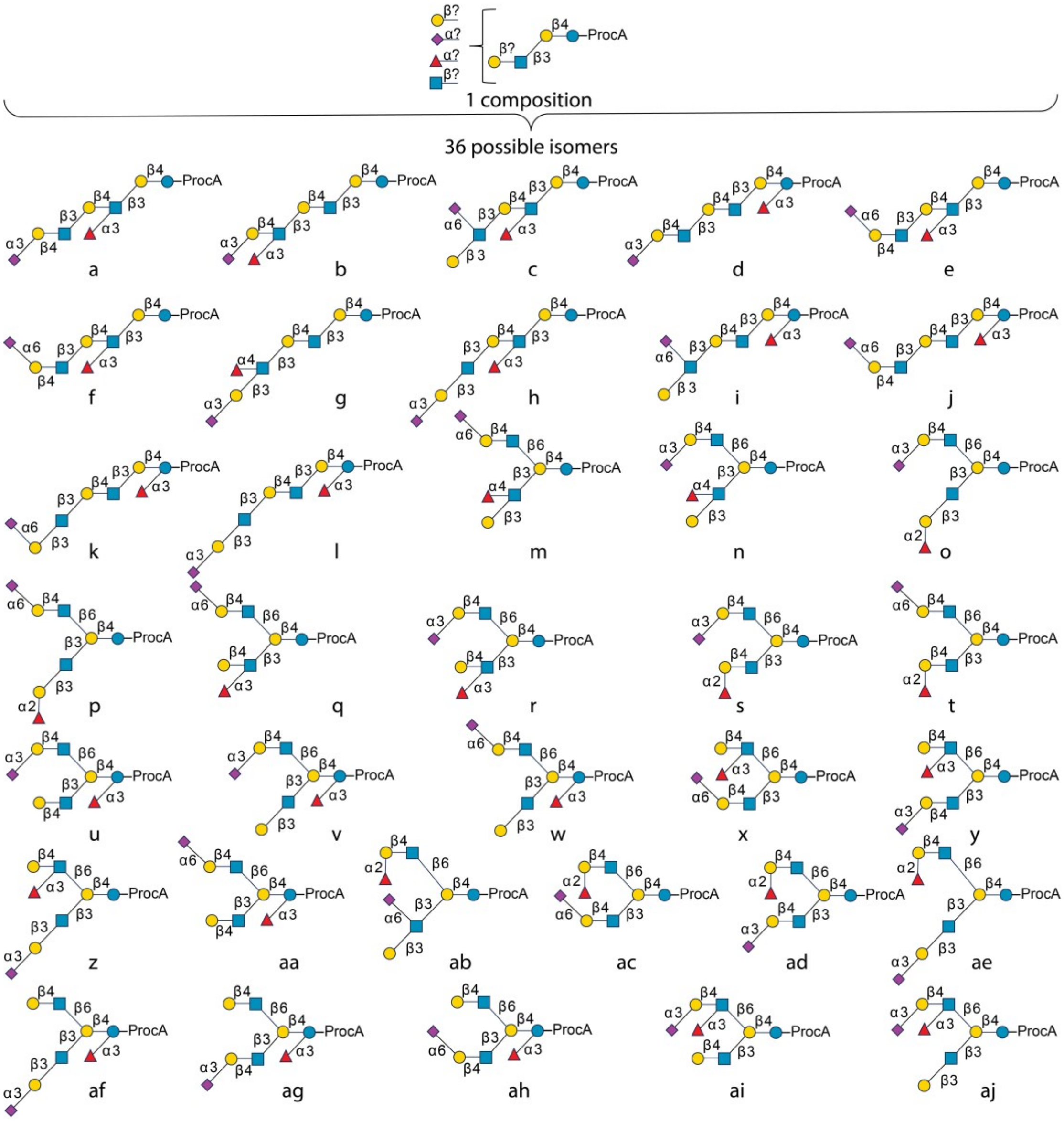


**Extended Data Fig. 1. A number of 36 different isomers are possible for an HMO with a composition consisting of 8 carbohydrate units.**

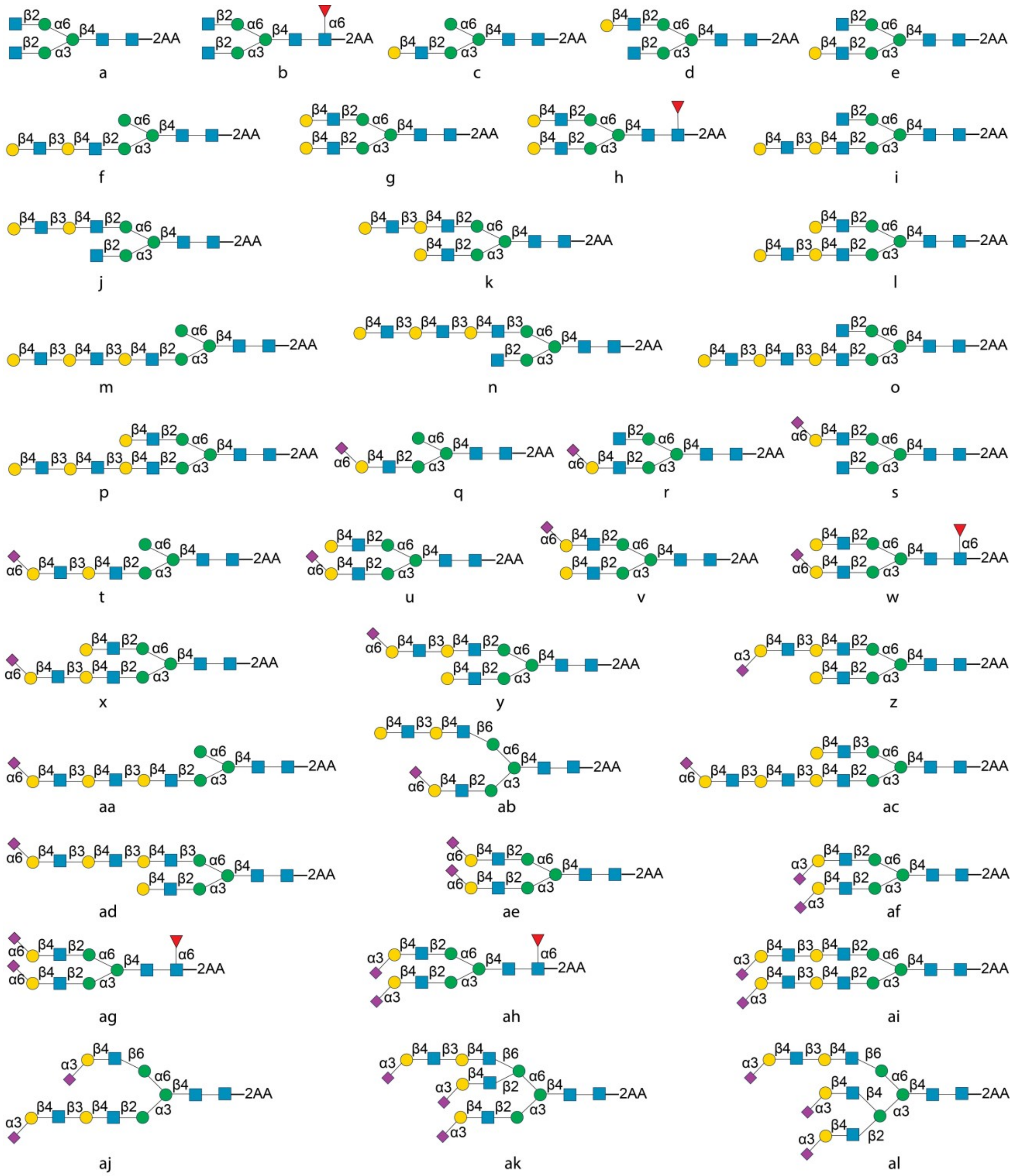


**Extended Data Fig. 2. Library of *N*-glycan standards subjected to IM-MS to expand the reference database with high resolution (HR-)ATDs.**
