## Supplementary Information for "*De novo* sequencing of complex glycans by ion mobility-mass spectrometry using a self-expanding reference database"

### 1. Synthesis of HMO structures

A collection of HMOs was prepared by exploiting an in-house generated library of glycosyltransferases that can install sialic acids and Lewis as well as ABO blood group epitopes. Glycan cores were generated by the consecutive actions of N-acetylglucosamine transferase B3GnT2 and Gal transferases B3GalT5 and B4GalT1 on lactose. Lewis- and blood group antigens were installed by the action of appropriate glycosyl transferases (Supplemental Information). The reaction progresses were determined by LC-MS with a SeQuant Zic HILIC column (20x2.1 mm, 5µm) using a gradient of ACN and water from 90% to 50% ACN over 5 min, followed by a 50% ACN isocratic elution. Detection was performed on a Bruker microTOF-QII MS (Billerica, MA). After each synthesis step the product was purified by size exclusion chromatography using Bio-Rad Bio-Gel P2 gel polyacrylamide beads by elution with water. Silica gel TLC was used to locate the fractions containing glycan compounds. The glycans were visualized by dipping the TLC plate in a 10% sulfuric acid solution in ethanol followed by charring. After size exclusion chromatography, preparative LC was conducted on a Shimadzu Prominence HPLC system (Kyoto, Japan), consisting of two LC-20AT pumps, a SIL-20A injector, a CBM-20A controller, an SPD-20AV UV detector and a FRC-10A fraction collector, connected to a Bruker micrOTOF MS. Separation was achieved on a 10x250 mm (5 µm) Waters Xbridge amide BEH prep column (Milford, MA). The final product was characterized by NMR.

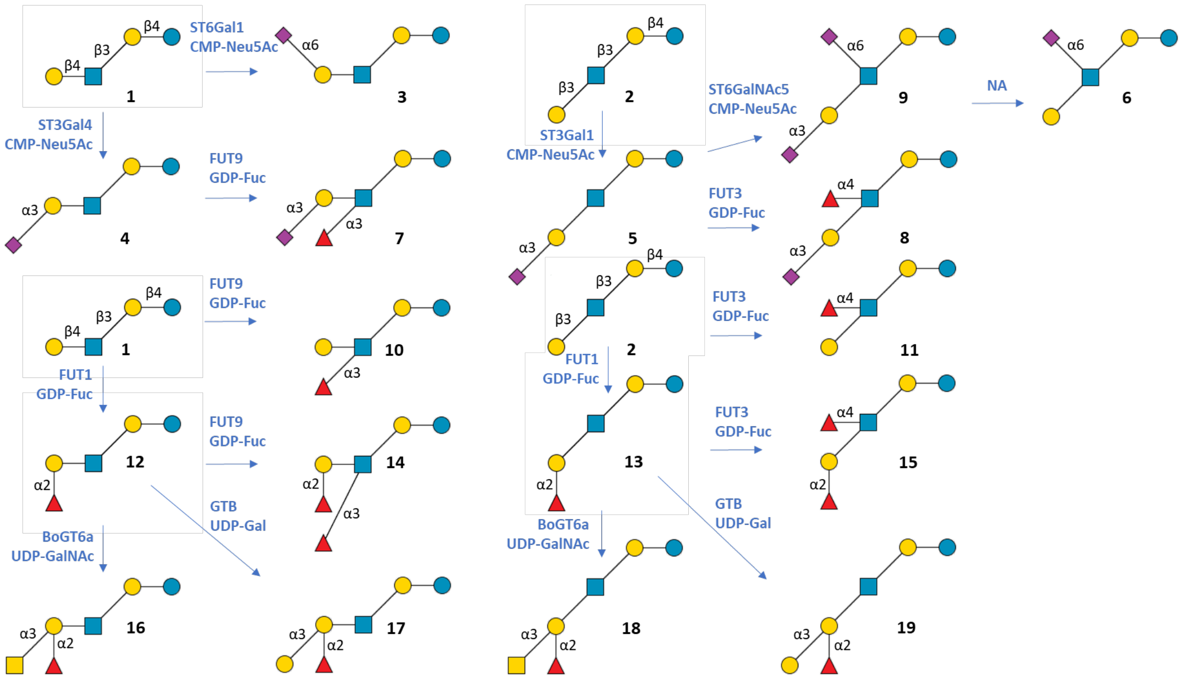

**Supplementary Figure 1**. Flowchart of enzymatic synthesis of HMO standards.

#### 1.1 Synthesis of lacto-*N*-neotetraose/LNnT (HMO1)

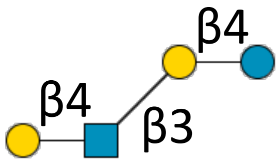

**1**

Lactose (30 mg, 69.4 mmol) was dissolved in HEPES buffer (50 mM HEPES, 25 mM KCl, 2 mM $MgCl_2$, 1 mM DTT, pH 7.3) at a concentration of 10 mM. To this, UDP-GlcNAc (1.5 eq, 15 mM), $B_3GnT2$ (10% v/v) and CIAP (1% v/v) were added, the mixture was incubated at 37 °C with gentle shaking and the reaction progress was monitored with LC-MS. The reaction mixture was passed through a Vivaspin® 20 10 kDa spinfilter by centrifugation at 4500 rpm. The filtrate was adjusted to a pH of 7-8 with NaOH (100 mM). UDP-gal (1.5 eq, 15mM), CIAP (1% v/v) and B4GalT1 (2% v/v) was added to the mixture, which was incubated at 37 °C with gentle shaking and the reaction progress was monitored with LC-MS. After completion, the mixture was freeze dried and ***1*** was purified using Bio-gel P2 size exclusion chromatography by elution with water. Fractions containing carbohydrate were identified by spotting 0.3 µl of each collected fraction on a TLC plate followed by a dip in 10% $H_2SO_4$ and charring. Fractions containing carbohydrate were analyzed by LC-MS and fractions with product were pooled and freeze dried. The solids were purified by preparative LC (yield: 19 mg, 26.9 mmol, 39% over 2 steps).

**$^1$H NMR (600 MHz, $D_2O$): δ (ppm).**

| | H1 | H2 | H3 | H4 | H5 | H6 | Ac |
|---|---|---|---|---|---|---|---|
| Glc(α) | 5.23 (d, J=3.8) | 3.58 | 3.84 | 3.65 | n/a | n/a | - |
| Glc(β) | 4.67 (d, J=8.0) | 3.28 (app. t) | 3.64 | 3.65 | n/a | n/a | - |
| Gal | 4.44 (d, J=7.82) | 3.60 | 3.73 | 4.16 (d, J=3.33) | n/a | n/a | - |
| GlcNAc | 4.71 (d, J=8.36) | 3.81 | 3.73 | 3.59 | n/a | n/a | 2.04 |
| Gal (2) | 4.48 (d, J=7.86) | 3.54 | 3.68 | 3.93 | n/a | n/a | - |

**$^{13}$C NMR derived from HSQC (150 MHz, $D_2O$): δ (ppm).**

| | C1 |
|---|---|
| Glc(α) | 91.63 |
| Glc(β) | 95.65 |
| Gal | 102.92 |
| GlcNAc | 102.64 |
| Gal (2) | 102.81 |

-ESI TOF-MS *m/z* calculated for $C_{26}H_{45}NO_{21}Na$ $(M + Na)^+$: 730.2376; measured: 730.2311.

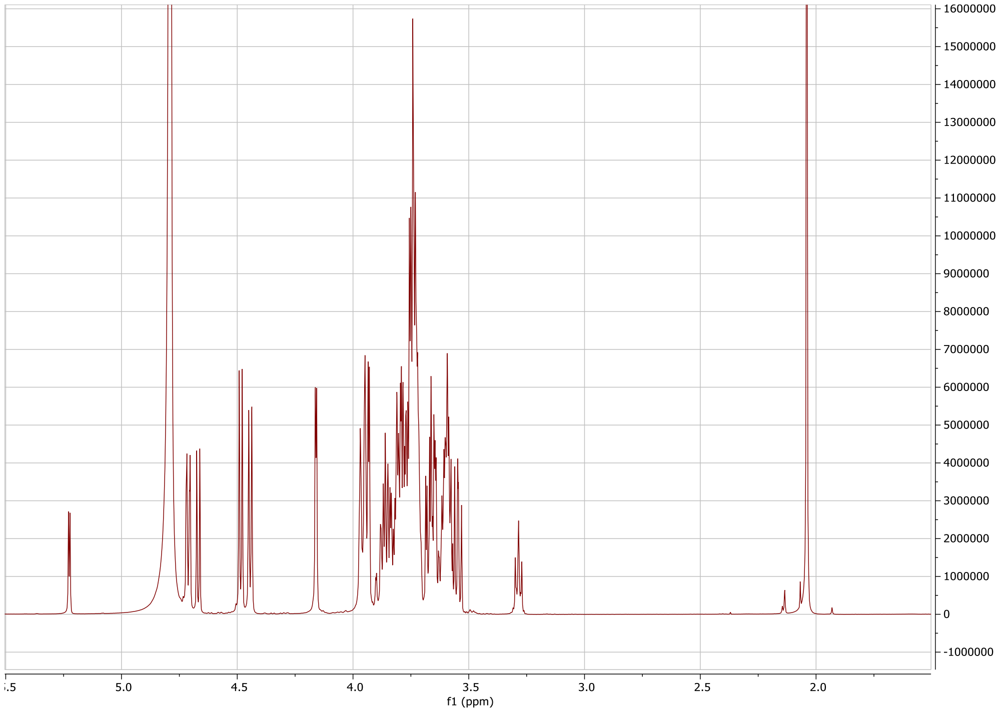

Proton spectrum of **1**.

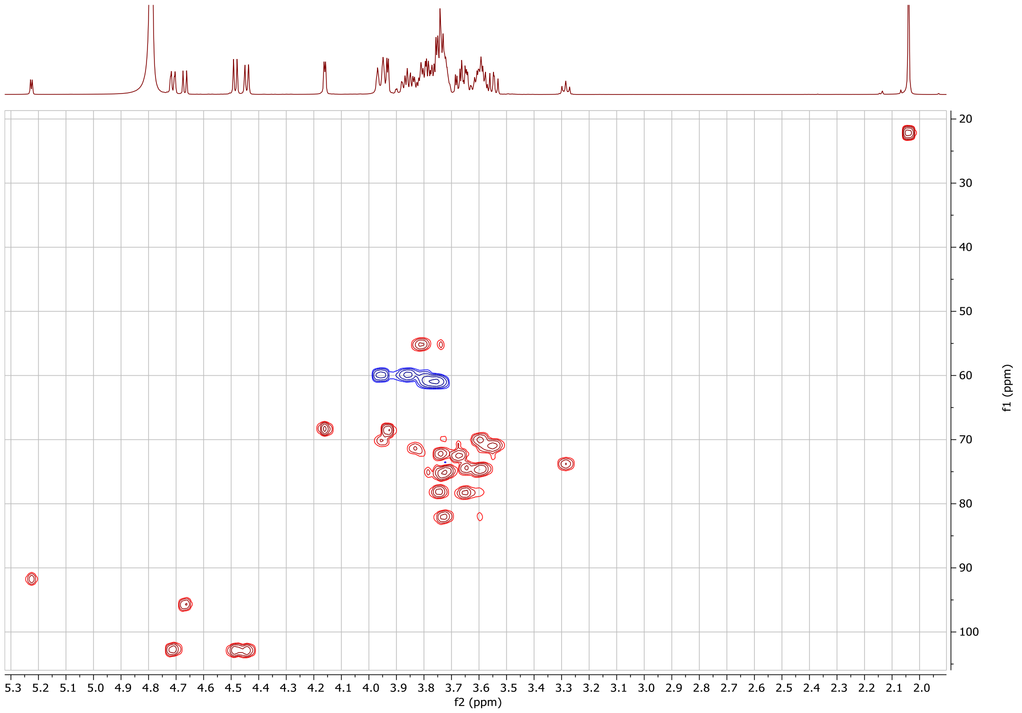

HSQC spectrum of **1**.

#### 1.2 Purification of lacto-*N*-tetraose/LNT (HMO 2)

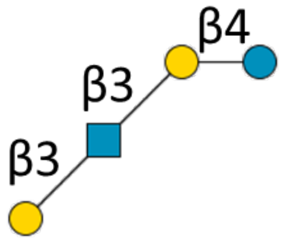

**2**

Crude Lacto-N-tetraose (100 mg) was purified by Bio-gel P2 size exclusion chromatography eluding with water to remove most lactose and lacto-*N*-triose impurities. Fractions containing only product were pooled and freeze dried (45 mg); 10 mg of the product was additionally purified by preparative LC resulting in 8 mg of high purity LNT.

**$^{1}$H NMR (600 MHz, $D_2O$): δ (ppm).**

| | H1 | H2 | H3 | H4 | H5 | H6 | Ac |
|---|---|---|---|---|---|---|---|
| Glc(α) | 5.23 (d, J=3.7) | 3.58 | 3.83 | 3.65 | n/a | n/a | - |
| Glc(β) | 4.58 (d, J=7.9) | 3.29 | 3.65 | 3.61 | n/a | n/a | - |
| Gal | 4.44 (d, J=7.9) | 3.61 | 3.74 | 4.16 | n/a | n/a | - |
| GlcNAc | 4.74 | 3.90 | 3.79 | 3.58 | n/a | n/a | 2.03 |
| Gal (2) | 4.44 (d, J=7.9) | 3.53 | 3.64 | 3.92 | n/a | n/a | - |

**$^{13}$C NMR derived from HSQC (150 MHz, $D_2O$): δ (ppm).**

| | C1 |
|---|---|
| Glc(α) | 95.73 |
| Glc(β) | 91.75 |
| Gal | 103.22 |
| GlcNAc | 102.63 |
| Gal (2) | 103.22 |

ESI TOF-MS *m/z* calculated for $C_{26}H_{45}NO_{21}Na$ $(M + Na)^+$: 730.2376; measured: 730.2318.

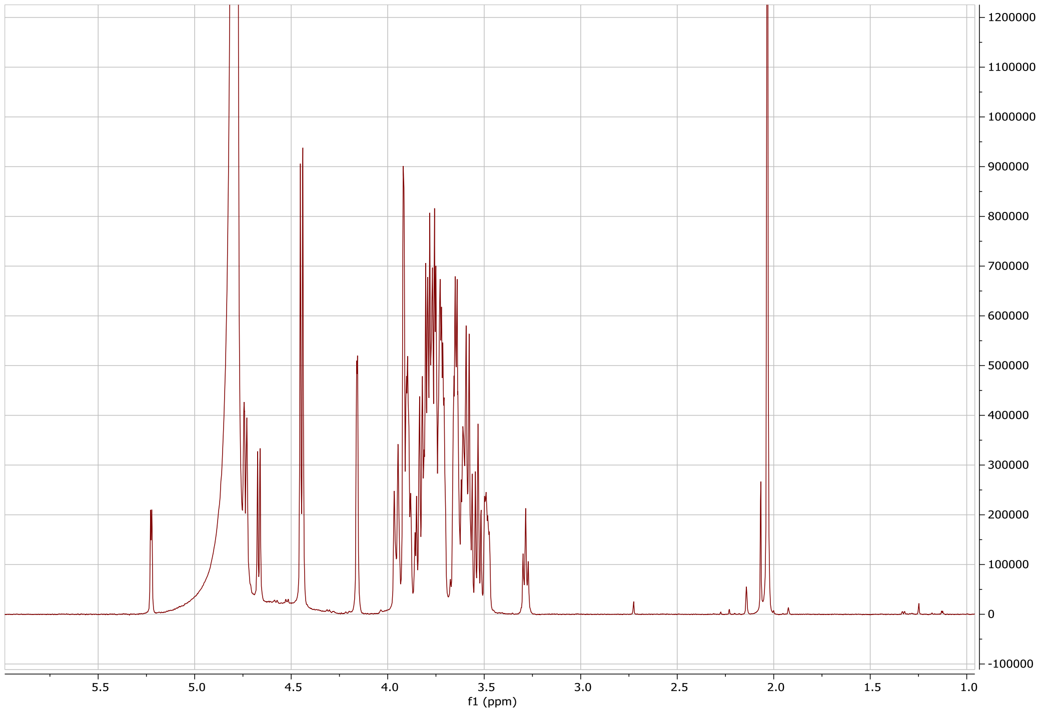

Proton spectrum of **2**.

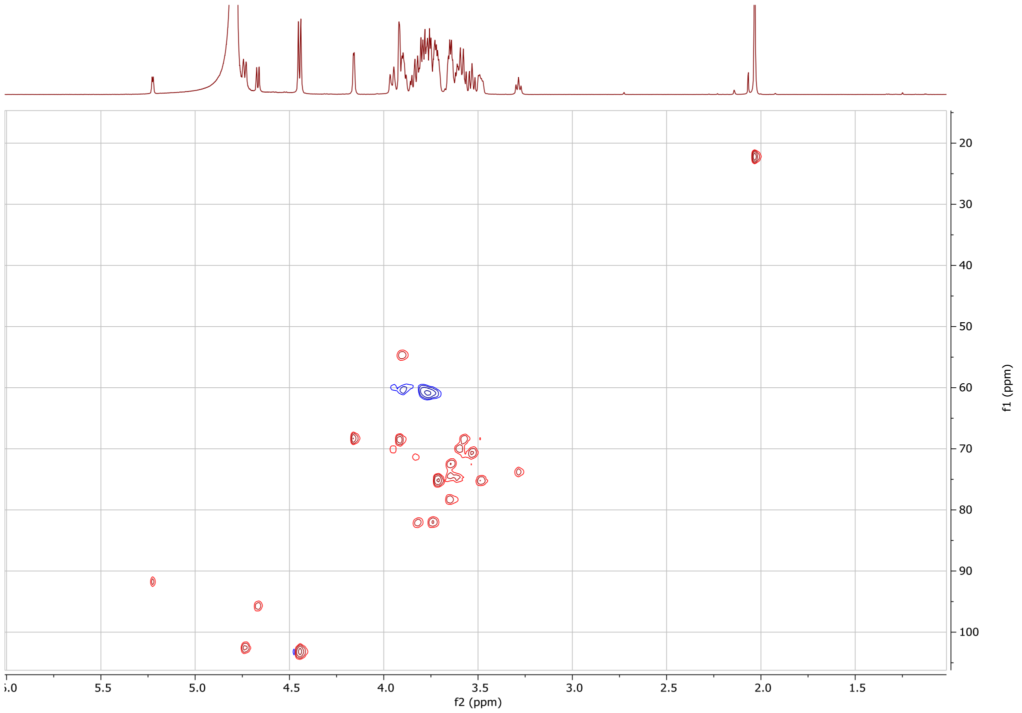

HSQC spectrum of **2**.

#### 1.3 Synthesis of α2-6 sialyllacto-*N*-neotetraose/ LSTc (HMO3)

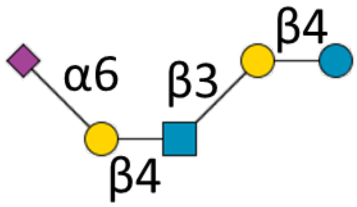

**3**

Compound **1** (LNnT; 2 mg, 2.8 µmol) was dissolved in 100 mM TRIS (pH 7.4) to a concentration of 10 mM and CMP-Sia (1.5 eq, 15 mM), bovine serum albumin (0.1% wt/v), CIAP (1% v/v) and ST6Gal1 (20% v/v) were added, after which the solution was incubated at 37 °C with gentle shaking. The reaction progress was monitored with LC-MS. Upon completion, the mixture was freeze dried and ***3*** was purified using size exclusion chromatography and preparative LC (yield: 2.4 mg, 86%, 2.4 µmol).

**$^{1}$H NMR (600 MHz, $D_2O$): δ (ppm)**

| | H1 | H2 | H3 | H4 | H5 | H6 | H7 | H8 | H9 | Ac |
|---|---|---|---|---|---|---|---|---|---|---|
| Glc(α) | 5.23 (d, J=3.80) | 3.57 | 3.83 | 3.65 | n/a | n/a | - | - | - | - |
| Glc(β) | 4.67 (d, J=7.98) | 3.28 (app. T) | 3.65 | 3.84 | n/a | n/a | - | - | - | - |
| Gal | 4.44 ( | 3.61 | 3.73 | 4.16 (d, J=3.31 | n/a | n/a | - | - | - | - |
| GlcNAc | 4.74 (d, J=6.64) | 3.81 | 3.61 | 3.84 | n/a | n/a | - | - | - | 2.03 |
| Gal (2) | 4.46 | 3.54 | 3.68 | 3.93 | n/a | n/a | - | - | - | - |
| Sia | - | - | 1.73 (t, J=12.19), 2.68 (dd, J=4.68/12.42) | 3.67 | 3.81 | 3.57 | n/a | n/a | n/a | 2.06 |

**$^{13}$C NMR derived from HSQC (150 MHz, $D_2O$): δ (ppm).**

| | C1 |
|---|---|
| Glc(α) | 91.68 |
| Glc(β) | 95.66 |
| Gal | 103.14 |
| GlcNAc | 102.52 |
| Gal (2) | 103.14 |

ESI TOF-MS *m/z* calculated for $C_{37}H_{62}N_2O_{29}Na$ $(M + Na)^+$: 1021.3330; measured: 1021.3239.

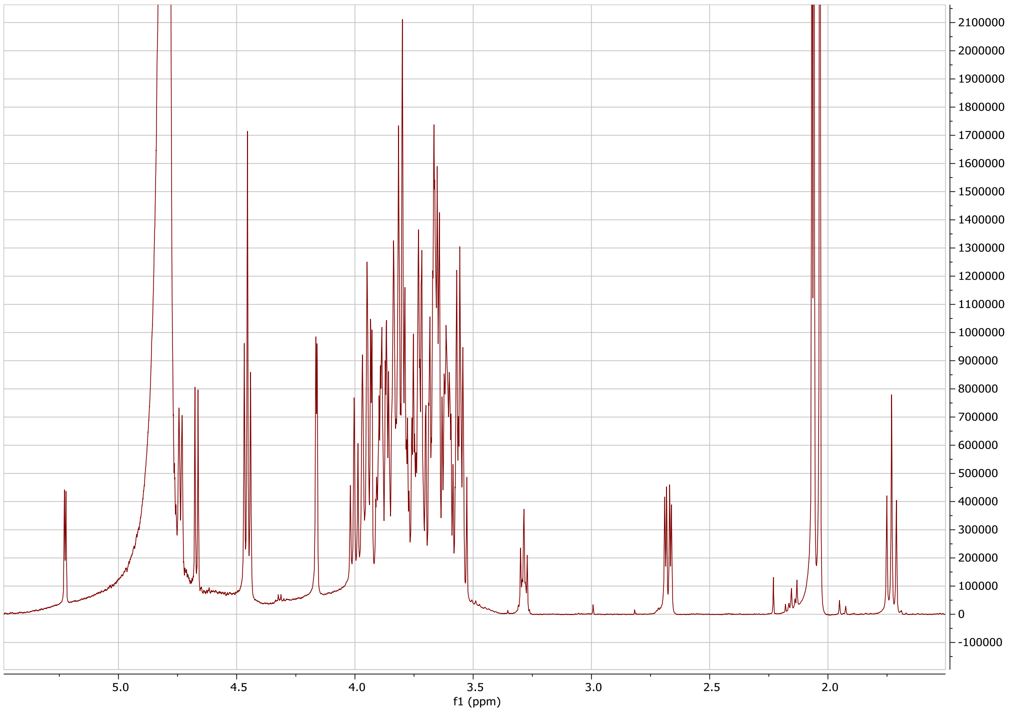

Proton spectrum of **3**.

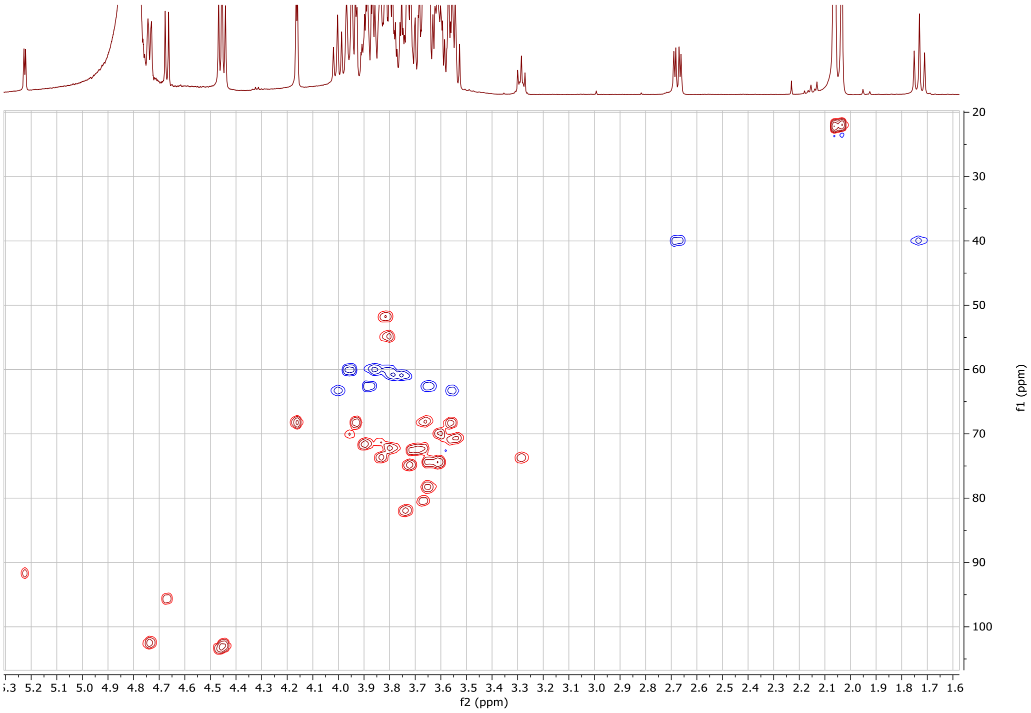

HSQC spectrum of **3**.

#### 1.4 Synthesis of α2-3 sialyllacto-*N*-neotetraose/LSTd (HMO4)

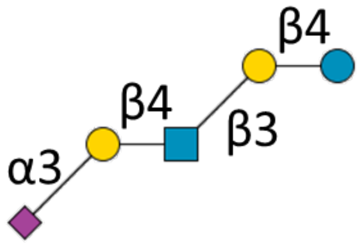

**4**

LNnT **1** (4 mg, 5.6 µmol) was dissolved in 100 mM TRIS (pH 7.4) to a concentration of 10 mM and CMP-Sia (1.5 eq, 15 mM), bovine serum albumin (0.1% wt/v), CIAP (1% v/v) and ST3Gal4 (30% v/v) were added, after which the solution was incubated at 37 °C with gentle shaking. The reaction progress was monitored with LC-MS. Upon reaction completion, the mixture was freeze dried and ***4*** was purified using size exclusion chromatography and preparative LC (yield: 3.9 mg, 70%, 3.9 µmol).

**$^{1}$H NMR (600 MHz, $D_2O$): δ (ppm).**

| | H1 | H2 | H3 | H4 | H5 | H6 | H7 | H8 | H9 | Ac |
|---|---|---|---|---|---|---|---|---|---|---|
| Glc(α) | 5.22 (d, J=3.80) | 3.58 | 3.84 | 3.65 | n/a | n/a | - | - | - | - |
| Glc(β) | 4.67 (d, J=7.90) | 3.28 (app. t) | 3.65 | 3.80 | n/a | n/a | - | - | - | - |
| Gal | 4.45 (d, J=7.90) | 3.60 | 3.73 | 4.17 (d, J=3.34) | n/a | n/a | - | - | - | - |
| GlcNAc | 4.71 (d, J=8.34) | 3.81 | 3.58 | 3.75 | n/a | n/a | - | - | - | 2.05 (6H) |
| Gal (2) | 4.57 (d, J=7.85) | 3.57 | 4.12 | 3.96 (dd, J=3.14/9.87) | 3.87 | n/a | - | - | - | - |
| Sia | - | - | 1.80 (t, J=12.11), 2.77 (dd, J=4.62/12.45) | 3.70 | 3.86 | 3.65 | n/a | n/a | n/a | 2.05 (6H) |

**$^{13}$C NMR derived from HSQC (150 MHz, $D_2O$): δ (ppm).**

| | C1 |
|---|---|
| Glc(α) | 91.73 |
| Glc(β) | 95.69 |
| Gal | 102.90 |
| GlcNAc | 102.79 |
| Gal (2) | 102.51 |

ESI TOF-MS *m/z* calculated for $C_{37}H_{62}N_2O_{29}Na$ $(M + Na)^+$: 1021.3330; measured: 1021.3226.

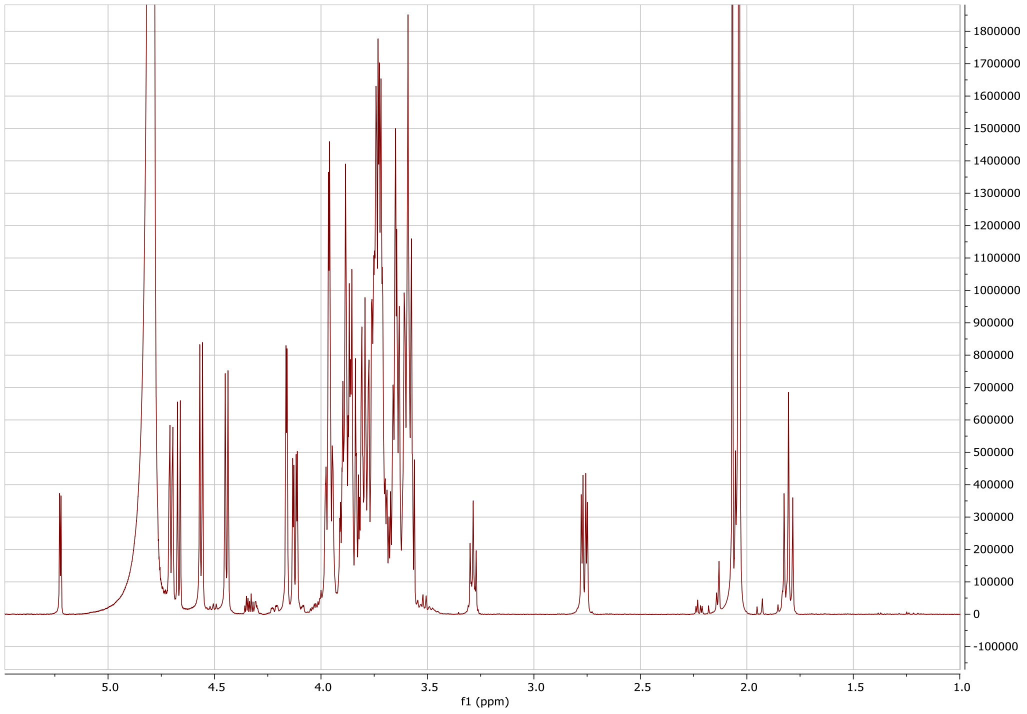

Proton spectrum of **4**.

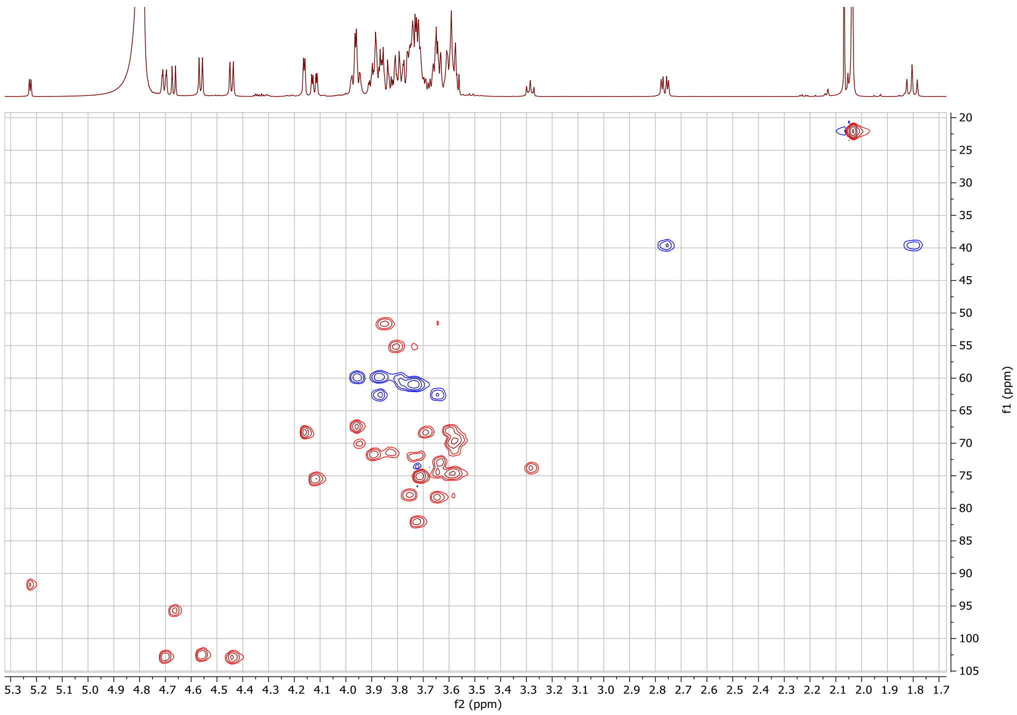

HSQC spectrum of **4**.

#### 1.5 Synthesis of α2-3 sialyllacto-*N*-tetraose/LSTa (HMO5)

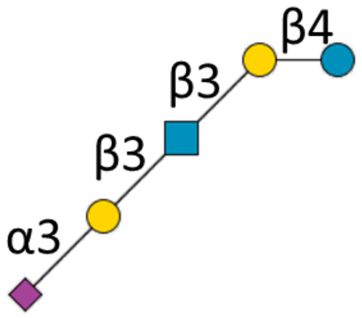

**5**

LNT **2** (7.5 mg, 10.6 µmol) was dissolved in 100 mM TRIS (pH 7.4) to a concentration of 10 mM, and CMP-Sia (1.5 eq, 15 mM), bovine serum albumin (0.1% wt/v), CIAP 1% (v/v) and ST3Gal1 (10% v/v) were added, after which the solution was incubated at 37 °C with gentle shaking. The reaction progress was monitored with LC-MS. Upon reaction completion, the mixture was freeze dried and ***5*** was purified using size exclusion chromatography and preparative LC (yield: 4 mg, 38%, 4.0 µmol).

**$^{1}$H NMR (600 MHz, $D_2O$): δ (ppm).**

| | H1 | H2 | H3 | H4 | H5 | H6 | H7 | H8 | H9 | Ac |
|---|---|---|---|---|---|---|---|---|---|---|
| Glc(α) | 5.23 (d, J=3.76) | 3.59 (app. t) | 3.84 | 3.65 | n/a | n/a | - | - | - | - |
| Glc(β) | 4.67 (d, J=7.95) | 3.28 | 3.65 | 3.96 | n/a | n/a | - | - | - | - |
| Gal | 4.45 (d, J=7.85) | 3.60 | 3.74 | 4.16 (d, J=3.33) | n/a | n/a | - | - | - | - |
| GlcNAc | 4.75 (d, J=8.48) | 3.90 | 3.81 | 3.49 | n/a | n/a | - | - | - | 2.05 (6H) |
| Gal (2) | 4.52 (d, J=7.76) | 3.55 | 4.09 (dd, J=3.13,9.81) | 3.94 | n/a | n/a | - | - | - | - |
| Sia | - | - | 1.79 (t, J=12.16), 2.77 (dd, J=4.61,12.45) | 3.69 | 3.85 | n/a | n/a | n/a | n/a | 2.05 (6H) |

**$^{13}$C NMR derived from HSQC (150 MHz, $D_2O$): δ (ppm).**

| | C1 |
|---|---|
| Glc(α) | 91.75 |
| Glc(β) | 95.67 |
| Gal | 102.93 |
| GlcNAc | 102.43 |
| Gal (2) | 103.33 |

ESI TOF-MS *m/z* calculated for $C_{37}H_{62}N_2O_{29}Na$ $(M + Na)^+$: 1021.3330; measured: 1021.3239.

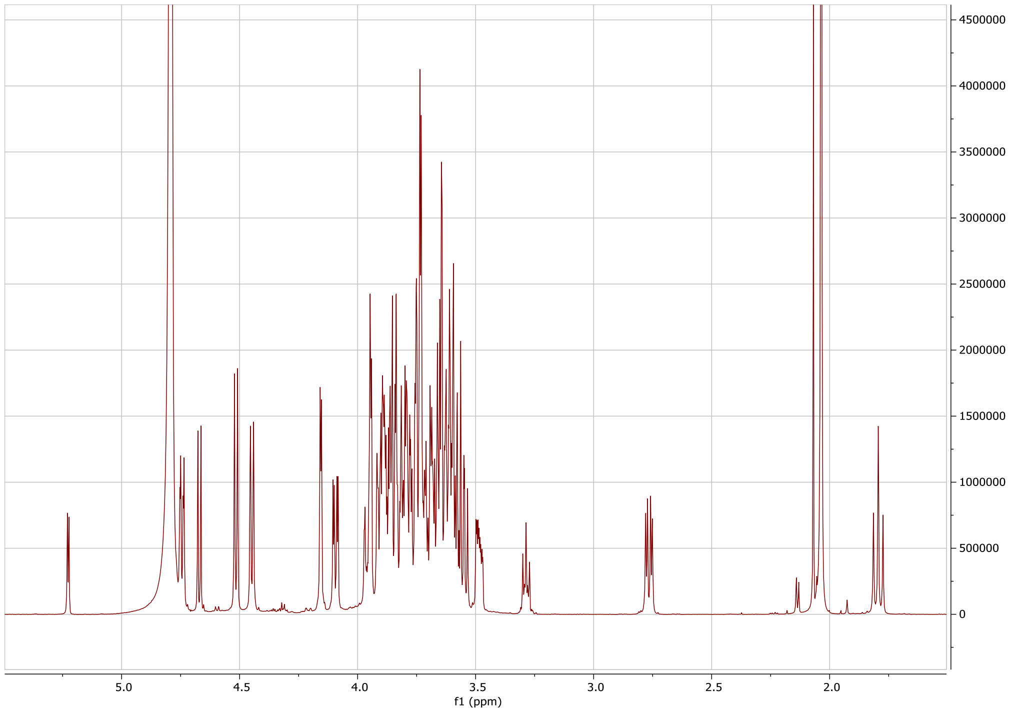

Proton spectrum of **5**.

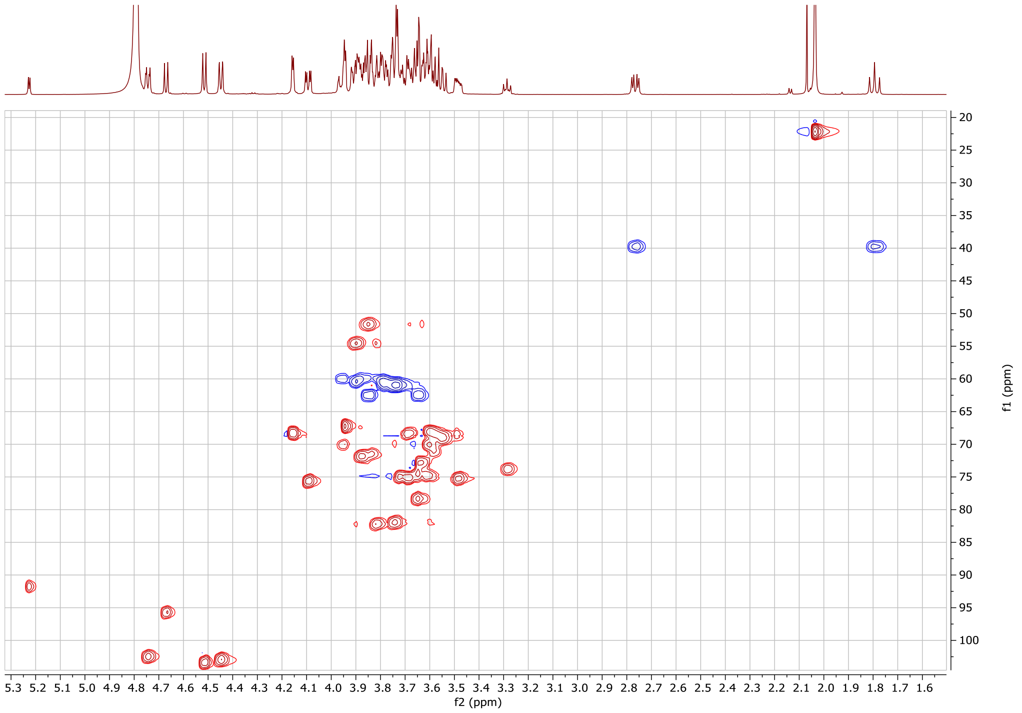

HSQC spectrum of **5**.

#### 1.6 Synthesis of *β*-Gal(1-3)-[*α*-sialyl(2-6)]-*β*-GlcNAc(1-3)-*β*-Gal(1-4)-Glc/LSTb (HMO6)

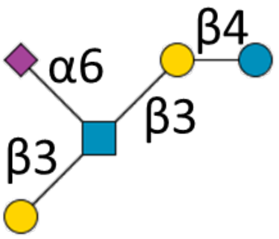

**6**

LNT **2** (4 mg, 5.6 µmol) was dissolved in 50 mM TRIS (pH 7.5) to a concentration of 10 mM and CMP-Sia (1.5 eq, 15 mM), bovine serum albumin (0.1 wt%), CIAP (1% v/v) and ST6GalNAc5 (50% v/v) were added, after which the solution was incubated at 37 °C with gentle shaking. The reaction progress was monitored with LC-MS. When no further reaction progress was observed after 2 days, the mixture was freeze dried and ***6*** was purified using size exclusion chromatography and preparative LC (yield: 2.6 mg, 46%, 2.6 µmol).

**$^{1}$H NMR (600 MHz, $D_2O$): δ (ppm).**

| | H1 | H2 | H3 | H4 | H5 | H6 | H7 | H8 | H9 | Ac |
|---|---|---|---|---|---|---|---|---|---|---|
| Glc(α) | 5.22 (d, J=3.78) | 3.59 | 3.81 | 3.65 | n/a | n/a | - | - | - | - |
| Glc(β) | 4.67 (d, J=8.03) | 3.28 | 3.65 | 3.61 | n/a | n/a | - | - | - | - |
| Gal | 4.44 (d, J=7.81) | 3.59 | 3.73 | 4.29 (d, J=3.26) | n/a | n/a | - | - | - | - |
| GlcNAc | 4.70 (d, J=8.52) | 3.91 | 3.81 | 3.64 | 3.56 | 3.77, 3.96 | - | - | - | 2.05 (6H) |
| Gal (2) | 4.44 (d, J=7.81) | 3.53 | 3.63 | 3.91 | n/a | n/a | - | - | - | - |
| Sia | - | - | 1.70 (t, J=12.41), 2.75 (dd, J=4.68/12.42) | 3.68 | 3.83 | n/a | n/a | n/a | n/a | 2.05 (6H) |

**$^{13}$C NMR derived from HSQC (150 MHz, $D_2O$): δ (ppm).**

| | C1 |
|---|---|
| Glc(α) | 91.71 |
| Glc(β) | 95.69 |
| Gal | 103.15 |
| GlcNAc | 102.59 |
| Gal (2) | 103.15 |

ESI TOF-MS m/z calculated for $C_{37}H_{62}N_2O_{29}Na$ $(M + Na)^+$: 1021.3330; measured: 1021.3237.

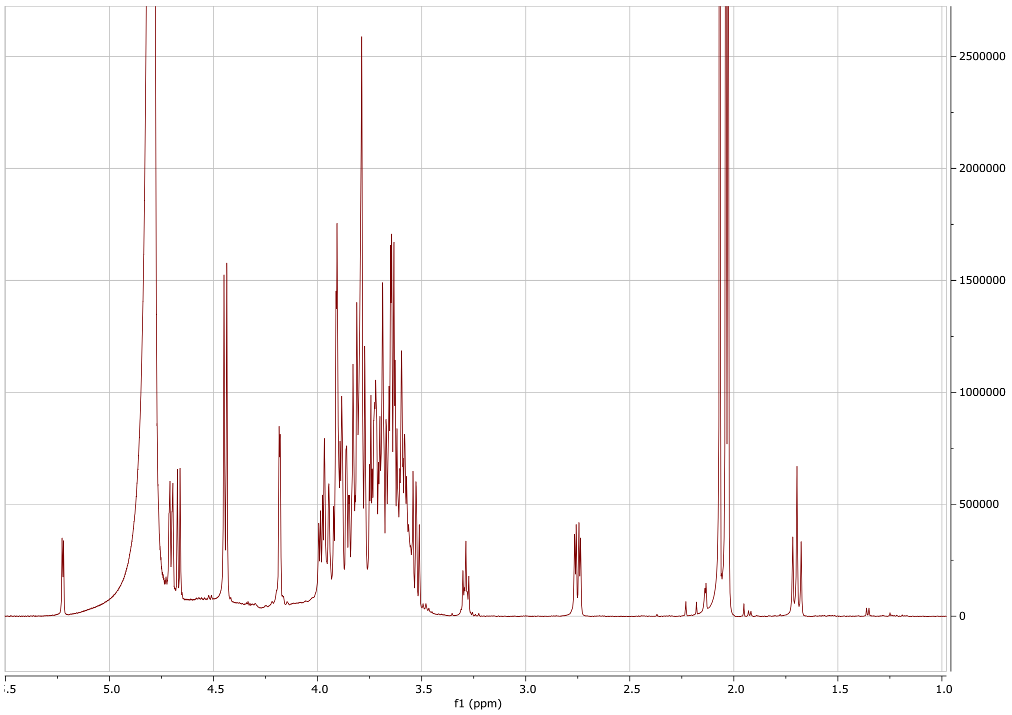

Proton spectrum of **6**.

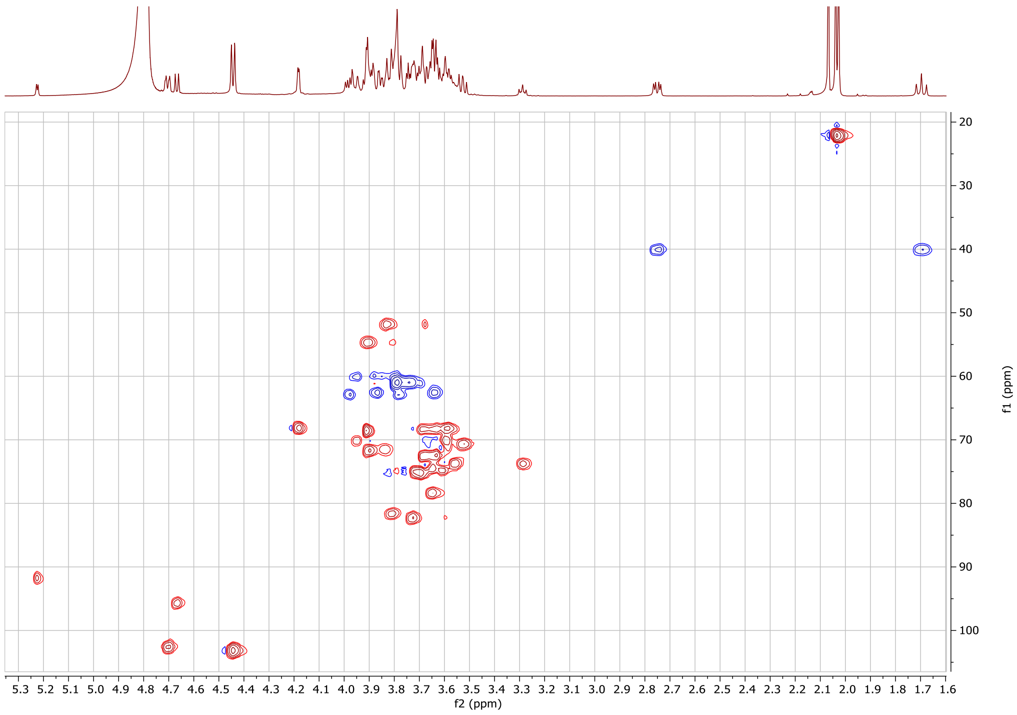

HSQC spectrum of **6**.

#### 1.7 Synthesis of sialyl Lewis x lacto-*N*-neohexose/SLeXLNnH (HMO7)

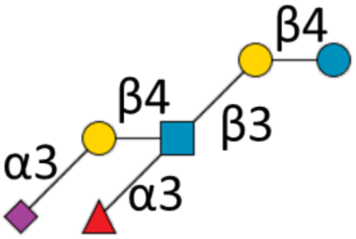

**7**

***4*** (1 mg, 1 µmol) was dissolved in 100 mM TRIS (pH 7.5) to a concentration of 10 mM, and GDP-Fuc (0.5 eq, 5 mM), $MnCl_2$ (10 mM, from 1 M solution at 1% v/v of reaction volume), CIAP (1% v/v) and FUT9 (10% v/v) were added, after which the solution was incubated at 37 °C with gentle shaking. The reaction progress was monitored with LC-MS. A volume of 0.1 eq. of GDP-Fucose was added each hour until the reaction went to completion, after which the mixture was freeze dried. Size exclusion chromatography and preparative LC were used to obtain ***7*** (yield: 0.4 mg 31%, 3.1 µmol).

**$^{1}$H NMR (600 MHz, $D_2O$): δ (ppm).**

| | H1 | H2 | H3 | H4 | H5 | H6 | H7 | H8 | H9 | Ac |
|---|---|---|---|---|---|---|---|---|---|---|
| Glc(α) | 5.23 (d, J=3.82) | 3.58 | 3.83 | 3.65 | n/a | n/a | - | - | - | - |
| Glc(β) | 4.67 (d, J=7.92) | 3.28 (app. t, J=8.49) | 3.64 | 3.80 | n/a | n/a | - | - | - | - |
| Gal | 4.44 (d, J=7.90) | 3.67 | 3.71 | 4.17 (d, J=3.32) | n/a | n/a | - | - | - | - |
| GlcNAc | 4.71 (d, J=9.28) | 3.97 | 3.88 | 3.60 | n/a | n/a | - | - | - | 2.05 (6H) |
| Gal (2) | 4.53 (d, J=7.79) | 3.53 | 4.09 (dd, J=3.19/ 9.84) | 3.94 | n/a | n/a | - | - | - | - |
| Fuc | 5.13 (d, J=4.04) | 3.68 | 3.90 | 3.78 | 4.83 | 1.17 (d, J=6.5 4) | - | - | - | - |
| Sia | - | - | 1.80 (t, J=12.17), 2.77 (dd, J=4.61/12.48) | 3.69 | 3.86 | n/a | n/a | n/a | n/a | 2.05 (6H) |

**$^{13}$C NMR derived from HSQC (150 MHz, $D_2O$): δ (ppm).**

| | C1 |
|---|---|
| Glc(α) | 91.83 |
| Glc(β) | 95.68 |
| Gal | 102.68 |
| GlcNAc | 102.53 |
| Gal (2) | 101.42 |
| Fuc | 98.46 |

ESI TOF-MS m/z calculated for $C_{43}H_{72}N_2O_{33}Na$ $(M + Na)^+$: 1167.3910; measured: 1167.3804.

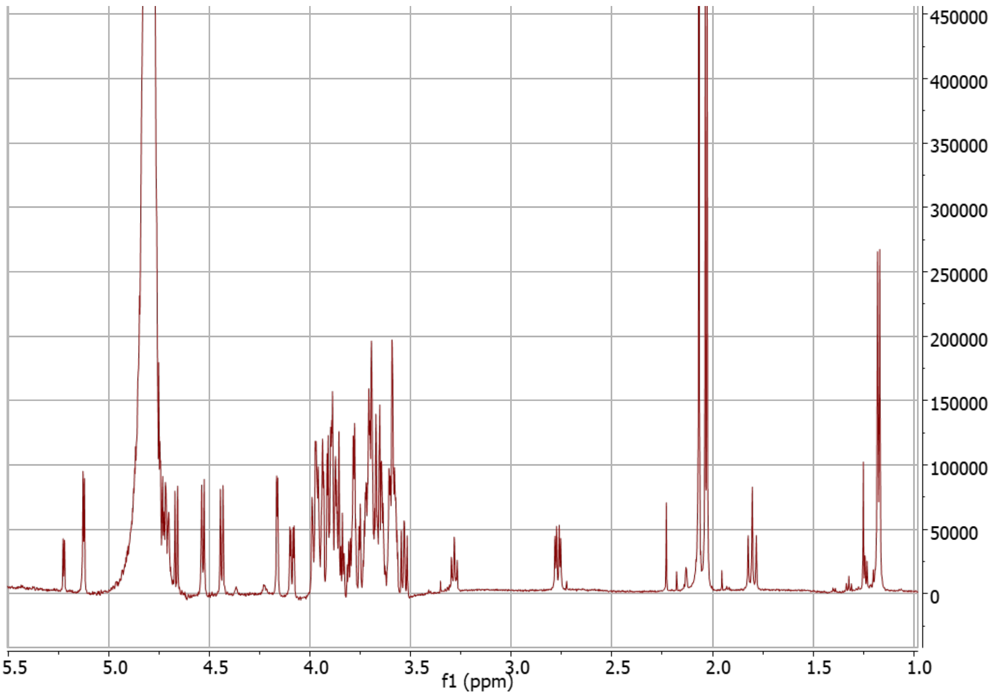

Proton spectrum of **7**.

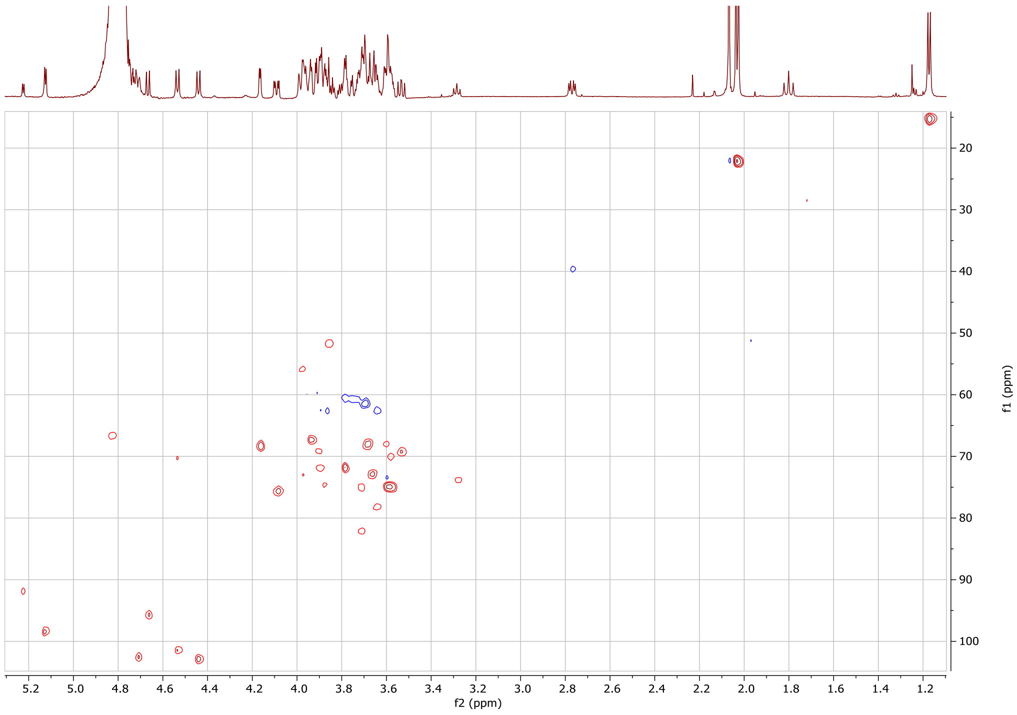

HSQC spectrum of **7**.

#### 1.8 Synthesis of sialyl Lewis a lacto-*N*-hexose/SLeALNH (HMO8)

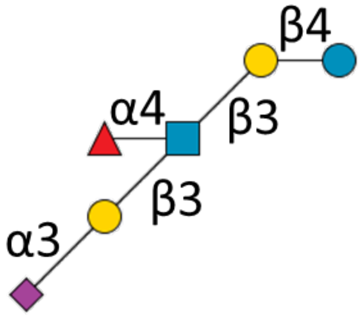

**8**

***5*** (1 mg, 1 µmol) was dissolved in 100 mM TRIS (pH 7.5) to a concentration of 10 mM, and GDP-Fuc (0.5 eq, 5 mM), $MnCl_2$ (10 mM, from 1 M solution at 1% v/v of reaction volume), CIAP (1% v/v) and FUT3 (10% v/v) were added, after which the solution was incubated at 37 °C with gentle shaking. The reaction progress was monitored with LC-MS. A volume of 0.1 eq. of GDP-Fucose was added each hour until the reaction went to completion, after which the mixture was freeze dried. Size exclusion chromatography and preparative LC were used to obtain **8** (yield: 1 mg 87%, 0.9 µmol).

**$^1$H NMR (600 MHz, $D_2O$): δ (ppm).**

| | H1 | H2 | H3 | H4 | H5 | H6 | H7 | H8 | H9 | Ac |
|---|---|---|---|---|---|---|---|---|---|---|
| Glc(α) | 5.23 (d, J=3.78) | 3.58 | 3.84 | 3.65 | n/a | n/a | - | - | - | - |
| Glc(β) | 4.67 (d, J=7.94) | 3.28 | 3.65 | n/a | n/a | n/a | - | - | - | - |
| Gal | 4.44 (d, J=7.88) | 3.59 | 3.72 | 4.16 (d, J=3.32) | n/a | n/a | - | - | - | - |
| GlcNAc | 4.71 (d, J=9.51) | 3.94 | 4.10 (t, J=9.83) | 3.73 | 3.53 | n/a | - | - | - | 2.05 (6H) |
| Gal (2) | 4.55 (d, J=7.70) | 3.51 | 4.06 (dd, J=3.14/9.78) | 3.92 | n/a | n/a | - | - | - | - |
| Fuc | 5.02 (d, J=4.00) | 3.79 | 3.88 | 3.78 | 4.88 (q, J=6.64) | 1.18 (3H, d, J=6.57) | - | - | - | - |
| Sia | - | - | 1.77 (t, J=12.81), 2.77 (dd, J=4.69/12.44) | 3.68 | 3.84 | n/a | n/a | n/a | n/a | 2.05 (6H) |

**$^{13}$C NMR derived from HSQC (150 MHz, $D_2O$): δ (ppm).**

| | C1 |
|---|---|
| Glc(α) | 91.77 |
| Glc(β) | 95.72 |
| Gal | 102.96 |
| GlcNAc | 102.47 |
| Gal (2) | 102.67 |
| Fuc | 97.93 |

ESI TOF-MS m/z calculated for $C_{43}H_{72}N_2O_{33}Na$ $(M + Na)^+$: 1167.3910; measured: 1167.3798.

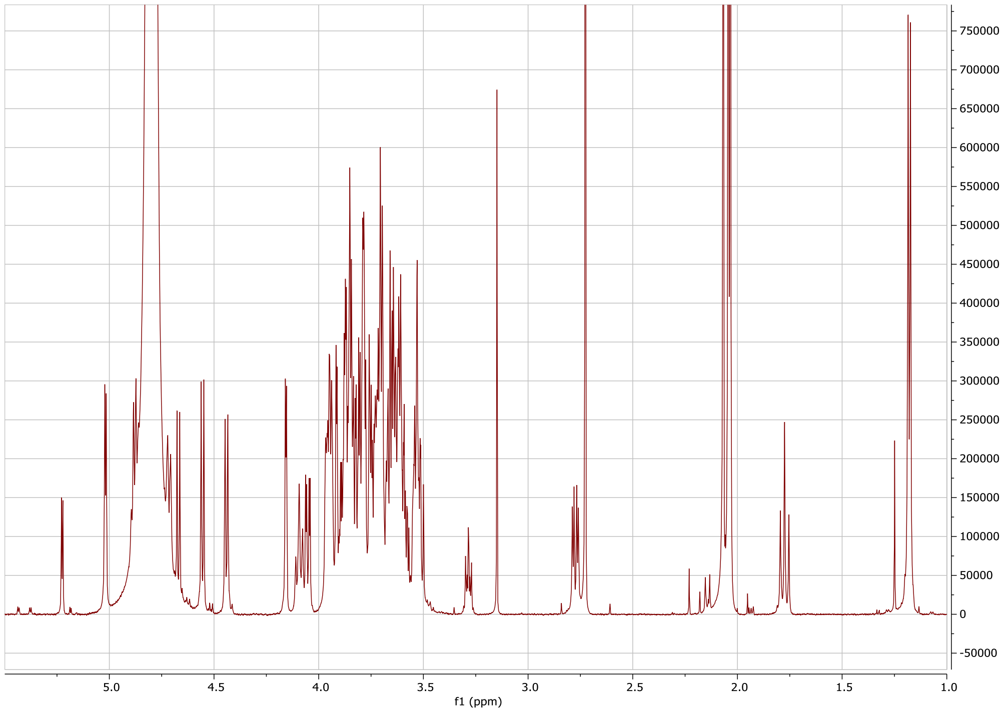

Proton spectrum of **8**.

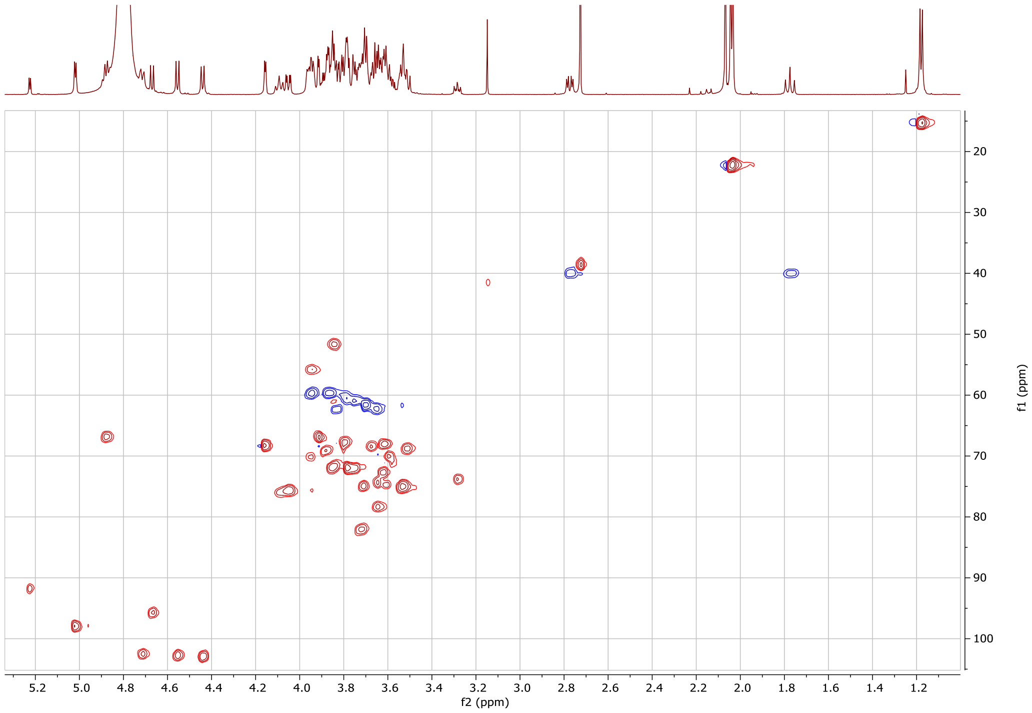

HSQC spectrum of **8**.

#### 1.9 Synthesis of disialyllacto-*N*-tetraose/DSLNT (HMO9)

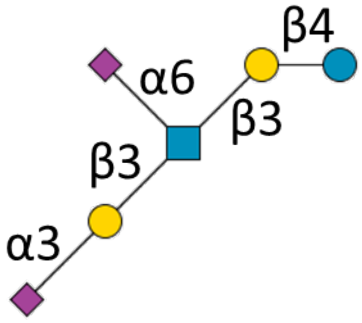

**9**

LNT **2** (10 mg, 14.1 µmol) was dissolved in 50 mM TRIS (pH 7.5) to a concentration of 10 mM, and CMP-Sia (3 eq, 30 mM), bovine serum albumin (0.1 wt%), CIAP (1% v/v) and ST3Gal1 (20% v/v) were added, after which the solution was incubated at 37 °C with gentle shaking. The reaction progress was monitored with LC-MS. When no starting material was detected, ST6GalNAc5 (10% v/v) was added, and the reaction was incubated again. Once no more mono-sialylated product was detected, the mixture was freeze dried and **9** was purified using size exclusion chromatography and preparative LC (yield: 11.7 mg, 64%, 9.0 µmol).

**$^{1}$H NMR (600 MHz, $D_2O$): δ (ppm).**

| | H1 | H2 | H3 | H4 | H5 | H6 | H7 | H8 | H9 | Ac |
|---|---|---|---|---|---|---|---|---|---|---|
| Glc(α) | 5.22 (d, J=3.77 | 3.59 (app. t) | 3.84 | 3.65 | n/a | n/a | - | - | - | - |
| Glc(β) | 4.67 (d, J=7.95) | 3.29 | 3.65 | 3.61 | n/a | n/a | - | - | - | - |
| Gal | 4.44 (d, J=7.87) | 3.60 | 3.73 | 4.17 (d, J=3.30) | n/a | n/a | - | - | - | - |
| GlcNAc | 4.71 (d, J=8.44) | 3.90 | 3.80 | 3.62 | 3.56 | 3.99 | - | - | - | 2.07 (3H) |
| Gal (2) | 4.51 (d, J=7.80) | 3.54 | 4.09 (d, J=9.84) | 3.94 | n/a | n/a | - | - | - | - |
| Sia | - | - | 1.70 (t, J=12.16), 2.75 (m) | 3.70 | 3.84 | n/a | n/a | n/a | n/a | 2.03 (6H) |
| Sia(2) | - | - | 1.80 (t, J=12.15), 2.77 (m) | 3.68 | 3.85 | n/a | n/a | n/a | n/a | 2.03 (6H) |

**$^{13}$C NMR derived from HSQC (150 MHz, $D_2O$): δ (ppm).**

| | C1 |
|---|---|
| Glc(α) | 91.81 |
| Glc(β) | 95.75 |
| Gal | 102.96 |
| GlcNAc | 102.51 |
| Gal (2) | 103.46 |

ESI TOF-MS m/z calculated for $C_{48}H_{79}N_3O_{37}Na$ $(M + Na)^+$: 1312.4285; measured: 1312.4184.

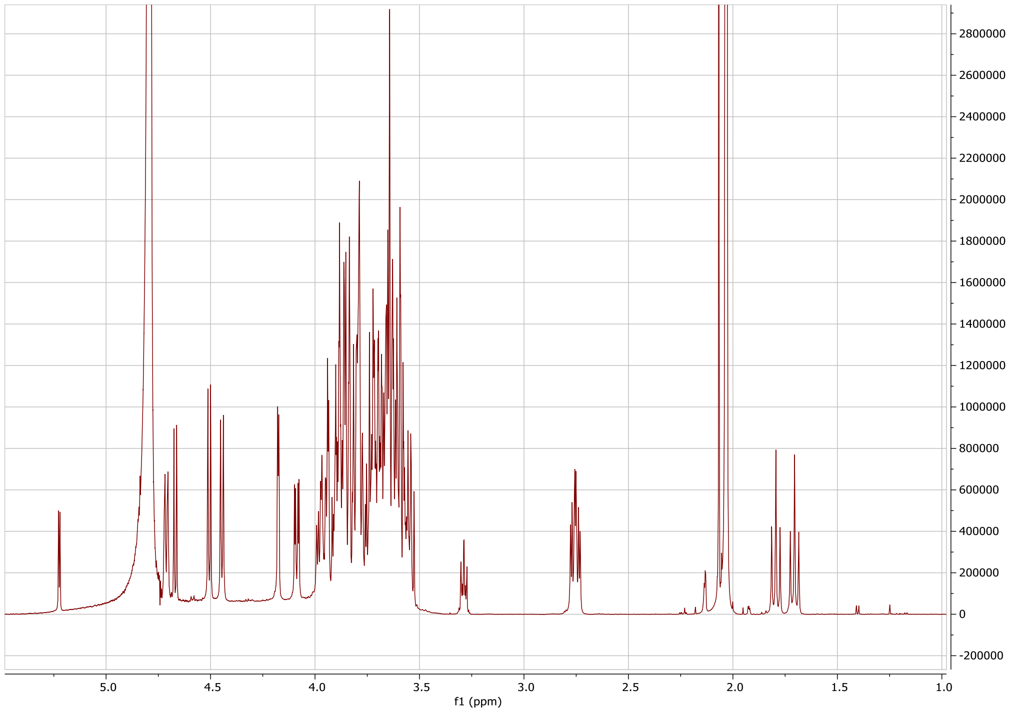

Proton spectrum of **9**.

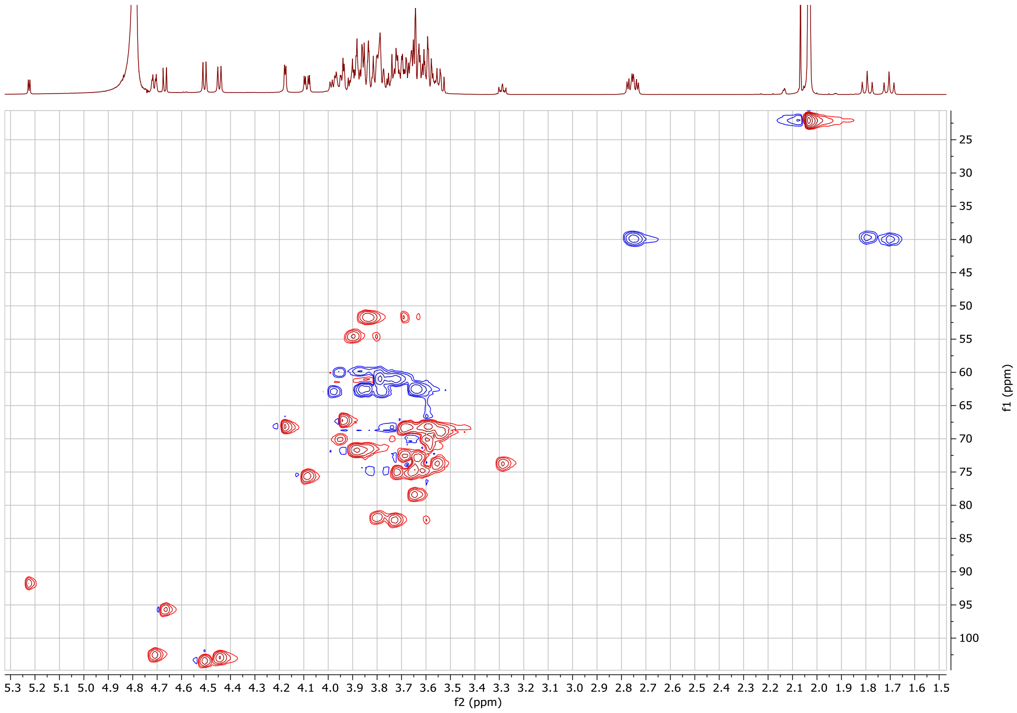

HSQC spectrum of **9**.

#### 1.10 Synthesis of lewis x lacto-*N*-neopentose/LNFPIII (HMO10)

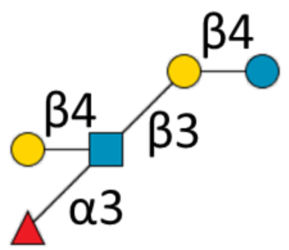

**10**

LnNT **1** (3 mg, 4.2 µmol) was dissolved in 100 mM TRIS (pH 7.5) to a concentration of 10 mM, and GDP-Fuc (0.5 eq, 5 mM), $MnCl_2$ (10 mM, from 1 M solution at 1% v/v of reaction volume), CIAP (1% v/v) and FUT9 (10% v/v) were added, after which the solution was incubated at 37 °C with gentle shaking. The reaction progress was monitored with LC-MS. A volume of 0.1 eq. of GDP-Fucose was added each hour until the reaction went to completion, after which the mixture was freeze dried. Size exclusion chromatography and preparative LC were used to obtain **10** (yield: 1.7 mg, 48%, 2.0 µmol).

**$^{1}$H NMR (600 MHz, $D_2O$): δ (ppm).**

| | H1 | H2 | H3 | H4 | H5 | H6 | Ac |
|---|---|---|---|---|---|---|---|
| Glc(α) | 5.23 (d, J=3.77) | 3.58 (app. t) | 3.83 | 3.65 | n/a | n/a | - |
| Glc(β) | 4.67 (d, J=7.91) | 3.28 | 3.64 | 3.61 | n/a | n/a | - |
| Gal | 4.44 (d, J=7.87) | 3.58 | 3.71 | 4.16 (d, J=3.38) | n/a | n/a | - |
| GlcNAc | 4.72 (d, J=8.35) | 3.97 | 3.89 | 3.59 | n/a | n/a | 2.03 (3H) |
| Gal (2) | 4.47 (d, J=7.83) | 3.50 (dd, J=7.80,9.84) | 3.66 | 3.91 | n/a | n/a | - |
| Fuc | 5.13 (d, J=4.03) | 3.70 | 3.91 | 3.81 | 4.84 | (1.18, 3H) | - |

**$^{13}$C NMR derived from HSQC (150 MHz, $D_2O$): δ (ppm).**

| | C1 |
|---|---|
| Glc(α) | 91.76 |
| Glc(β) | 95.66 |
| Gal | 102.81 |
| GlcNAc | 102.49 |
| Gal (2) | 101.73 |
| Fuc | 98.49 |

ESI TOF-MS m/z calculated for $C_{32}H_{55}NO_{25}Na$ $(M + Na)^+$: 876.2955; measured: 876.2888.

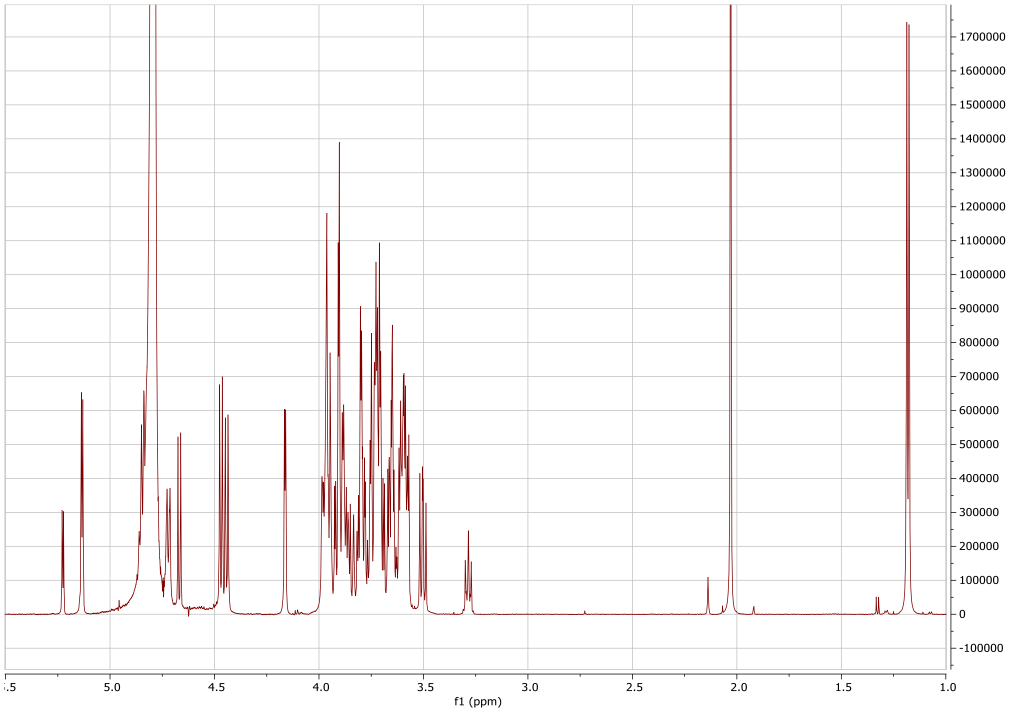

Proton spectrum of **10**.

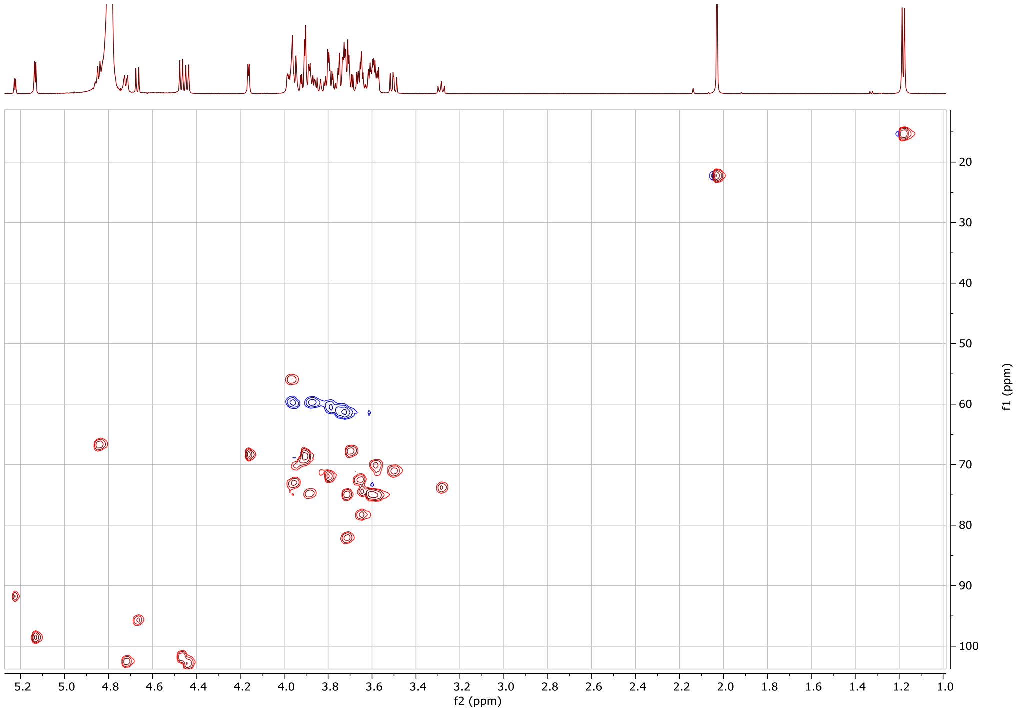

HSQC spectrum of **10**.

#### 1.11 Synthesis of Lewis a lacto-*N*-pentose/LNFPII (HMO11)

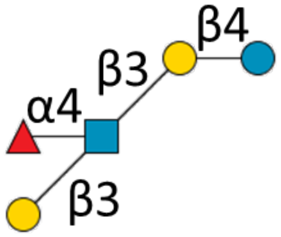

**11**

LNT **2** (3 mg, 5.6 µmol) was dissolved in 100 mM TRIS (pH 7.5) to a concentration of 10 mM, and GDP-Fuc (0.5 eq, 5 mM), $MnCl_2$ (10 mM, from 1 M solution at 1% v/v of reaction volume), CIAP (1% v/v) and FUT3 (10% v/v) were added, after which the solution was incubated at 37 °C with gentle shaking. The reaction progress was monitored with LC-MS. A volume of 0.1 eq. of GDP-Fucose was added each hour until only limited amount of starting material was observed, after which the mixture was freeze dried. NMR after size exclusion chromatography revealed a ratio of 1:1 fucosylation of GlcNAc and glucose. FUT5 led to a ratio of 1:9 and FUT6 and FUT 9 were not able to fucosylate type 1 LacNAc. The mono-fucosylated isomers were purified by preparative LC to obtain **11** (yield: 1.2 mg, 25%, 1.4 µmol).

**$^{1}$H NMR (600 MHz, $D_2O$): δ (ppm).**

| | H1 | H2 | H3 | H4 | H5 | H6 | Ac |
|---|---|---|---|---|---|---|---|
| Glc(α) | 5.22 (d, J=3.78) | 3.58 | 3.84 | 3.65 | n/a | n/a | - |
| Glc(β) | 4.67 (d, 7.88) | 3.28 (app. t, J=8.48) | 3.64 | 3.95 | n/a | n/a | - |
| Gal | 4.44 (d, J=7.87) | 3.60 | 3.72 | 4.16 (d, J=3.33) | n/a | n/a | - |
| GlcNAc | 4.71 (d, J=8.54) | 3.95 | 4.09 (t, J=9.73) | 3.76 | 3.55 | 3.88 | 2.07 (3H) |
| Gal (2) | 4.51 (d, J=7.66) | 3.49 | 3.63 | 3.89 | n/a | n/a | - |
| Fuc | 5.03 (d, J=4.01) | 3.81 | 3.96 | 3.90 | 4.88 | (1.18, 3H) | - |

**$^{13}$C NMR derived from HSQC (150 MHz, $D_2O$): δ (ppm).**

| | C1 |
|---|---|
| Glc(α) | 91.74 |
| Glc(β) | 95.83 |
| Gal | 102.90 |
| GlcNAc | 102.63 |
| Gal (2) | 102.76 |
| Fuc | 97.92 |

ESI TOF-MS m/z calculated for $C_{32}H_{55}NO_{25}Na$ $(M + Na)^{+}$: 876.2955; measured: 876.2281.

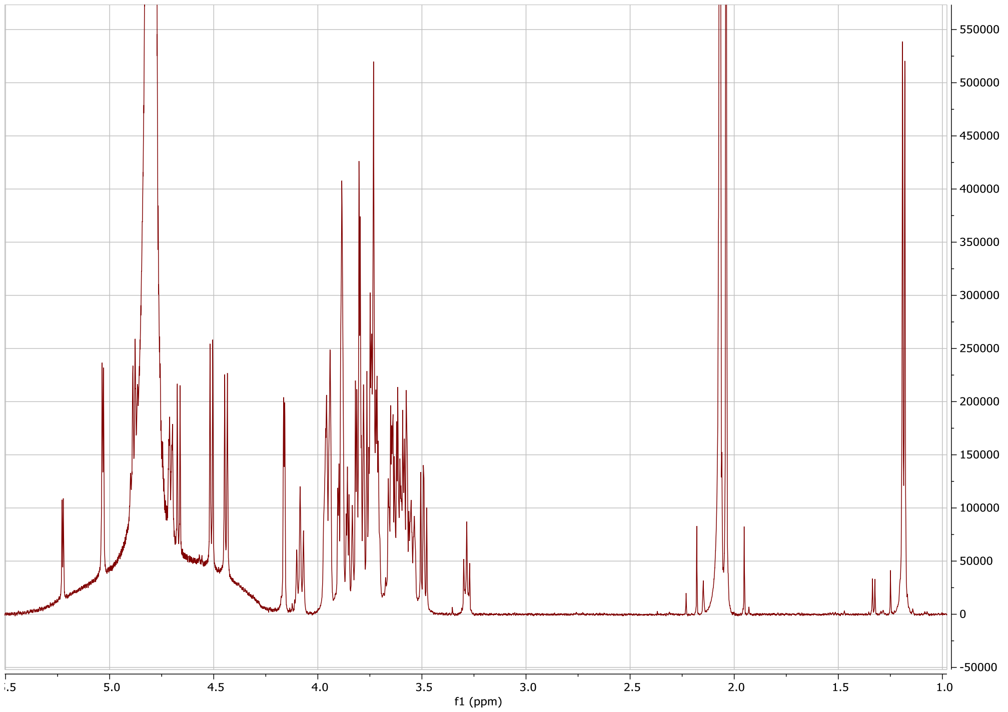

Proton spectrum of **11**.

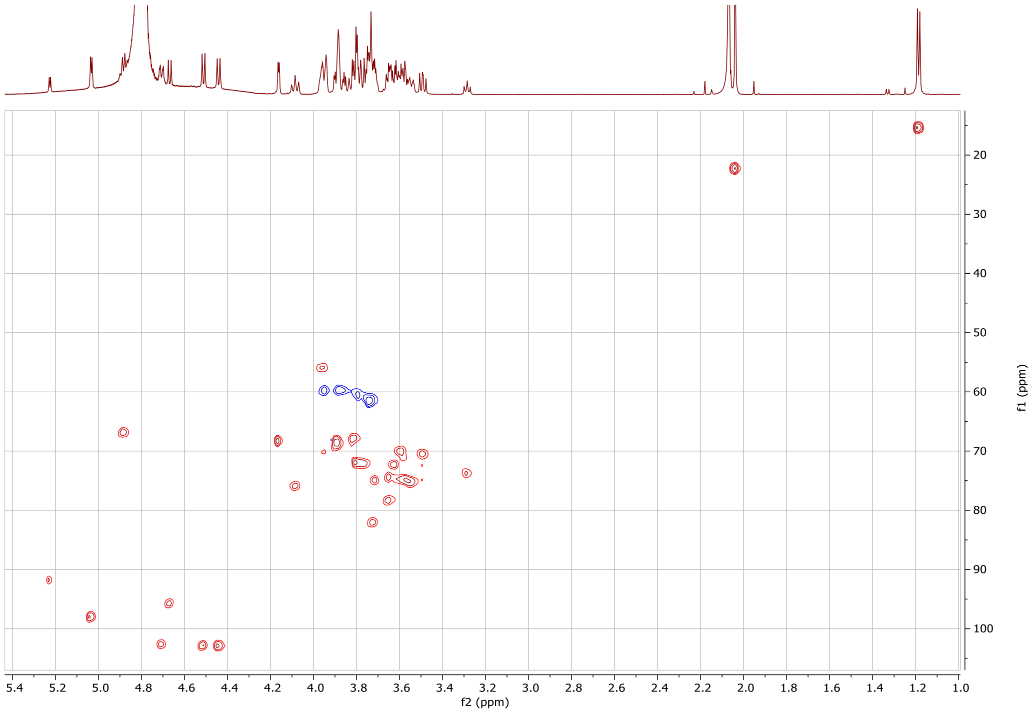

HSQC spectrum of **11**.

#### 1.12 Synthesis of H type 2 lacto-*N*-neopentose/LNnFPI (HMO12)

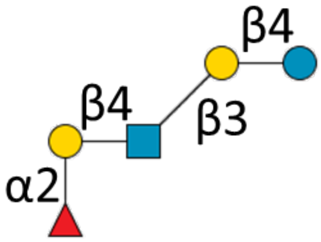

**12**

LNnT **1** (10 mg, 14.1 µmol) was dissolved in 50 mM TRIS (pH 7.5) to a concentration of 10 mM, and GDP-Fuc (1.5 eq, 15 mM), $MnCl_2$ (10 mM, from 1 M solution at 1% v/v of reaction volume), CIAP (1% v/v) and FUT1 (10% v/v) were added, after which the solution was incubated at 37 °C with gentle shaking. The reaction progress was monitored with LC-MS. The products were purified using size exclusion chromatography and preparative LC to obtain **12** (yield: 7 mg, 58%, 8.2 µmol).

**$^{1}$H NMR (600 MHz, $D_2O$): δ (ppm).**

| | H1 | H2 | H3 | H4 | H5 | H6 | Ac |
|---|---|---|---|---|---|---|---|
| Glc(α) | 5.23 (d, J=3.77) | 3.58 | 3.81 | 3.65 | n/a | n/a | - |
| Glc(β) | 4.67 (d, J=7.97) | 3.28 (app. t) | 3.64 | 3.96 | n/a | n/a | - |
| Gal | 4.45 (d, J=7.74) | 3.60 | 3.71 | 4.15 (d, J=3.34) | n/a | n/a | - |
| GlcNAc | 4.71 (d, J=8.39) | 3.82 | 3.47 | 3.97 | n/a | n/a | 2.06 (3H) |
| Gal (2) | 4.55 (d, J=7.79) | 3.68 | 3.88 | n/a (3.88) | n/a | n/a | - |
| Fuc | 5.31 (d, J=3.06) | 3.80 | 3.96 | 3.83 | 4.22 (q, J=6.65) | (1.23, 3H) | - |

**$^{13}$C NMR derived from HSQC (150 MHz, $D_2O$): δ (ppm).**

| | C1 |
|---|---|
| Glc(α) | 91.73 |
| Glc(β) | 95.73 |
| Gal | 102.88 |
| GlcNAc | 102.75 |
| Gal (2) | 100.23 |
| Fuc | 99.36 |

ESI TOF-MS m/z calculated for $C_{32}H_{55}NO_{25}Na$ $(M + Na)^{+}$: 876.2955; measured: 876.2881.

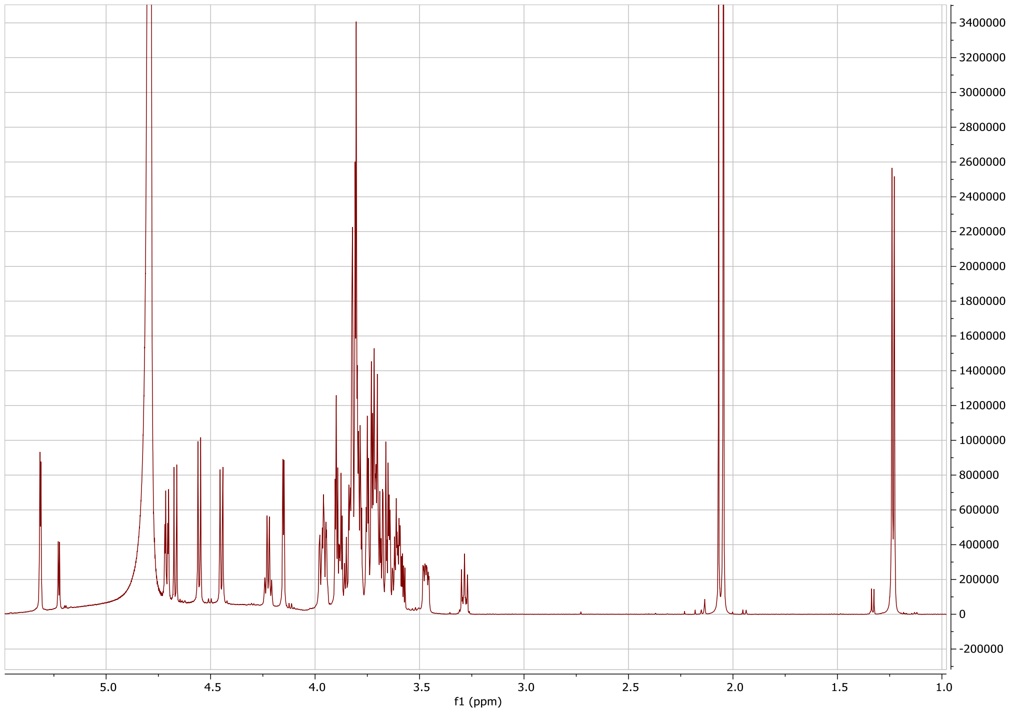

Proton spectrum of **12**.

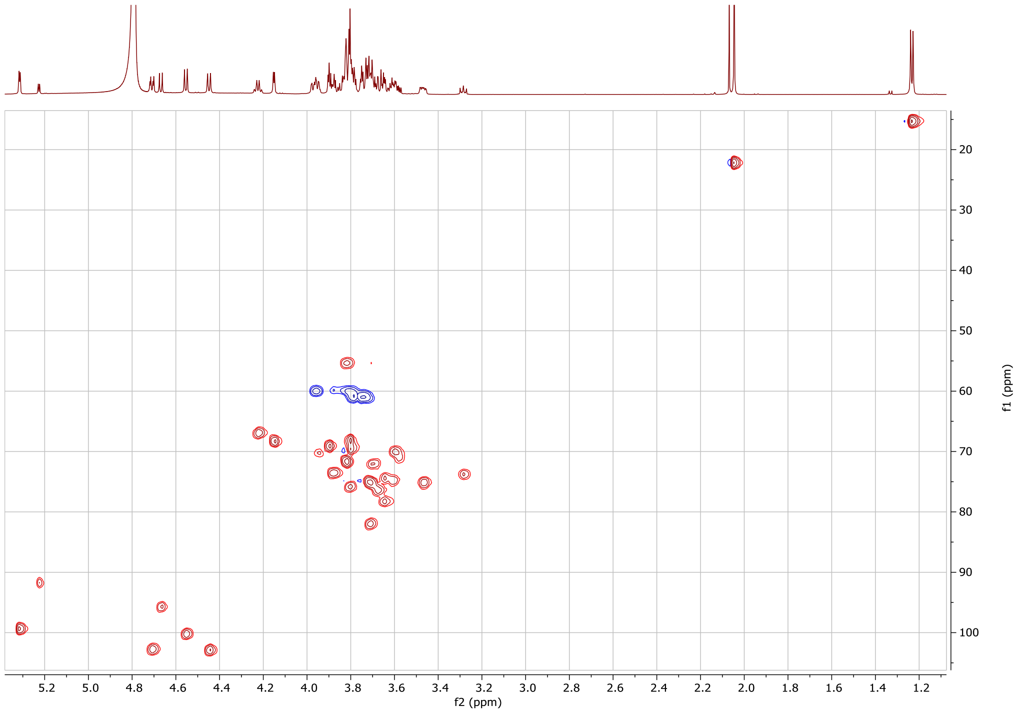

HSQC spectrum of **12**.

#### 1.13 Synthesis of H type 1 lacto-*N*-pentose/LNFPI (HMO13)

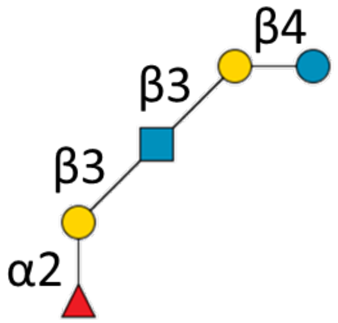

**13**

LNT **2** (7.5 mg, 10.6 μmol) was dissolved in 50 mM TRIS (pH 7.5) to a concentration of 10 mM, and GDP-Fuc (1.5 eq, 15 mM), $MnCl_2$ (10 mM, from 1 M solution at 1% v/v of reaction volume), CIAP (1% v/v) and FUT1 (10% v/v) were added, after which the solution was incubated at 37 °C with gentle shaking. The reaction progress was monitored with LC-MS. The products were purified using size exclusion chromatography and preparative LC to obtain **13** (yield: 8 mg, 89%, 9.4 μmol).

**$^{1}$H NMR (600 MHz, $D_2O$): δ (ppm).**

| | H1 | H2 | H3 | H4 | H5 | H6 | Ac |
|---|---|---|---|---|---|---|---|
| Glc(α) | 5.22 (d, J=3.80) | 3.58 | 3.83 | n/a | n/a | n/a | - |
| Glc(β) | 4.67 (d, J=7.74) | 3.28 | 3.63 | 3.95 | n/a | n/a | - |
| Gal | 4.43 (d, J=7.89) | 3.57 | 3.72 | 4.15 (d, J=3.32) | n/a | n/a | - |
| GlcNAc | 4.63 (d, J=8.41) | 3.80 | 3.51 | 3.55 | n/a | n/a | 2.06 (3H) |
| Gal (2) | 4.65 (d, J=7.84) | 3.60 | 3.84 | 3.89 | n/a | n/a | - |
| Fuc | 5.19 (d, J=4.11) | 3.77 | 3.67 | 3.74 | 4.30 (q, J=6.7) | 1.24($CH_3$) | - |

**$^{13}$C NMR derived from HSQC (150 MHz, $D_2O$): δ (ppm).**

| | C1 |
|---|---|
| Glc(α) | 91.76 |
| Glc(β) | 95.68 |
| Gal | 102.95 |
| GlcNAc | 103.23 |
| Gal (2) | 100.19 |
| Fuc | 99.45 |

ESI TOF-MS m/z calculated for $C_{32}H_{55}NO_{25}Na$ $(M + Na)^+$: 876.2955; measured: 876.2880.

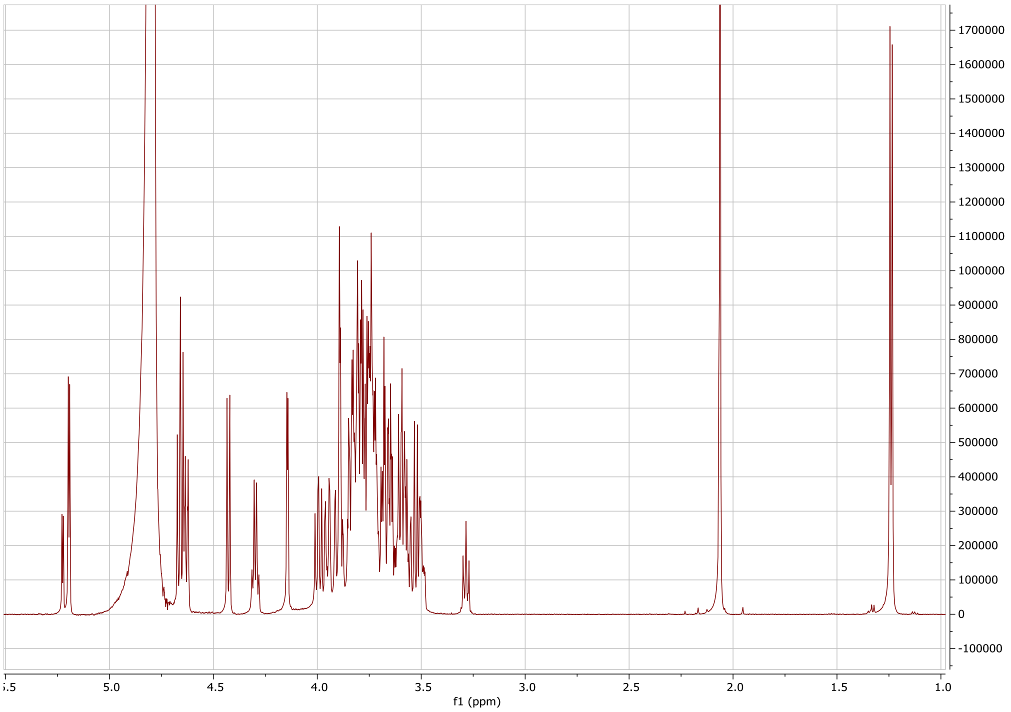

Proton spectrum of **13**.

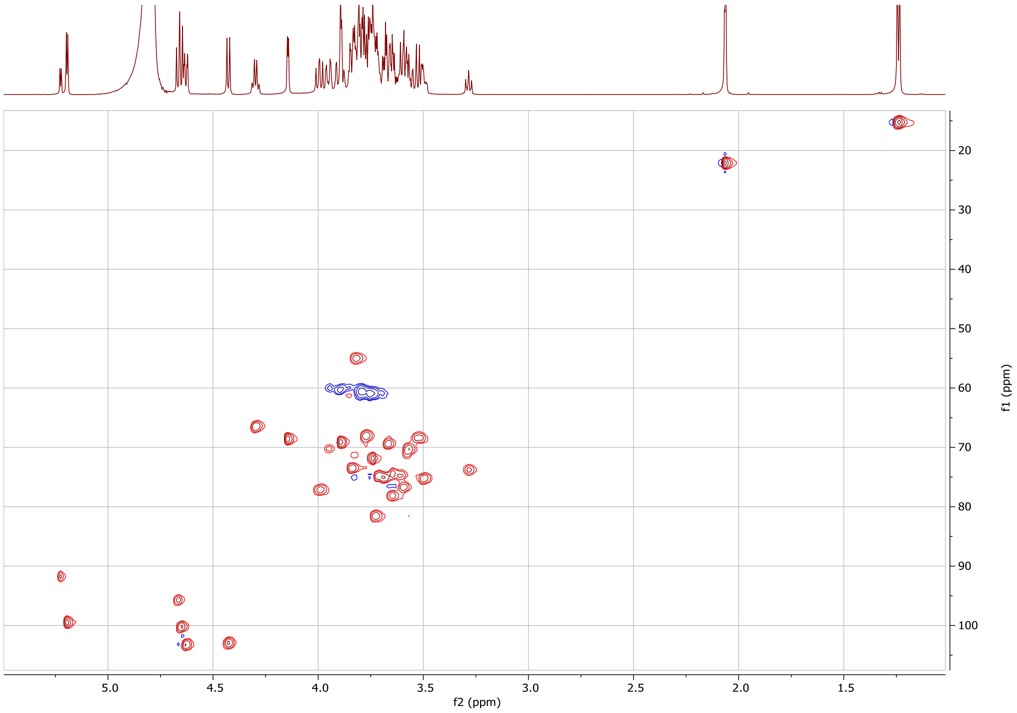

HSQC spectrum of **13**.

#### 1.14 Synthesis of lewis y lacto-*N*-neohexose/LNnDFHI (HMO14)

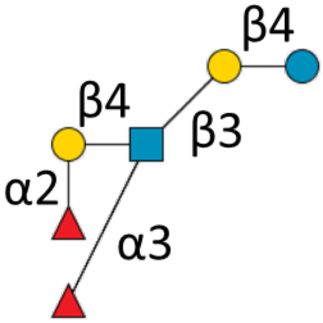

**14**

**12** (3 mg, 3.5 µmol) was dissolved in 100 mM TRIS (pH 7.5) to a concentration of 10 mM, and GDP-Fuc (0.5 eq, 5 mM), $MnCl_2$ (10 mM, from 1 M solution at 1% v/v of reaction volume), CIAP (1% v/v) and FUT9 (10% v/v) were added, after which the solution was incubated at 37 °C with gentle shaking. The reaction progress was monitored with LC-MS. 0.1 eq. of GDP-Fucose was added each hour until the reaction went to completion, after which the mixture was freeze dried. Size exclusion chromatography and preparative LC were used to obtain **14** (yield: 2.3 mg 66%, 2.3 µmol).

**$^{1}$H NMR (600 MHz, $D_2O$): δ (ppm).**

| | H1 | H2 | H3 | H4 | H5 | H6 | Ac |
|---|---|---|---|---|---|---|---|
| Glc(α) | 5.22 (d, J=3.76) | 3.58 | 3.85 | 3.65 | n/a | n/a | - |
| Glc(β) | 4.67 (d, J=8.02) | 3.28 | 3.65 | 3.61 | n/a | n/a | - |
| Gal | 4.45 (d, J=7.73) | 3.60 | 3.70 | 4.15 (d, J=3.30) | n/a | n/a | - |
| GlcNAc | 4.72 (d, J=8.55 | 3.96 | 3.46 | 3.87 | n/a | n/a | 2.03 (3H) |
| Gal (2) | 4.52 (d, J=7.82) | 3.66 | 3.86 | 3.86 | n/a | n/a | - |
| Fuc glcnac | 5.12 (d, J=3.99) | 3.70 | 3.93 | 3.81 | 4.89 (q, J=6.84/7.18) | (1.24, 3H) | - |
| Fuc(2) gal | 5.28 (d, J=3.45) | 3.80 | 3.77 | 3.83 | 4.62 (q, J=6.68/6.75) | (1.27, 3H) | - |

**$^{13}$C NMR derived from HSQC (150 MHz, $D_2O$): δ (ppm).**

| | C1 |
|---|---|
| Glc(α) | 91.75 |
| Glc(β) | 95.72 |
| Gal | 102.90 |
| GlcNAc | 102.49 |
| Gal (2) | 100.18 |
| Fuc | 98.56 |
| Fuc(2) | 99.37 |

ESI TOF-MS m/z calculated for $C_{38}H_{65}NO_{29}Na$ $(M + Na)^{+}$: 1022.3534; measured: 1022.3450.

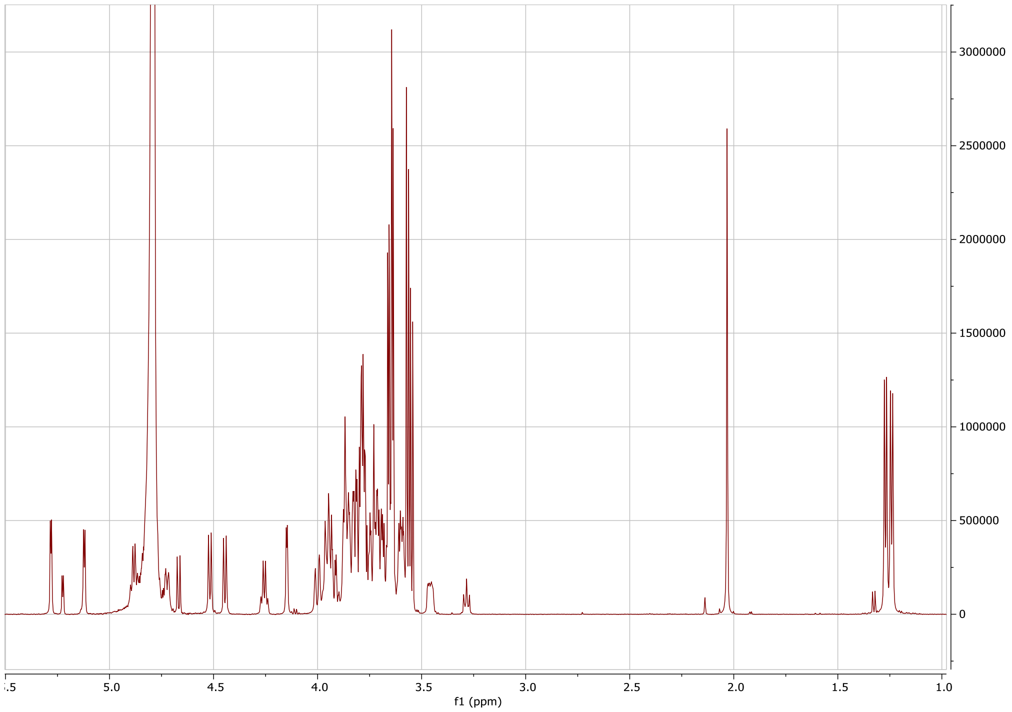

Proton spectrum of **14**.

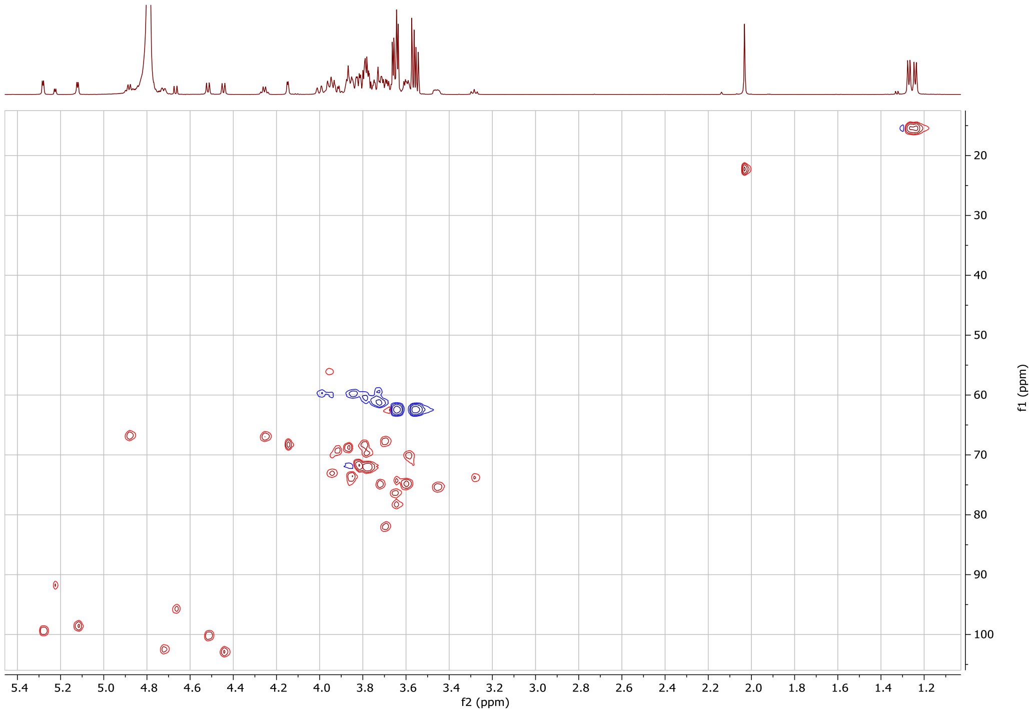

HSQC spectrum of **14**.

#### 1.15 Synthesis of lewis b lacto-*N*-hexose/LNDFHI (HMO15)

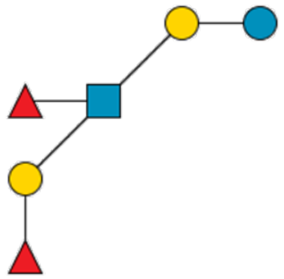

**15**

**13** (1.5 mg, 1.8 µmol) was dissolved in 100 mM TRIS (pH 7.5) to a concentration of 10 mM, and GDP-Fuc (0.5 eq, 5 mM), $MnCl_2$ (10 mM, from 1 M solution at 1% v/v of reaction volume), CIAP (1% v/v) and FUT3 (10% v/v) were added, after which the solution was incubated at 37 °C with gentle shaking. The reaction progress was monitored with LC-MS. A volume of 0.1 eq. of GDP-Fucose was added each hour until the reaction went to completion, after which the mixture was freeze dried. Size exclusion chromatography and preparative LC were used to obtain **15** (yield: 0.8 mg 44%, 0.8 µmol).

**$^1$H NMR (600 MHz, $D_2O$): δ (ppm).**

| | H1 | H2 | H3 | H4 | H5 | H6 | Ac |
|---|---|---|---|---|---|---|---|
| Glc(α) | 5.23 (d, J=3.75) | 3.56 | 3.82 | n/a | n/a | n/a | - |
| Glc(β) | 4.67 (2H, m) | 3.26 (app. t, J=8.46) | 3.65 | 3.95 | n/a | n/a | - |
| Gal | 4.42 (d, J=7.90) | 3.57 | 3.73 | 4.15 (m, 2H) | n/a | n/a | - |
| GlcNAc | 4.66 (2H, m) | 3.60 | n/a | n/a | n/a | n/a | 2.07 (3H) |
| Gal (2) | 4.61 (d, J=8.16) | 3.85 | 3.74 | 4.15 (m, 2H) | n/a | n/a | - |
| Fuc | 5.03 (d, J=3.94) | 3.81 | 3.93 | 3.83 | 4.87 | 1.26 (3H, d, J=6.52) | - |
| Fuc(2) | 5.16 (d, J=4.06) | 3.75 | 3.68 | 3.77 | 4.35 (q, J=6.65) | 1.28 (3H, d, J=6.77) | - |

**$^{13}$C NMR derived from HSQC (150 MHz, $D_2O$): δ (ppm).**

| | C1 |
|---|---|
| Glc(α) | 91.67 |
| Glc(β) | 95.72 |
| Gal | 102.95 |
| GlcNAc | 100.61 |
| Gal (2) | 103.27 |
| Fuc | 97.78 |
| Fuc(2) | 99.49 |

ESI TOF-MS m/z calculated for $C_{38}H_{65}NO_{29}Na$ $(M + Na)^+$: 1022.3534; measured: 1022.3456.

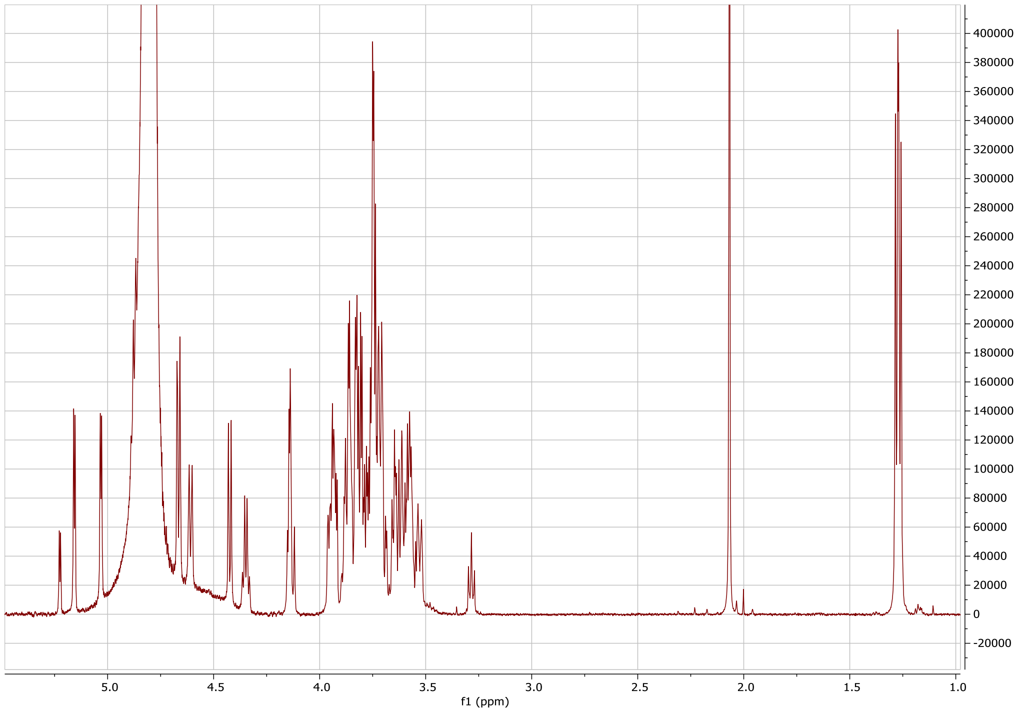

Proton spectrum of **15**.

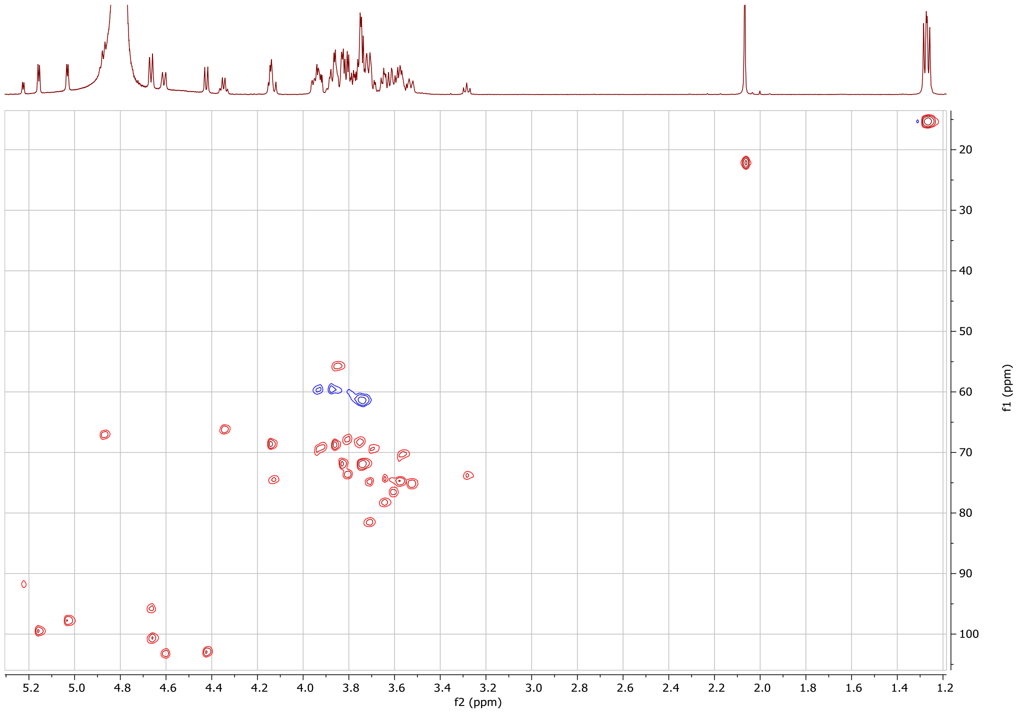

HSQC spectrum of **15**.

#### 1.16 Synthesis of A type 2 lacto-*N*-neohexose (HMO16)

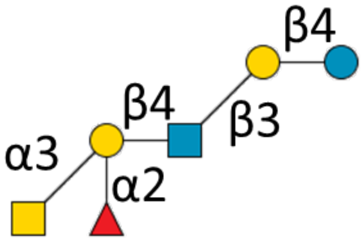

**16**

**12** (0.4 mg, 0.5 µmol) was dissolved in 100 mM TRIS (pH 7) to a concentration of 10 mM, and UDP-GalNAc (1.5 eq, 15 mM), CIAP (1% v/v) and Bo6GTA (5% v/v) were added, after which the solution was incubated at 37 °C with gentle shaking. The reaction process was monitored with LC-MS. Size exclusion chromatography and preparative HPLC were used to obtain **16** (yield: 0.2 mg , 40% , 0.2 µmol).

**$^{1}$H NMR (600 MHz, $D_2O$): δ (ppm).**

| | H1 | H2 | H3 | H4 | H5 | H6 | Ac |
|---|---|---|---|---|---|---|---|
| Glc(α) | 5.23 (d, J=3.80) | 3.58 | 3.83 | 3.66 | n/a | n/a | - |
| Glc(β) | 4.67 (d, J=7.98) | 3.29 (app. t, J=8.50) | 3.65 | 3.81 | n/a | n/a | - |
| Gal | 4.45 (d, J=7.81) | 3.60 | 3.68 | 4.15 (d, J=3.31) | n/a | n/a | - |
| GlcNAc | 4.71 (d, J=8.32) | 3.80 | 3.45 | 3.68 | n/a | n/a | 2.05 (6H) |
| Gal (2) | 4.62 (d, J=7.67) | 3.91 | 3.76 | 4.22 | n/a | n/a | - |
| Fuc | 5.36 (d, J=4.13) | 3.79 | 3.72 | 3.83 | 4.32 (q, J=6.71) | 1.25 (d, J=6.63) | - |
| GalNAc | 5.18 (d, J=3.87) | 4.24 | 3.90 | 4.00 | n/a | n/a | 2.05 (6H) |

**$^{13}$C NMR derived from HSQC (150 MHz, $D_2O$): δ (ppm).**

| | C1 |
|---|---|
| Glc(α) | 91.95 |
| Glc(β) | 95.74 |
| Gal | 102.96 |
| GlcNAc | 102.68 |
| Gal (2) | 99.99 |
| Fuc | 98.61 |
| GalNAc | 91.22 |

ESI TOF-MS m/z calculated for $C_{40}H_{68}N_2O_{30}Na$ $(M + Na)^+$: 1079.3749; measured: 1079.3710.

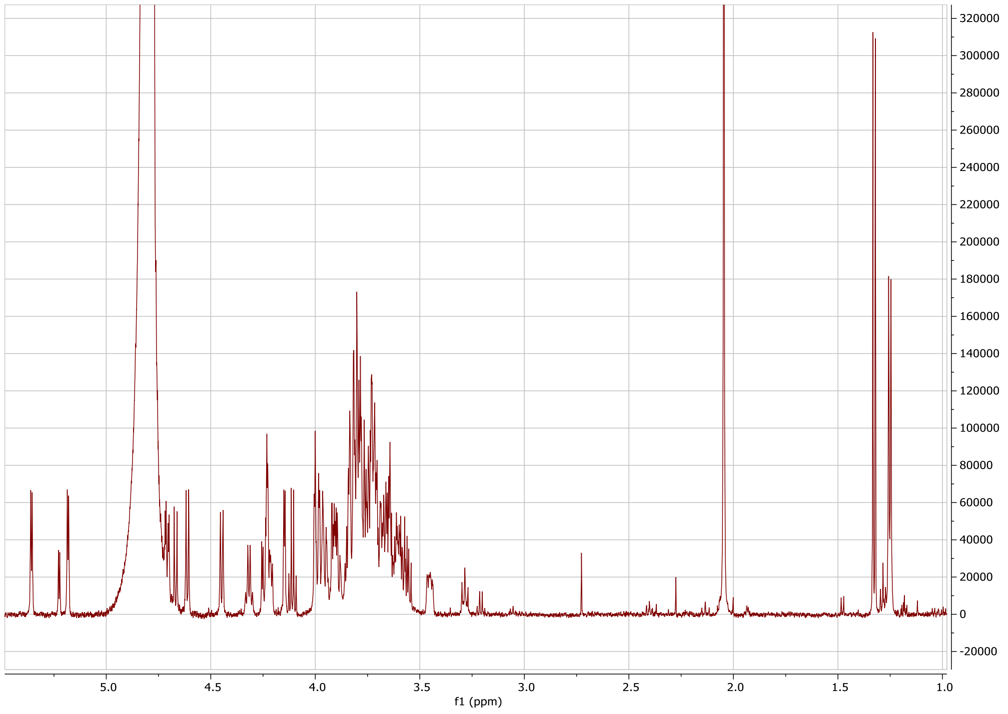

Proton spectrum of **16**.

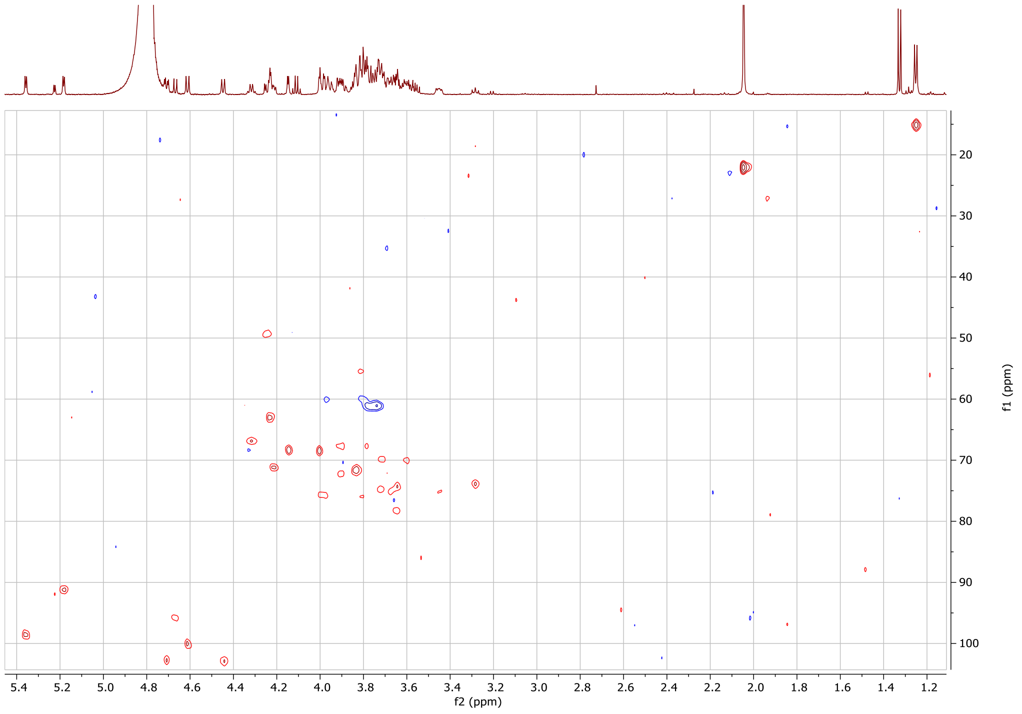

HSQC spectrum of **16**.

#### 1.17 Synthesis of B type 2 lacto-*N*-neohexose (HMO17)

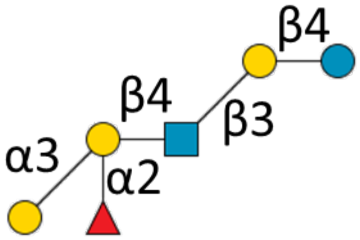

**17**

**12** (0.7 mg, 0.8 µmol) was dissolved in 100 mM TRIS (pH 7) to a concentration of 10 mM, and UDP-GalNAc (1.5 eq, 15 mM), CIAP (1% v/v) and GTB (50% v/v) were added, after which the solution was incubated at 37 °C with gentle shaking. The reaction process was monitored with LC-MS. Size exclusion chromatography and preparative LC were used to obtain **17** (yield: 0.2 mg , 23% , 0.2 µmol).

**$^{1}$H NMR (600 MHz, $D_2O$): δ (ppm).**

| | H1 | H2 | H3 | H4 | H5 | H6 | Ac |
|---|---|---|---|---|---|---|---|
| Glc(α) | 5.23 (d, J=3.93) | 3.59 (app. t, J=8.47) | 3.83 | 3.66 | n/a | n/a | - |
| Glc(β) | 4.67 (d, J=8.02) | 3.29 | 3.65 | 3.81 | n/a | n/a | - |
| Gal | 4.45 (d, J=7.86) | 3.61 | 3.72 | 4.15 (d, J=3.31) | n/a | n/a | - |
| GlcNAc | 4.71 (d, J=7.81) | 3.82 | 3.45 | 3.70 | 3.98 | n/a | 2.05 (3H) |
| Gal (2) | 4.64 (d, J=7.65) | 3.93 | 4.30 | 4.01 | n/a | n/a | - |
| Fuc | 5.35 (d, J=4.12) | 3.79 | 3.73 | 3.82 | 4.31 | 1.24 (d, J=6.64) | - |
| Gal (3) | 5.26 (app. s) | 3.89 | 3.99 | 4.20 (t, J=6.20) | 3.75 | n/a | - |

**$^{13}$C NMR derived from HSQC (150 MHz, $D_2O$): δ (ppm).**

| | C1 |
|---|---|
| Glc(α) | 91.87 |
| Glc(β) | 95.64 |
| Gal | 102.81 |
| GlcNAc | 100.10 |
| Gal (2) | 102.75 |
| Fuc | 98.67 |
| Gal (3) | 92.92 |

ESI TOF-MS m/z calculated for $C_{38}H_{65}NO_{30}Na$ $(M + Na)^{+}$: 1038.3484; measured: 1038.3449.

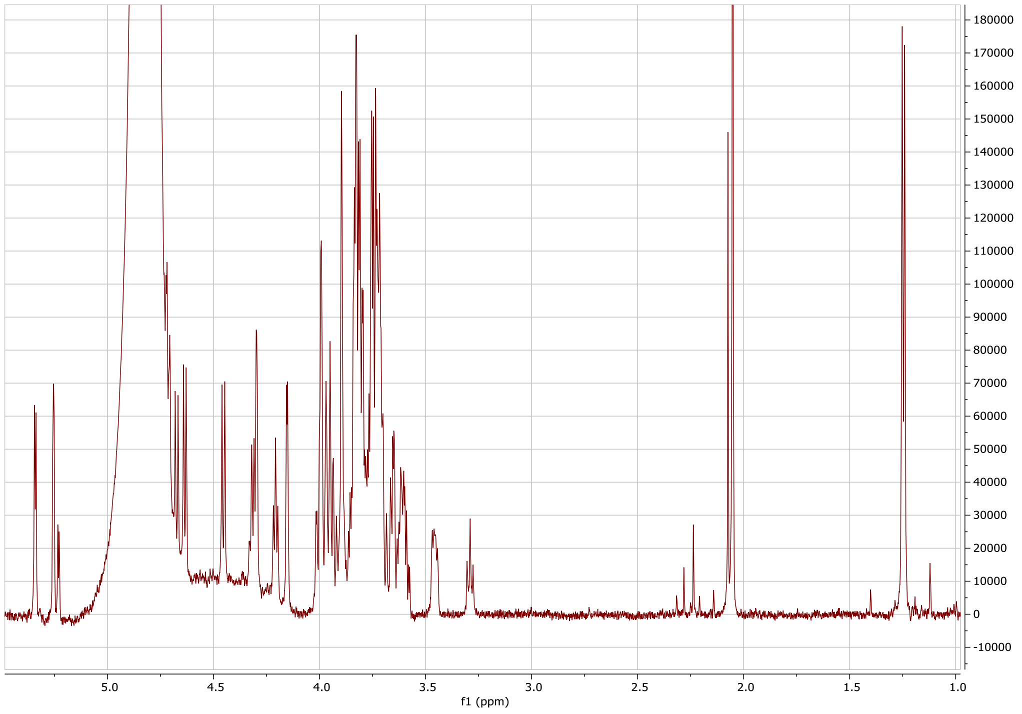

Proton spectrum of **17**.

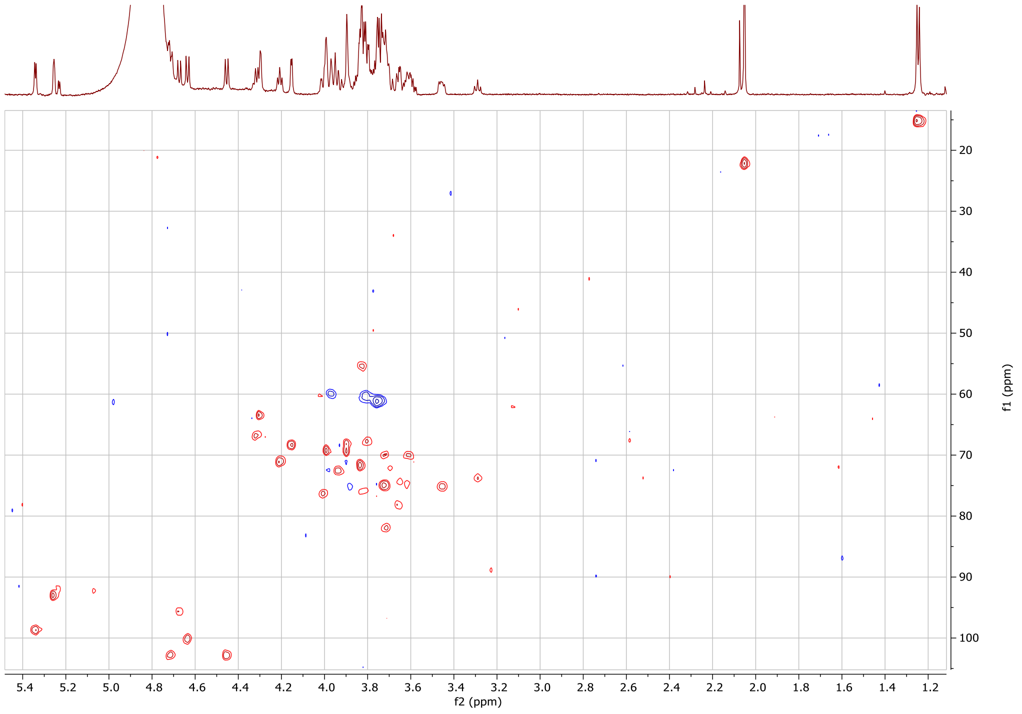

HSQC spectrum of **17**.

#### 1.18 Synthesis of A type 1 lacto-*N*-hexose (HMO18)

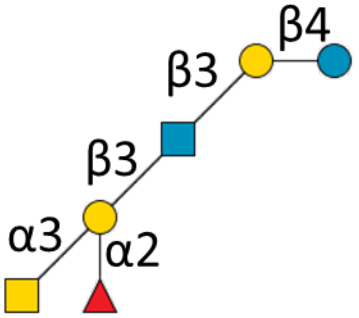

**18**

**13** (0.25 mg, 0.3 µmol) was dissolved in 100 mM TRIS (pH 7) to a concentration of 10 mM, and UDP-GalNAc (1.5 eq, 15 mM), CIAP (1% v/v) and Bo6GTA (5% v/v) were added, after which the solution was incubated at 37 °C with gentle shaking. The reaction process was monitored with LC-MS. Size exclusion chromatography and preparative LC were used to obtain **18** (yield: 0.2 mg , 65% , 0.2 µmol).

**$^{1}$H NMR (600 MHz, $D_2O$): δ (ppm).**

| | H1 | H2 | H3 | H4 | H5 | H6 | Ac |
|---|---|---|---|---|---|---|---|
| Glc(α) | 5.23 (d, J=3.77) | 3.58 | 3.85 | 3.64 | n/a | n/a | - |
| Glc(β) | 4.67 (d, J=8.00) | 3.28 (app. t) | 3.65 | 3.96 | n/a | n/a | - |
| Gal | 4.43 (d, J=7.89) | 3.57 | 3.72 | 4.15 (d, J=3.33) | n/a | n/a | - |
| GlcNAc | 4.63 (d, J=8.46) | 3.83 | 4.02 | 3.54 | n/a | n/a | 2.06 (6H) |
| Gal (2) | 4.71 (d, J=7.67) | 3.82 | 3.95 | 4.24 | n/a | n/a | - |
| Fuc | 5.26 (d, J=4.25) | 3.76 (2H) | 3.63 | 3.76 (2H) | 4.36 | 1.25 (3H) (d, J=6.54) | - |
| GalNAc | 5.19 (d, J=3.83) | 4.23 | 3.97 (2H) | 3.97 (2H) | n/a | n/a | 2.06 (6H) |

**$^{13}$C NMR derived from HSQC (150 MHz, $D_2O$): δ (ppm).**

| | C1 |
|---|---|
| Glc(α) | 91.91 |
| Glc(β) | 95.61 |
| Gal | 103.00 |
| GlcNAc | 103.22 |
| Gal (2) | 100.09 |
| Fuc | 99.01 |
| GalNAc | 91.10 |

ESI TOF-MS m/z calculated for $C_{40}H_{68}N_2O_{30}Na$ $(M + Na)^+$: 1079.3749; measured: 1079.3704.

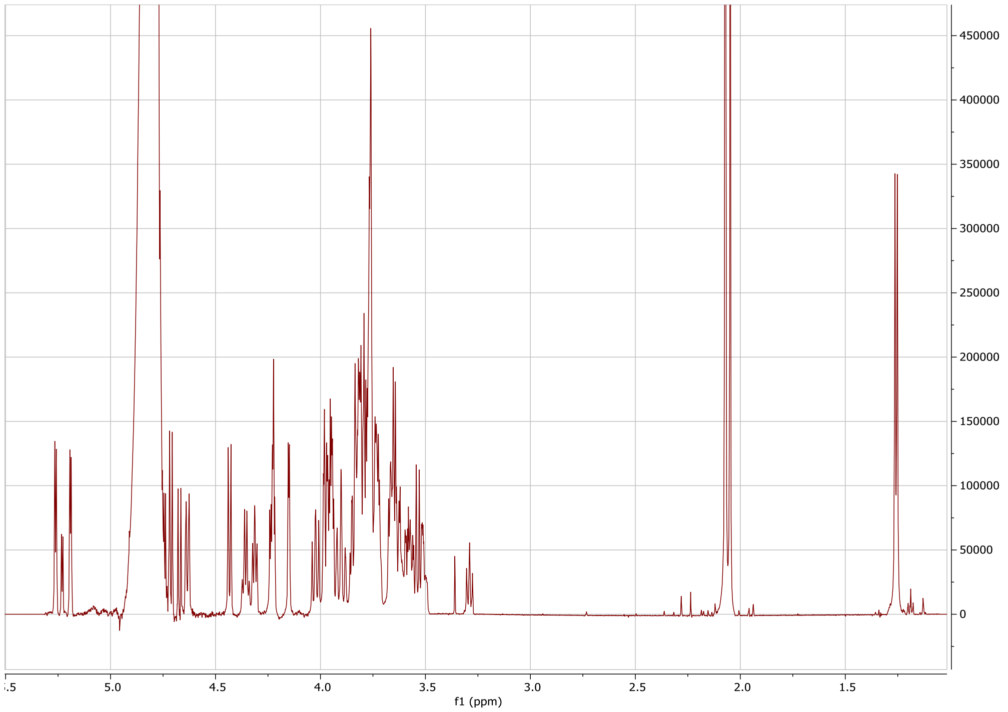

Proton spectrum of **18**

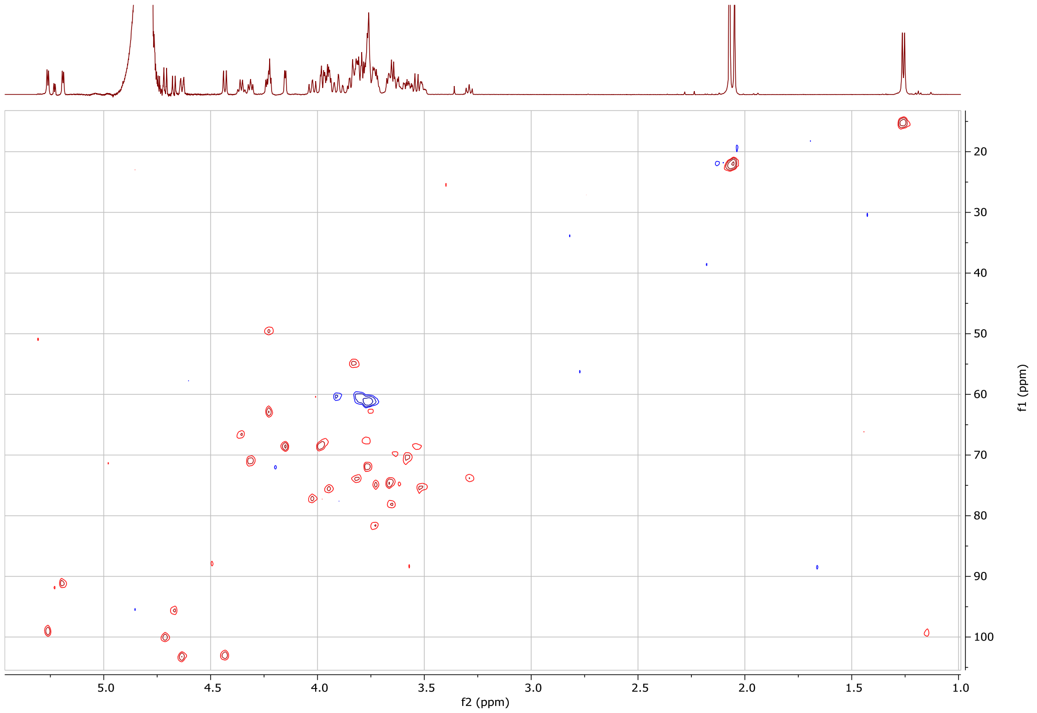

HSQC spectrum of **18**.

#### 1.19 Synthesis of B type 1 lacto-N-hexose (HMO19)

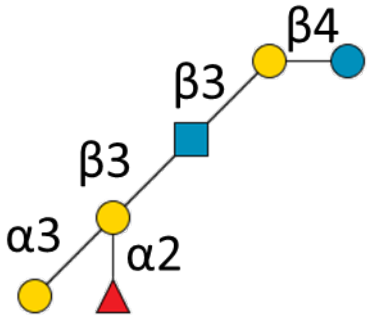

**19**

**13** (0.25 mg, 0.3 µmol) was dissolved in 100 mM TRIS (pH 7) to a concentration of 10 mM, and UDP-GalNAc (1.5 eq, 15 mM), CIAP (1% v/v) and GTB (50% v/v) were added, after which the solution was incubated at 37 °C with gentle shaking. The reaction process was monitored with LC-MS. Size exclusion chromatography and preparative LC were used to obtain **19** (yield: 0.1 mg , 31% , 0.2 µmol).

**$^1$H NMR (600 MHz, $D_2O$): δ (ppm).**

| | H1 | H2 | H3 | H4 | H5 | H6 | Ac |
|---|---|---|---|---|---|---|---|
| Glc(α) | 5.22 (q, 3H, J=4.21-5.30) | 3.60 (app. t, J=8.27) | 3.83 | n/a | n/a | n/a | - |
| Glc(β) | 4.67 (d, J=8.11) | 3.29 | 3.65 | 3.81 | n/a | n/a | - |
| Gal | 4.43 (d, J=7.91) | 3.57 | 3.73 | 4.15 (d, J=3.32) | n/a | n/a | - |
| GlcNAc | 4.64 (d, J=8.27) | 3.83 | 4.02 (app. t, J=9.5) | 3.53 | n/a | n/a | 2.07 (3H) |
| Gal (2) | 4.73 (d, J=7.72) | 3.84 | 3.96 | 4.29 | n/a | n/a | - |
| Fuc | 5.23 (q, 3H, J=4.21-5.30) | 3.77 | 3.63 | 3.75 | 4.36 (q, J=6.72) | 1.25 (3H, d, J=6.61) | - |
| Gal (3) | 5.24 (q, 3H, J=4.21-5.30) | 3.89 | 3.95 | 4.28 | 3.74 | n/a | - |

**$^{13}$C NMR derived from HSQC (150 MHz, $D_2O$): δ (ppm).**

| | C1 |
|---|---|
| Glc(α) | 92.09 |
| Glc(β) | 95.64 |
| Gal | 102.91 |
| GlcNAc | 103.31 |
| Gal (2) | 100.16 |
| Fuc | 99.26 |
| Gal (3) | 92.95 |

ESI TOF-MS m/z calculated for $C_{38}H_{65}NO_{30}Na$ (M + Na)$^+$: 1038.3484; measured: 1038.3446.

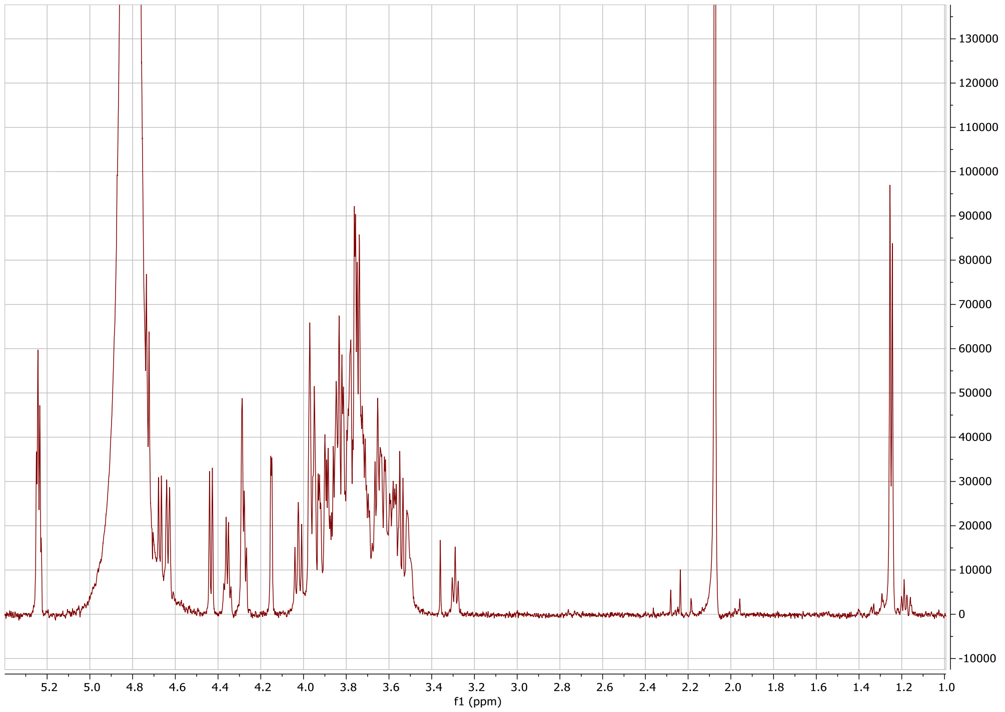

Proton spectrum of **19**.

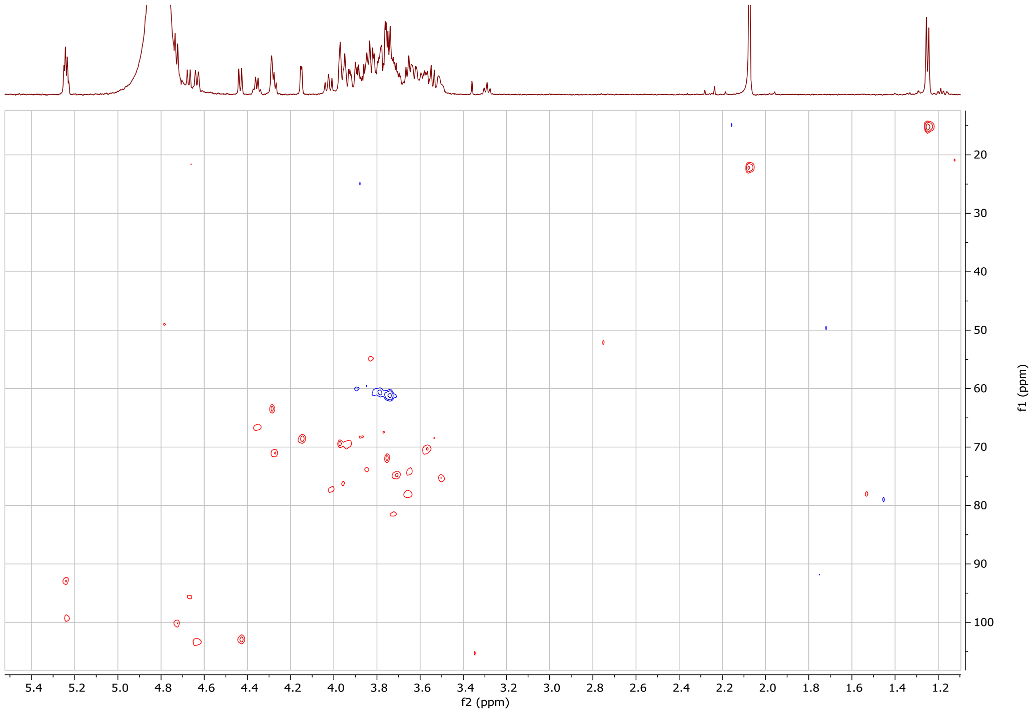

HSQC spectrum of **19**.

#### 1.20 Synthesis of B1,6-Glc-NAc i-branched LNT (HMO20)

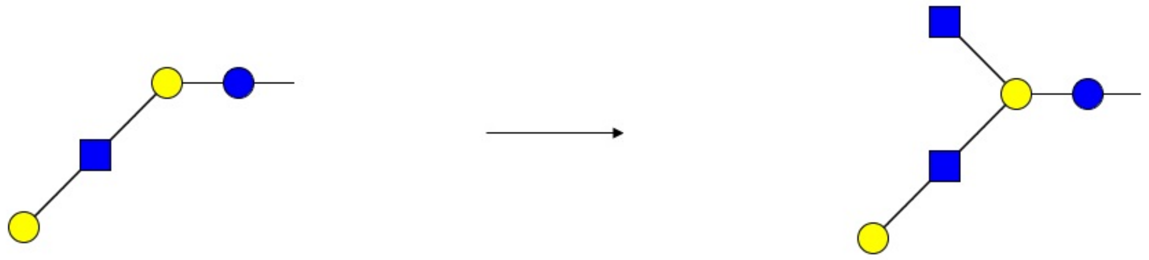

**20**

Amounts of 2.93 µmol **2** and 3.22 µmol UDP-GlcNAc were dissolved in 250 µL 50 mM Lgtb buffer. A volume of 6 µL 10% BSA, 6 µL 1U/µL CIAP, 6 µL 1 M $MnCl_2$, 6 µL 1M $MgCl_2$ and 25 µL GCNT2 were added and the reaction mixture was incubated at 37 °C under gentle shaking. The reaction was monitored using positive mode LC-MS. Over the course of 2 days, 50 µL GCNT2, 250 µL GCNT2 buffer and 25 µL 1 M $Na_2EDTA$ were added. The final estimated conversion after 7 days was approximately 10-20%. The reaction mixture was freeze-dried and then loaded onto a Biogel P2 column using 50 mM $NH_4HCO_3$ as eluent. Fractions containing product were pooled, freeze-dried, purified by preparative LC and freeze dried (yield 0.5 mg, 0.6 µmol.

#### 1.21 Synthesis of UDP-UL-$^{13}$C-galactose (HMO21)

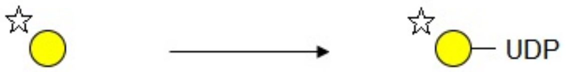

**21**

UL-$^{13}$C-galactose ($^{13}$C-Gal, 53.73 µmol) was dissolved in 200 µL TRIS-HCl buffer. Amounts of 64.48 µmol ATP, 64.48 µmol UTP and 10 mM $MgCl_2$ were added. The pH of the solution was adjusted to 7.5 before addition of the enzymes BiGalK (2.5% wt/wt), BLUSP (2.5% wt/wt), and inorganic phosphatase (2.5% wt/wt). The mixture was incubation at 37 °C under gentle shaking. The reaction was monitored using TLC with ceric molybdate staining until no starting material was visible anymore. The reaction mixture was freeze-dried, then purified using column chromatography. Fractions were checked using negative mode LC-MS. Fractions containing product were pooled and freeze-dried, then further purified using preparative LC and freeze-dried (yield: 2.1 mg, 3.6 µmol).

#### 1.22 Synthesis of 13C-Lacto-N-hexaose LNnH (HMO22)

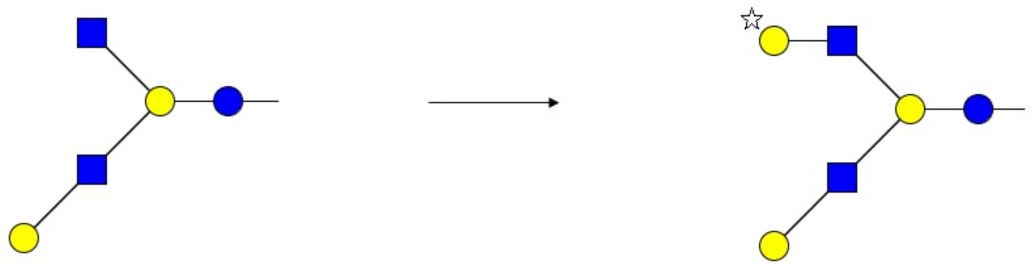

**22**

To a solution of 0.5 mg (0.6 µmol) **20** in 200 µL, 50 mM Lgtb buffer, 0.1 mg **21** (UDP-$^{13}$C-Gal), 1% BSA, 10 mM $MgCl_2$, 10 mM $MnCl_2$ and 2U CIAP were added. The reaction was incubated at 37 °C under gentle shaking and monitored using positive mode LC-MS. More UDP-$^{13}$C-Gal was added until no starting material was detected anymore. The reaction mixture was directly applied to a P6 column and the fractions containing product were combined and then freeze-dried (yield: weight too low to determine).

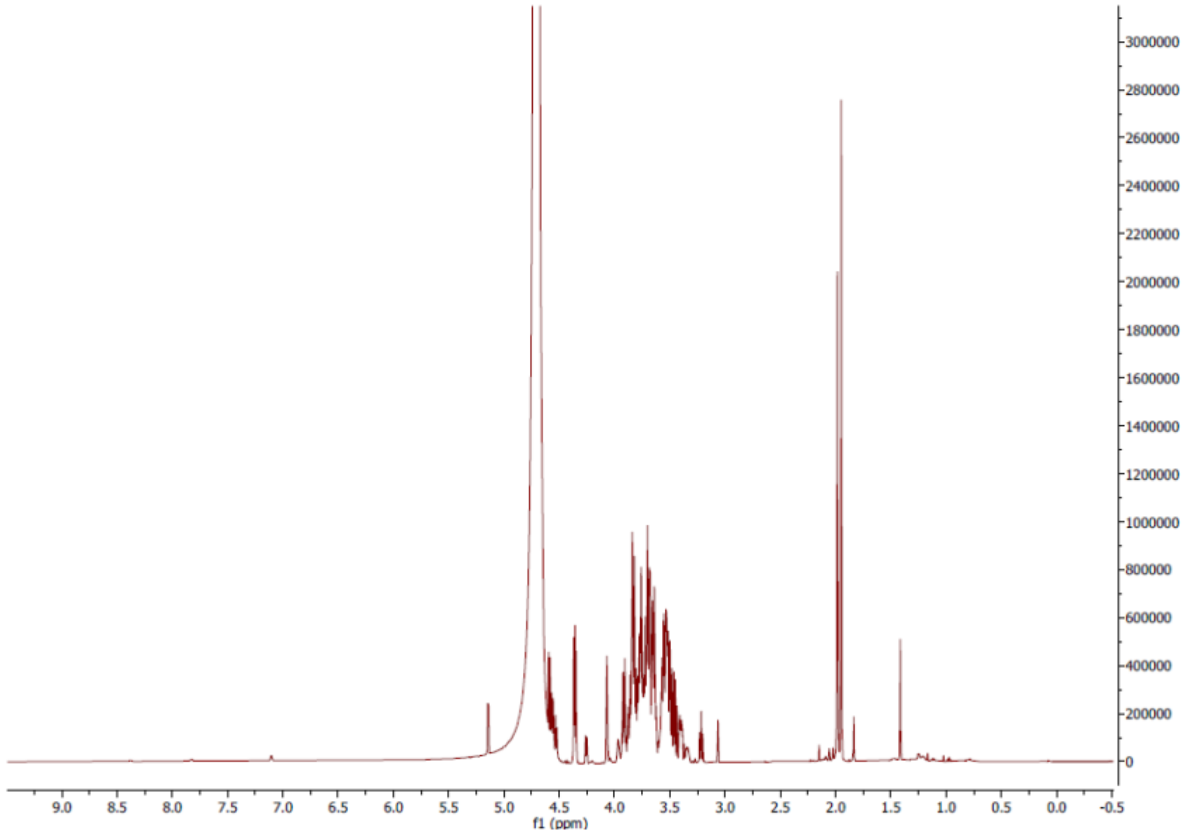

Proton spectrum of **22**.

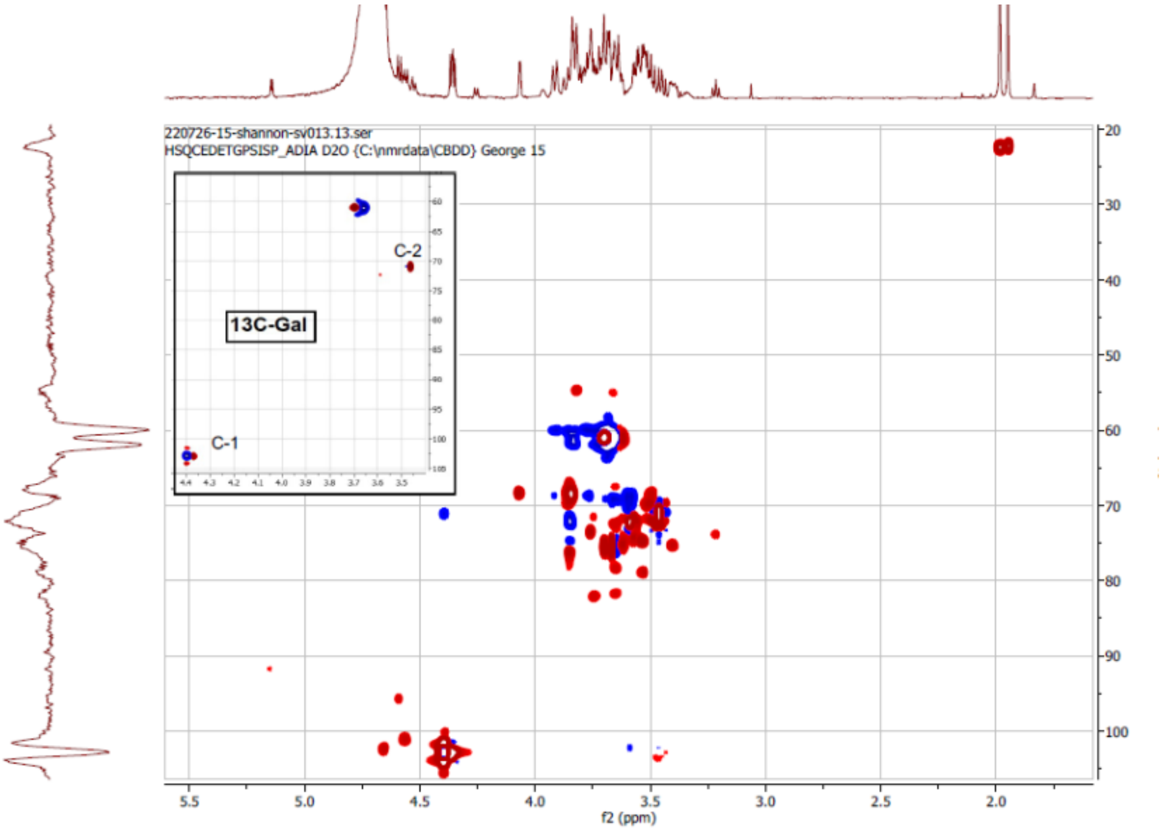

HSQC spectrum of **22**.

### 2. Analysis results of HMOs in breast milk.

#### 2.1 Acidic Fraction from Donor 1.

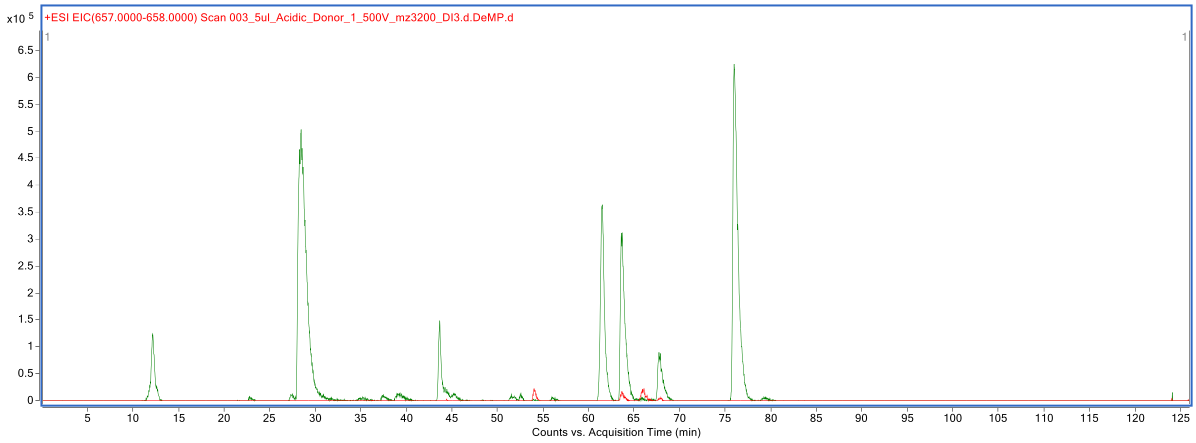

**Supplementary Figure 2. Extracted ion chromatogram at *m/z* 657-658 & and *m/z* 562-563 of HMOs labeled with procainamide.**

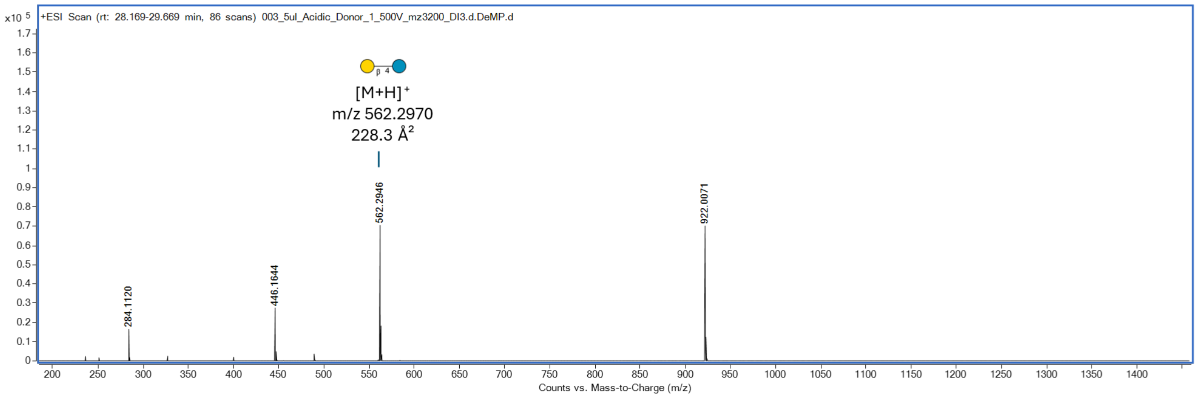

**Supplementary Figure 3. MS spectrum showing an identified HMO structure eluting at 28.5 min.**

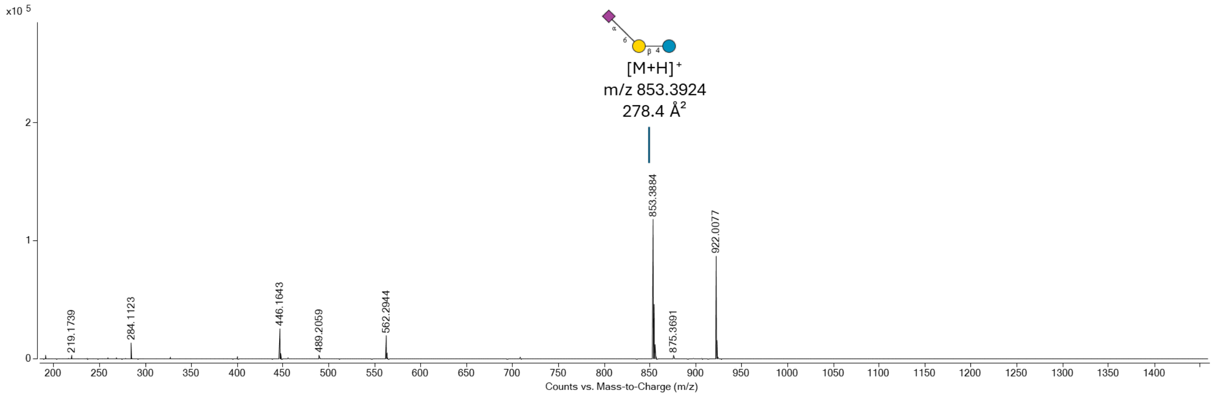

**Supplementary Figure 4. MS spectrum showing an identified HMO structure eluting at 43.7 min.**

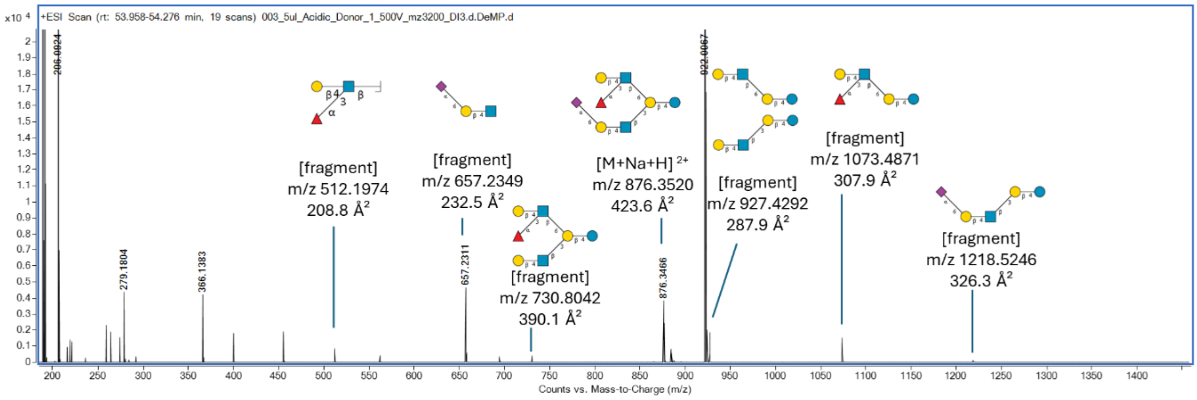

**Supplementary Figure 5. MS spectrum showing an identified HMO structure and its fragment ions eluting at 54.0 min.**

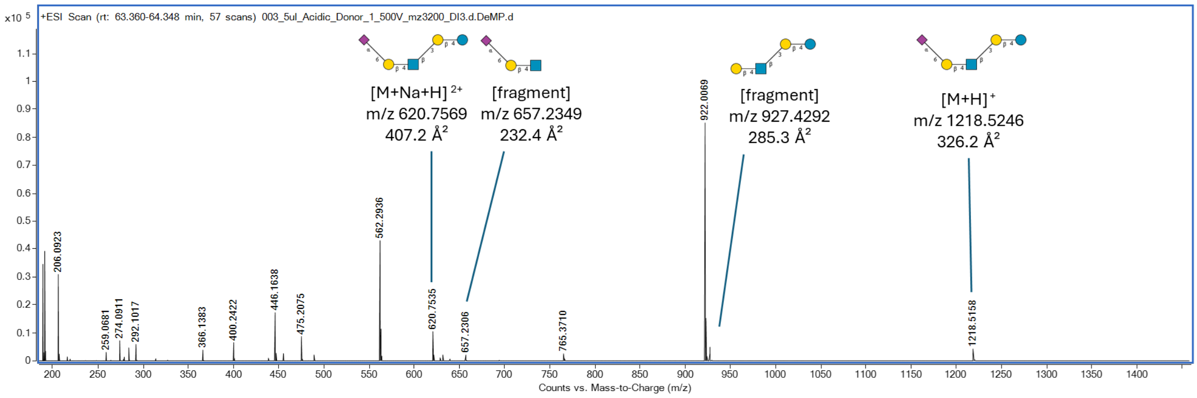

**Supplementary Figure 6. MS spectrum showing an identified HMO structure and its fragment ions eluting at 63.7 min.**

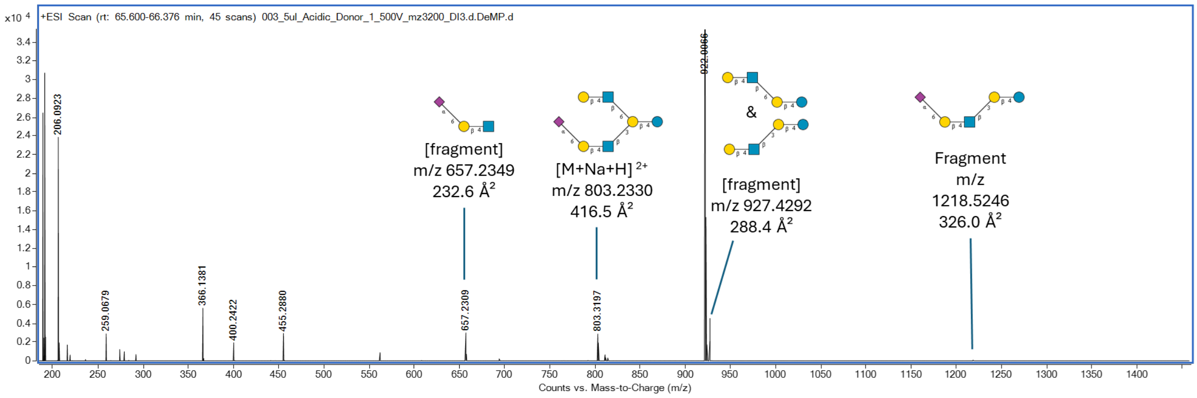

**Supplementary Figure 7. MS spectrum showing an identified HMO structure and its fragment ions eluting at 66.1 min.**

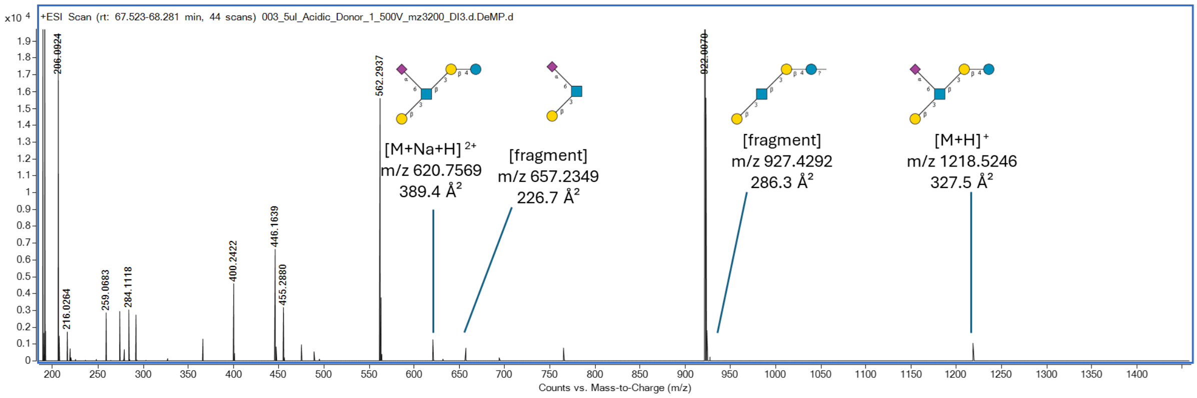

**Supplementary Figure 8. MS spectrum showing an identified HMO structure and its fragment ions eluting at 67.7 min.**

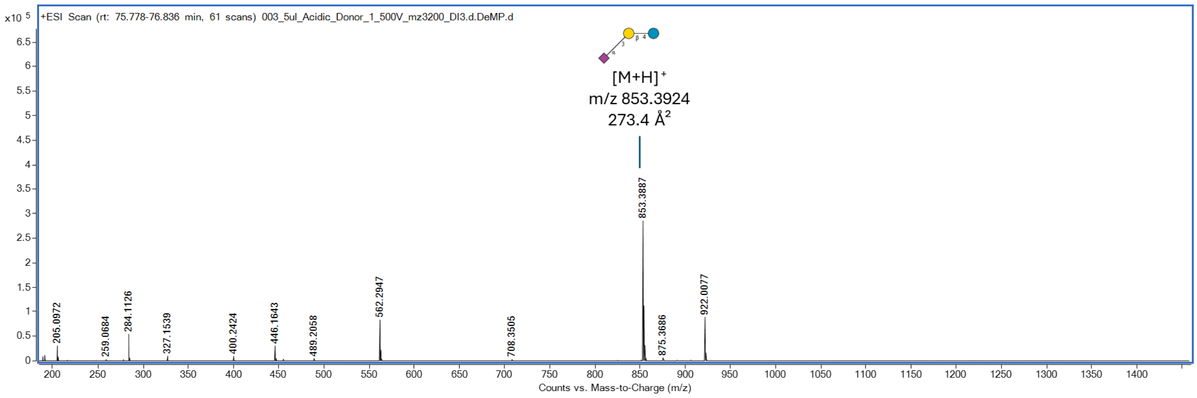

**Supplementary Figure 9. MS spectrum showing an identified HMO structure and its fragment ions eluting at 76.0 min.**

#### 2.2 Acidic Fraction from Donor 2.

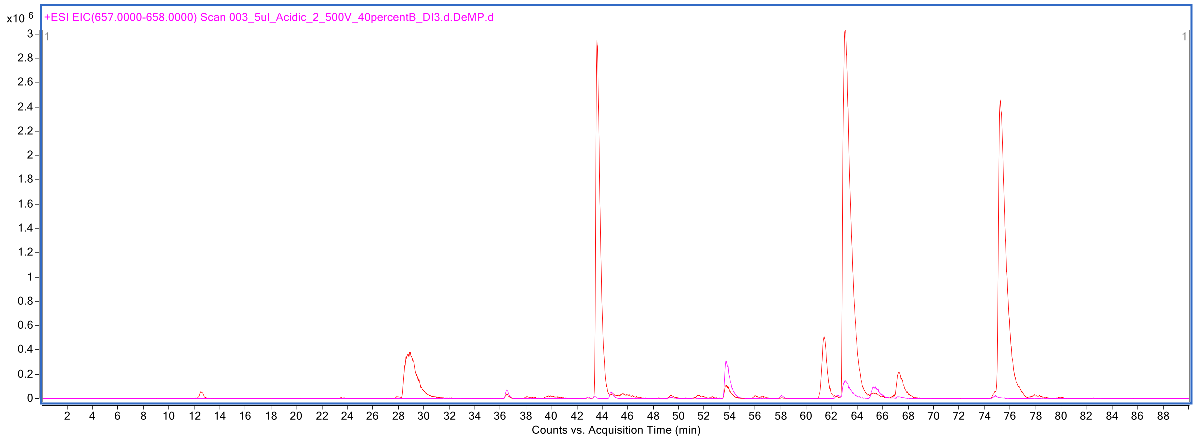

**Supplementary Figure 10. Extracted ion chromatogram at *m/z* 657-658 & and *m/z* 562-563 of HMOs labeled with procainamide.**

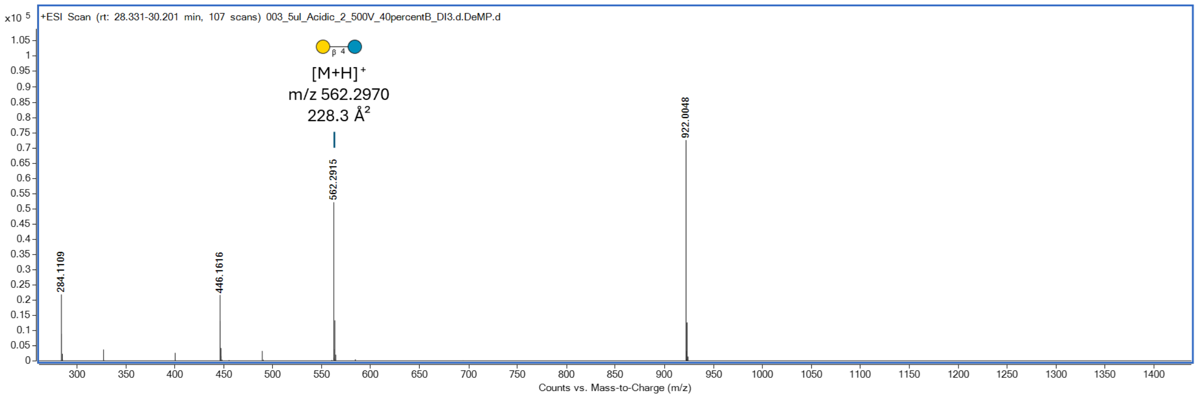

**Supplementary Figure 11. MS spectrum showing an identified HMO structure and its fragment ions eluting at 28.9 min.**

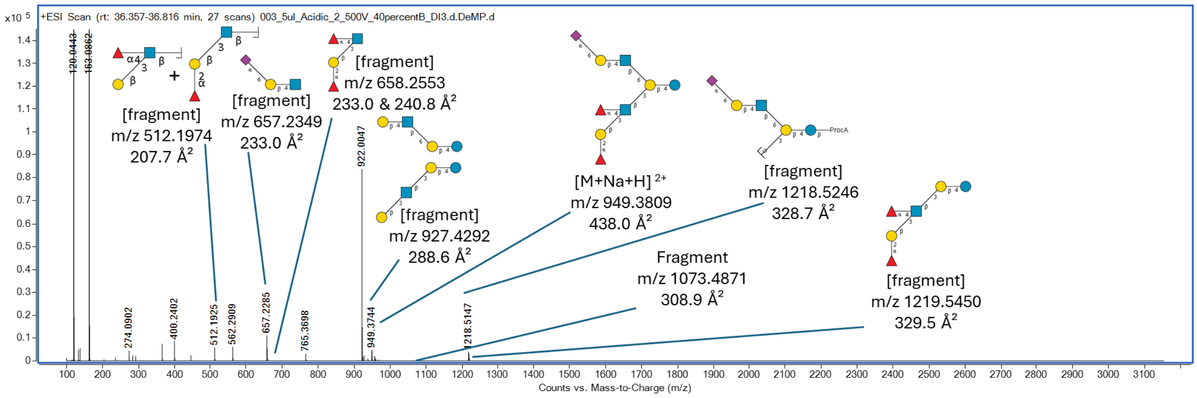

**Supplementary Figure 12. MS spectrum showing an identified HMO structure and its fragment ions eluting at 36.5 min.**

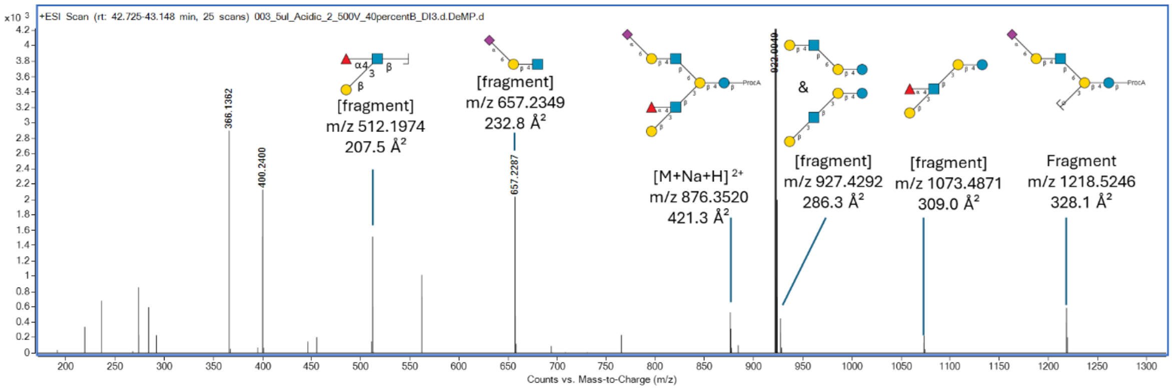

**Supplementary Figure 13. MS spectrum showing an identified HMO structure and its fragment ions eluting at 42.9 min.**

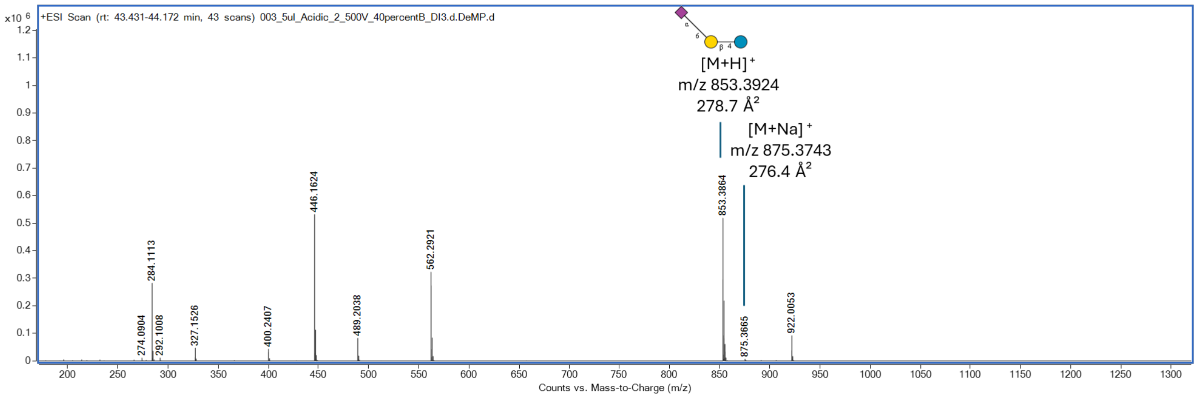

**Supplementary Figure 14. MS spectrum showing an identified HMO structure and its fragment ions eluting at 43.6 min.**

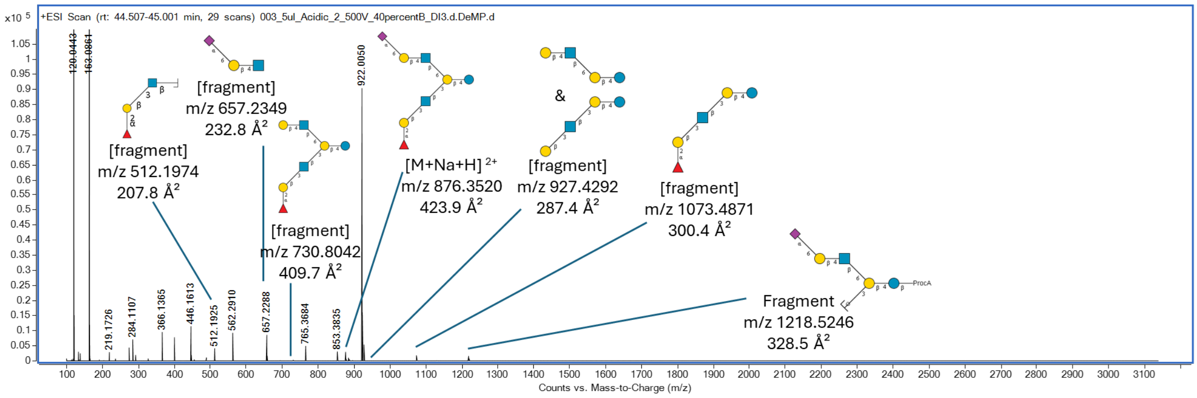

**Supplementary Figure 15. MS spectrum showing an identified HMO structure and its fragment ions eluting at 44.7 min.**

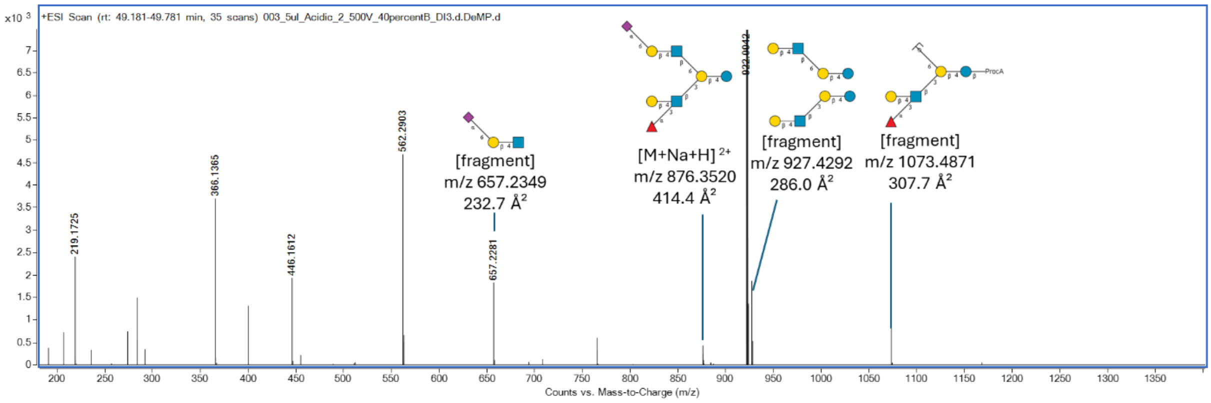

**Supplementary Figure 16. MS spectrum showing an identified HMO structure and its fragment ions eluting at 49.9 min.**

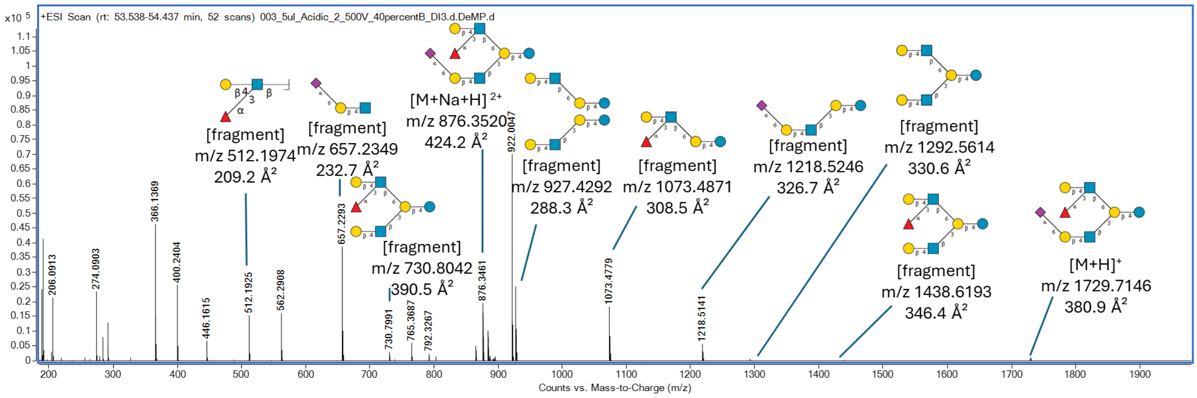

**Supplementary Figure 17. MS spectrum showing an identified HMO structure and its fragment ions eluting at 53.7 min.**

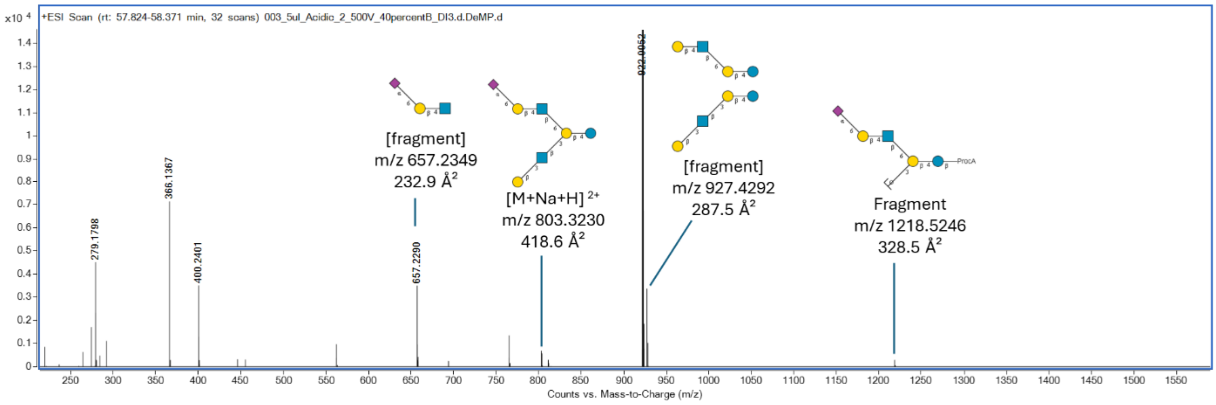

**Supplementary Figure 18. MS spectrum showing an identified HMO structure and its fragment ions eluting at 58.1 min.**

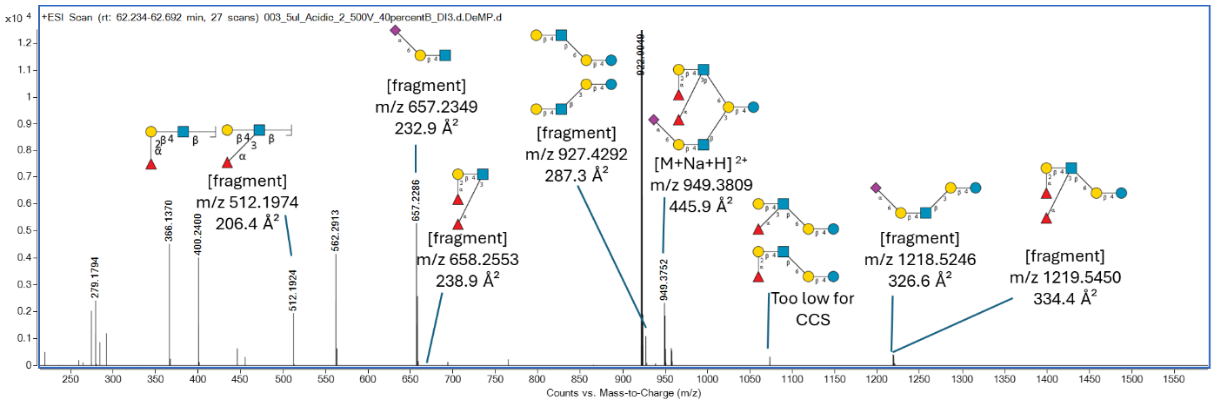

**Supplementary Figure 19. MS spectrum showing an identified HMO structure and its fragment ions eluting at 62.4 min.**

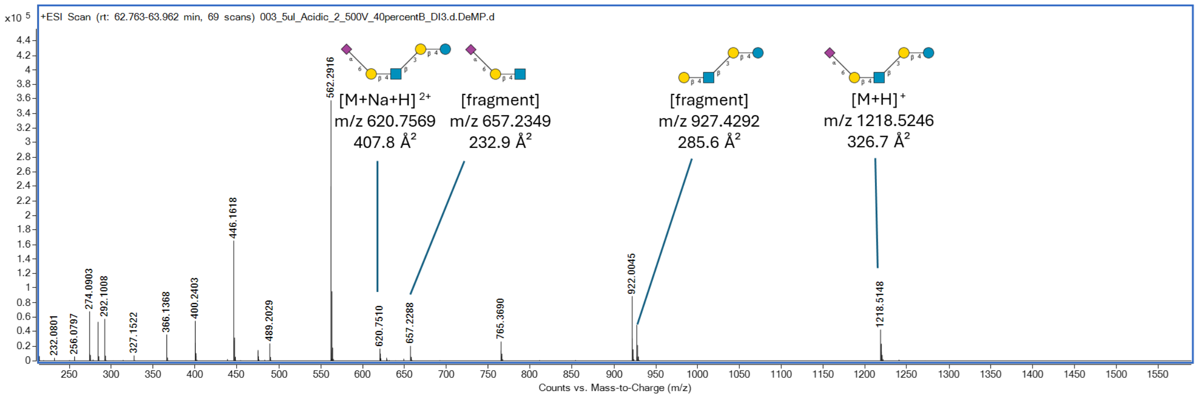

**Supplementary Figure 20. MS spectrum showing an identified HMO structure and its fragment ions eluting at 63.1 min.**

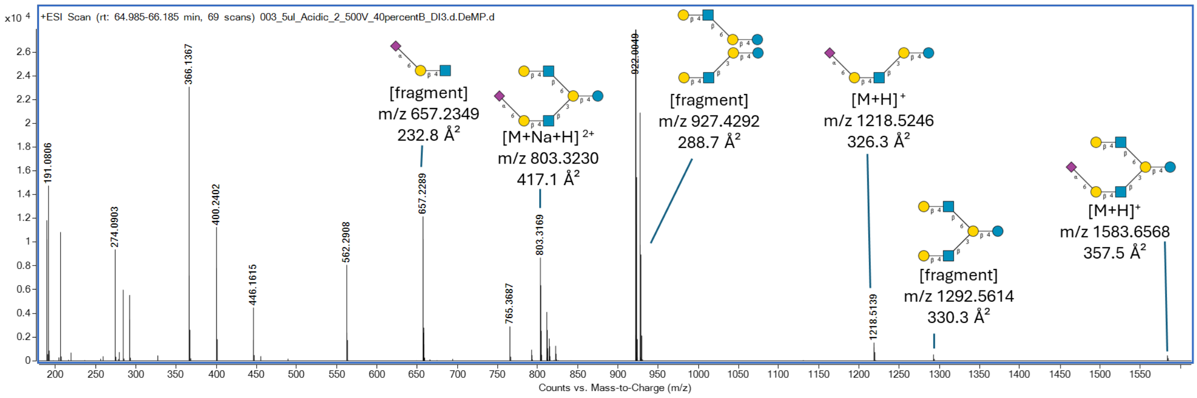

**Supplementary Figure 21. MS spectrum showing an identified HMO structure and its fragment ions eluting at 65.4 min.**

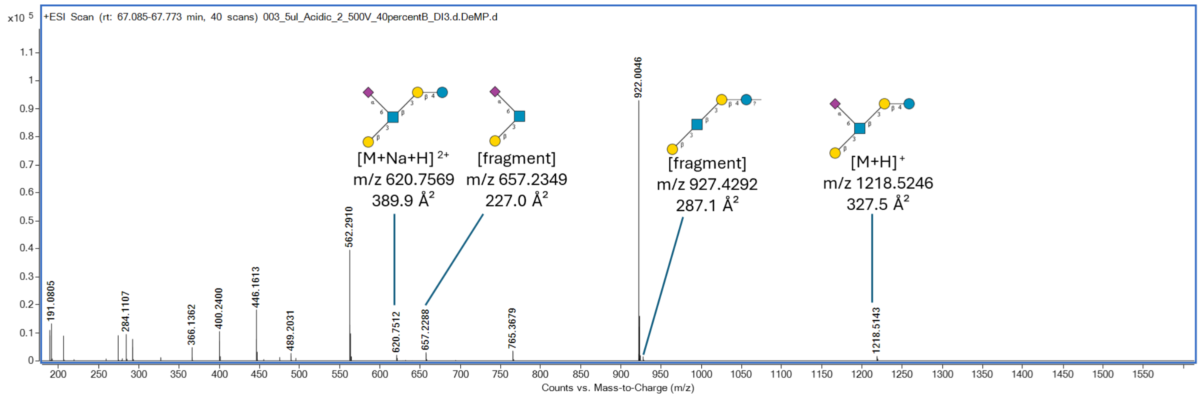

**Supplementary Figure 22. MS spectrum showing an identified HMO structure and its fragment ions eluting at 67.3 min.**

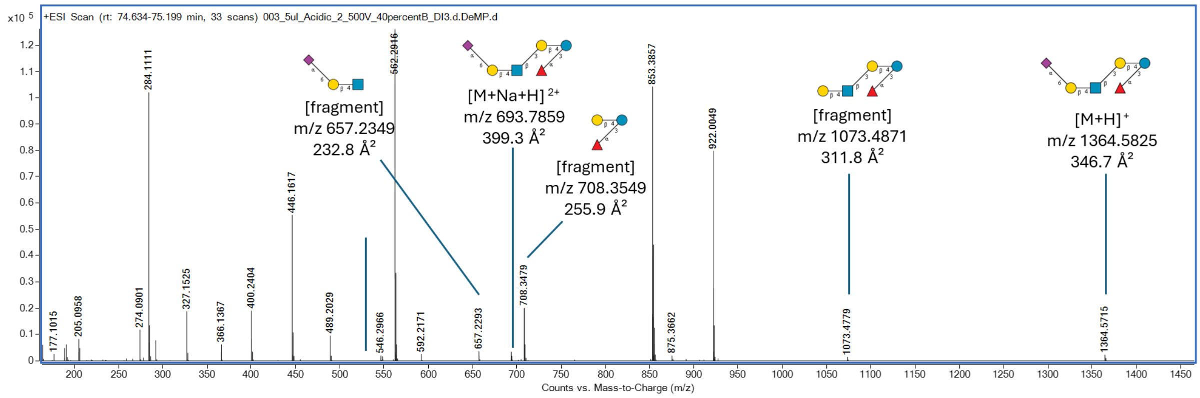

**Supplementary Figure 23. MS spectrum showing an identified HMO structure and its fragment ions eluting at 74.9 min.**

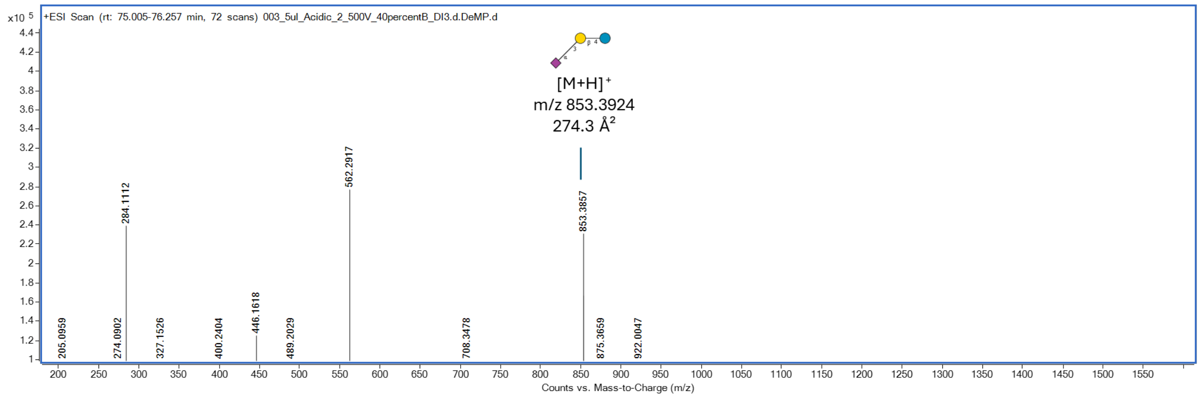

**Supplementary Figure 24. MS spectrum showing an identified HMO structure and its fragment ions eluting at 75.3 min.**

#### 2.3 Acidic Fraction from Donor 3.

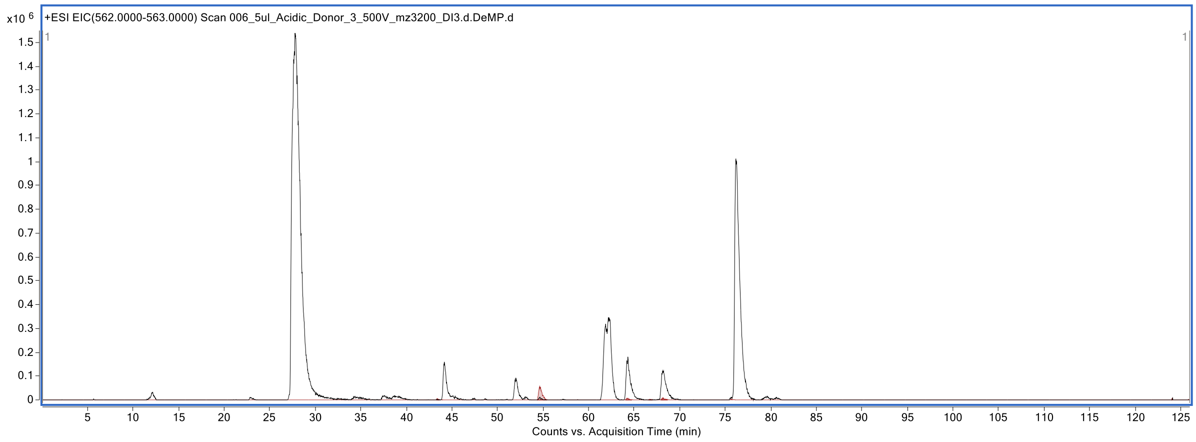

**Supplementary Figure 25. Extracted ion chromatogram at *m/z* 657-658 & and *m/z* 562-563 of HMOs labeled with procainamide.**

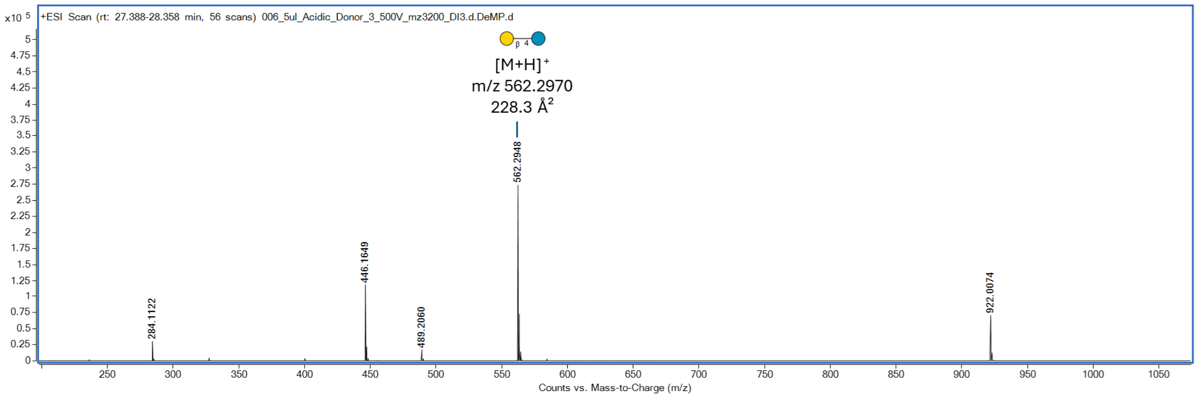

**Supplementary Figure 26. MS spectrum showing an identified HMO structure and its fragment ions eluting at 22.8 min.**

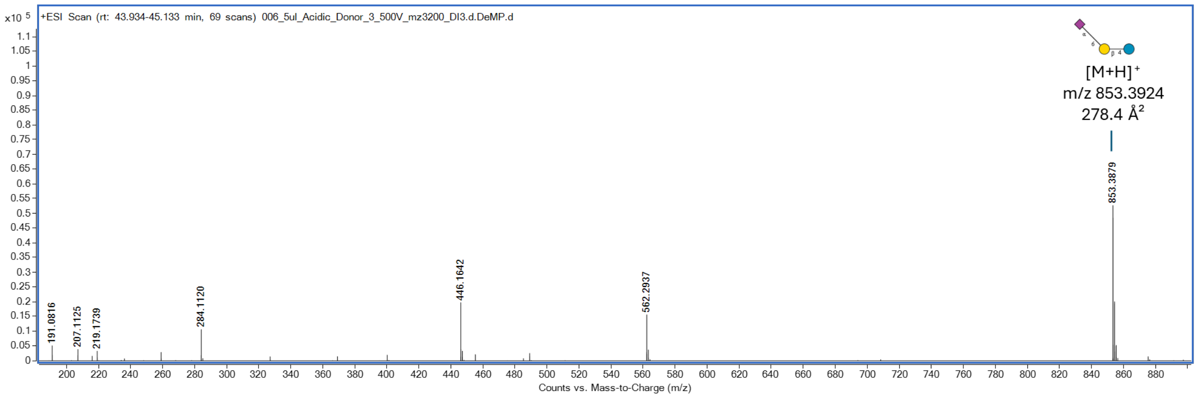

**Supplementary Figure 27. MS spectrum showing an identified HMO structure and its fragment ions eluting at 44.2 min.**

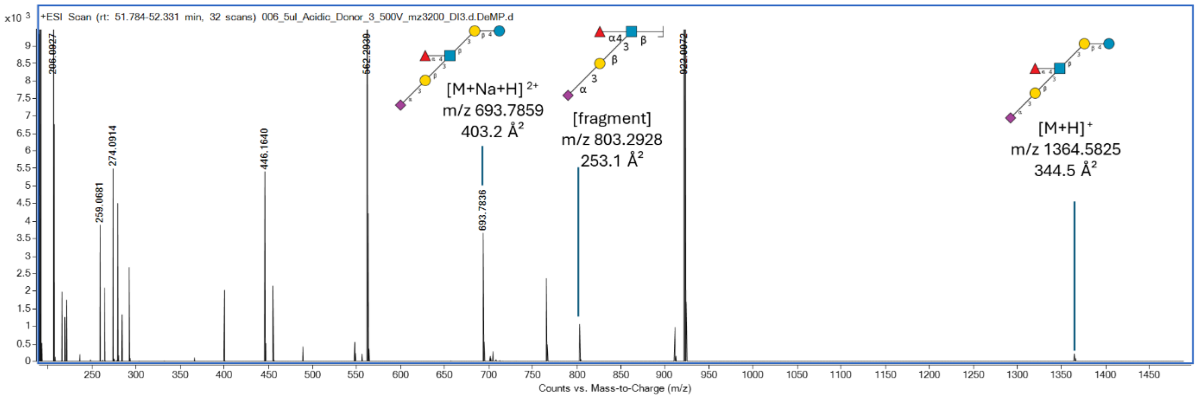

**Supplementary Figure 28. MS spectrum showing an identified HMO structure and its fragment ions eluting at 52.0 min.**

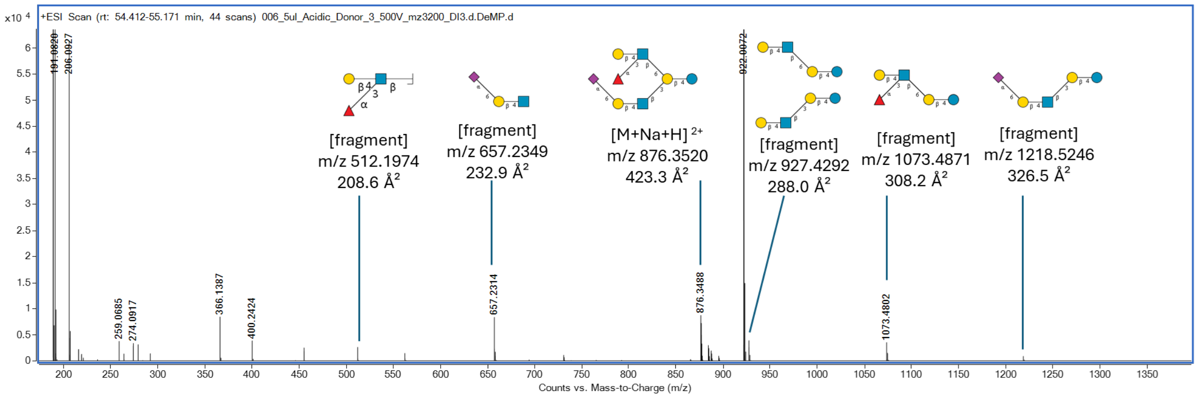

**Supplementary Figure 29. MS spectrum showing an identified HMO structure and its fragment ions eluting at 54.6 min.**

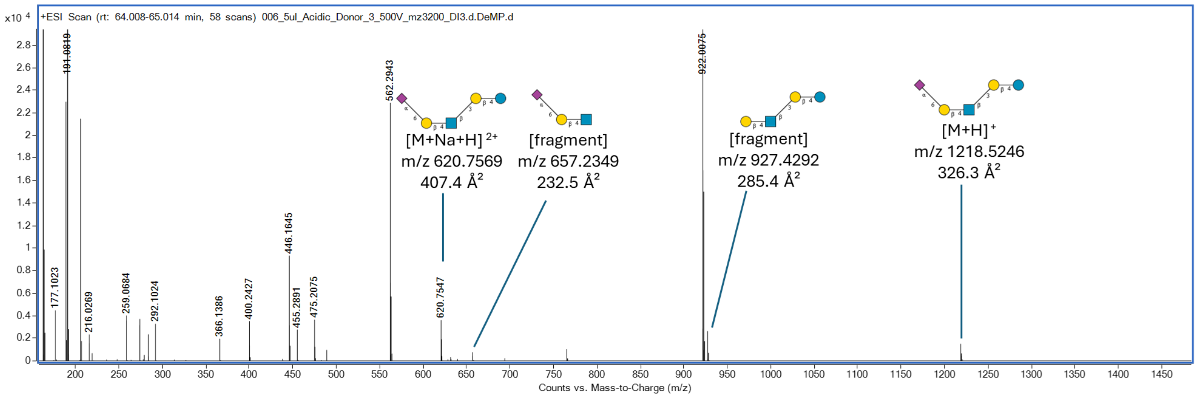

**Supplementary Figure 30. MS spectrum showing an identified HMO structure and its fragment ions eluting at 64.3 min.**

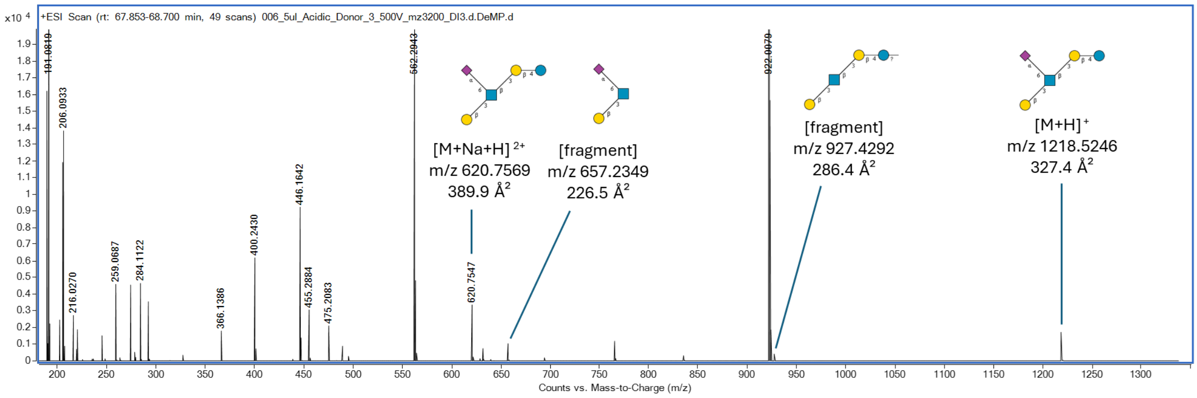

**Supplementary Figure 31. MS spectrum showing an identified HMO structure and its fragment ions eluting at 68.2 min.**

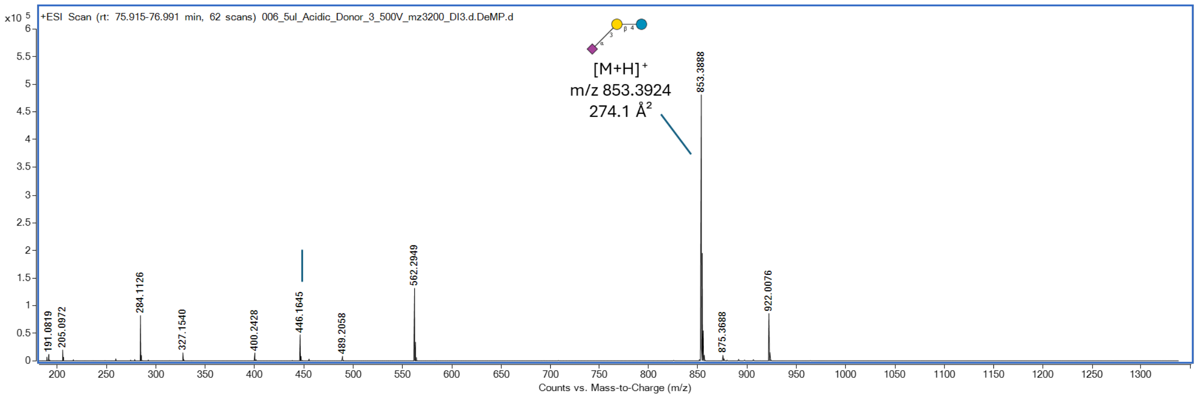

**Supplementary Figure 32. MS spectrum showing an identified HMO structure and its fragment ions eluting at 76.2 min.**

#### 2.4 Acidic Fraction from Donor 4.

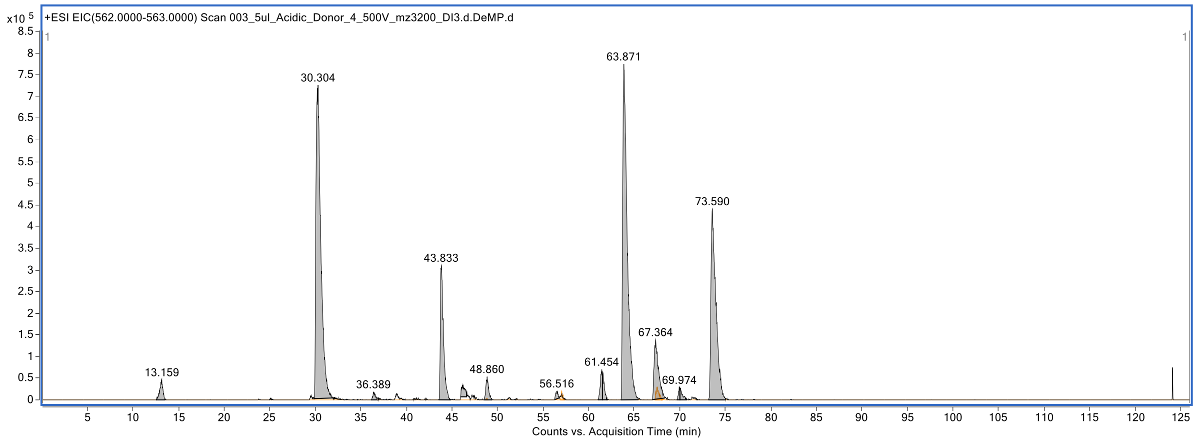

**Supplementary Figure 33. Extracted ion chromatogram at *m/z* 657-658 & and *m/z* 562-563 of HMOs labeled with procainamide.**

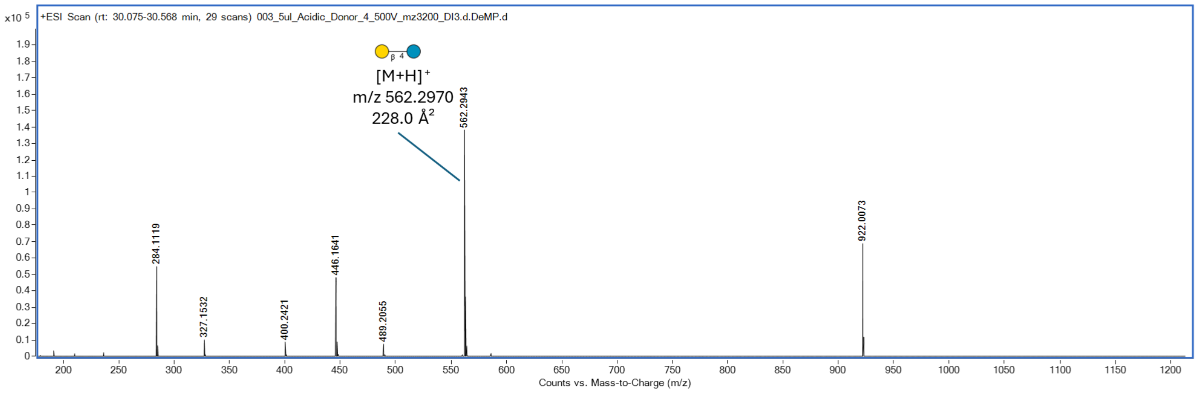

**Supplementary Figure 34. MS spectrum showing an identified HMO structure and its fragment ions eluting at 30.3 min.**

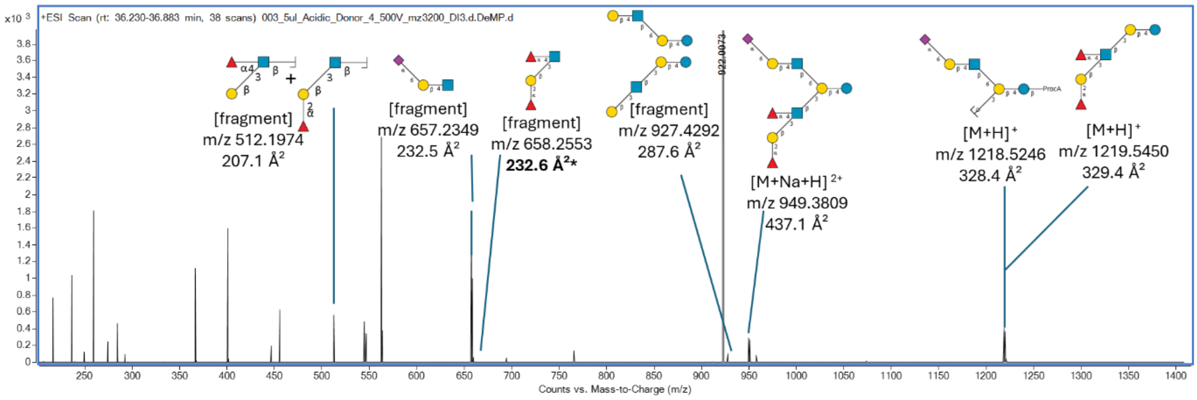

**Supplementary Figure 35. MS spectrum showing an identified HMO structure and its fragment ions eluting at 36.4 min.**

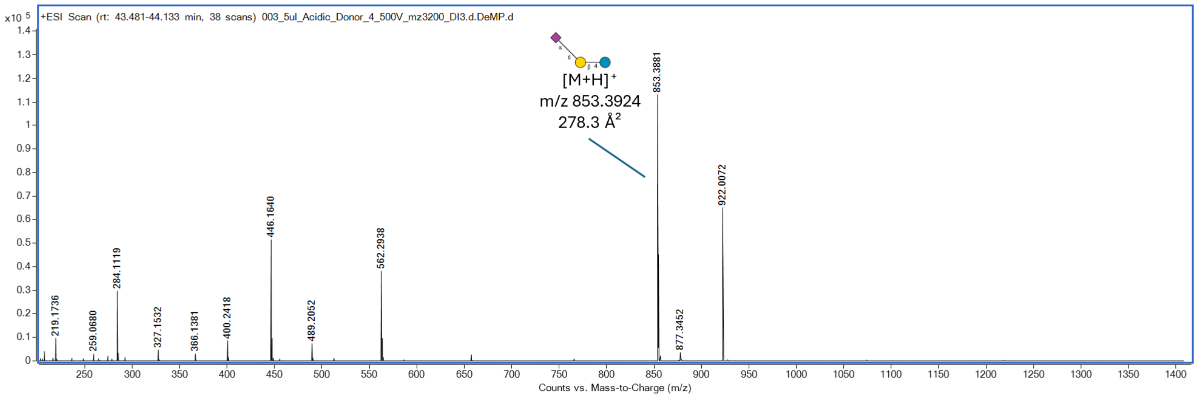

**Supplementary Figure 36. MS spectrum showing an identified HMO structure and its fragment ions eluting at 43.9 min.**

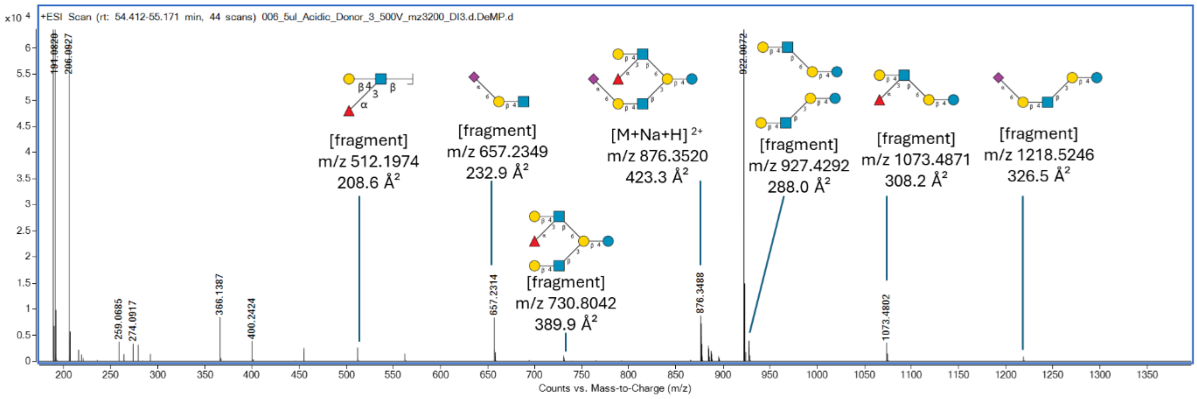

**Supplementary Figure 37. MS spectrum showing an identified HMO structure and its fragment ions eluting at 56.4 min.**

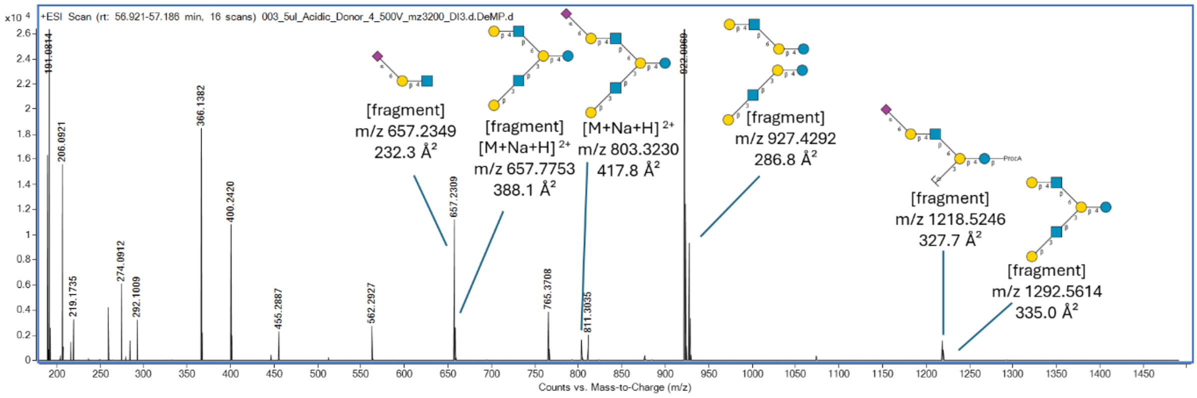

**Supplementary Figure 38. MS spectrum showing an identified HMO structure and its fragment ions eluting at 57.0 min.**

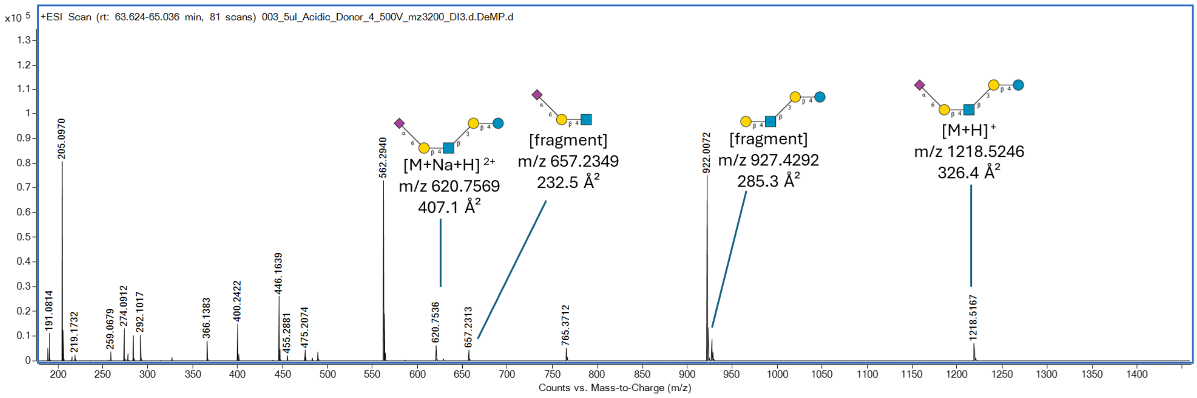

**Supplementary Figure 39. MS spectrum showing an identified HMO structure and its fragment ions eluting at 63.9 min.**

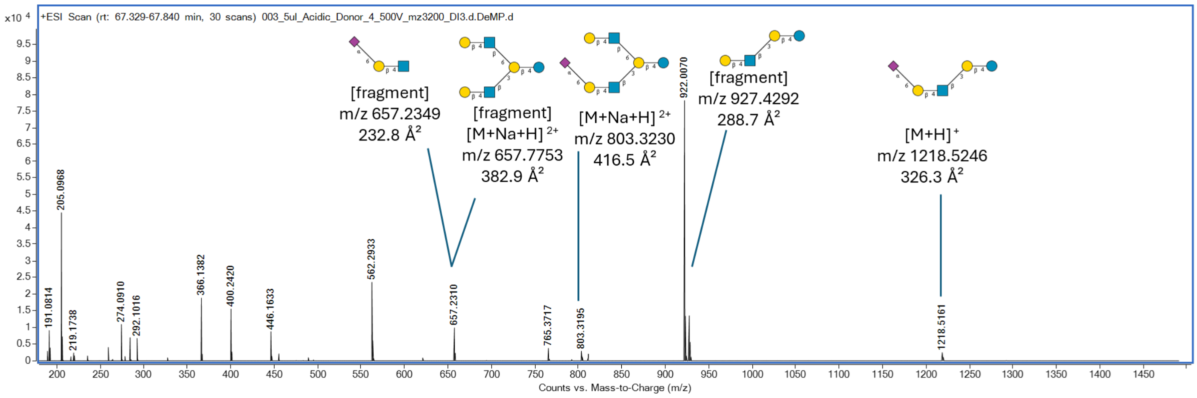

**Supplementary Figure 40. MS spectrum showing an identified HMO structure and its fragment ions eluting at 67.4 min.**

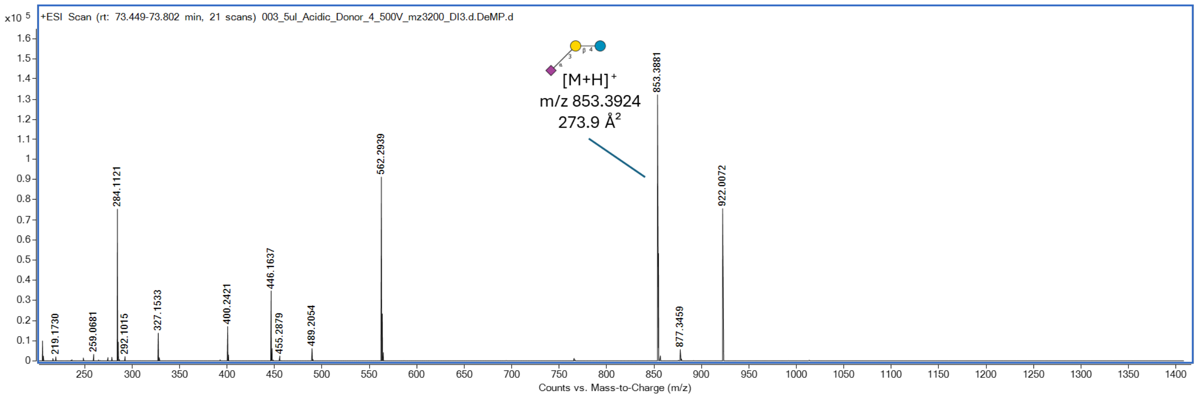

**Supplementary Figure 41. MS spectrum showing an identified HMO structure and its fragment ions eluting at 73.6 min.**

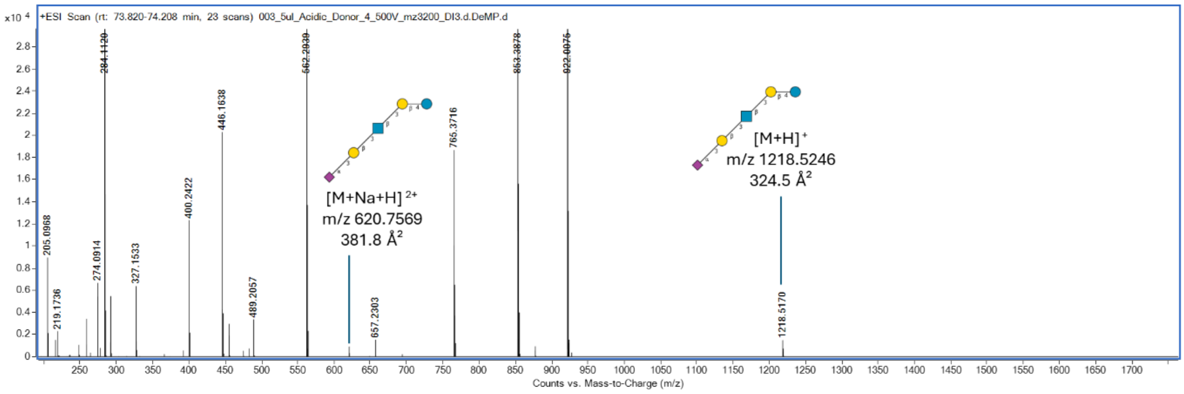

**Supplementary Figure 42. MS spectrum showing an identified HMO structure and its fragment ions eluting at 74.0 min.**

#### 2.5 Acidic Fraction from Donor 5.

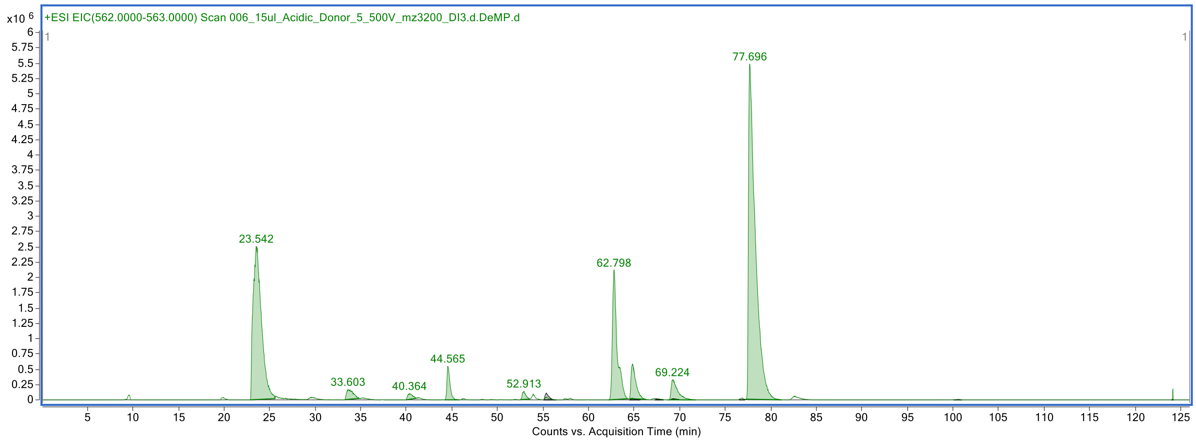

**Supplementary Figure 43. Extracted ion chromatogram at *m/z* 657-658 & and *m/z* 562-563 of HMOs labeled with procainamide.**

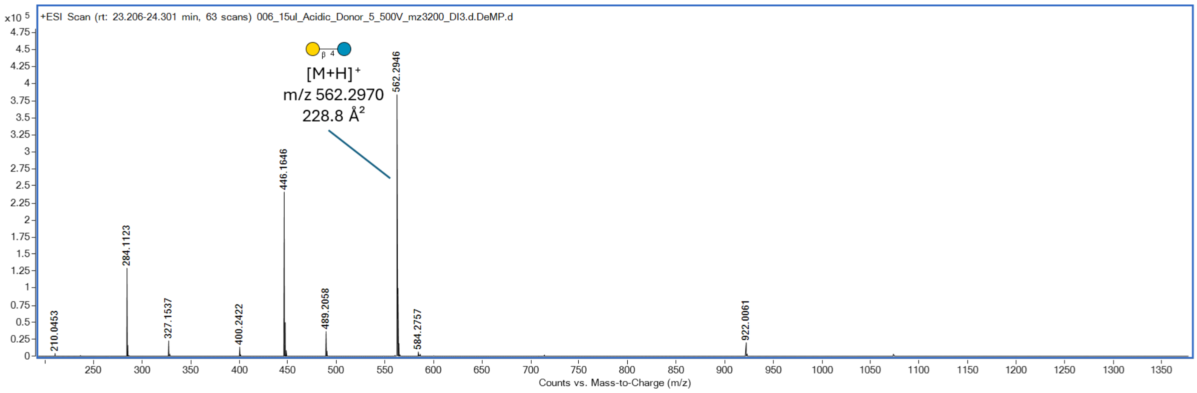

**Supplementary Figure 44. MS spectrum showing an identified HMO structure and its fragment ions eluting at 23.5 min.**

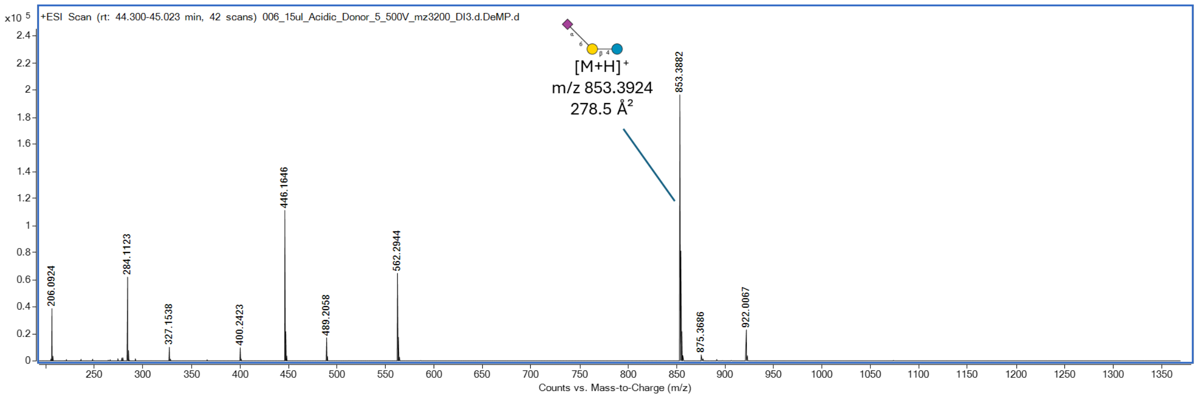

**Supplementary Figure 45. MS spectrum showing an identified HMO structure and its fragment ions eluting at 44.6 min.**

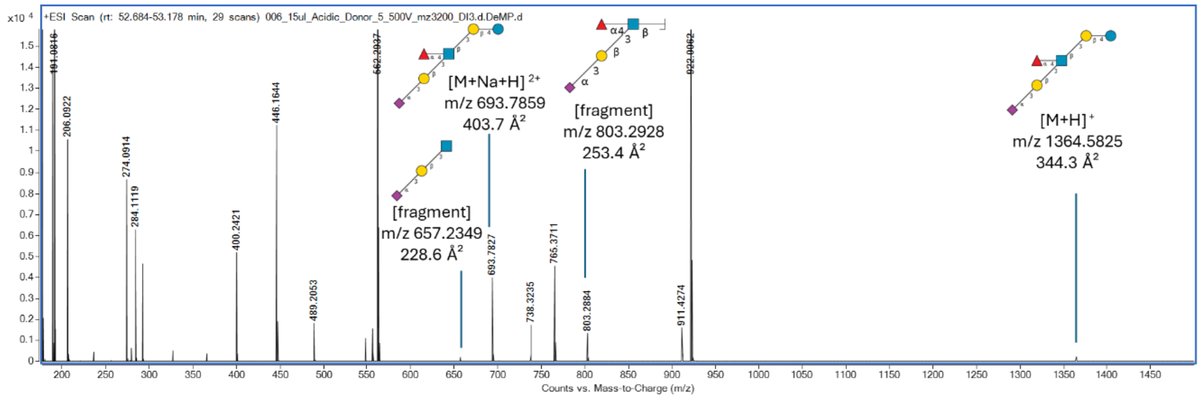

**Supplementary Figure 46. MS spectrum showing an identified HMO structure and its fragment ions eluting at 52.9 min.**

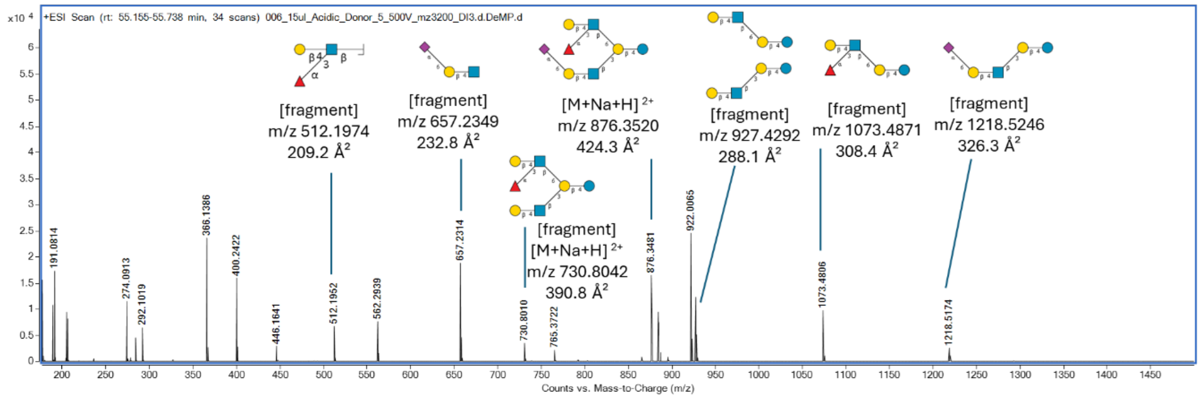

**Supplementary Figure 47. MS spectrum showing an identified HMO structure and its fragment ions eluting at 55.4 min.**

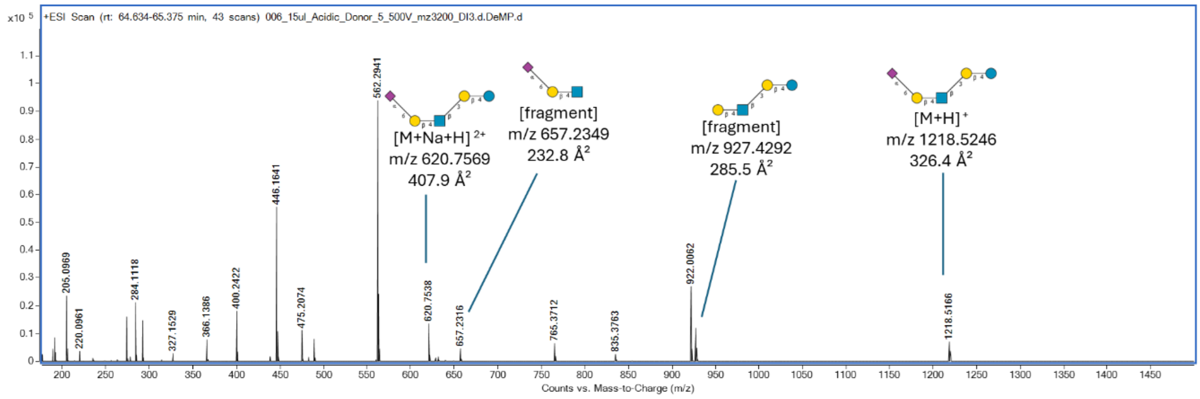

**Supplementary Figure 48. MS spectrum showing an identified HMO structure and its fragment ions eluting at 64.8 min.**

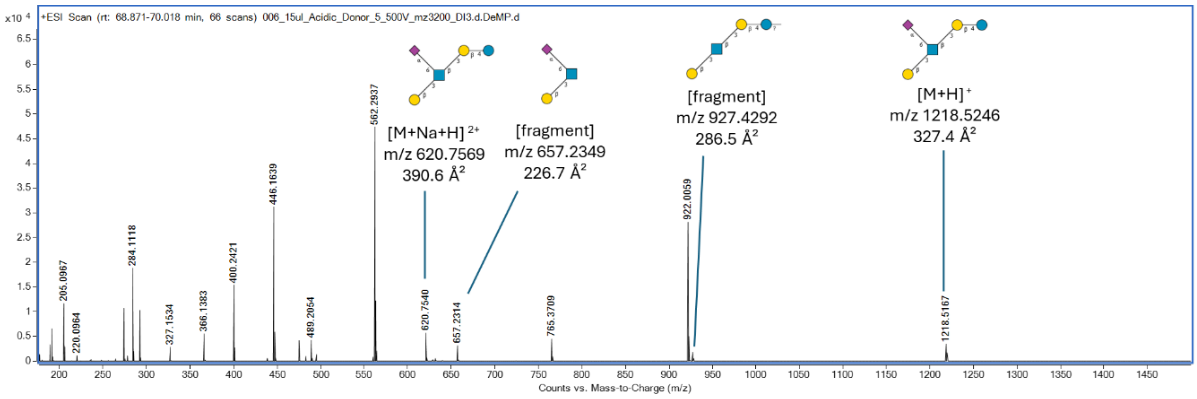

**Supplementary Figure 49. MS spectrum showing an identified HMO structure and its fragment ions eluting at 69.2 min.**

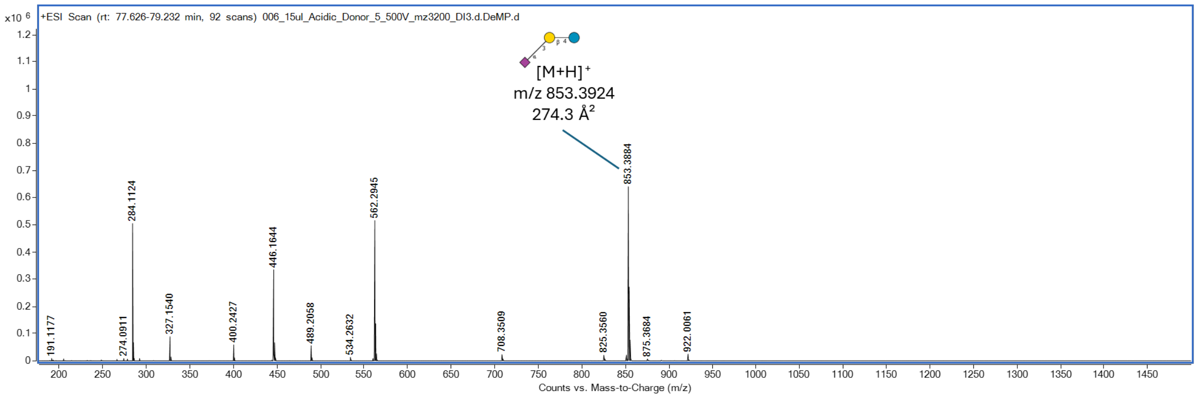

**Supplementary Figure 50. MS spectrum showing an identified HMO structure and its fragment ions eluting at 77.7 min.**

#### 2.6 Neutral Fraction from Donor 1.

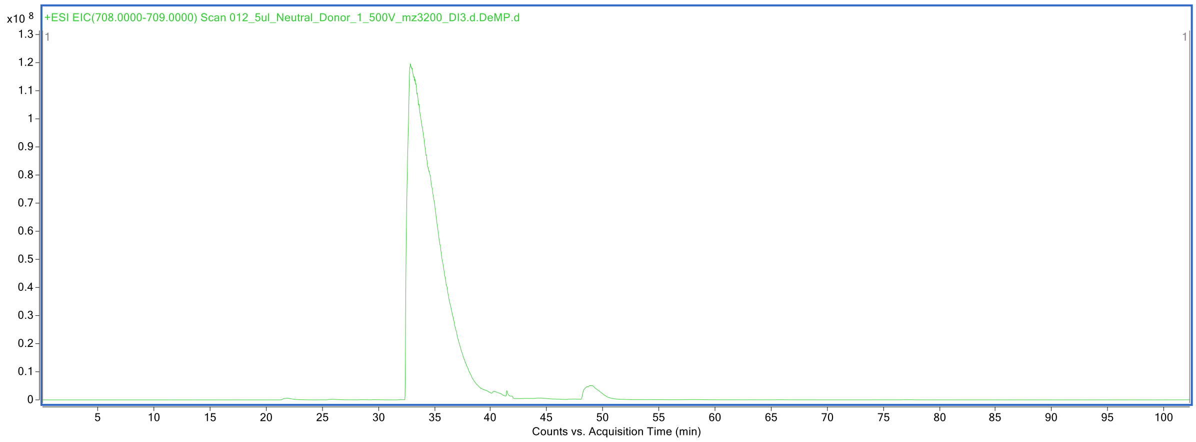

**Supplementary Figure 51. Extracted ion chromatogram at *m/z* 708-709 of HMOs labeled with procainamide.**

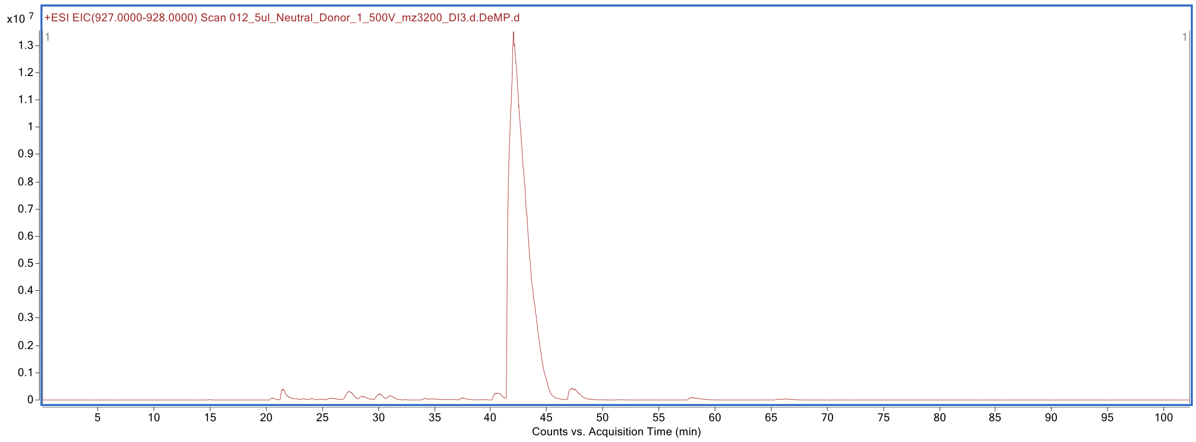

**Supplementary Figure 52. Extracted ion chromatogram at *m/z* 927-928 of HMOs labeled with procainamide.**

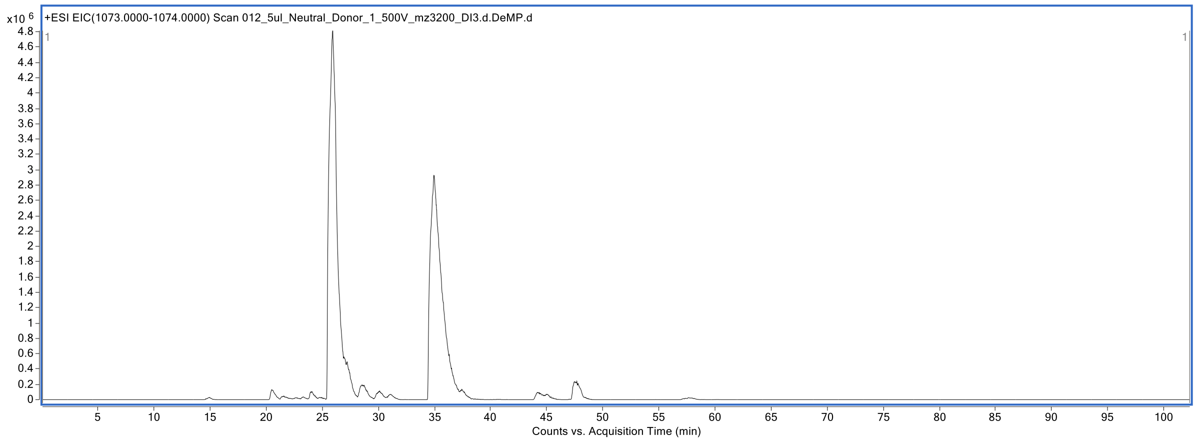

**Supplementary Figure 53. Extracted ion chromatogram at *m/z* 1073-1074 of HMOs labeled with procainamide.**

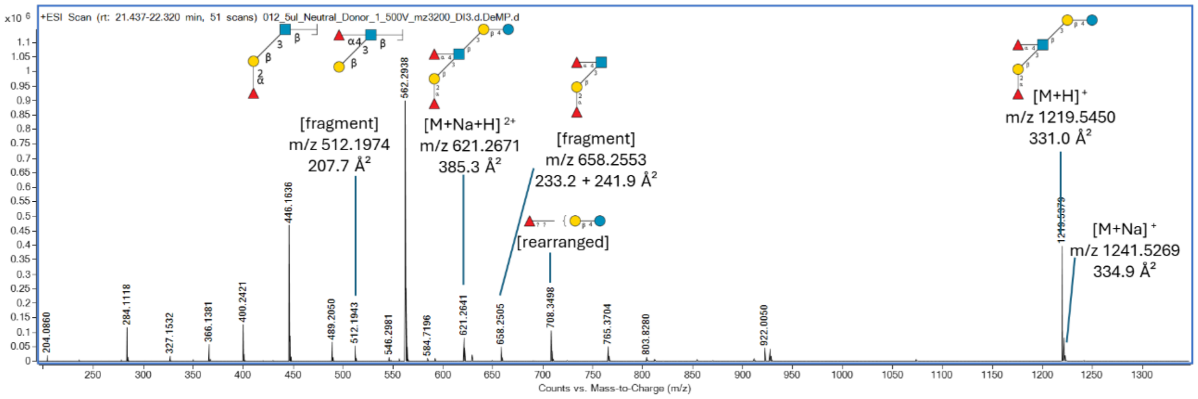

**Supplementary Figure 54. MS spectrum showing an identified HMO structure and its fragment ions eluting at 21.8 min.**

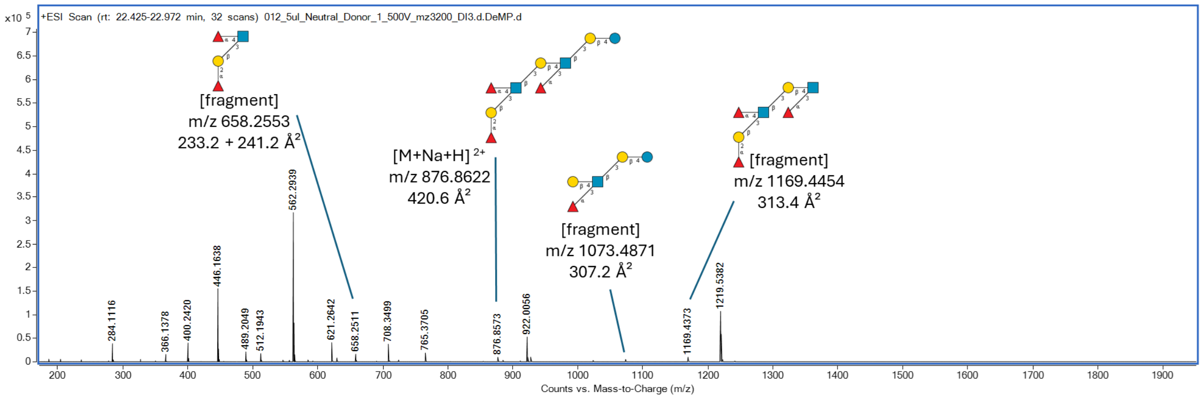

**Supplementary Figure 55. MS spectrum showing an identified HMO structure and its fragment ions eluting at 22.7 min.**

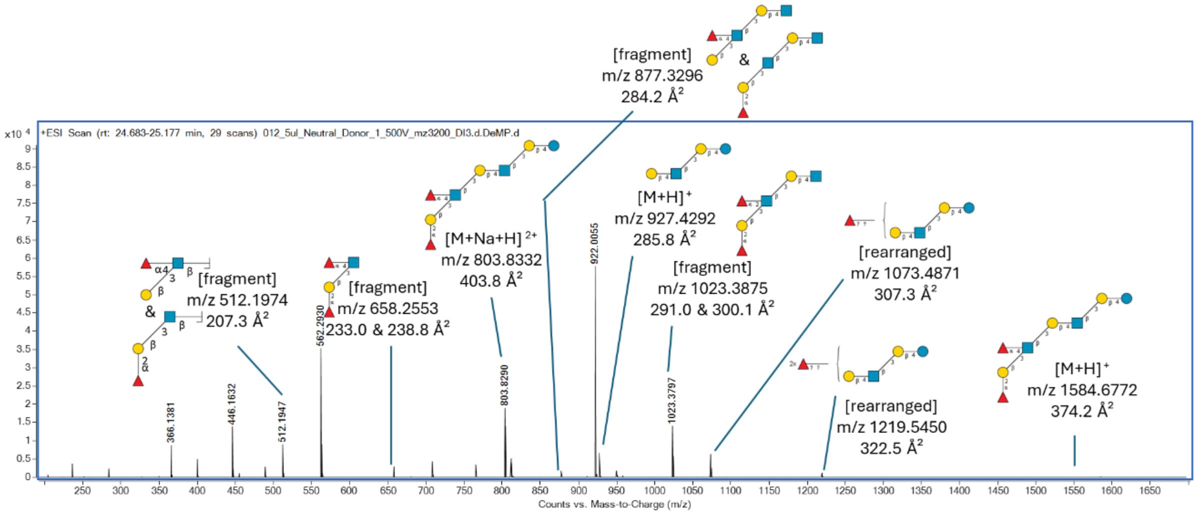

**Supplementary Figure 56. MS spectrum showing an identified HMO structure and its fragment ions eluting at 24.8 min.**

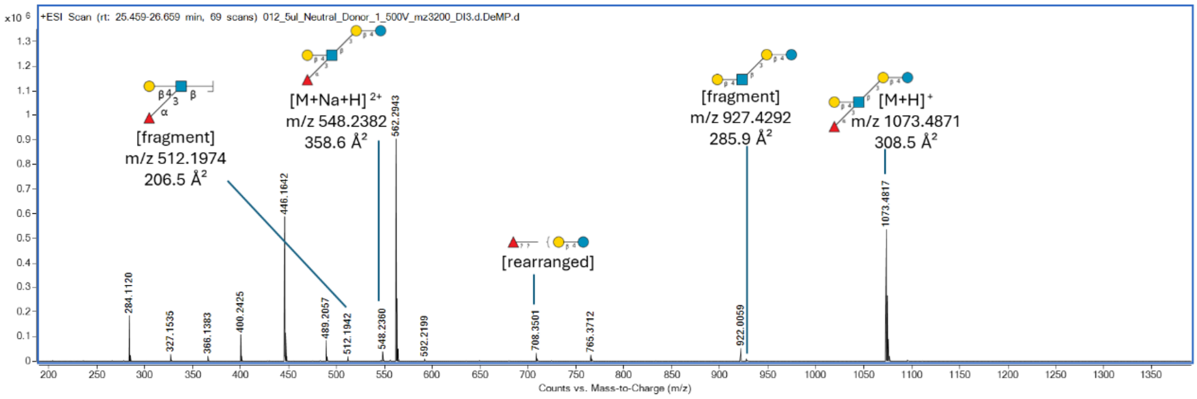

**Supplementary Figure 57. MS spectrum showing an identified HMO structure and its fragment ions eluting at 25.9 min.**

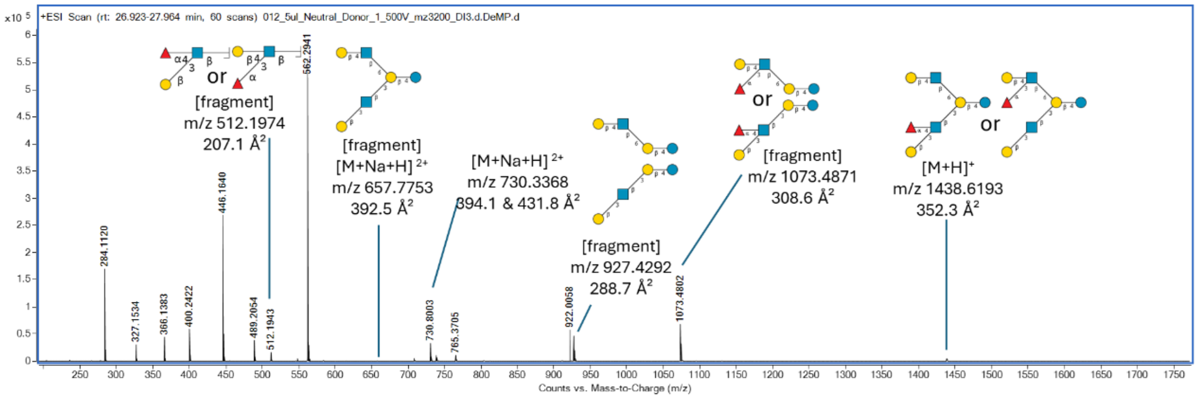

**Supplementary Figure 58. MS spectrum showing an identified HMO structure and its fragment ions eluting at 27.5 min.**

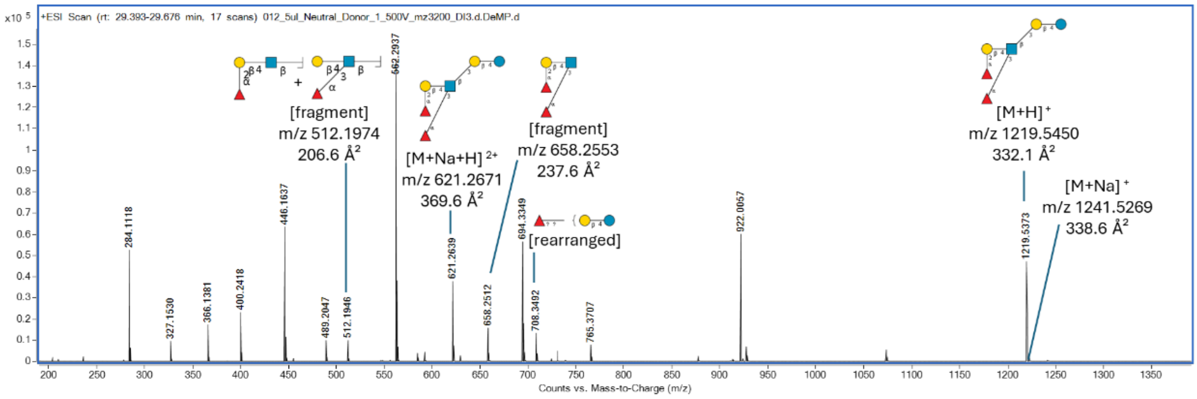

**Supplementary Figure 59. MS spectrum showing an identified HMO structure and its fragment ions eluting at 29.6 min.**

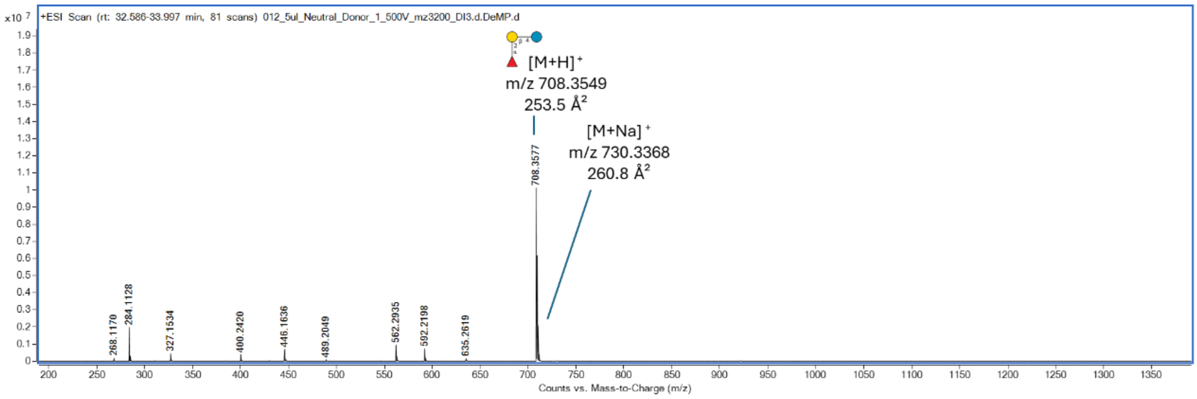

**Supplementary Figure 60. MS spectrum showing an identified HMO structure and its fragment ions eluting at 32.9 min.**

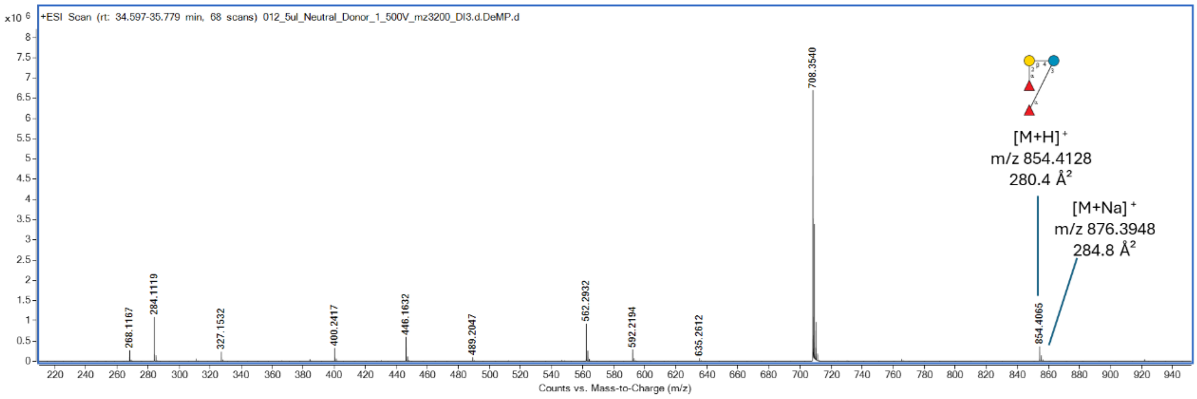

**Supplementary Figure 61. MS spectrum showing an identified HMO structure eluting at 34.9 min.**

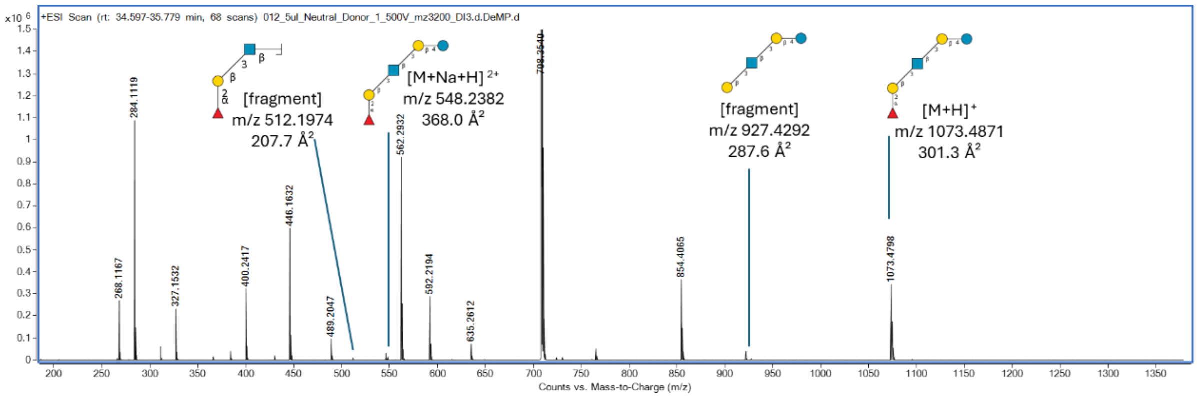

**Supplementary Figure 62. MS spectrum showing an identified HMO structure and its fragment ions eluting at 35.0 min.**

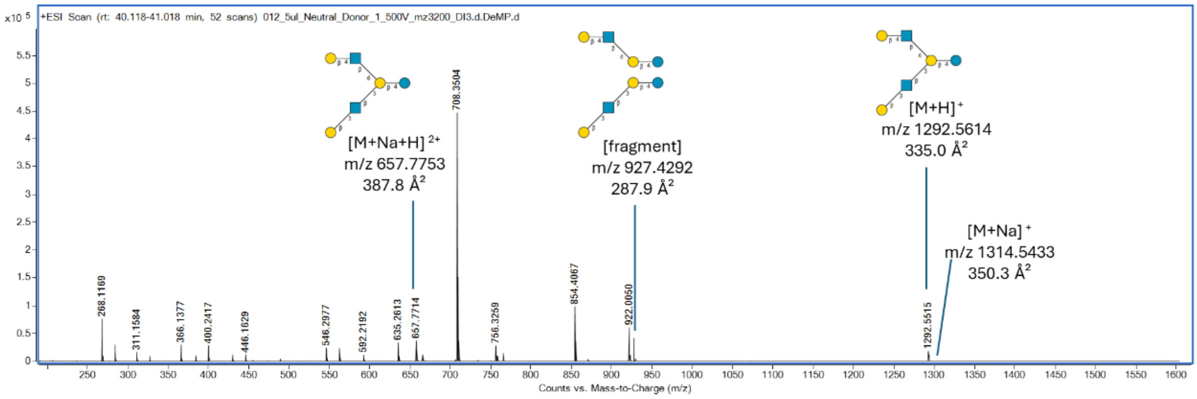

**Supplementary Figure 63. MS spectrum showing an identified HMO structure and its fragment ions eluting at 40.5 min.**

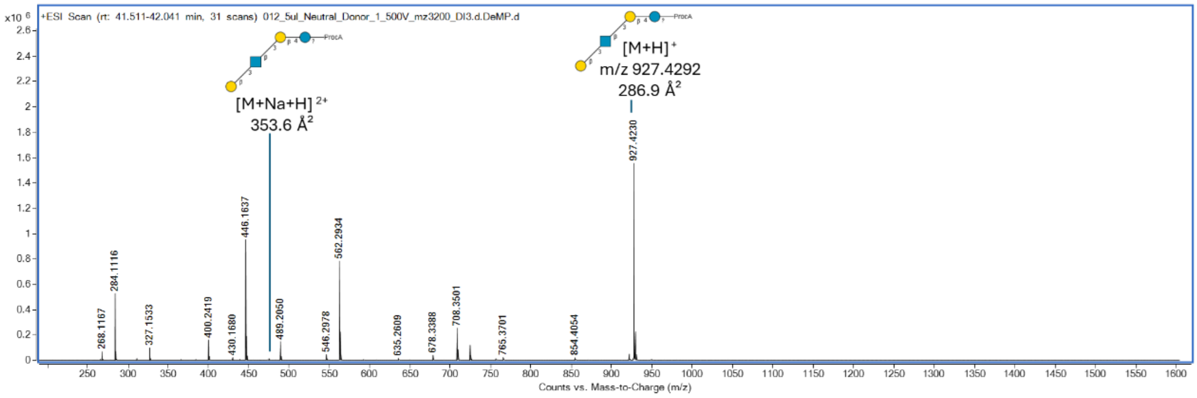

**Supplementary Figure 64. MS spectrum showing an identified HMO structure and its fragment ions eluting at 42.0 min (start of chromatographic peak).**

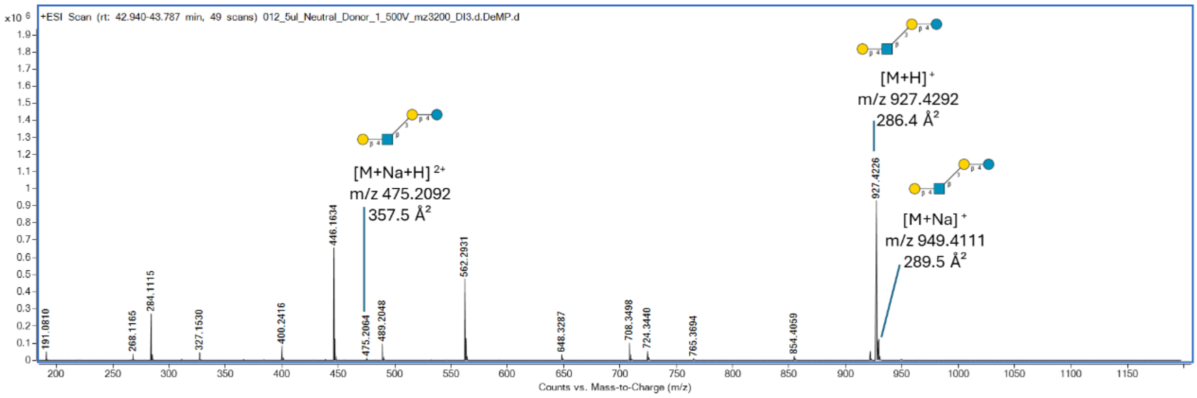

**Supplementary Figure 65. MS spectrum showing an identified HMO structure and its fragment ions eluting at 42 min (end of chromatographic peak).**

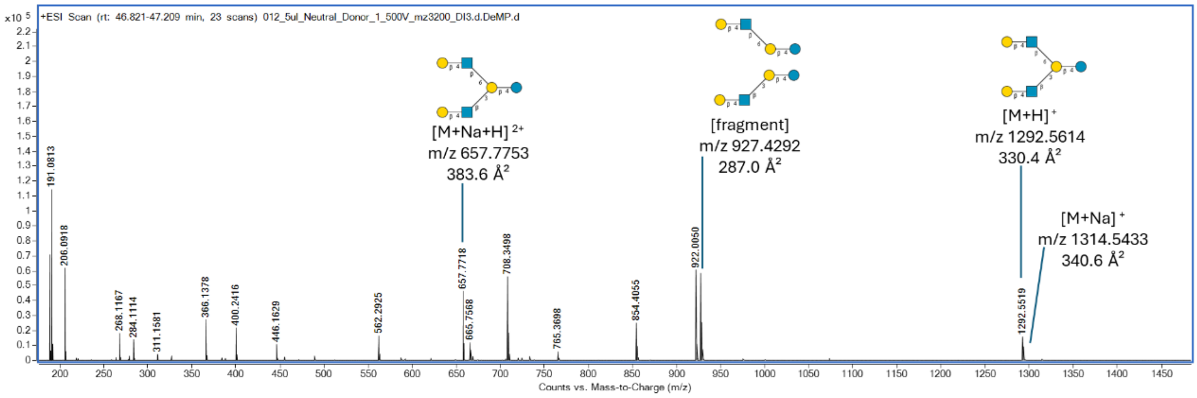

**Supplementary Figure 66. MS spectrum showing an identified HMO structure and its fragment ions eluting at 47.3 min.**

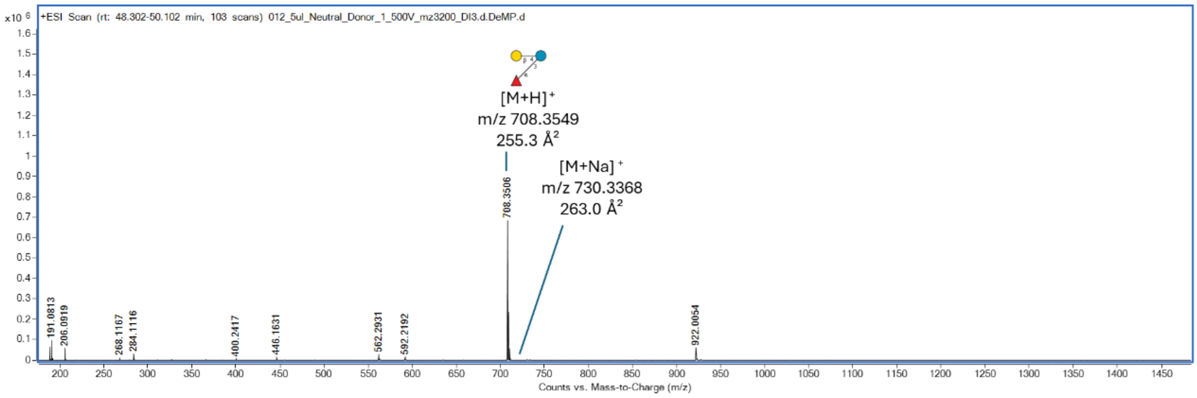

**Supplementary Figure 67. MS spectrum showing an identified HMO structure and its fragment ions eluting at 49.0 min.**

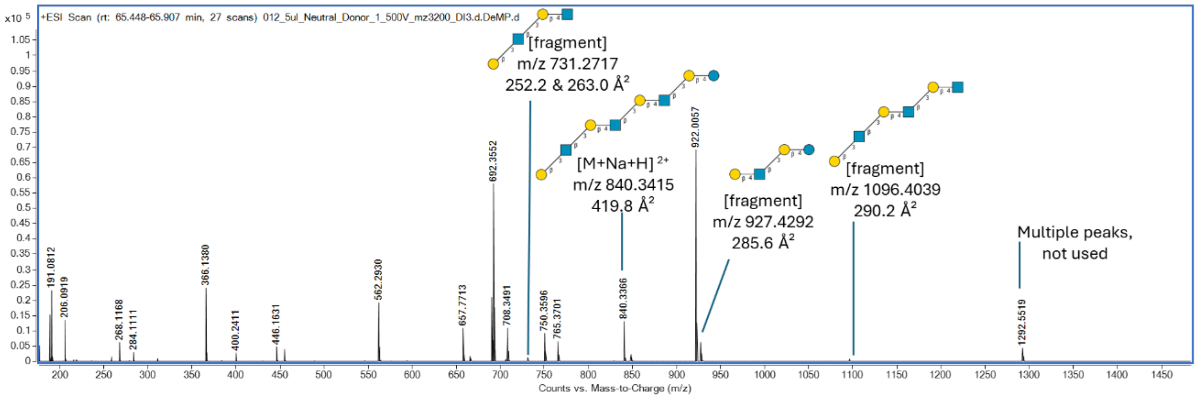

**Supplementary Figure 68. MS spectrum showing an identified HMO structure and its fragment ions eluting at 65.6 min.**

#### 2.7 Neutral Fraction from Donor 2.

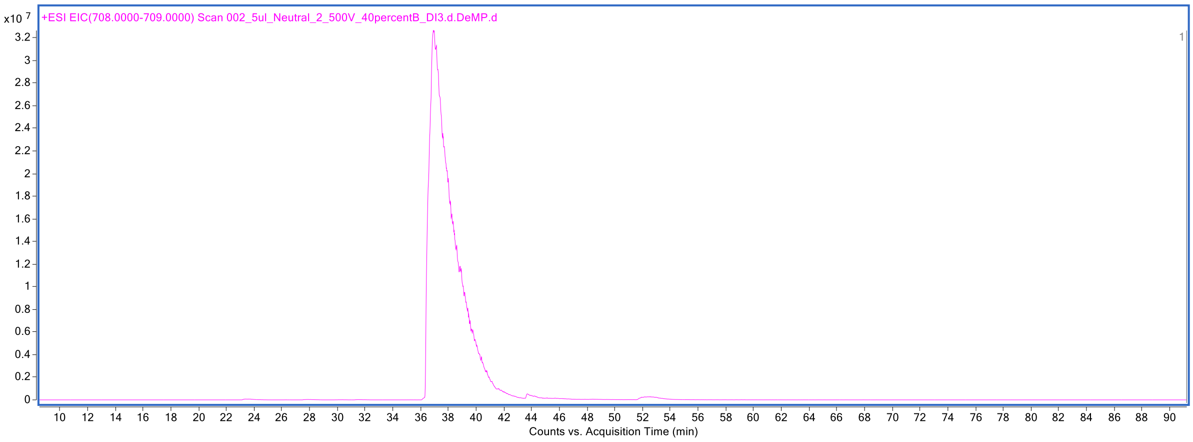

**Supplementary Figure 69. Extracted ion chromatogram at *m/z* 708-709 of HMOs labeled with procainamide.**

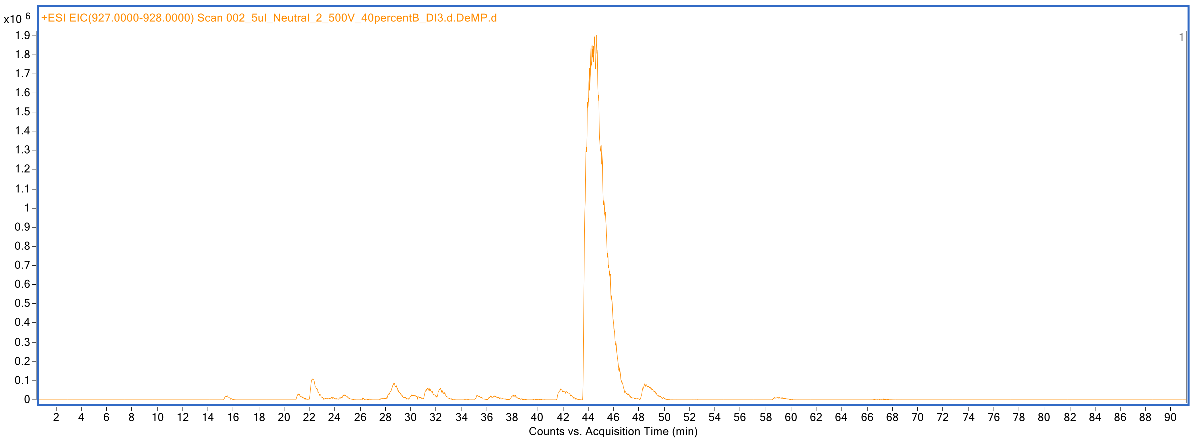

**Supplementary Figure 70. Extracted ion chromatogram at *m/z* 927-928 of HMOs labeled with procainamide.**

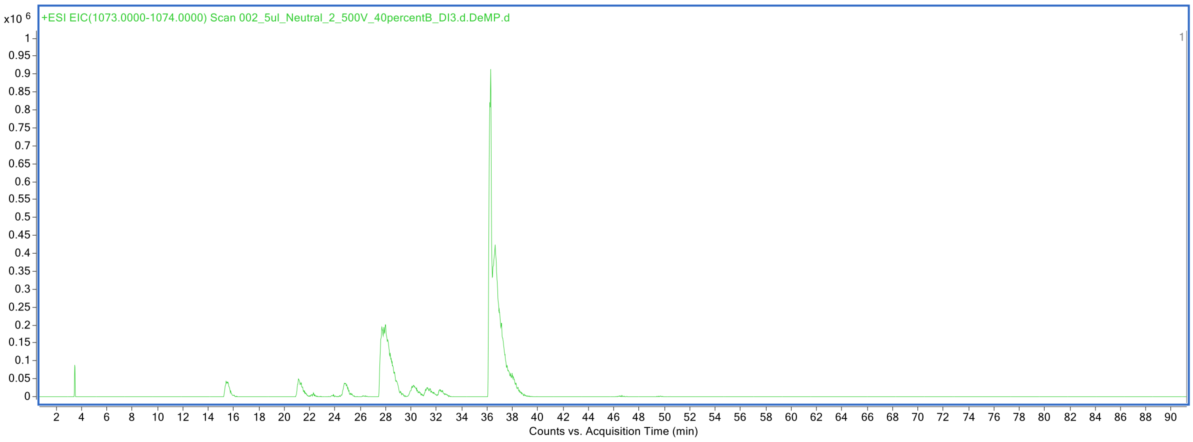

**Supplementary Figure 71. Extracted ion chromatogram at *m/z* 1073-1074 of HMOs labeled with procainamide.**

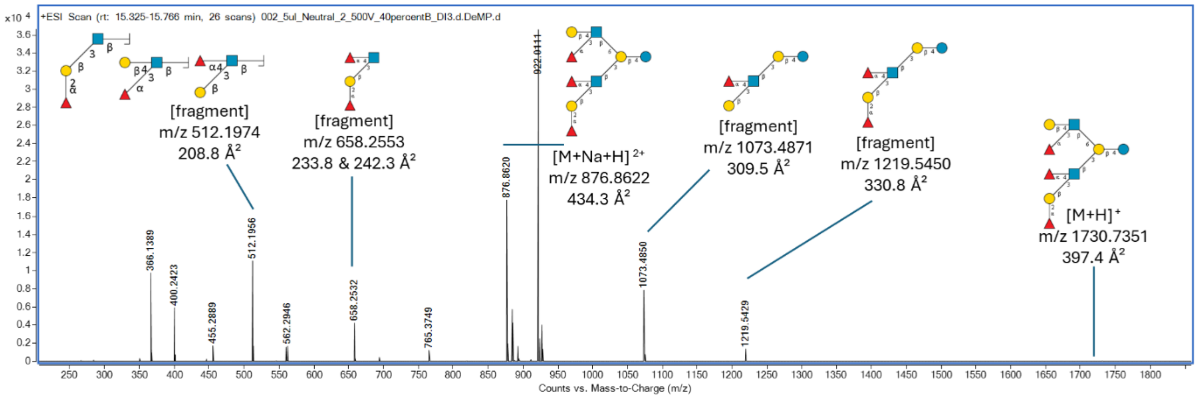

**Supplementary Figure 72. MS spectrum showing an identified HMO structure and its fragment ions eluting at 15.5 min.**

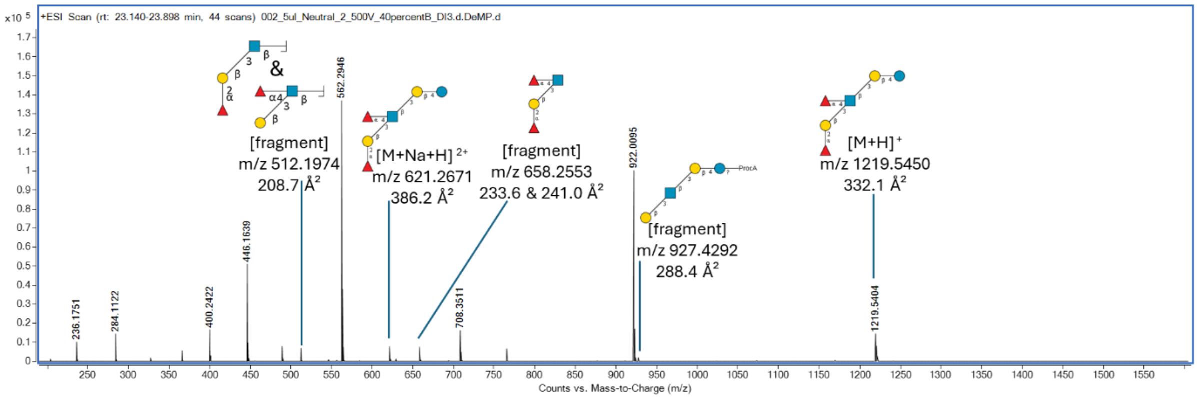

**Supplementary Figure 73. MS spectrum showing an identified HMO structure and its fragment ions eluting at 23.4 min.**

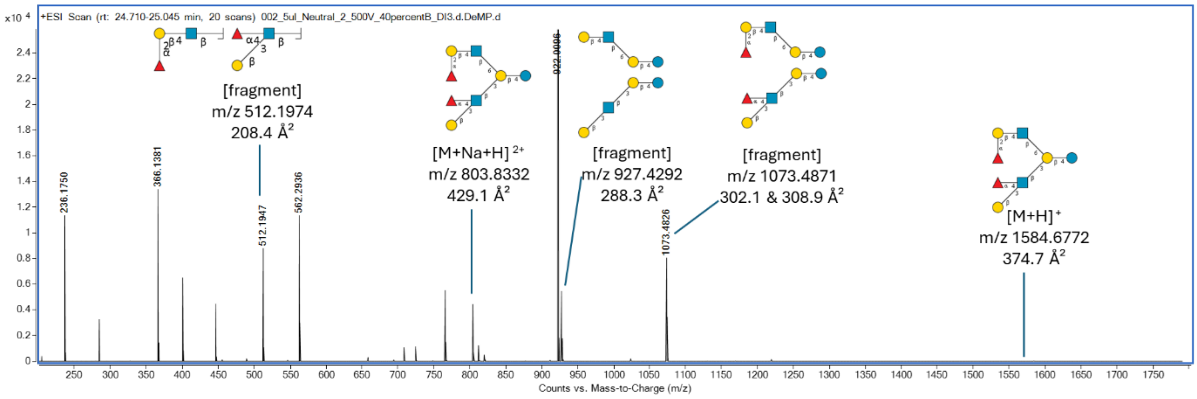

**Supplementary Figure 74. MS spectrum showing an identified HMO structure and its fragment ions eluting at 24.7 min.**

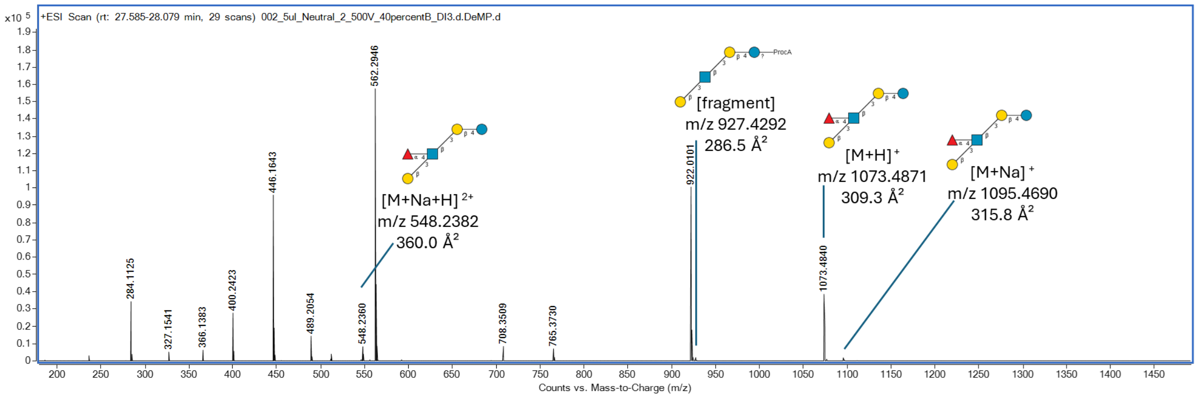

**Supplementary Figure 75. MS spectrum showing an identified HMO structure and its fragment ions eluting at 28.0 min.**

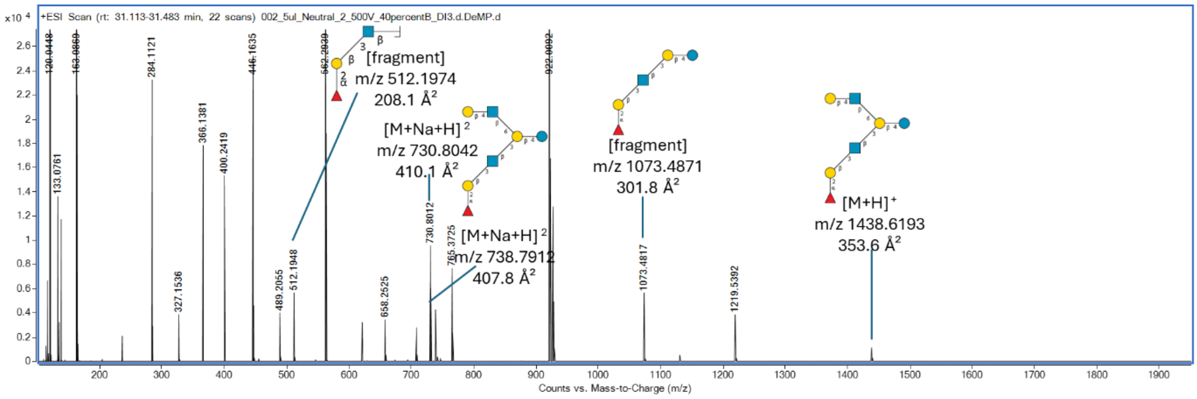

**Supplementary Figure 76. MS spectrum showing an identified HMO structure and its fragment ions eluting at 31.2 min.**

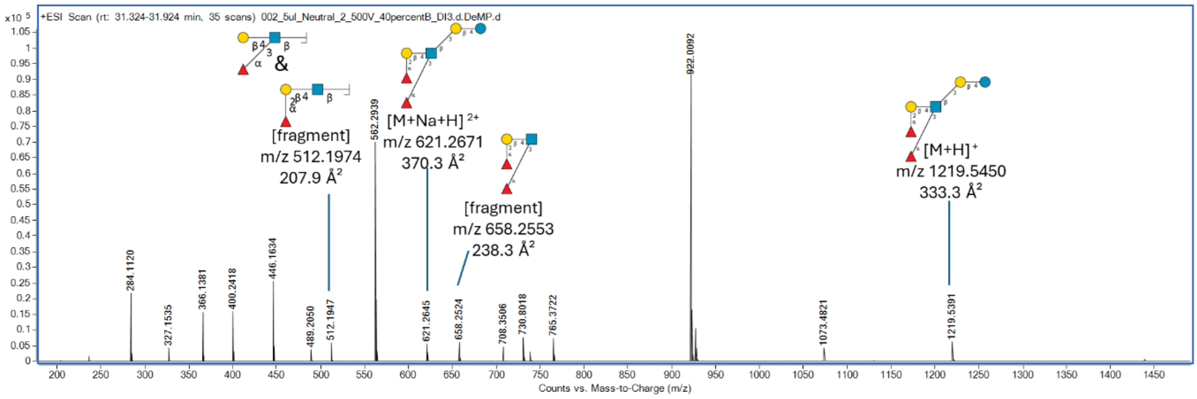

**Supplementary Figure 77. MS spectrum showing an identified HMO structure and its fragment ions eluting at 31.4 min.**

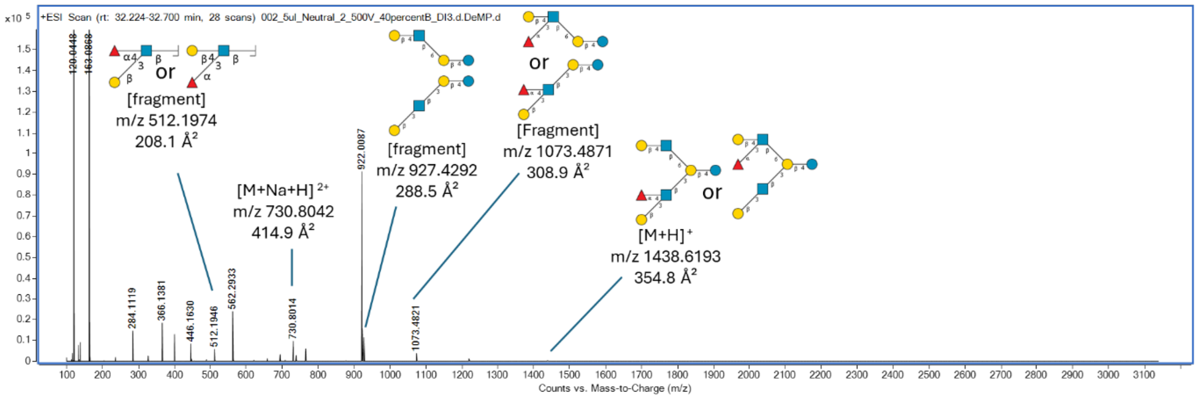

**Supplementary Figure 78. MS spectrum showing an identified HMO structure and its fragment ions eluting at 32.3 min.**

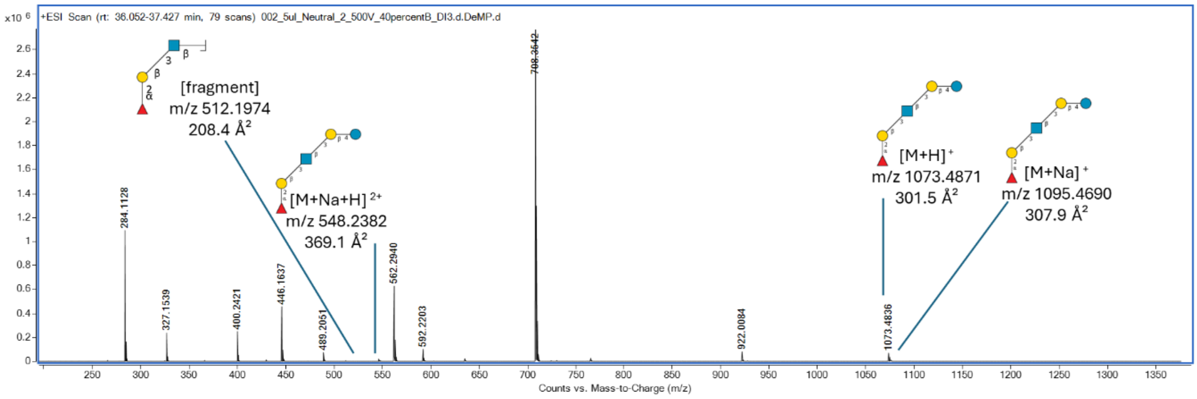

**Supplementary Figure 79. MS spectrum showing an identified HMO structure and its fragment ions eluting at 36.3 min.**

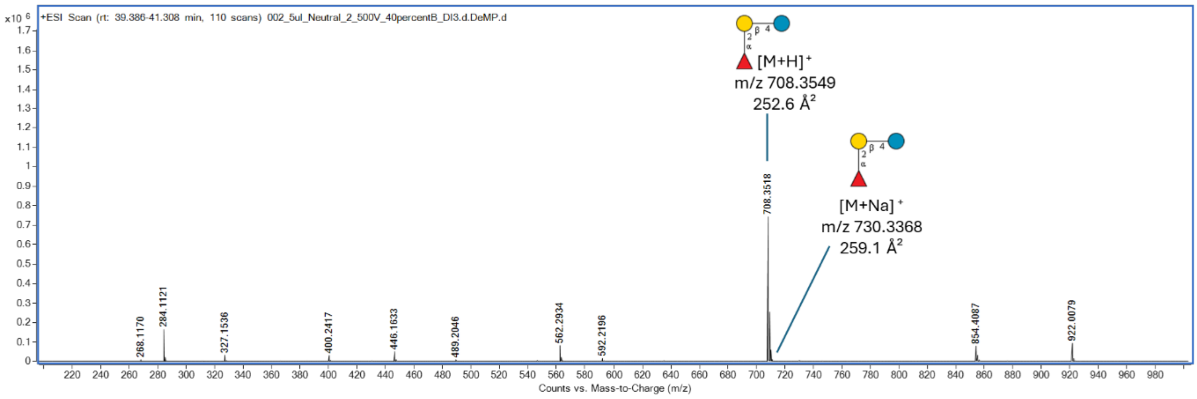

**Supplementary Figure 80. MS spectrum showing an identified HMO structure and its fragment ions eluting at 37.0 min.**

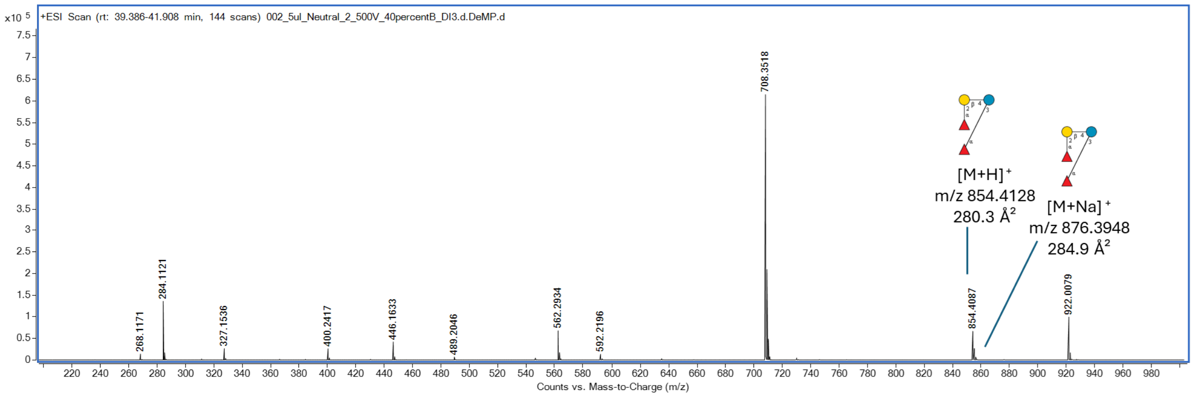

**Supplementary Figure 81. MS spectrum showing an identified HMO structure and its fragment ions eluting at 40.3 min.**

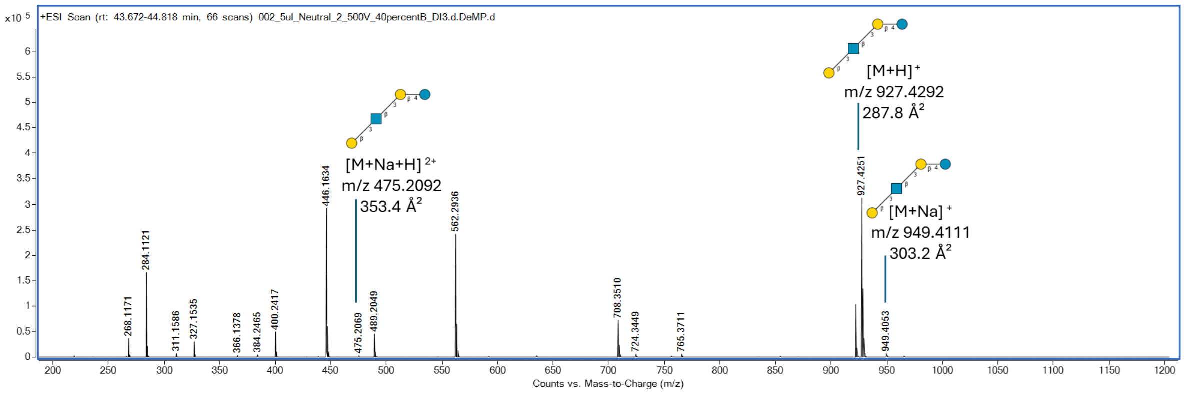

**Supplementary Figure 82. MS spectrum showing an identified HMO structure and its fragment ions eluting at 44.6 min.**

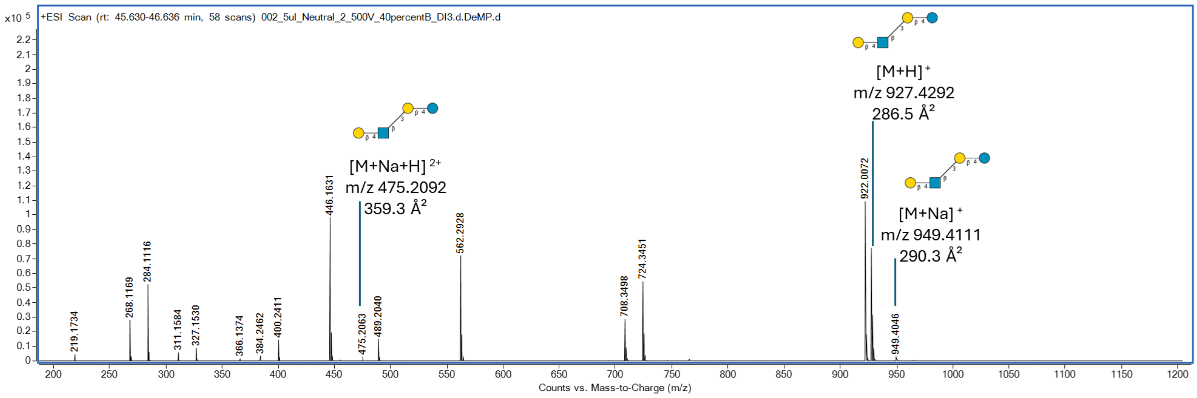

**Supplementary Figure 83. MS spectrum showing an identified HMO structure and its fragment ions eluting at 44.6 min.**

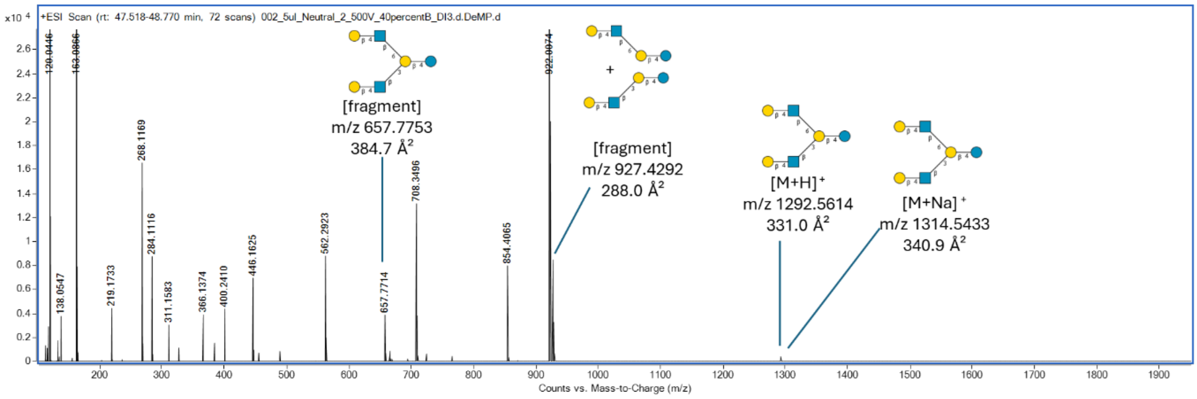

**Supplementary Figure 84. MS spectrum showing an identified HMO structure and its fragment ions eluting at 48.5 min.**

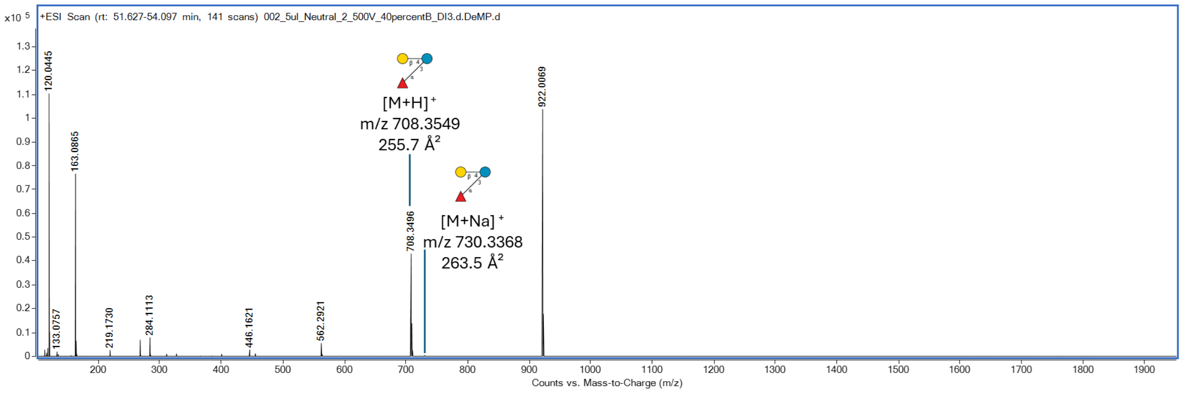

**Supplementary Figure 85. MS spectrum showing an identified HMO structure and its fragment ions eluting at 52.4 min.**

#### 2.8 Neutral Fraction from Donor 3.

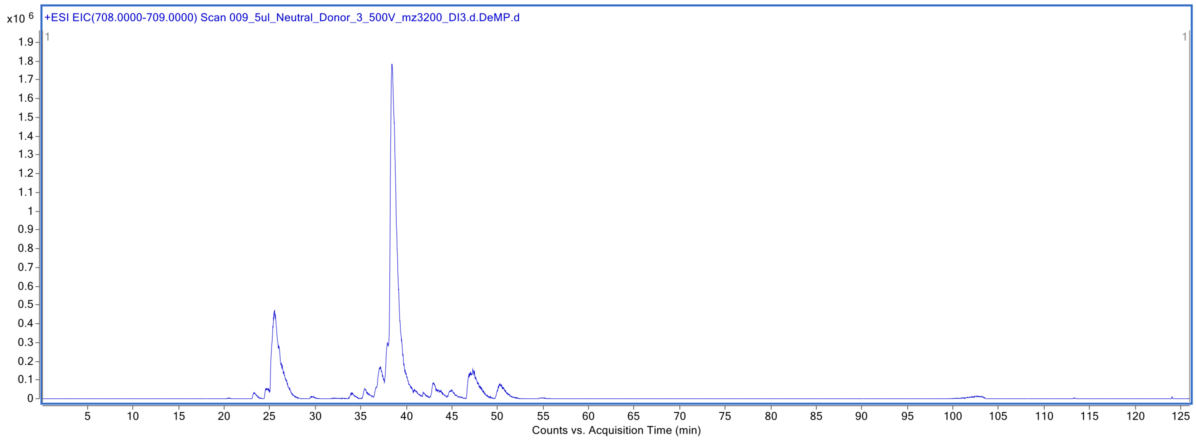

**Supplementary Figure 86. Extracted ion chromatogram at *m/z* 708-709 of HMOs labeled with procainamide.**

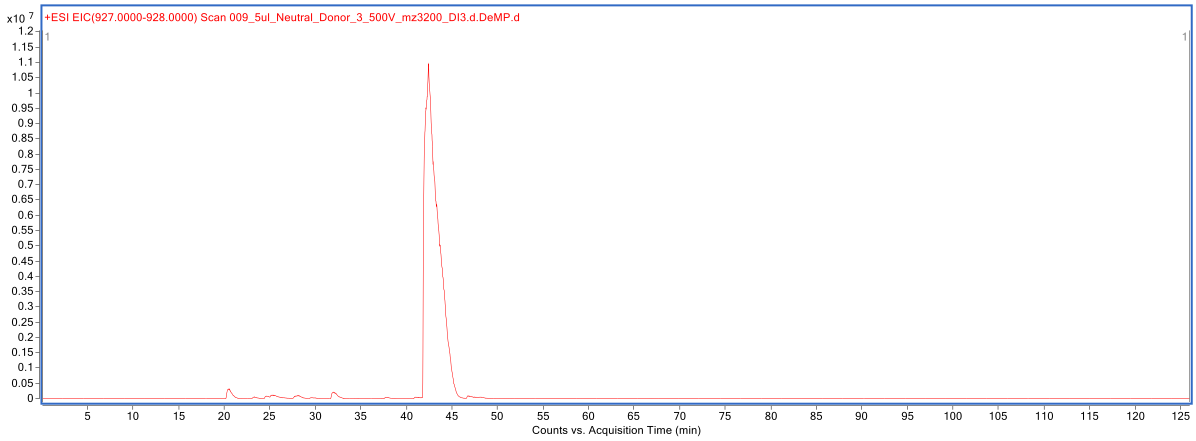

**Supplementary Figure 87. Extracted ion chromatogram at *m/z* 927-928 of HMOs labeled with procainamide.**

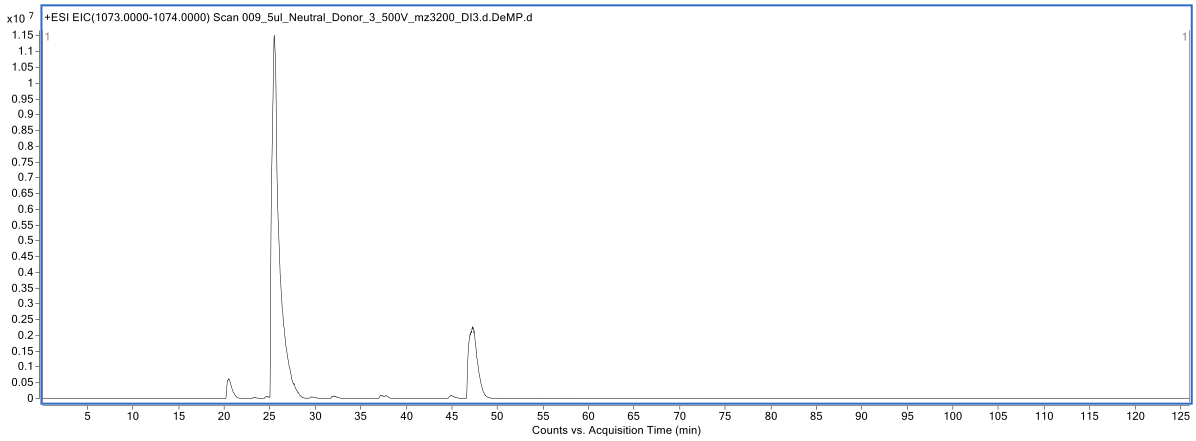

**Supplementary Figure 88. Extracted ion chromatogram at *m/z* 1073-1074 of HMOs labeled with procainamide.**

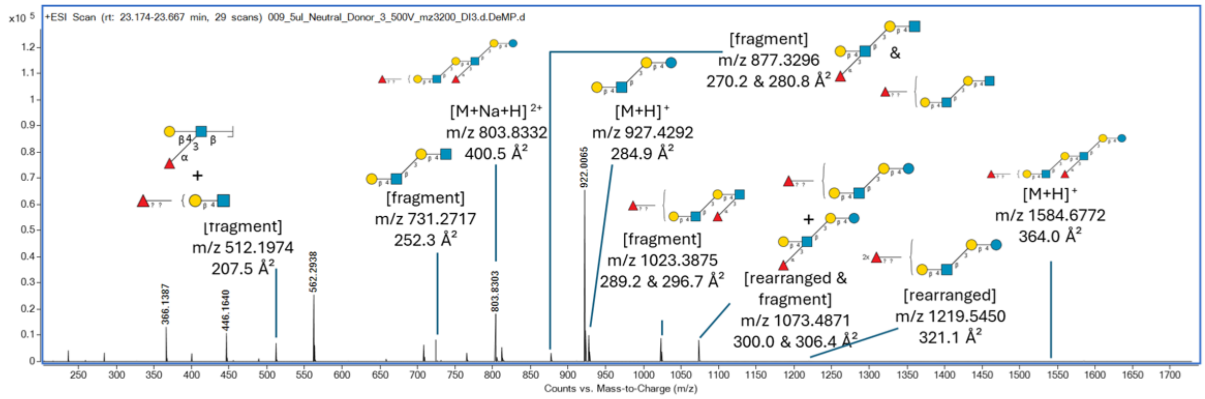

**Supplementary Figure 89. MS spectrum showing an identified HMO structure and its fragment ions eluting at 23.3 min.**

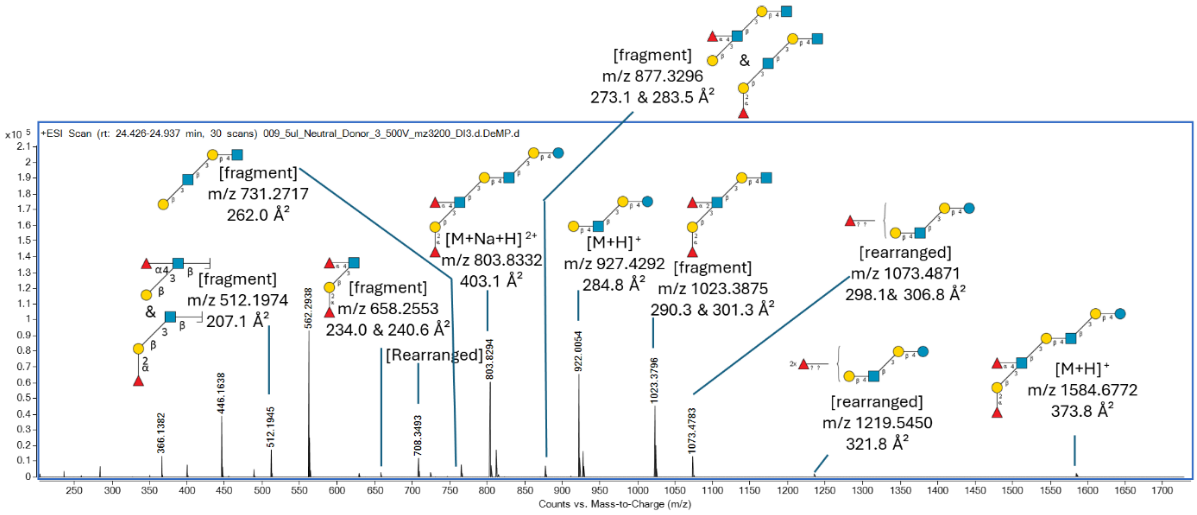

**Supplementary Figure 90. MS spectrum showing an identified HMO structure and its fragment ions eluting at 24.6 min.**

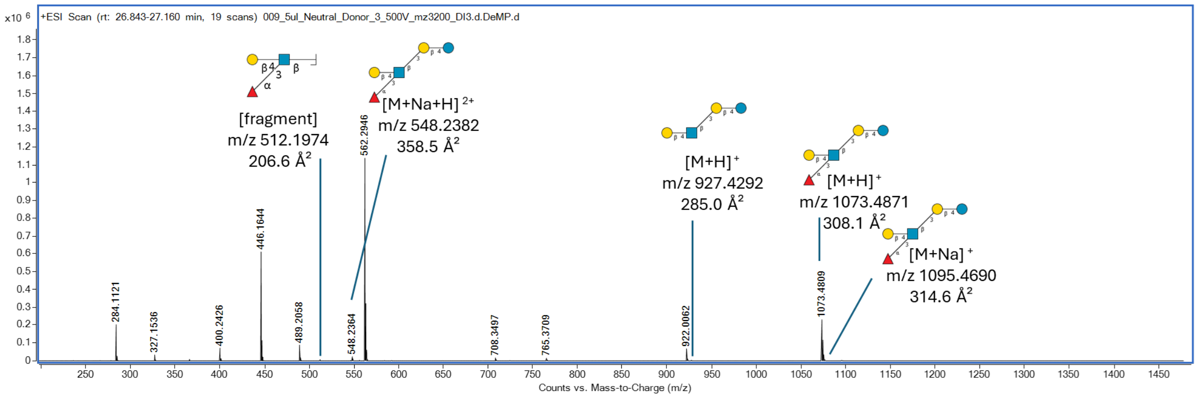

**Supplementary Figure 91. MS spectrum showing an identified HMO structure and its fragment ions eluting at 25.5 min.**

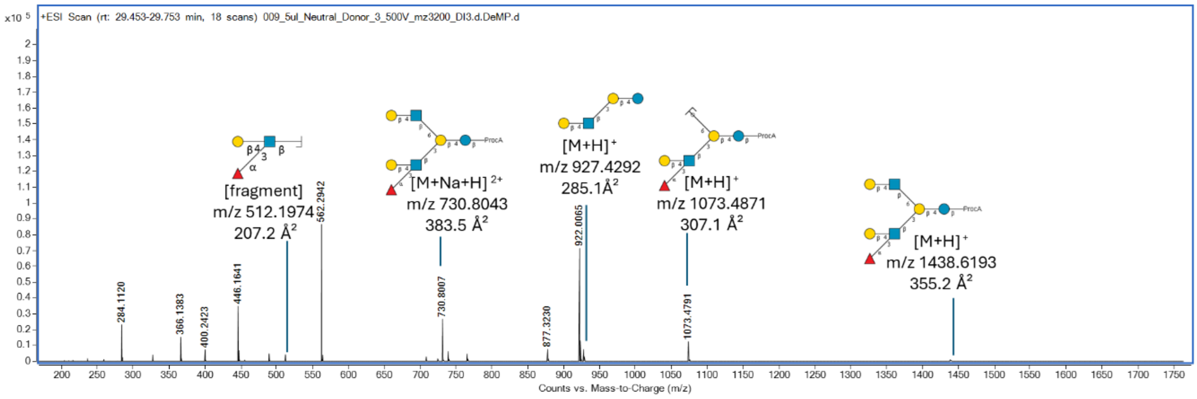

**Supplementary Figure 92. MS spectrum showing an identified HMO structure and its fragment ions eluting at 29.6 min.**

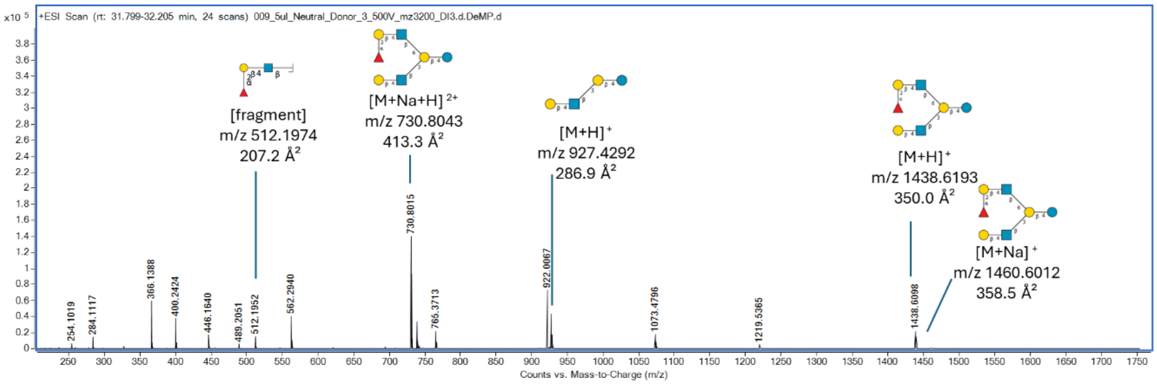

**Supplementary Figure 93. MS spectrum showing an identified HMO structure and its fragment ions eluting at 31.9 min.**

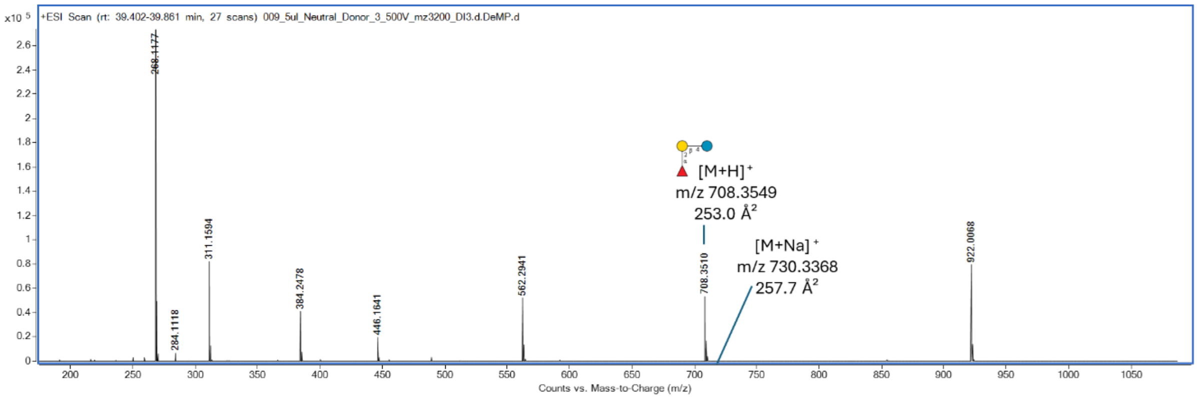

**Supplementary Figure 94. MS spectrum showing an identified HMO structure and its fragment ions eluting at 39.4 min.**

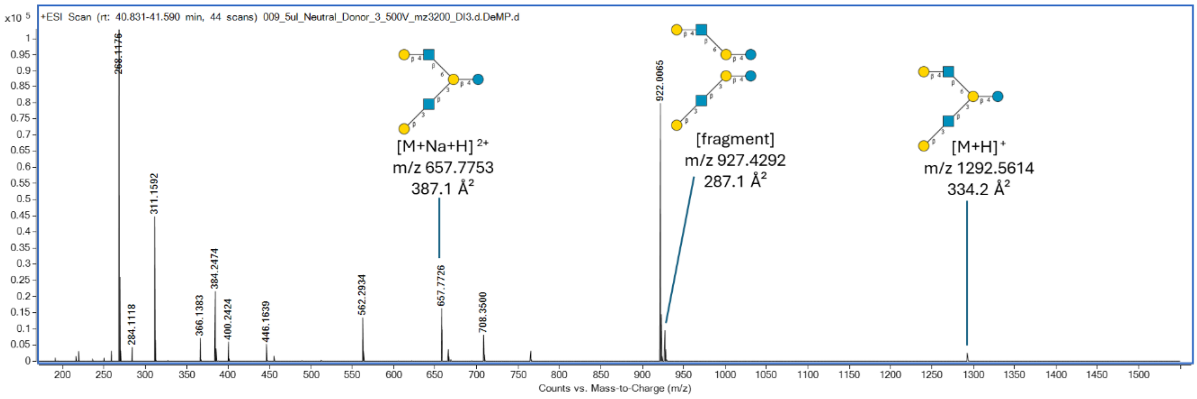

**Supplementary Figure 95. MS spectrum showing an identified HMO structure and its fragment ions eluting at 41.1 min.**

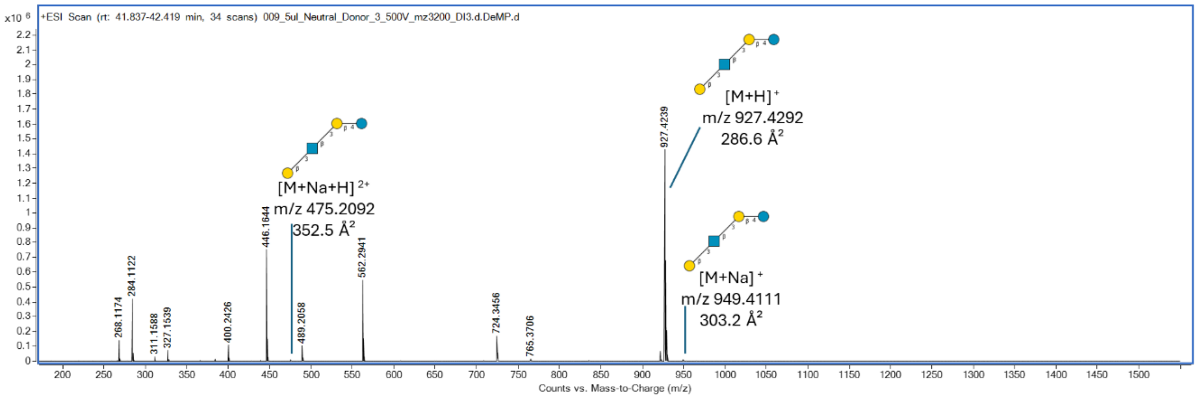

**Supplementary Figure 96. MS spectrum showing an identified HMO structure and its fragment ions eluting at 42.4 min (at the start of chromatographic peak).**

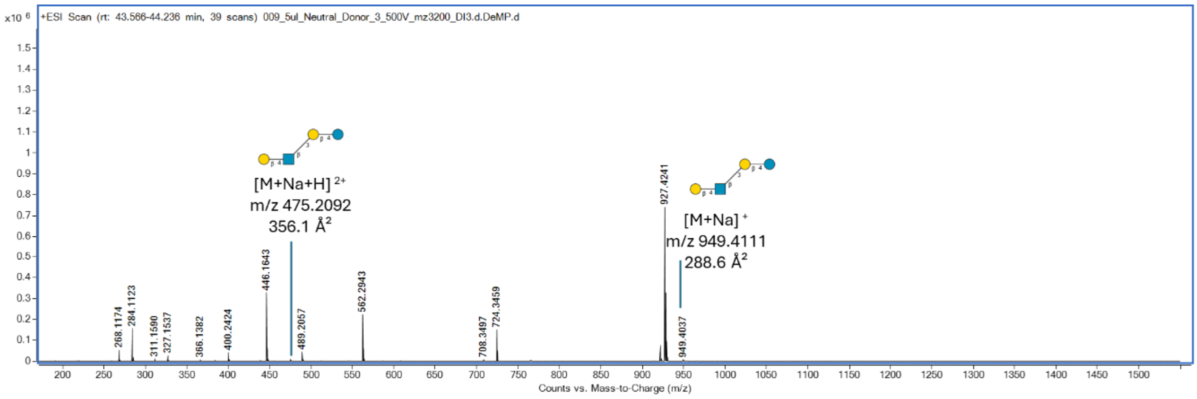

**Supplementary Figure 97. MS spectrum showing an identified HMO structure and its fragment ions eluting at 42.4 min (at the end of the chromatographic peak).**

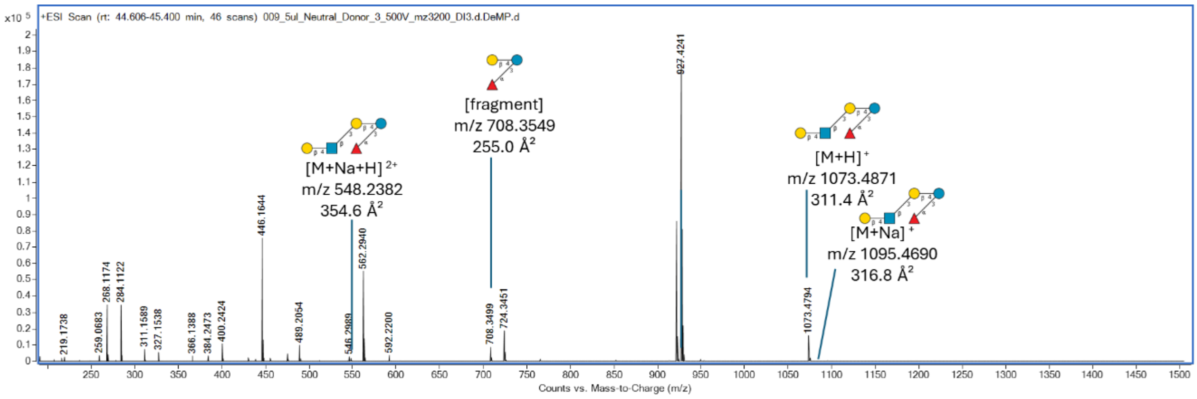

**Supplementary Figure 98. MS spectrum showing an identified HMO structure and its fragment ions eluting at 45.0 min.**

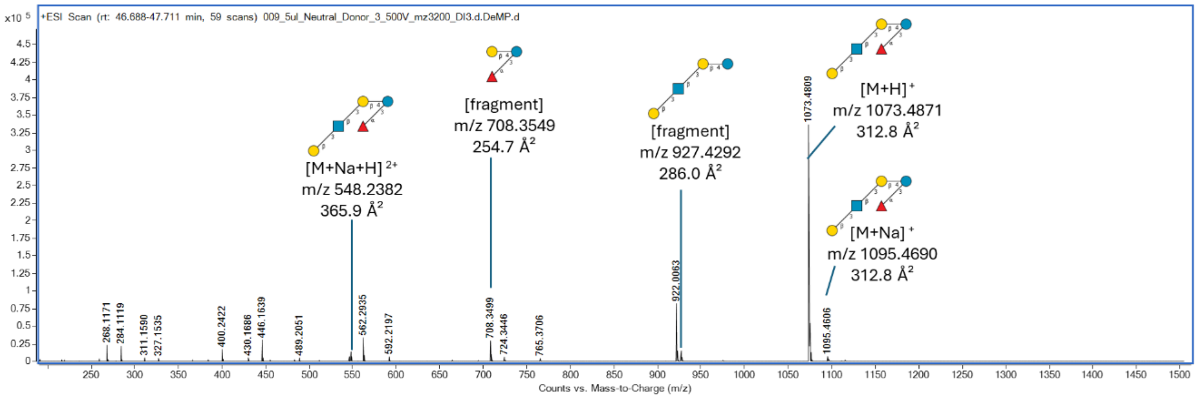

**Supplementary Figure 99. MS spectrum showing an identified HMO structure and its fragment ions eluting at 47.2 min.**

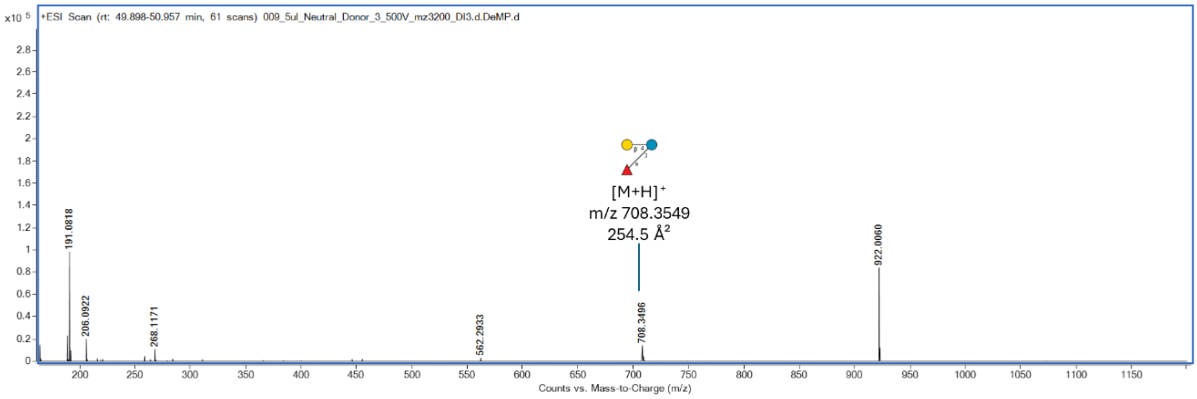

**Supplementary Figure 100. MS spectrum showing an identified HMO structure and its fragment ions eluting at 50.3 min.**

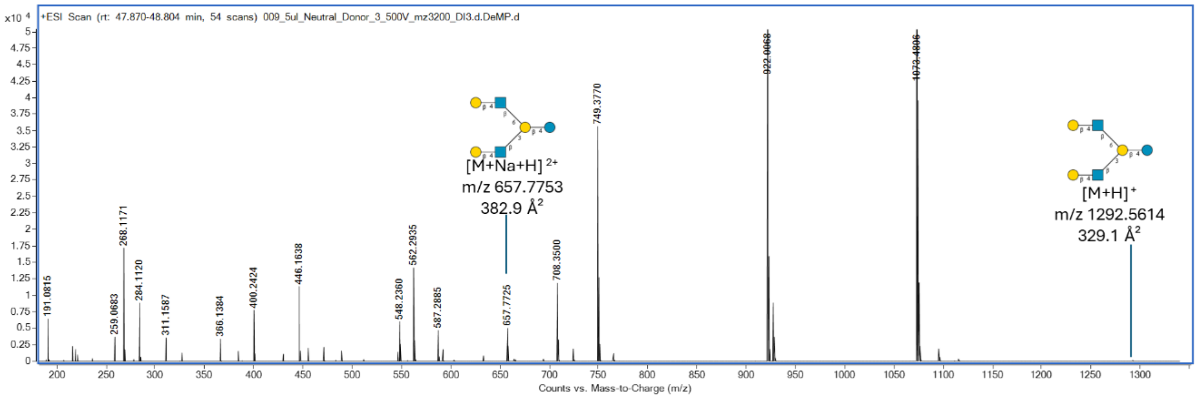

**Supplementary Figure 101. MS spectrum showing an identified HMO structure and its fragment ions eluting at 48.3 min.**

#### 2.9 Neutral Fraction from Donor 4.

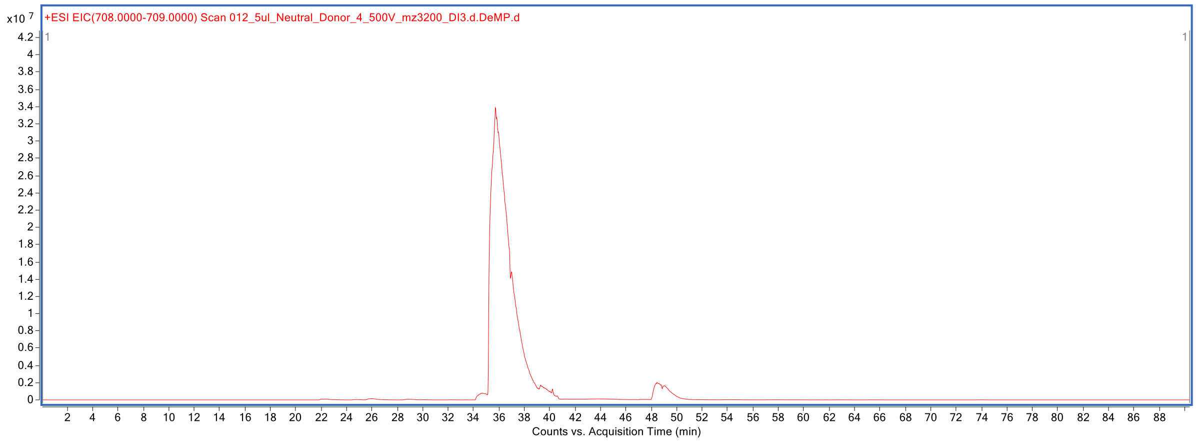

**Supplementary Figure 102. Extracted ion chromatogram at *m/z* 708-709 of HMOs labeled with procainamide.**

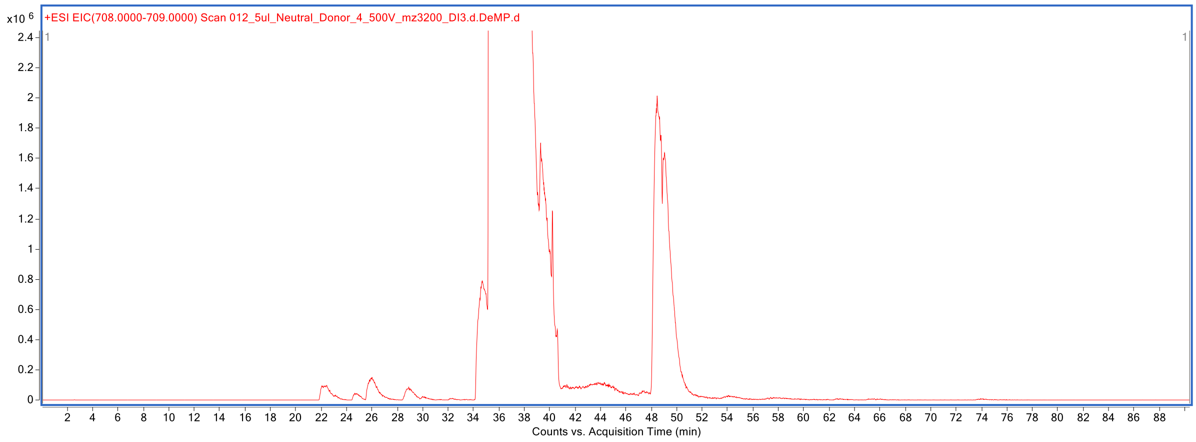

**Supplementary Figure 103. Zoomed-in Extracted ion chromatogram at *m/z* 708-709 of HMOs labeled with procainamide.**

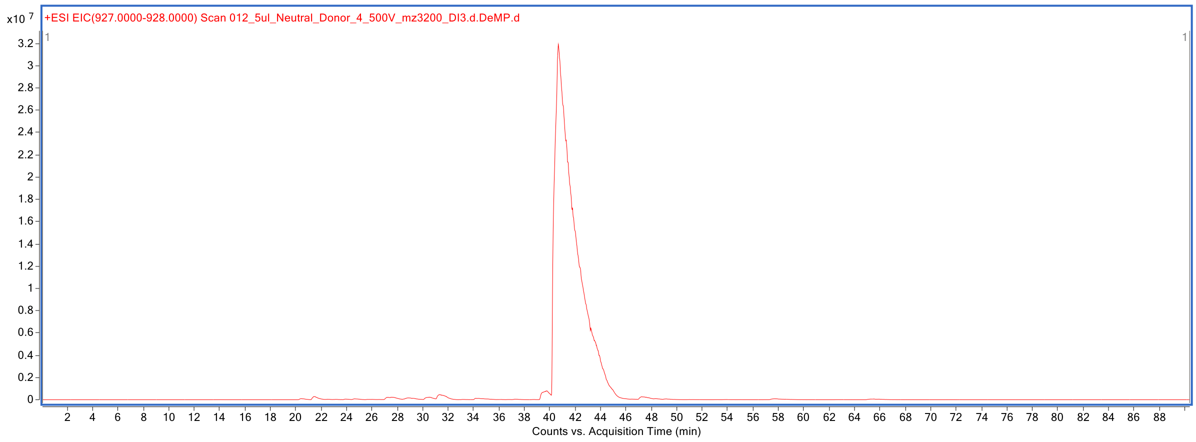

**Supplementary Figure 104. Extracted ion chromatogram at *m/z* 927-928 of HMOs labeled with procainamide.**

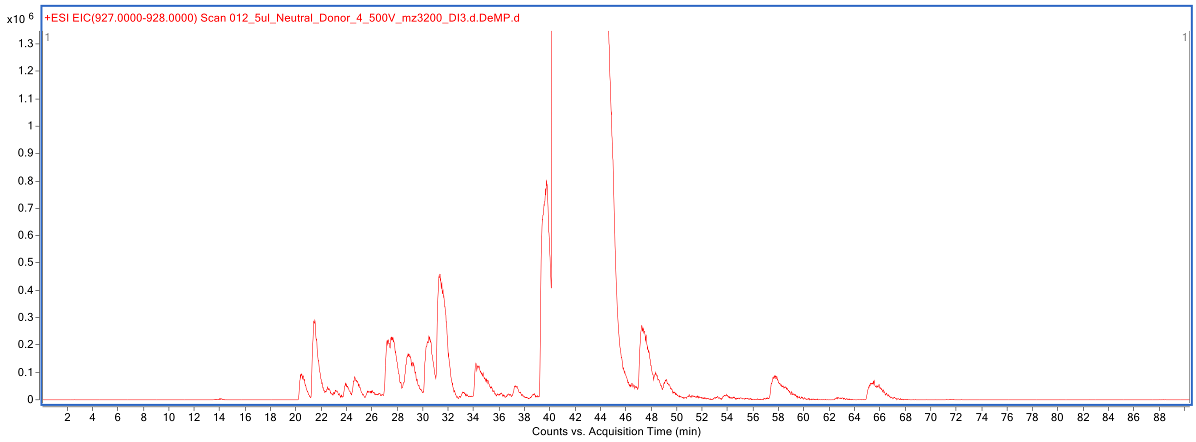

**Supplementary Figure 105. Zoomed-in Extracted ion chromatogram at *m/z* 927-928 of HMOs labeled with procainamide.**

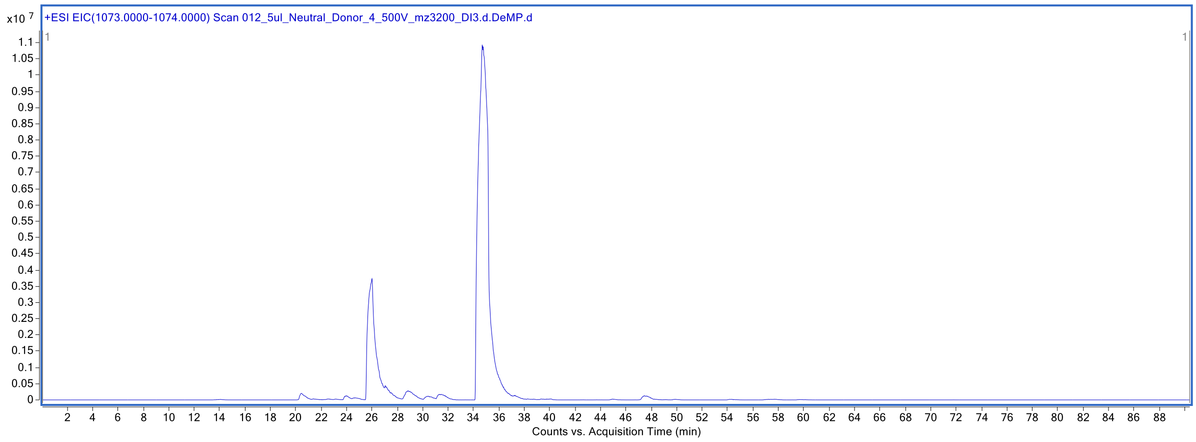

**Supplementary Figure 106. Extracted ion chromatogram at *m/z* 1073-1074 of HMOs labeled with procainamide.**

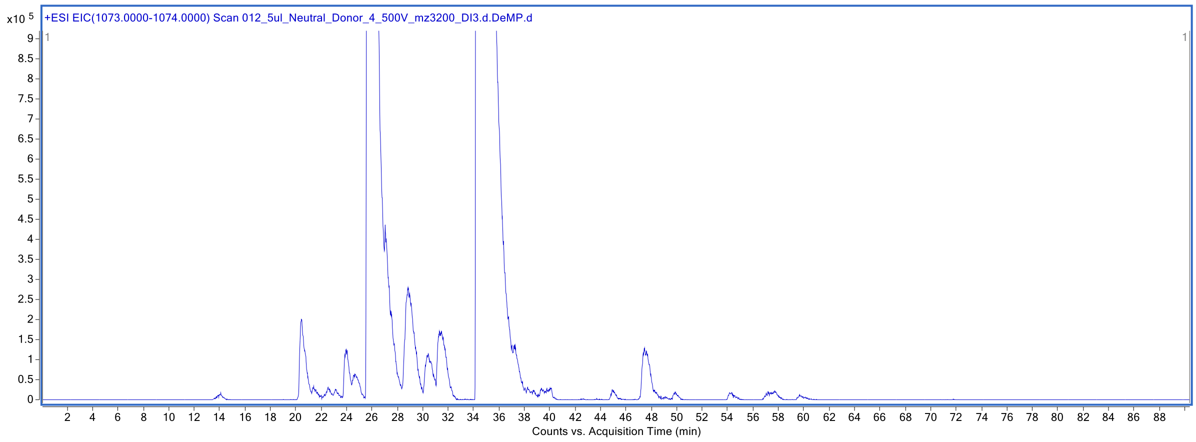

**Supplementary Figure 107. Zoomed-in Extracted ion chromatogram at *m/z* 1073-1074 of HMOs labeled with procainamide.**

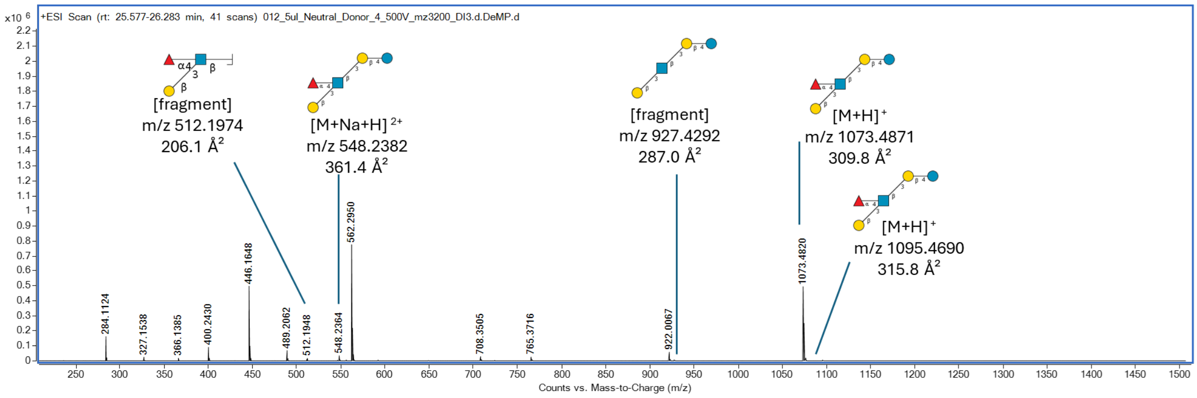

**Supplementary Figure 108. MS spectrum showing an identified HMO structure and its fragment ions eluting at 26.0 min.**

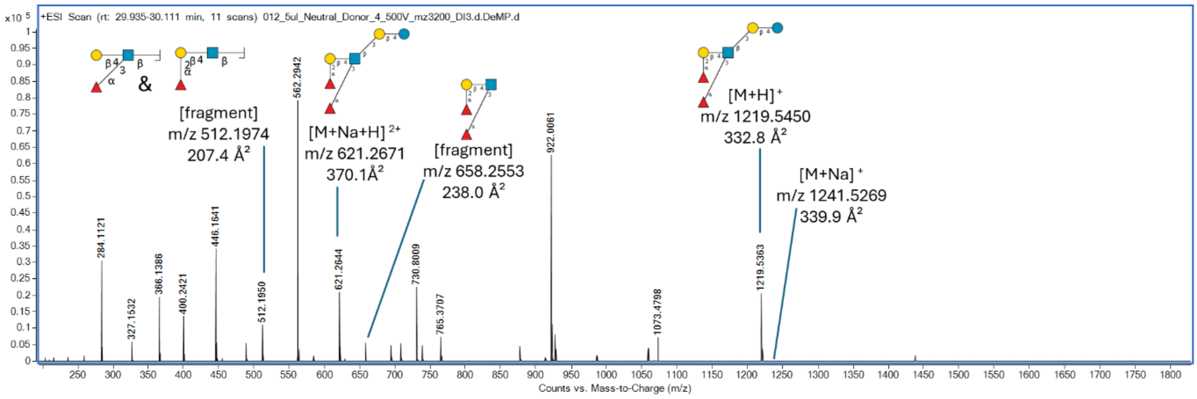

**Supplementary Figure 109. MS spectrum showing an identified HMO structure and its fragment ions eluting at 30.2 min.**

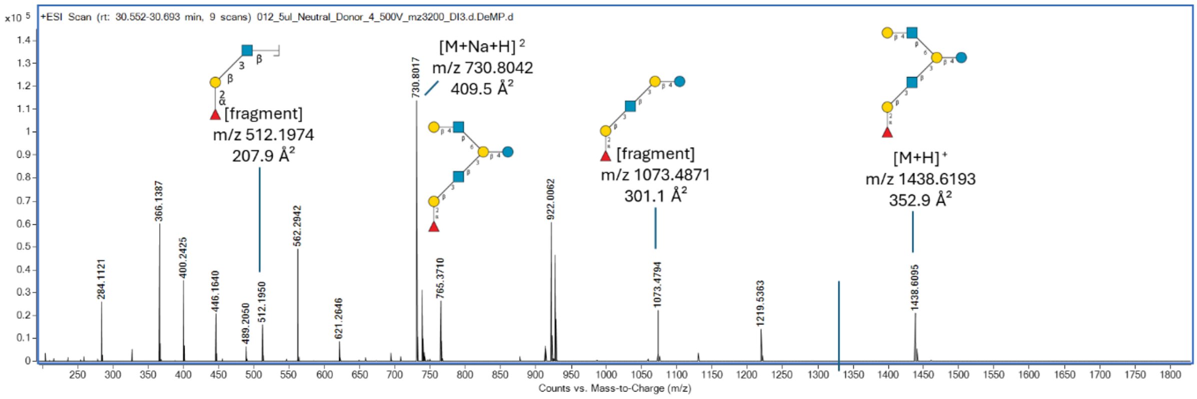

**Supplementary Figure 110. MS spectrum showing an identified HMO structure and its fragment ions eluting at 30.4 min.**

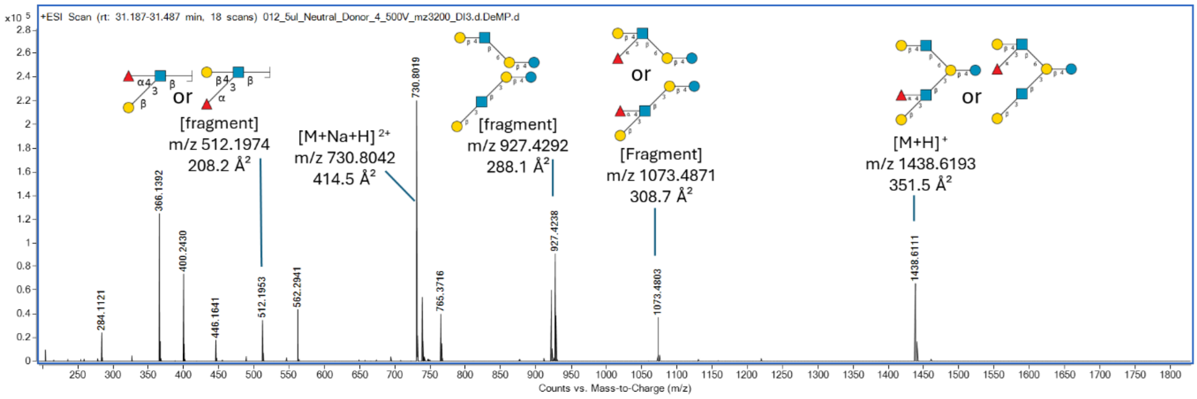

**Supplementary Figure 111. MS spectrum showing an identified HMO structure and its fragment ions eluting at 31.4 min.**

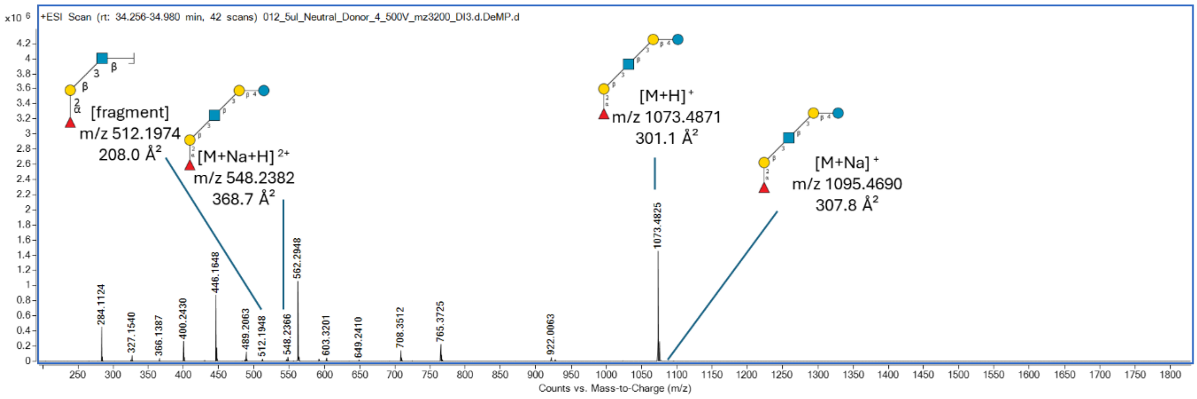

**Supplementary Figure 112. MS spectrum showing an identified HMO structure and its fragment ions eluting at 34.7 min.**

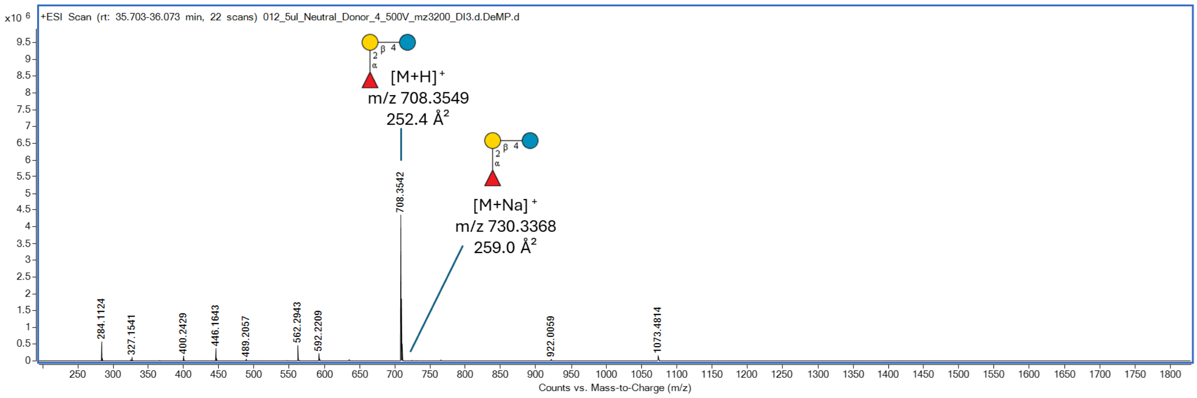

**Supplementary Figure 113. MS spectrum showing an identified HMO structure and its fragment ions eluting at 35.7 min.**

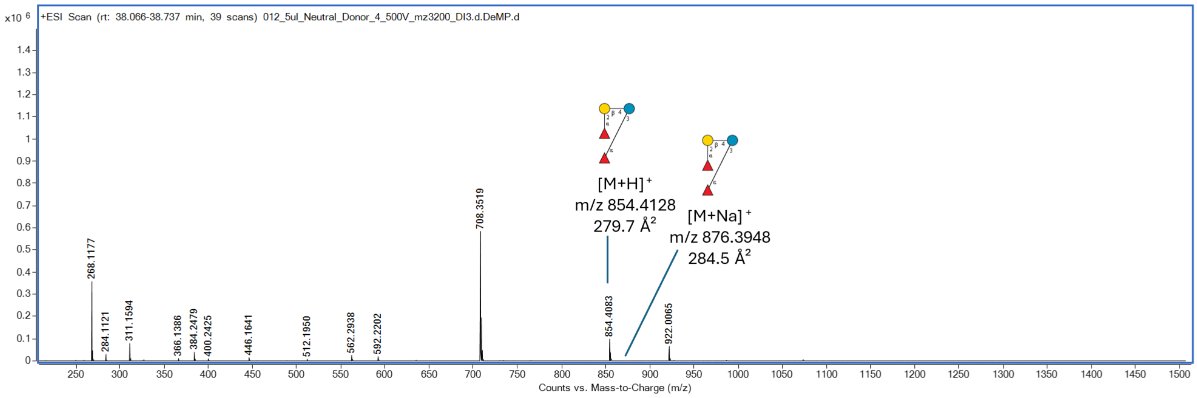

**Supplementary Figure 114. MS spectrum showing an identified HMO structure and its fragment ions eluting at 38.4 min.**

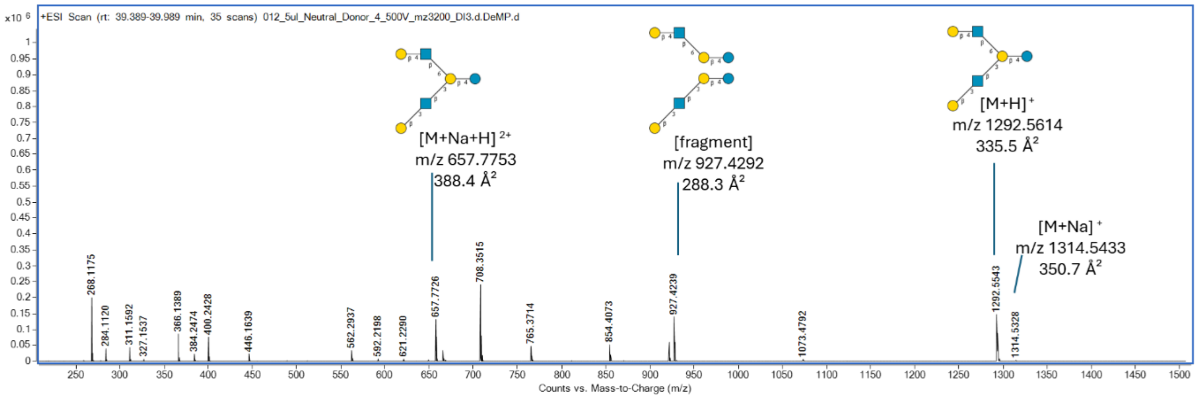

**Supplementary Figure 115. MS spectrum showing an identified HMO structure and its fragment ions eluting at 39.9 min.**

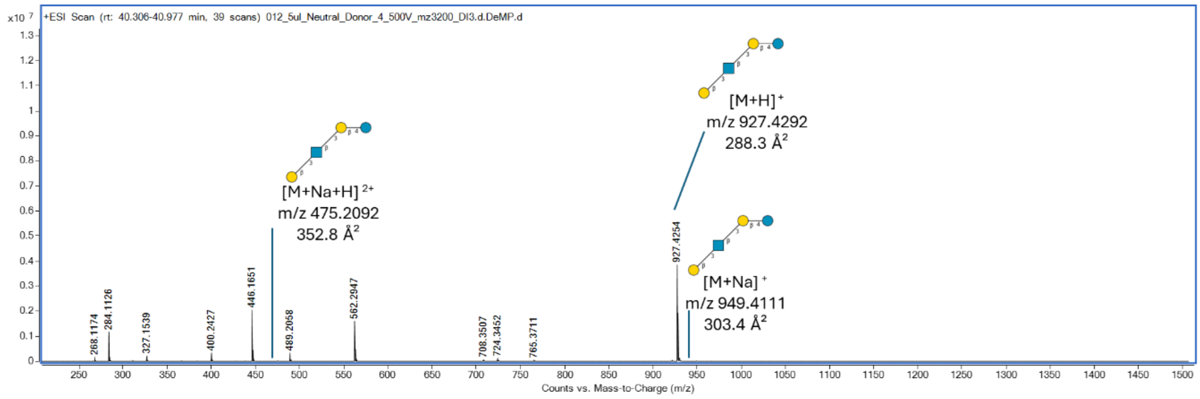

**Supplementary Figure 116. MS spectrum showing an identified HMO structure and its fragment ions eluting at 40.7 min.**

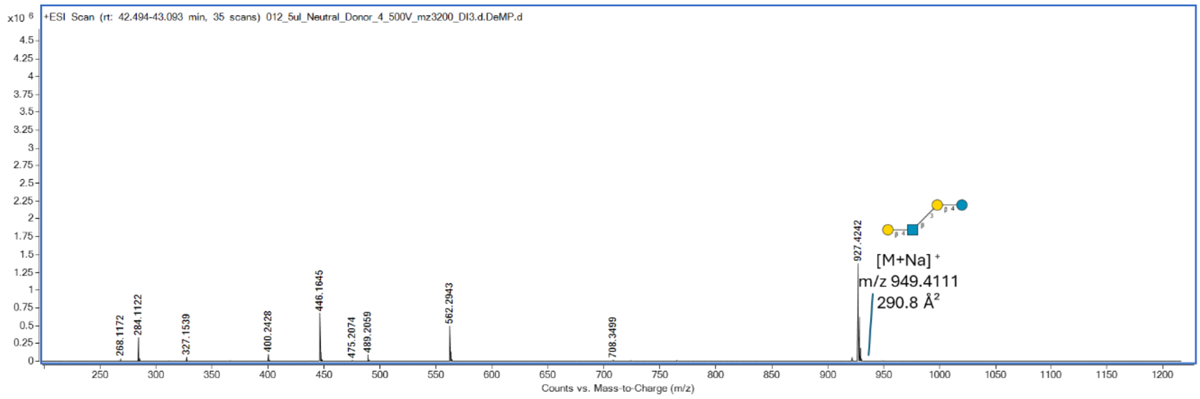

**Supplementary Figure 117. MS spectrum showing an identified HMO structure and its fragment ions eluting at 40.7 min.**

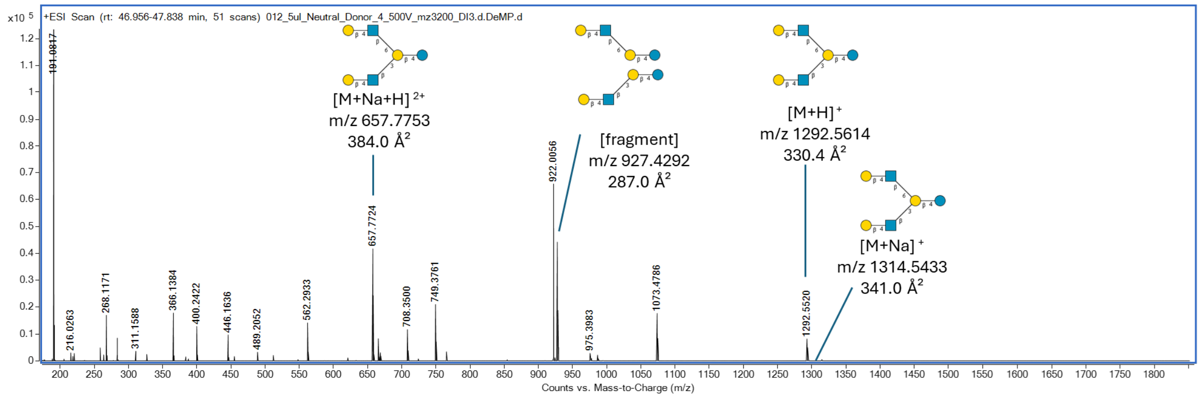

**Supplementary Figure 118. MS spectrum showing an identified HMO structure and its fragment ions eluting at 47.3 min.**

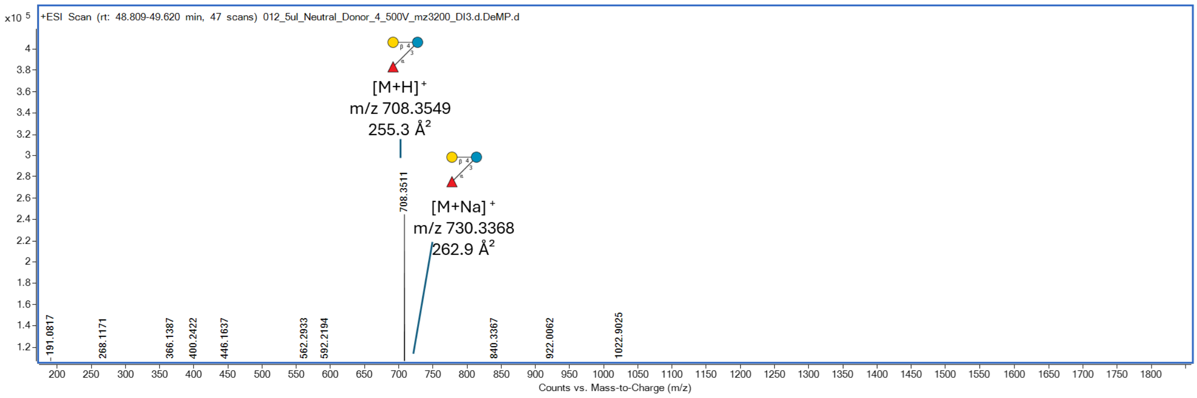

**Supplementary Figure 119. MS spectrum showing an identified HMO structure and its fragment ions eluting at 48.5 min.**

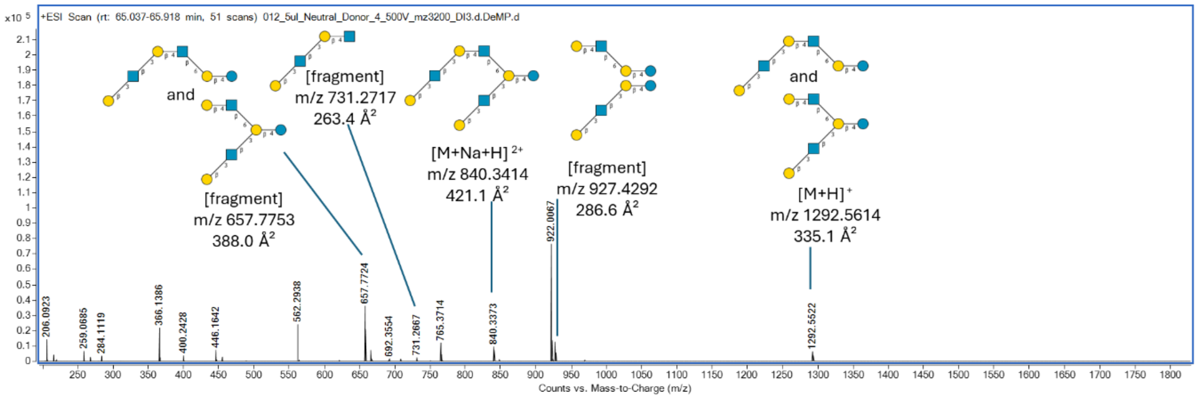

**Supplementary Figure 120. MS spectrum showing an identified HMO structure and its fragment ions eluting at 65.5 min.**

#### 2.10 Neutral Fraction from Donor 5.

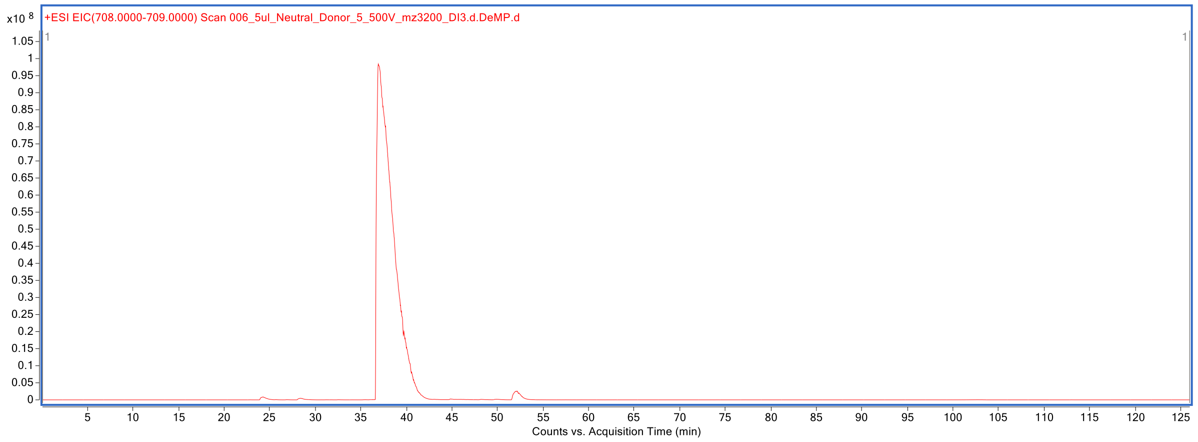

**Supplementary Figure 121. Extracted ion chromatogram at *m/z* 708-709 of HMOs labeled with procainamide.**

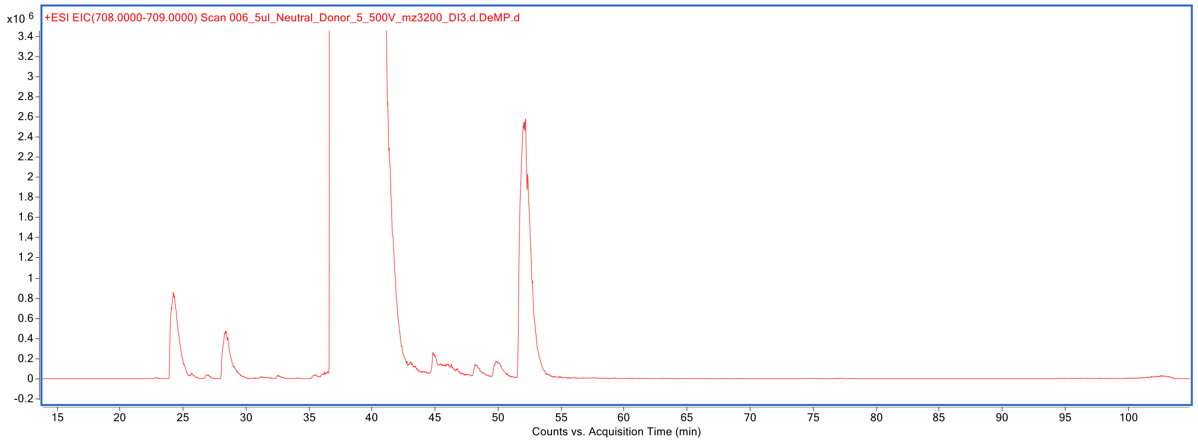

**Supplementary Figure 122. Zoomed-in Extracted ion chromatogram at *m/z* 708-709 of HMOs labeled with procainamide.**

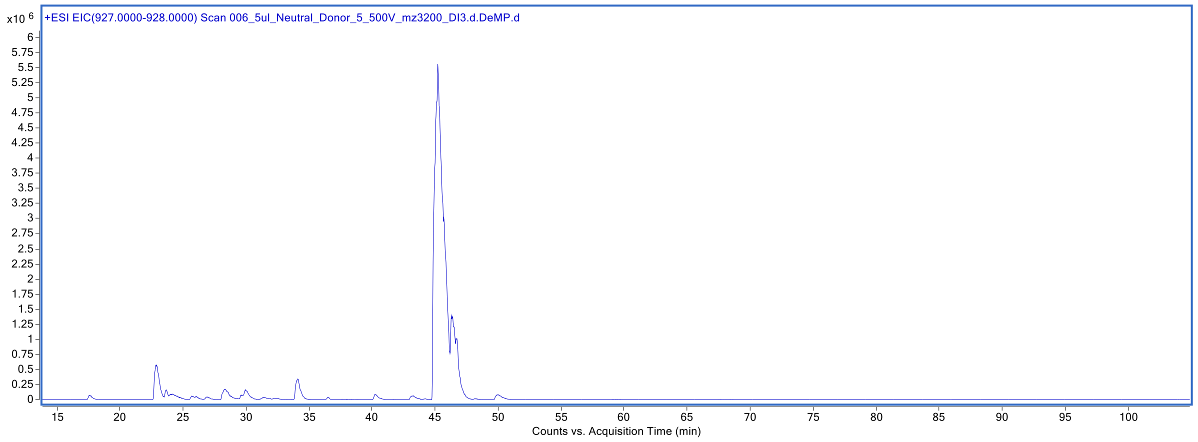

**Supplementary Figure 123. Extracted ion chromatogram at *m/z* 927-928 of HMOs labeled with procainamide.**

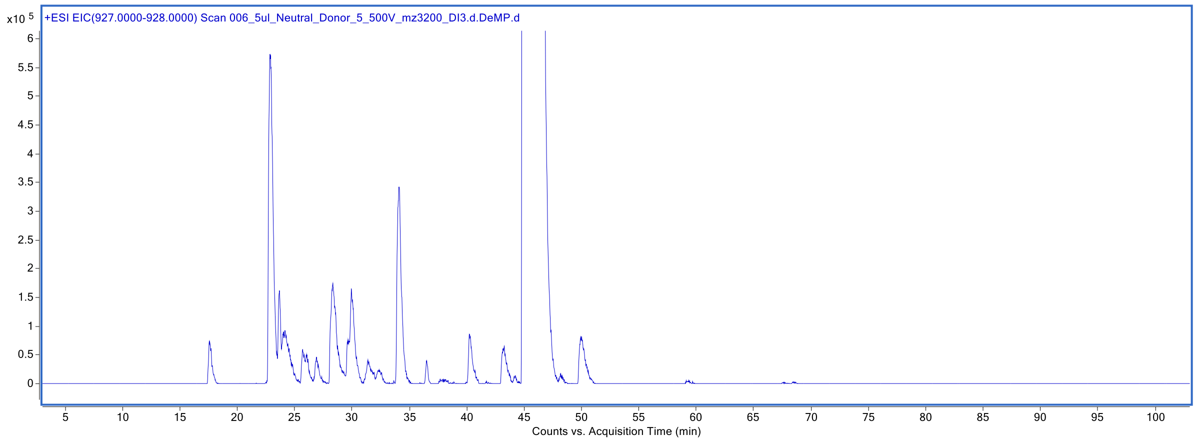

**Supplementary Figure 124. Zoomed-in Extracted ion chromatogram at *m/z* 927-928 of HMOs labeled with procainamide.**

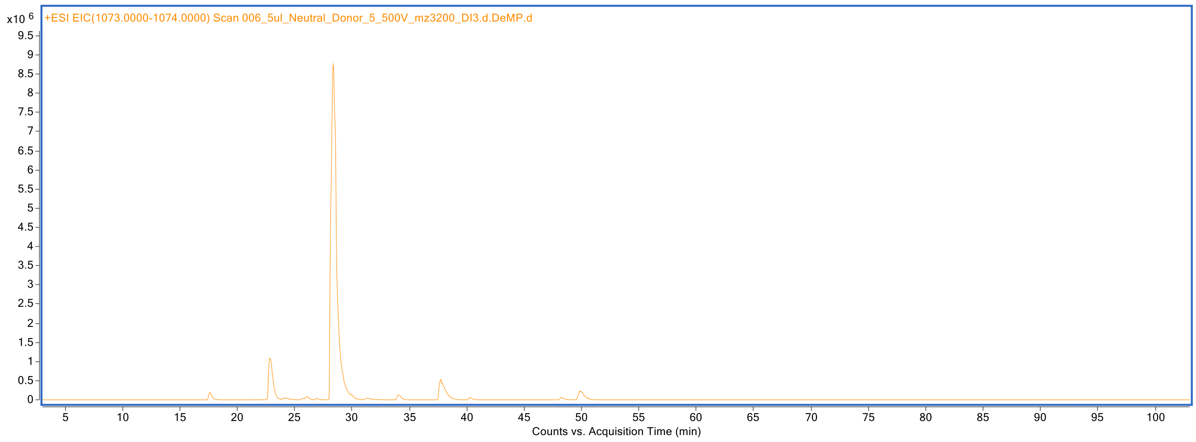

**Supplementary Figure 125. Extracted ion chromatogram at *m/z* 1073-1074 of HMOs labeled with procainamide.**

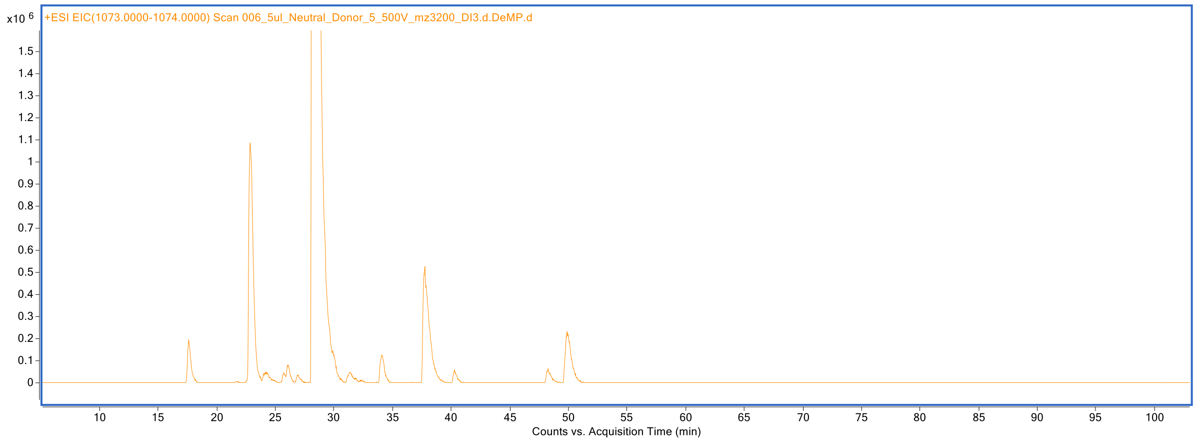

**Supplementary Figure 126. Zoomed-in Extracted ion chromatogram at *m/z* 1073-1074 of HMOs labeled with procainamide.**

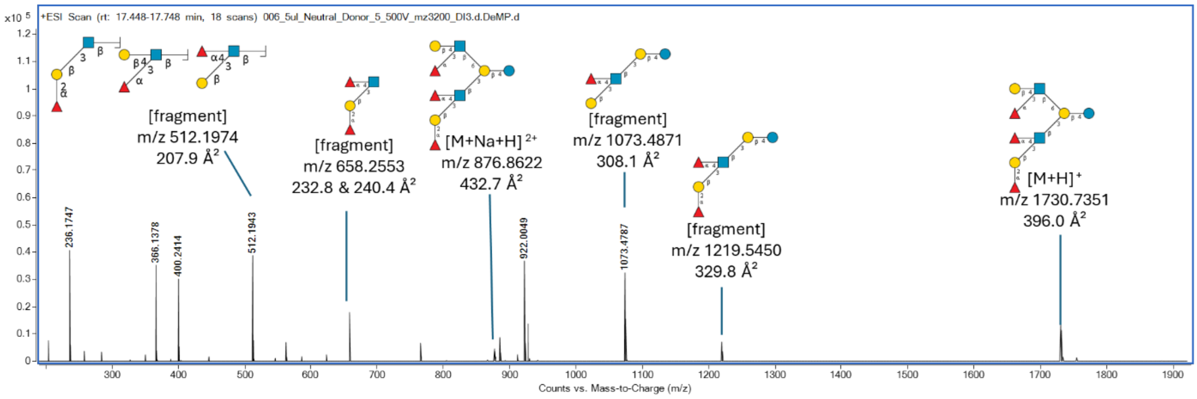

**Supplementary Figure 127. MS spectrum showing an identified HMO structure and its fragment ions eluting at 17.6 min.**

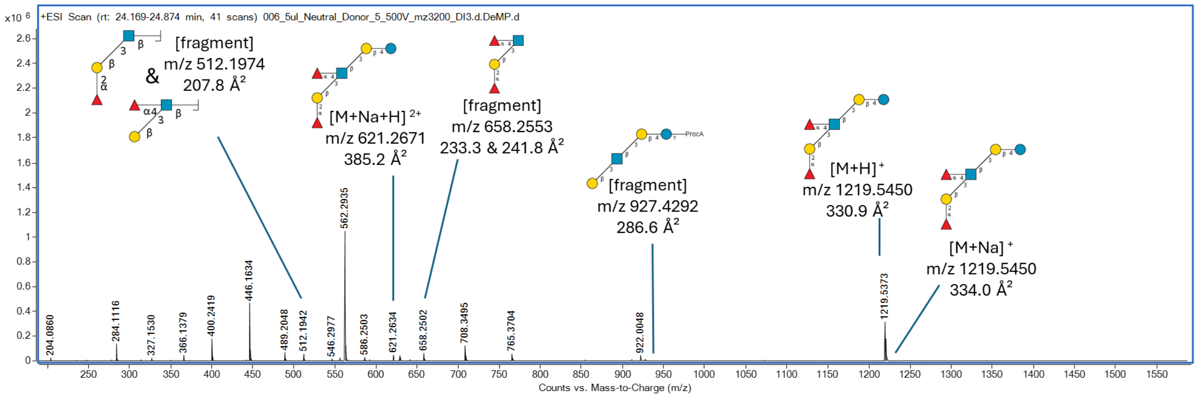

**Supplementary Figure 128. MS spectrum showing an identified HMO structure and its fragment ions eluting at 24.3 min.**

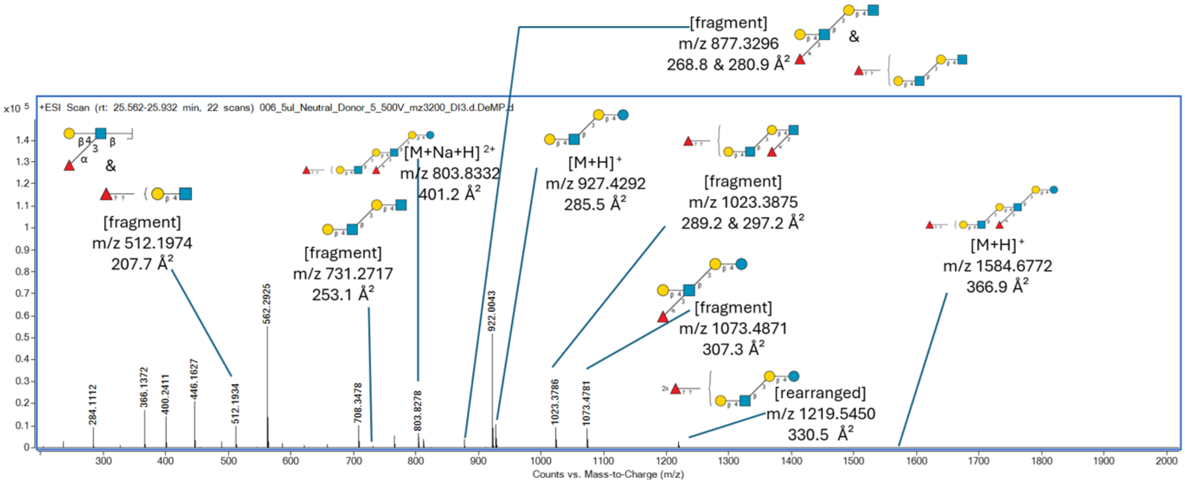

**Supplementary Figure 129. MS spectrum showing an identified HMO structure and its fragment ions eluting at 25.7 min.**

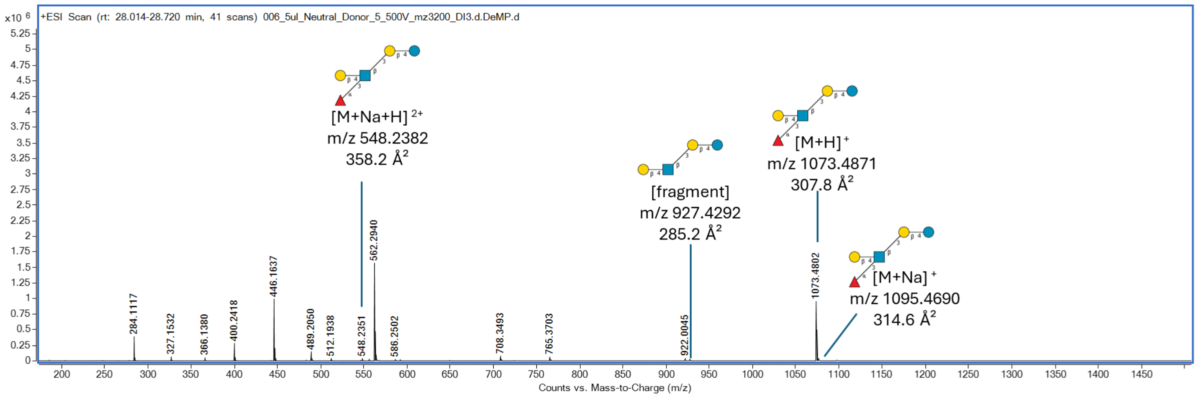

**Supplementary Figure 130. MS spectrum showing an identified HMO structure and its fragment ions eluting at 28.3 min.**

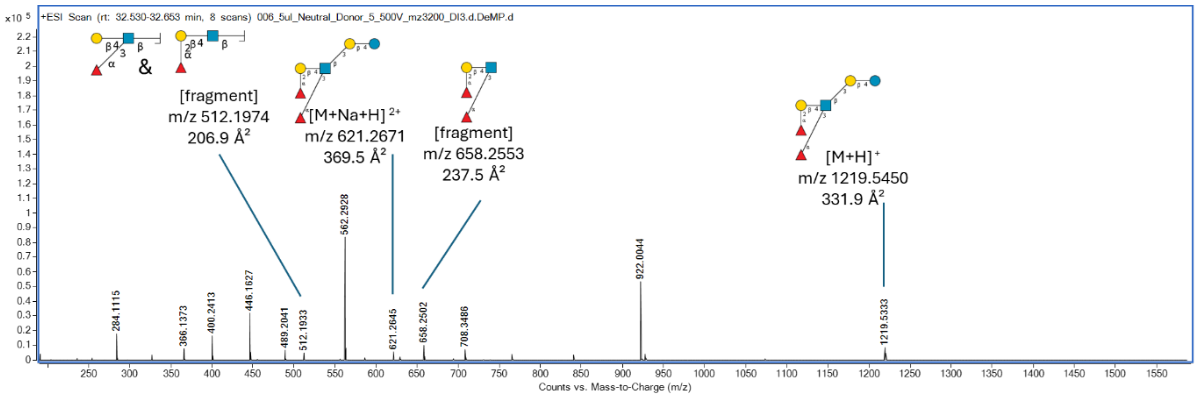

**Supplementary Figure 131. MS spectrum showing an identified HMO structure and its fragment ions eluting at 32.6 min.**

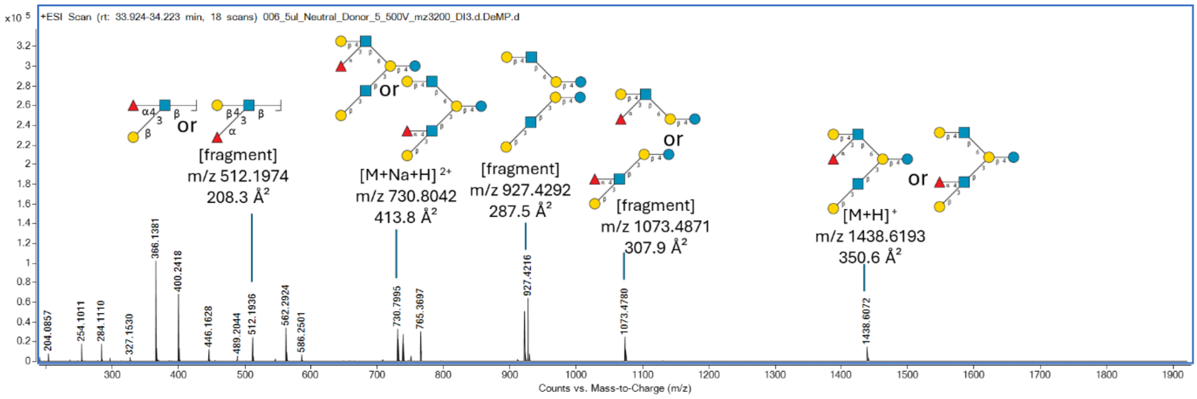

**Supplementary Figure 132. MS spectrum showing an identified HMO structure and its fragment ions eluting at 34.0 min.**

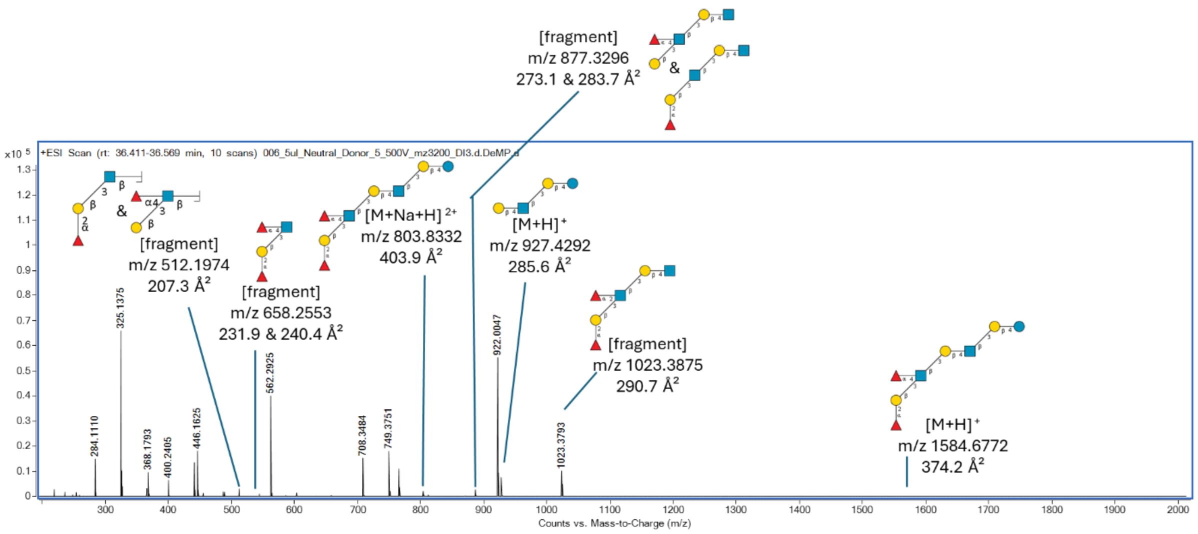

**Supplementary Figure 133. MS spectrum showing an identified HMO structure and its fragment ions eluting at 36.5 min.**

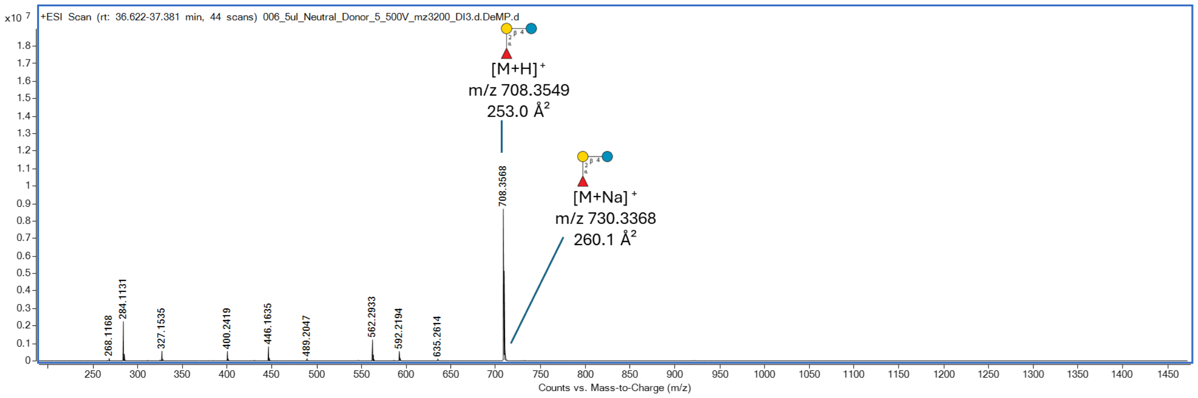

**Supplementary Figure 134. MS spectrum showing an identified HMO structure and its fragment ions eluting at 36.9 min.**

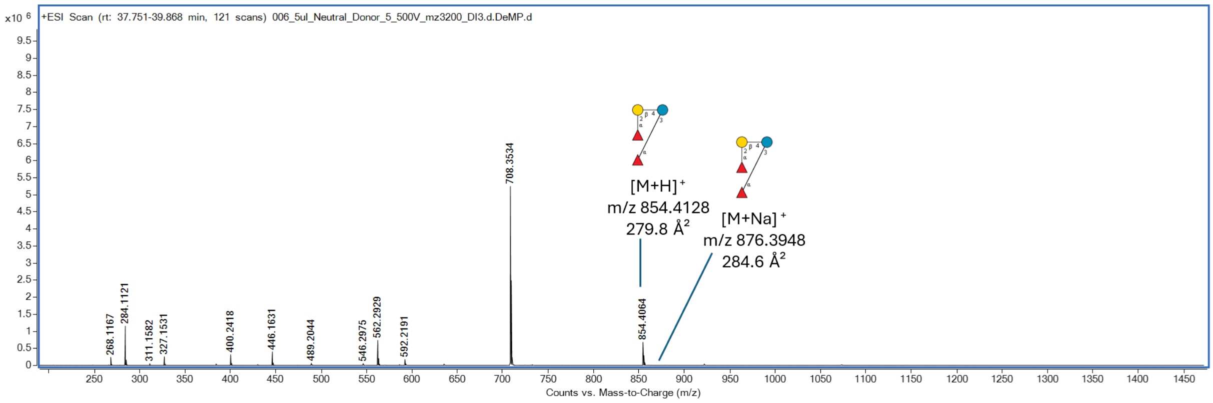

**Supplementary Figure 135. MS spectrum showing an identified HMO structure and its fragment ions eluting at 39.3 min.**

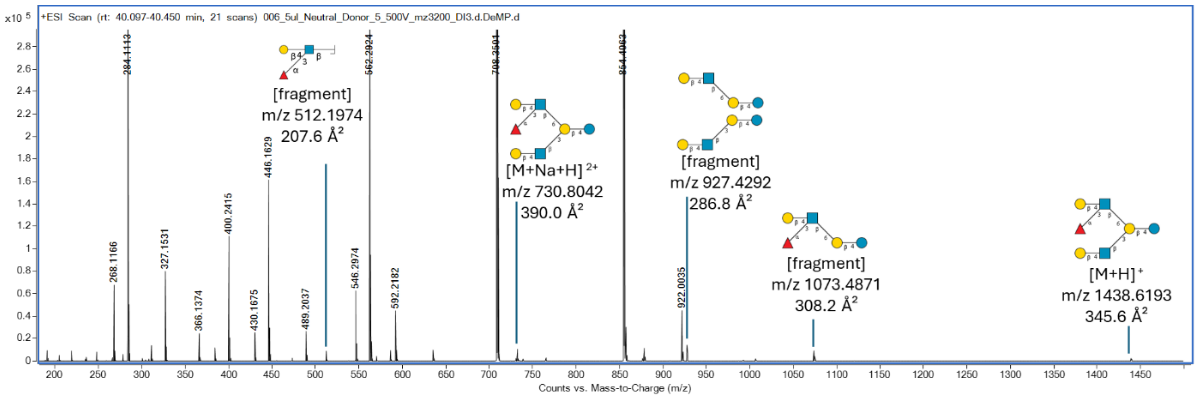

**Supplementary Figure 136. MS spectrum showing an identified HMO structure and its fragment ions eluting at 40.4 min.**

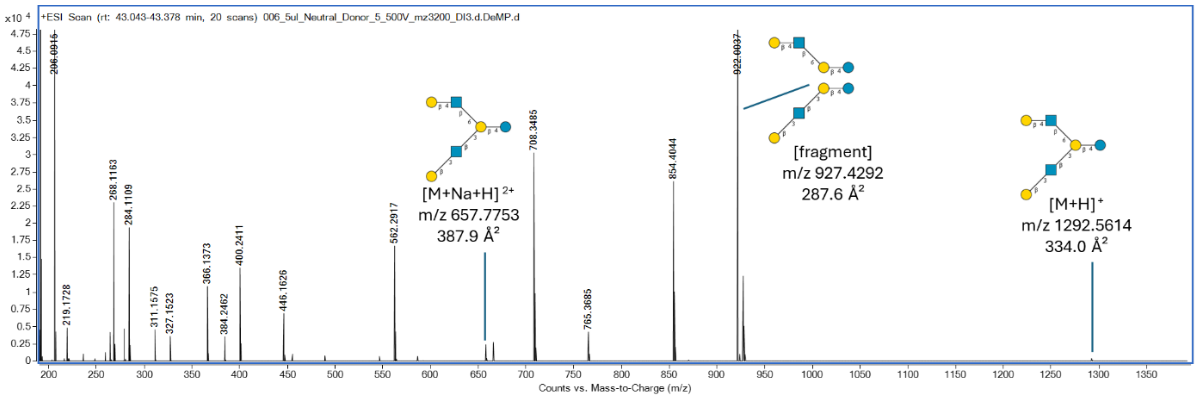

**Supplementary Figure 137. MS spectrum showing an identified HMO structure and its fragment ions eluting at 42.2 min.**

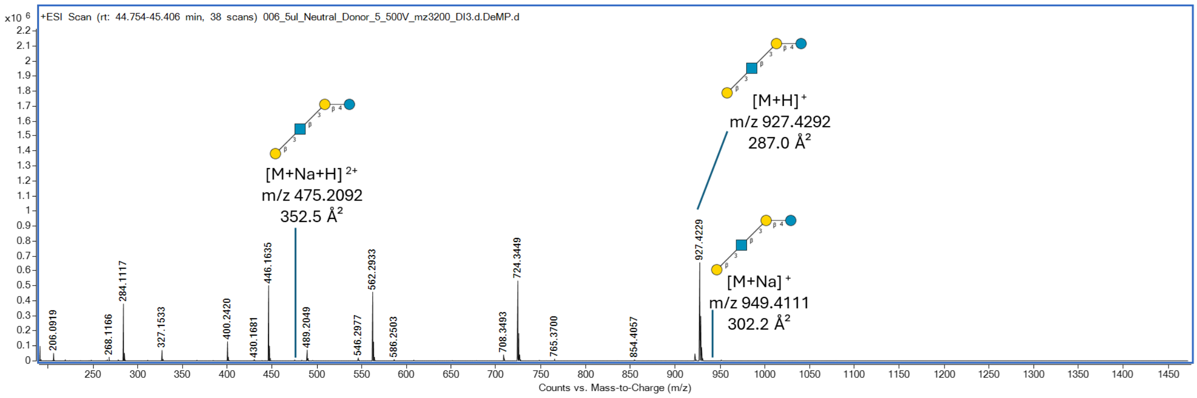

**Supplementary Figure 138. MS spectrum showing an identified HMO structure and its fragment ions eluting at 45.2 min.**

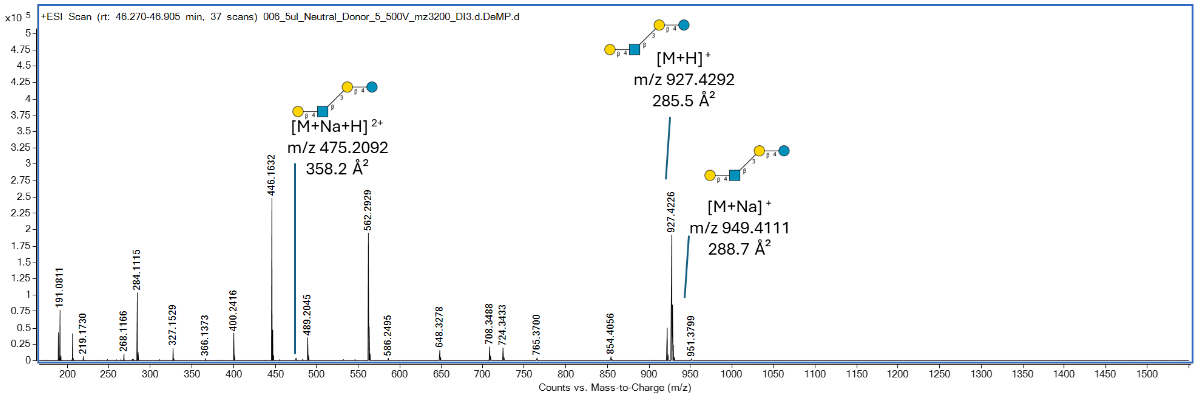

**Supplementary Figure 139. MS spectrum showing an identified HMO structure and its fragment ions eluting at 46.3 min.**

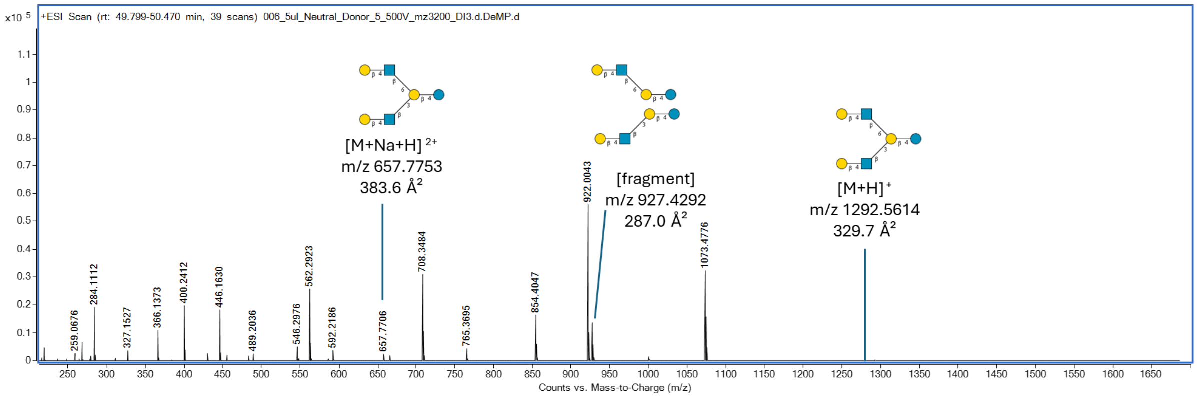

**Supplementary Figure 140. MS spectrum showing an identified HMO structure and its fragment ions eluting at 50.1 min.**

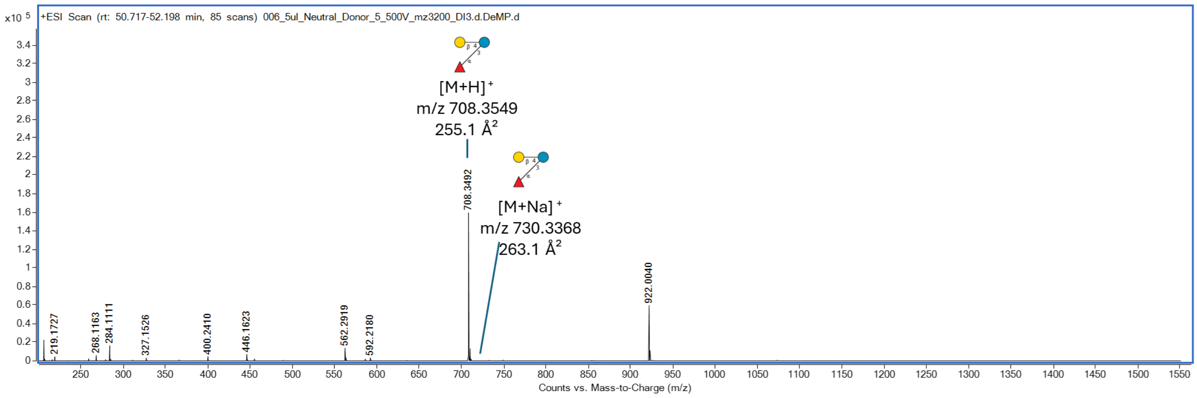

**Supplementary Figure 141. MS spectrum showing an identified HMO structure and its fragment ions eluting at 52.2 min.**

### 3. HMO standards for validation of assigned structures in breast milk.

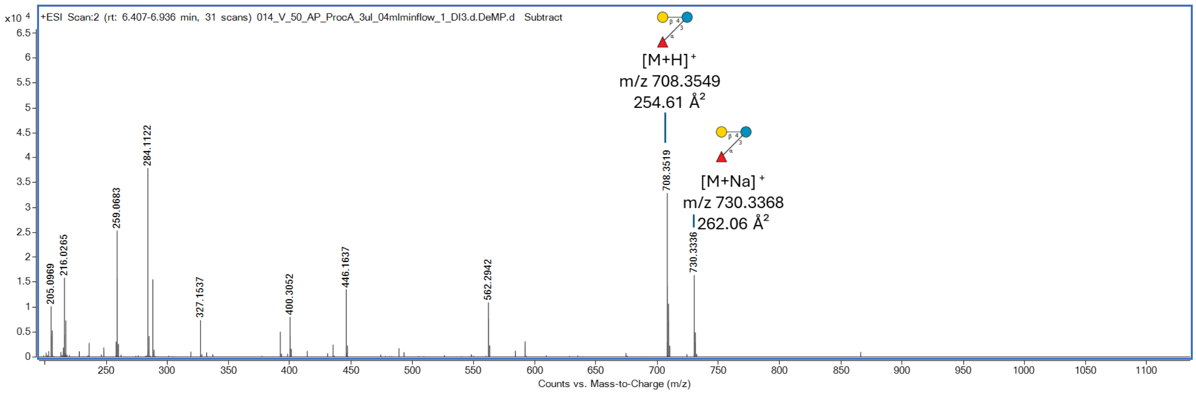

**Supplementary Figure 142. MS spectrum of the synthetic standard HMO23 showing intact and fragment ions with their theoretical *m/z* and experimentally determined CCS values (n=3 measurements).**

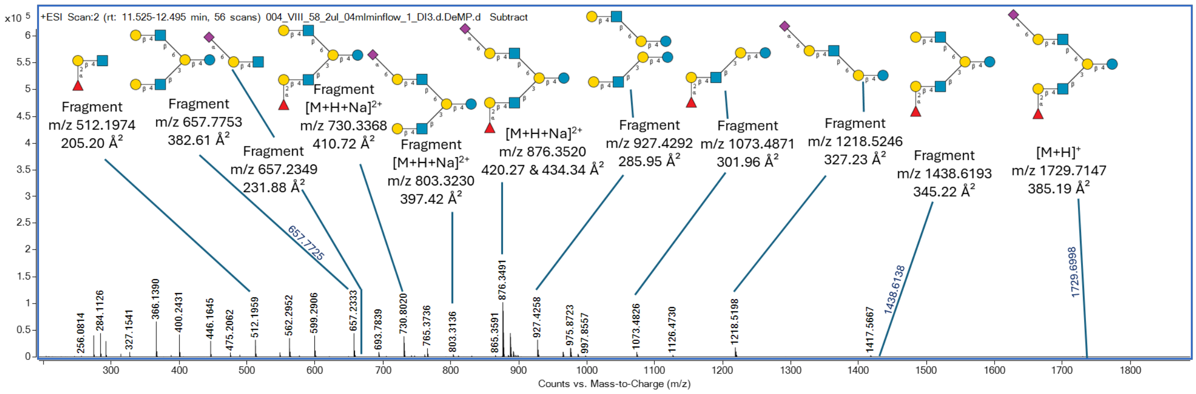

**Supplementary Figure 143. MS spectrum of the synthetic standard HMO24 showing intact and fragment ions with their theoretical *m/z* and experimentally determined CCS values (n=3 measurements).**

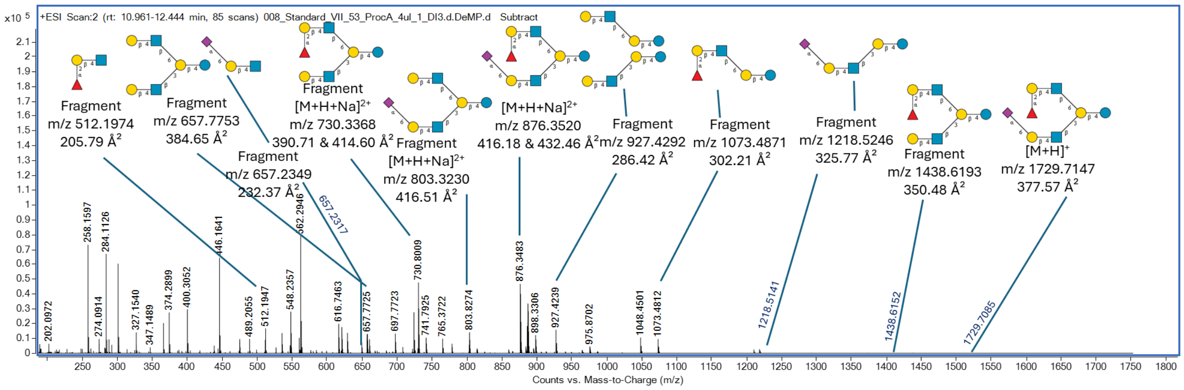

**Supplementary Figure 144. MS spectrum of the synthetic standard HMO25 showing intact and fragment ions with their theoretical *m/z* and experimentally determined CCS values (n=3 measurements).**

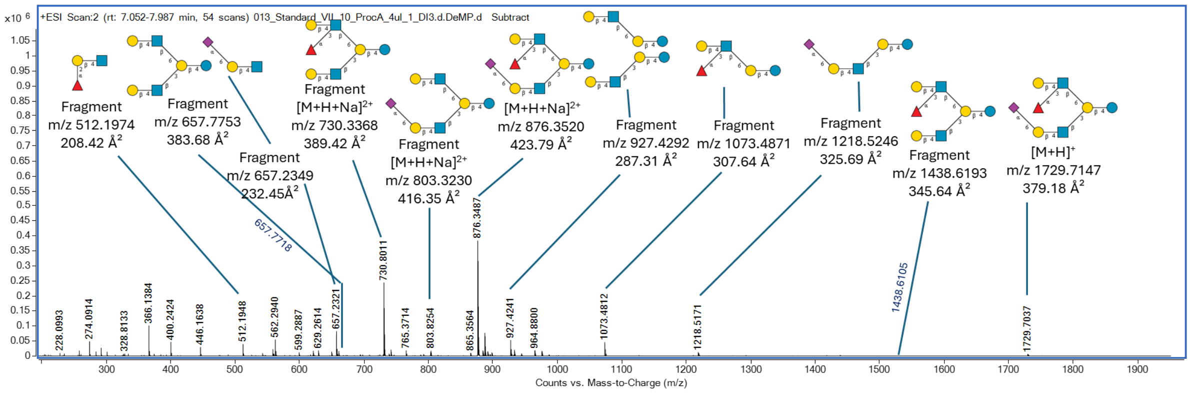

**Supplementary Figure 145. MS spectrum of the synthetic standard HMO26 showing intact and fragment ions with their theoretical *m/z* and experimentally determined CCS values (n=3 measurements).**

### 4. Possible isomeric structures of a human milk octasaccharide.

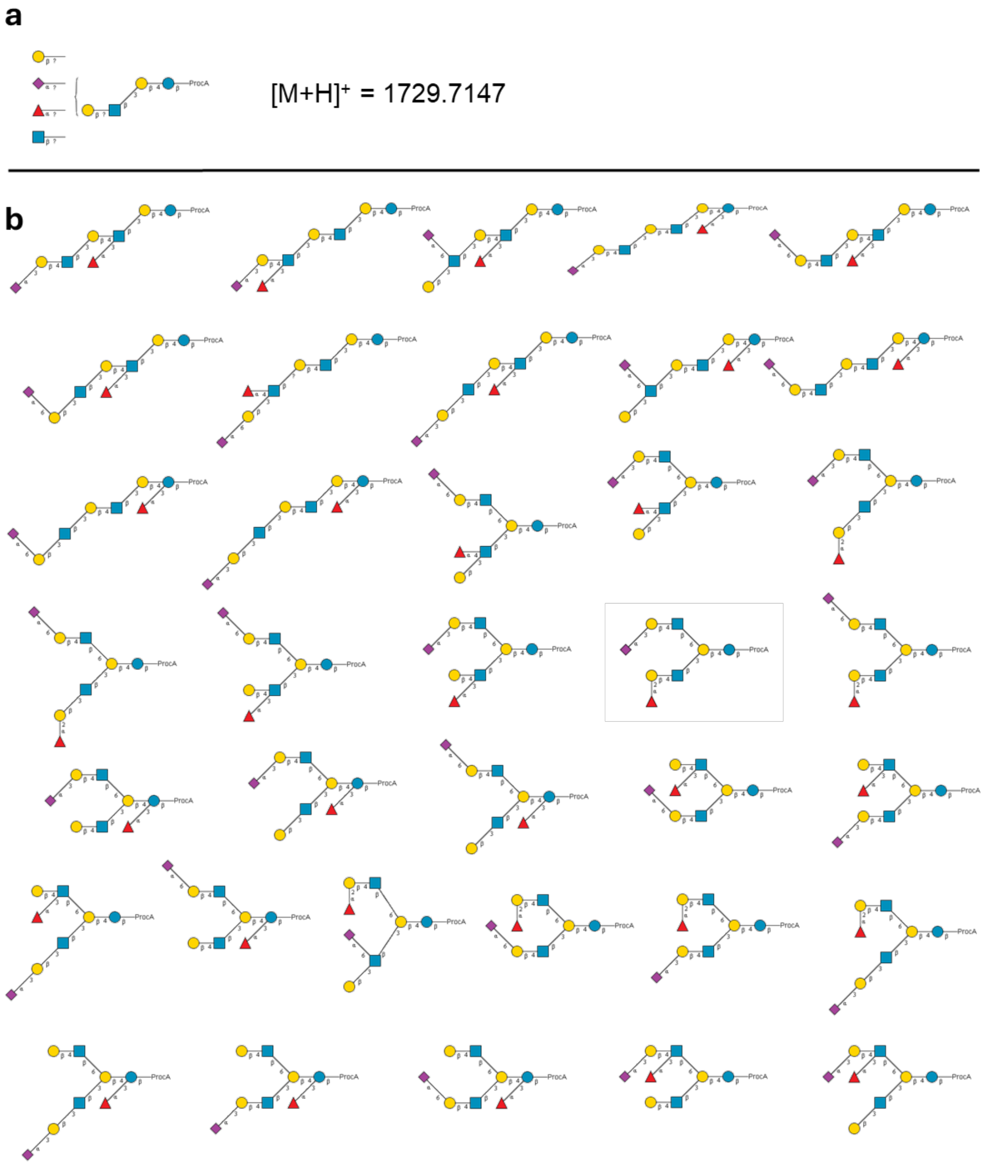

**Supplementary Figure 146. Composition of an octasaccharide with an *m/z* value of 1729.7147 (a) and 36 biosynthetically feasible isomeric structures with this composition (b).**

### 5. Fingerprinting identification of glycans.

#### 5.1 ATD fingerprints of synthetic *N*-glycan standards.

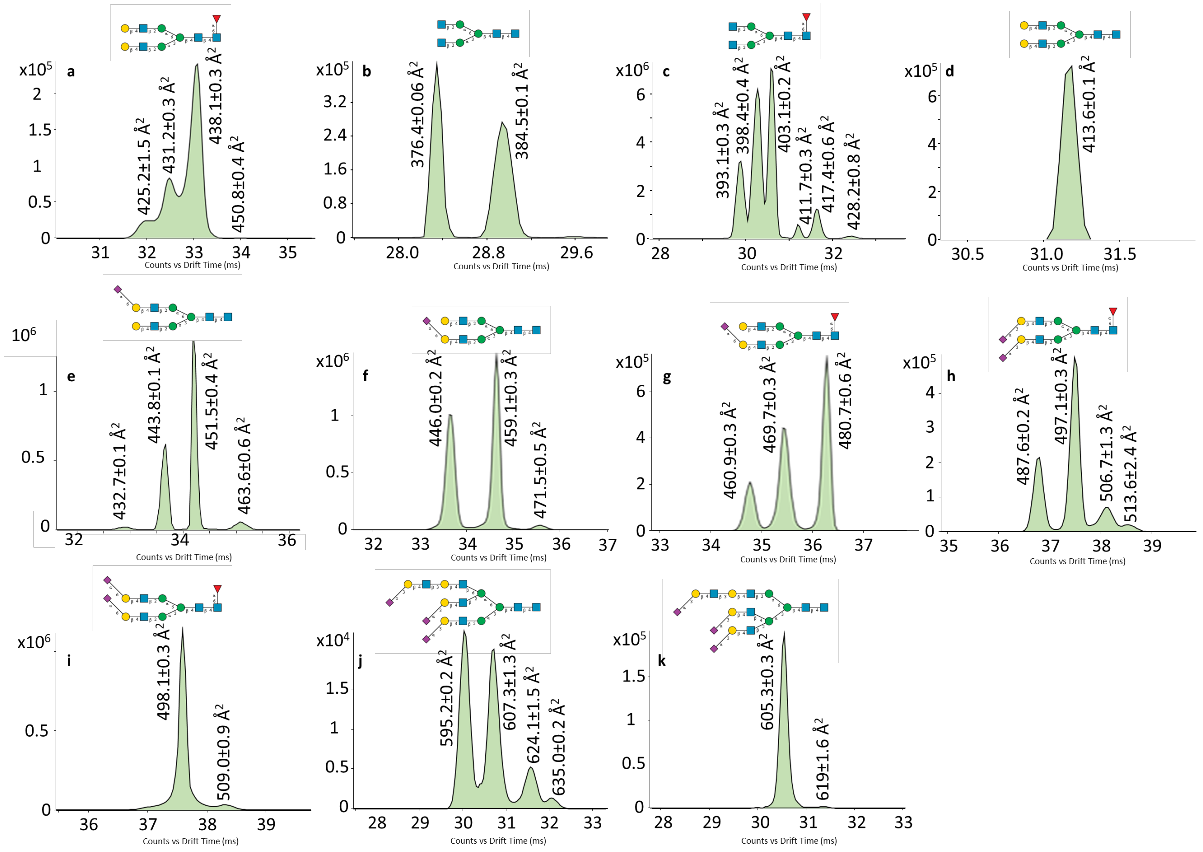

**Supplementary Figure 147. ATDs of synthetic *N*-glycan standards set1 (a-i as $[M-2H]^{2-}$ ions and j-k as $[M-3H]^{3-}$ ions), derivatized with 2AA by reductive amination at the reducing end (n=3 measurements).**

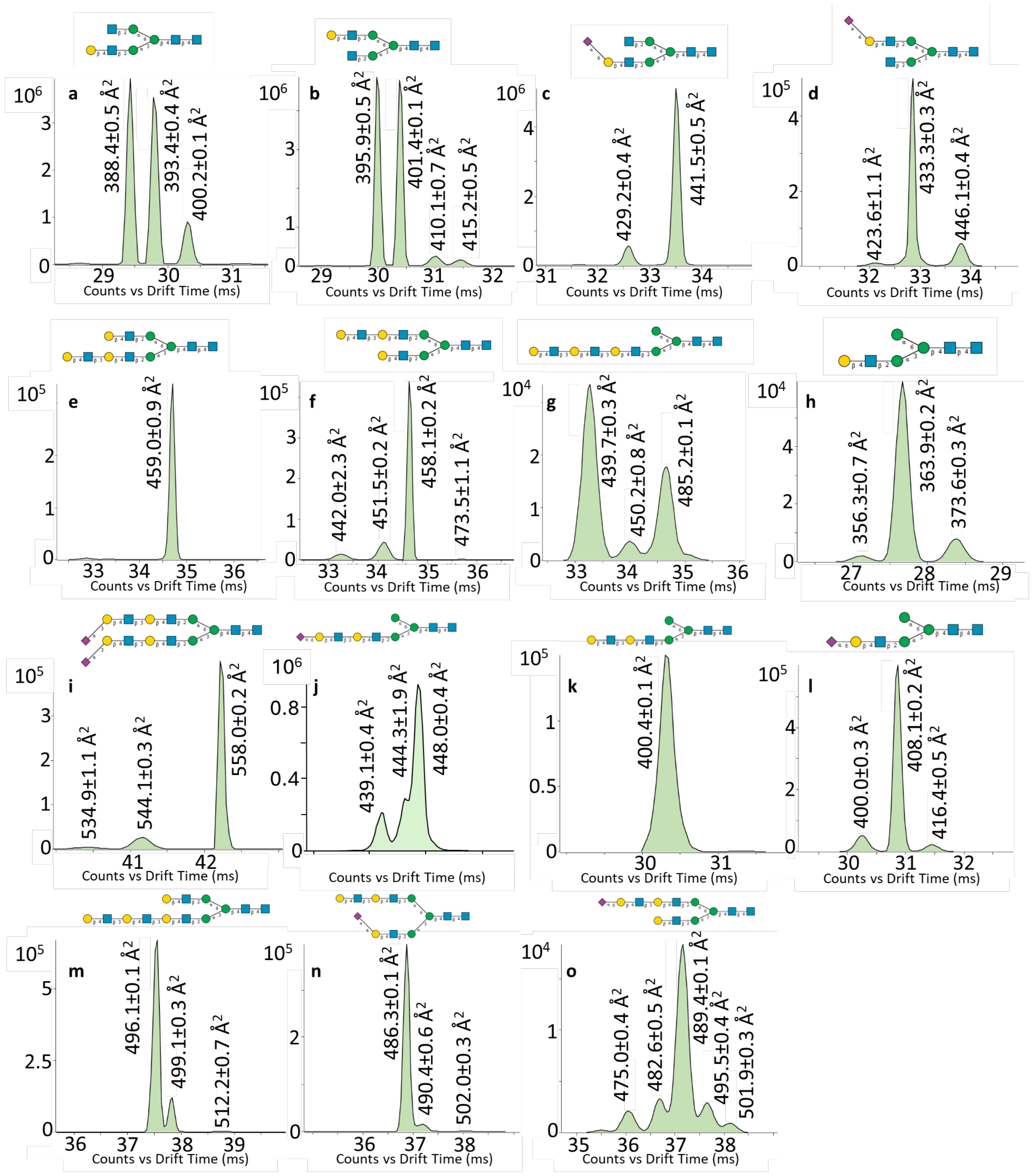

**Supplementary Figure 148. ATDs of synthetic *N*-glycan standards set2 (as $[M\text{-}2H]^{2-}$ ions), derivatized with 2AA by reductive amination at the reducing end (n=3 measurements).**

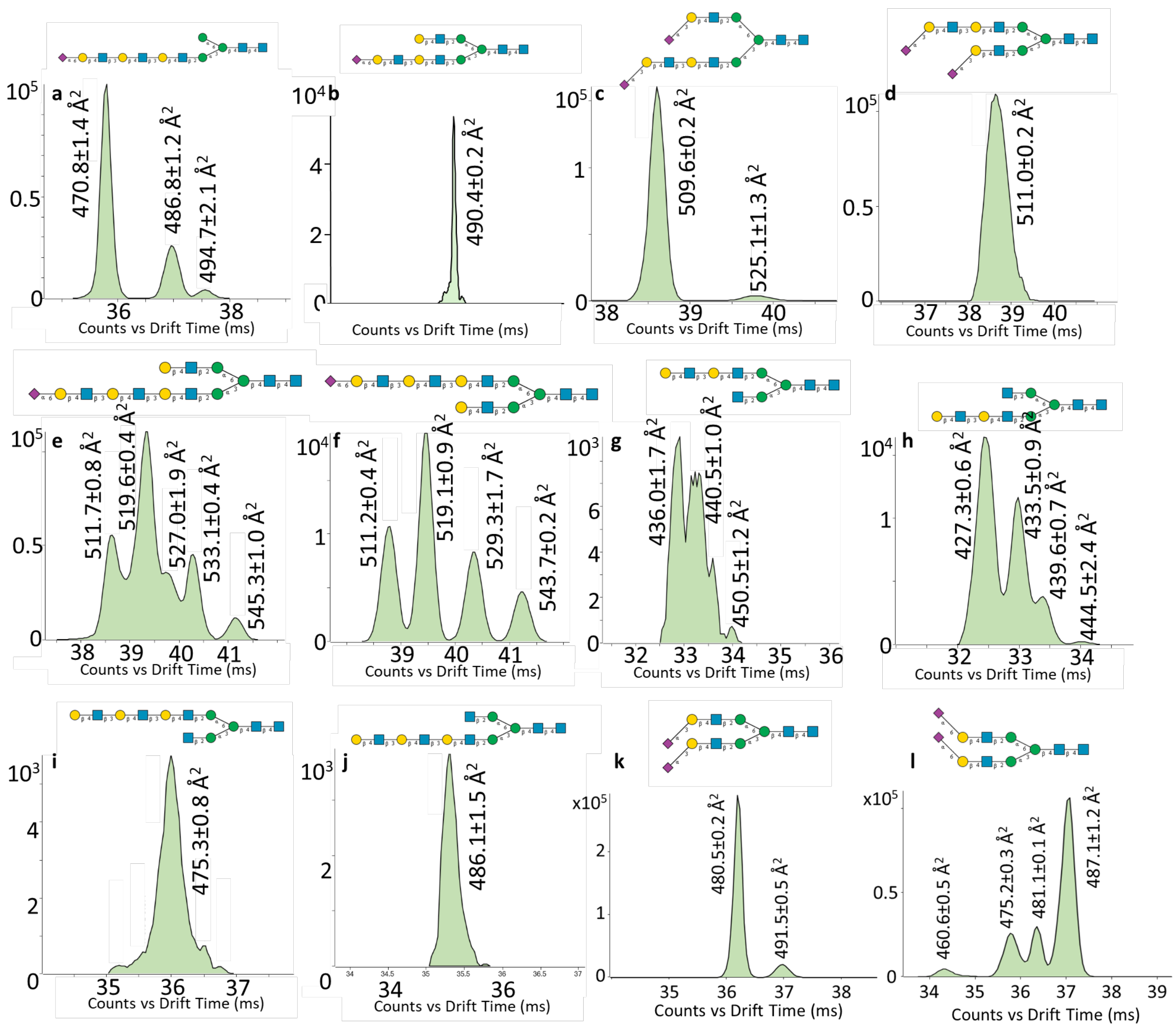

**Supplementary Figure 149. ATDs of synthetic *N*-glycan standards set3 (as $[M\text{-}2H]^{2-}$ ions), derivatized with 2AA by reductive amination at the reducing end (n=3 measurements).**

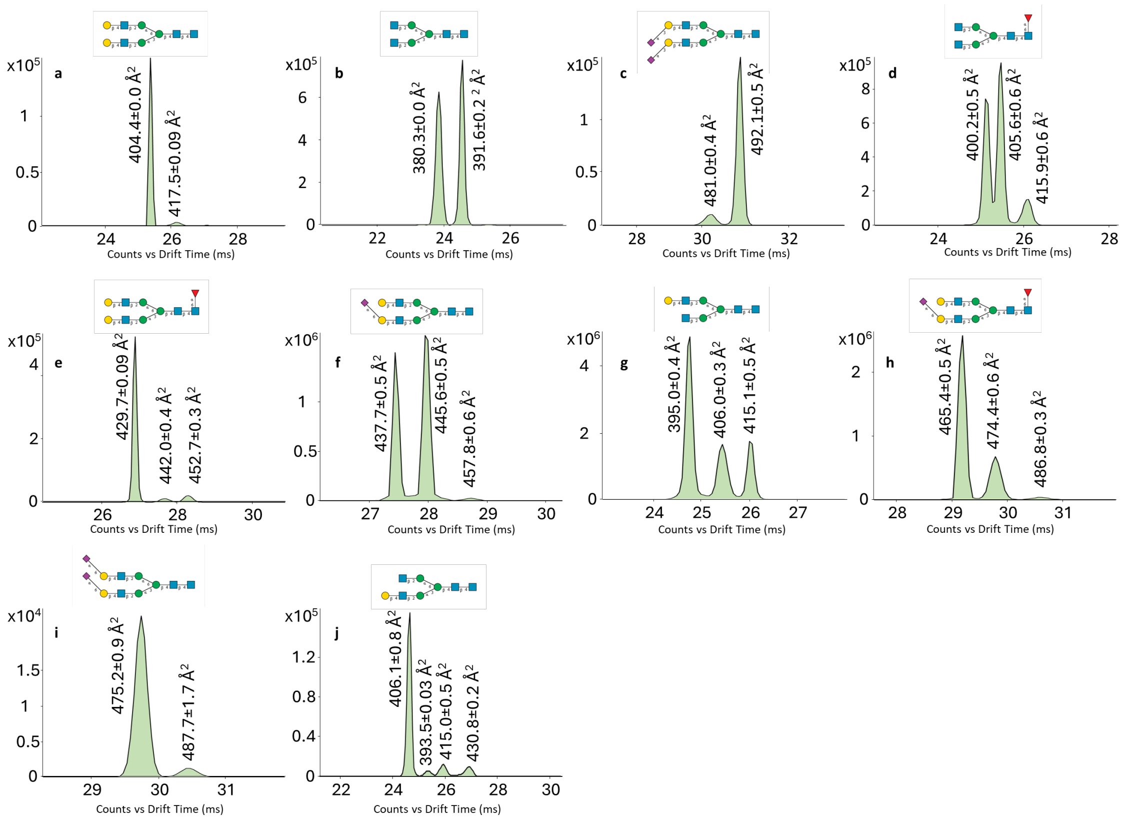

**Supplementary Figure 150. ATDs of synthetic *N*-glycan standards (as $[M+2H]^{2+}$ ions) derivatized with 2AA by reductive amination at the reducing end (n=3 measurements).**

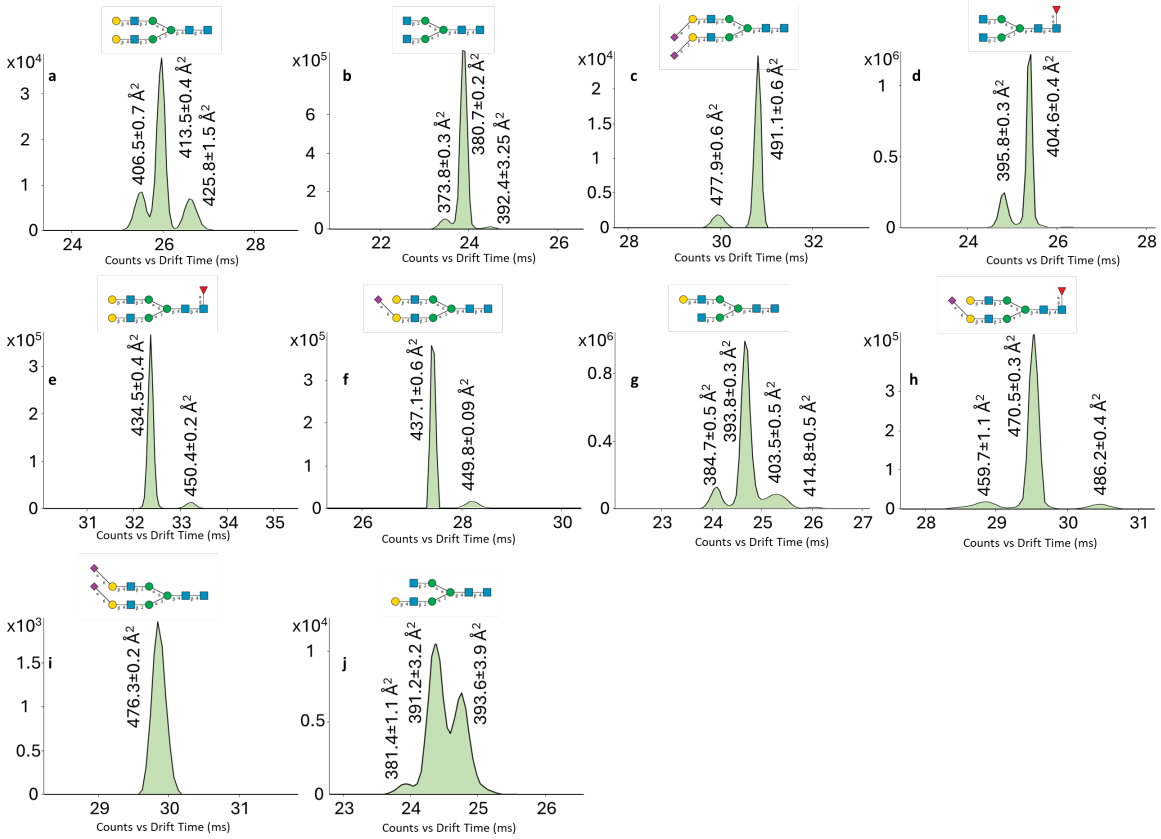

**Supplementary Figure 151. ATDs of synthetic *N*-glycan standards (as [M+H+Na]$^{2+}$ ions) derivatized with 2AA by reductive amination at the reducing end (n=3 measurements).**

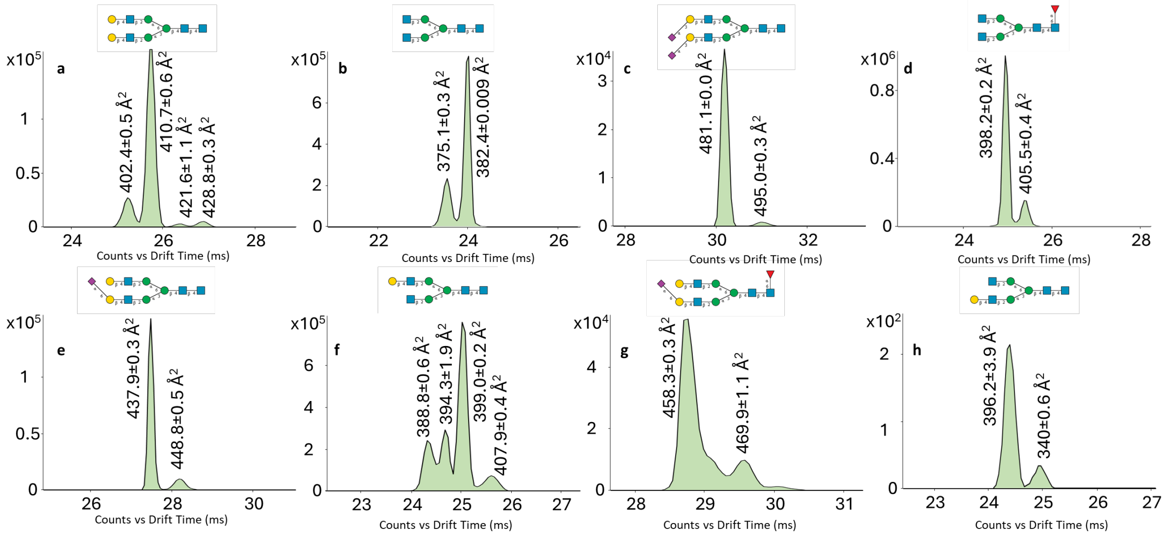

**Supplementary Figure 152. ATDs of *N*-glycan standards (as [M+H+K]$^{2+}$ ions) derivatized with 2AA by reductive amination at the reducing end (n=3 measurements).**

#### 5.2 ATD fingerprints obtained with different ionization sources

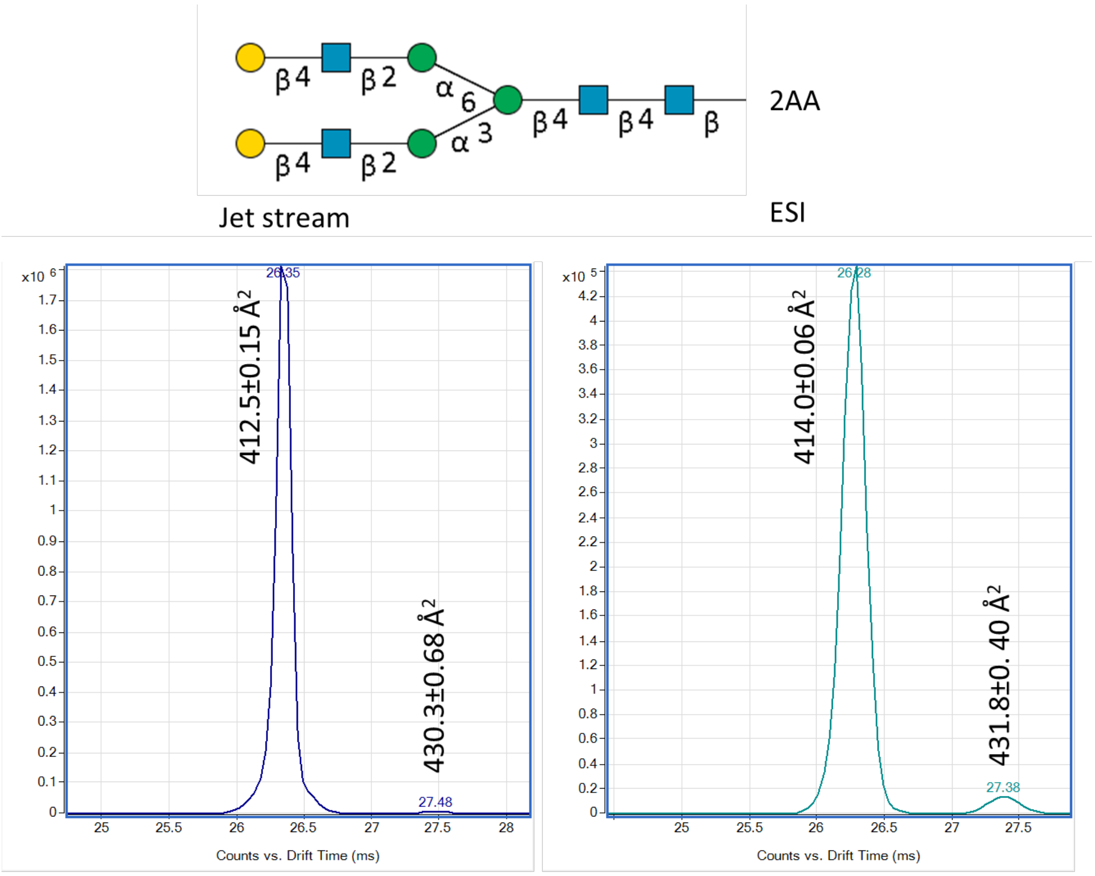

**Supplementary Figure 153. ATDs of synthetic *N*-glycan standards (as $[M\text{-}2H]^{2-}$ ions) derivatized with 2AA by reductive amination at the reducing end with 2AA (n=3 measurements). The ATDs were obtained with either a Jet Stream interface with superheated gas or the standard ESI interface.**

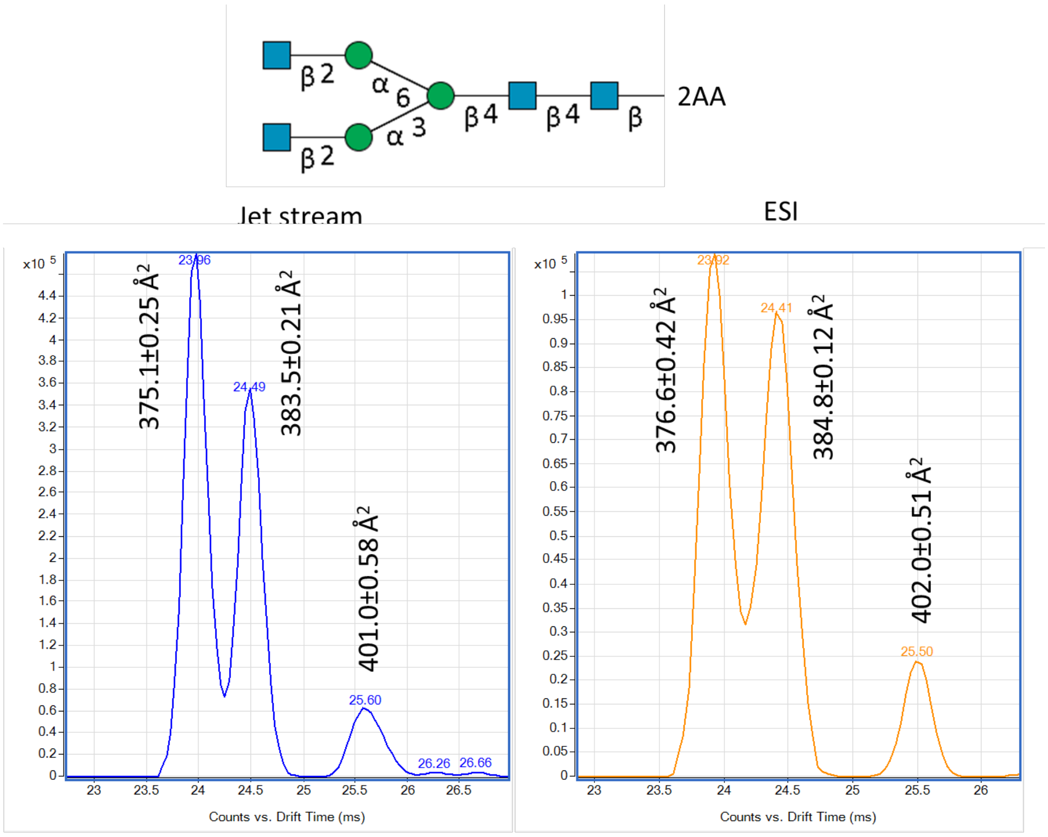

**Supplementary Figure 154. ATDs of synthetic *N*-glycan standards (as [M-2H]$^{2-}$ ions) derivatized with 2AA by reductive amination at the reducing end with 2AA (n=3 measurements). The ATDs were obtained with either a Jet Stream interface with superheated gas or the standard ESI interface.**

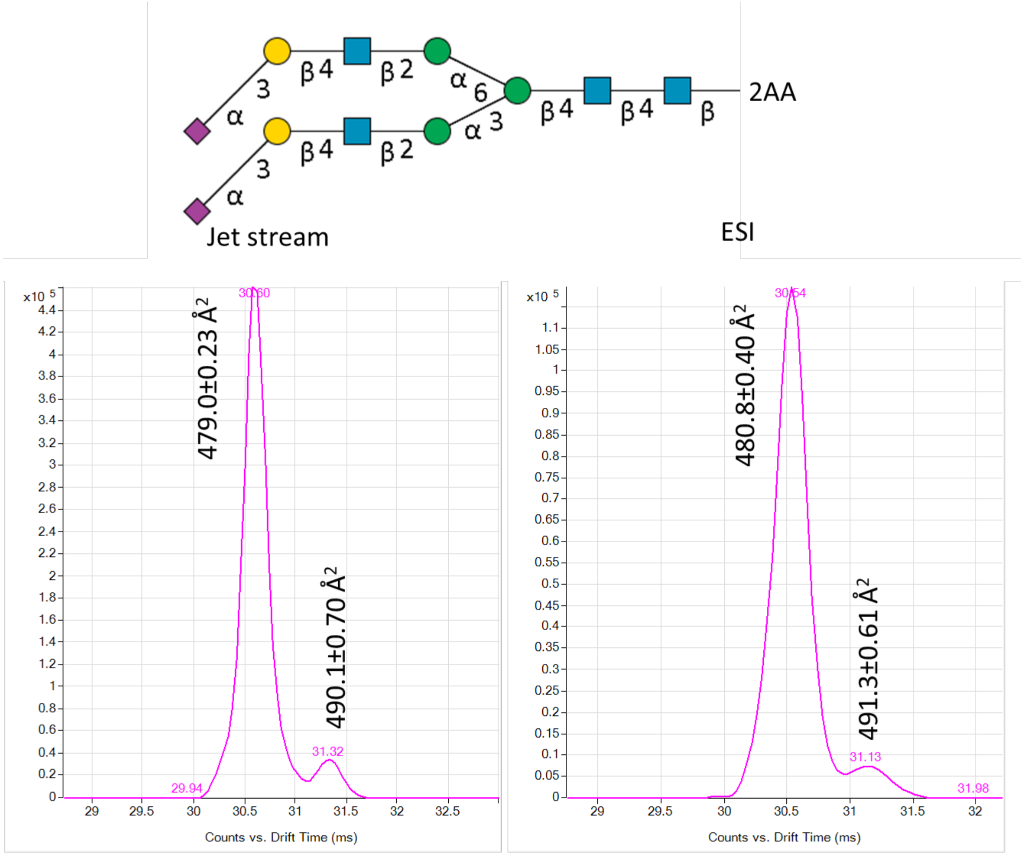

**Supplementary Figure 155. ATDs of synthetic *N*-glycan standards (as $[M-2H]^{2-}$ ions) derivatized with 2AA by reductive amination at the reducing end with 2AA (n=3 measurements). The ATDs were obtained with either a Jet Stream interface with superheated gas or the standard ESI interface.**

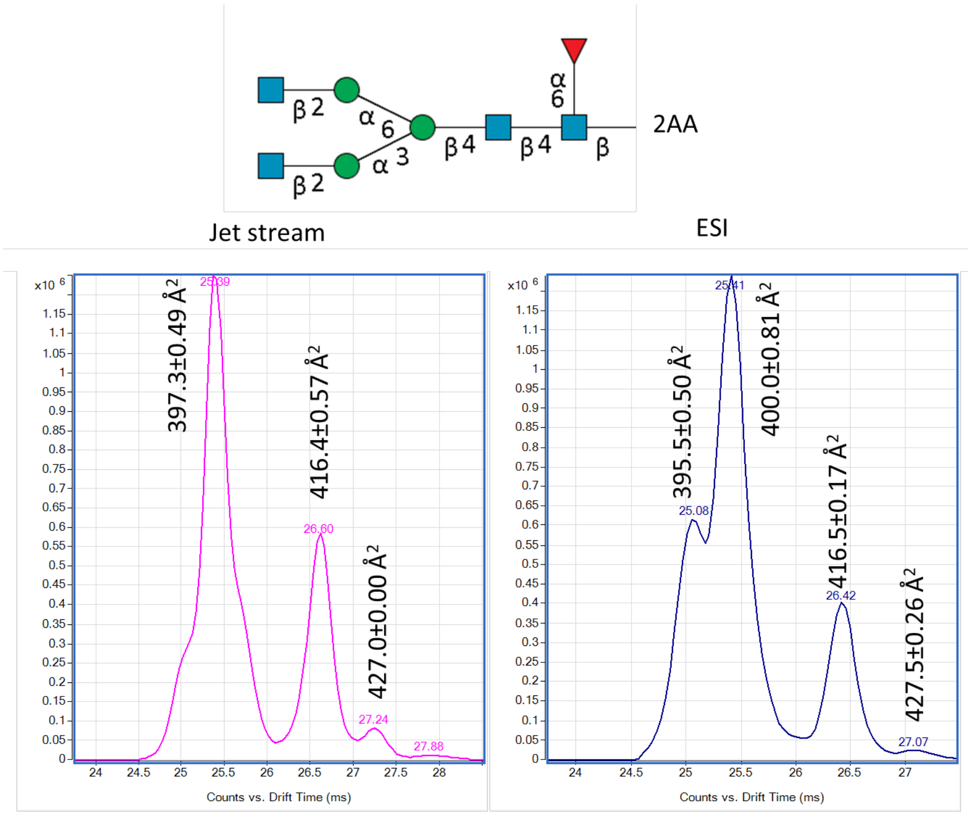

**Supplementary Figure 156. ATDs of synthetic *N*-glycan standards (as [M-2H]$^{2-}$ ions) derivatized with 2AA by reductive amination at the reducing end with 2AA (n=3 measurements). The ATDs were obtained with either a Jet Stream interface with superheated gas or the standard ESI interface.**

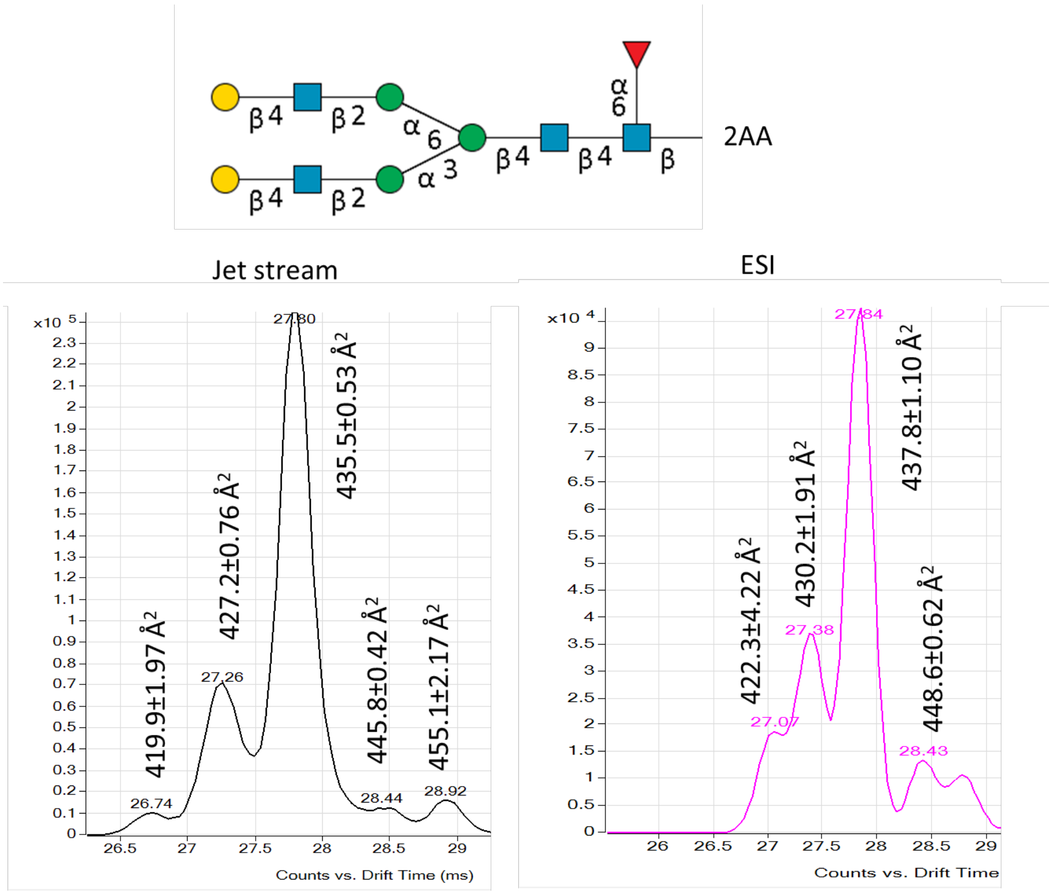

**Supplementary Figure 157. ATDs of synthetic *N*-glycan standards (as [M-2H]$^{2-}$ ions) derivatized with 2AA by reductive amination at the reducing end with 2AA (n=3 measurements). The ATDs were obtained with either a Jet Stream interface with superheated gas or the standard ESI interface.**

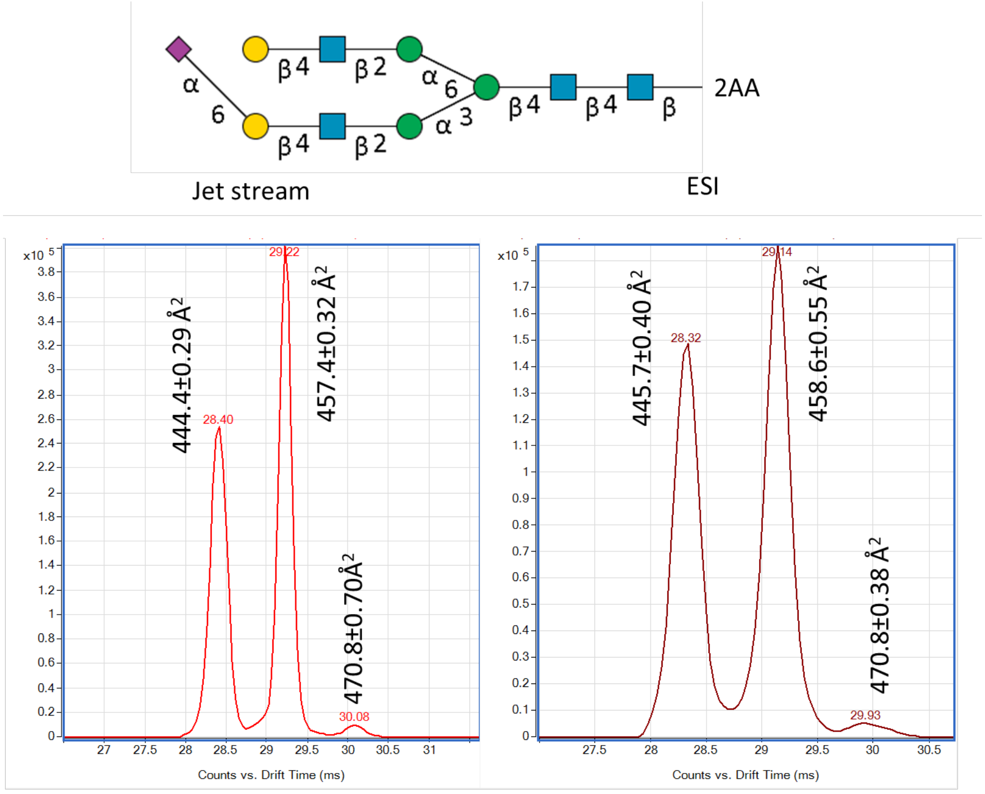

**Supplementary Figure 158. ATDs of synthetic *N*-glycan standards (as [M-2H]$^{2-}$ ions) derivatized with 2AA by reductive amination at the reducing end with 2AA (n=3 measurements). The ATDs were obtained with either a Jet Stream interface with superheated gas or the standard ESI interface.**

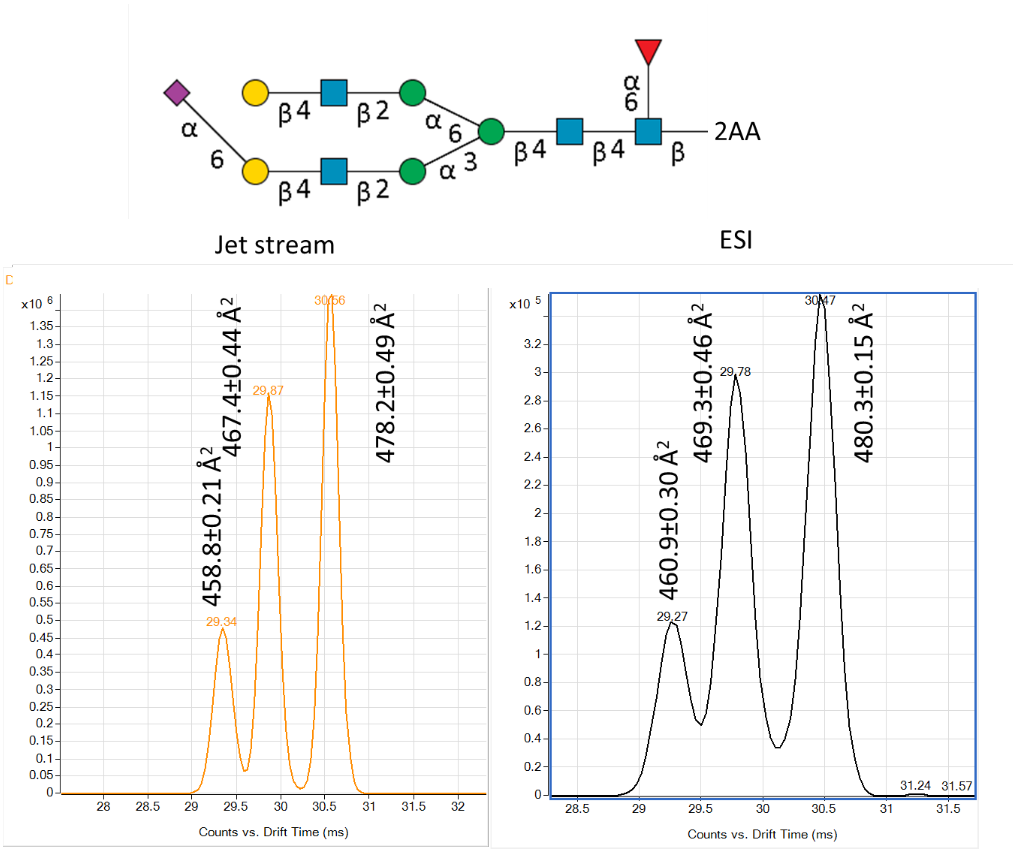

**Supplementary Figure 159. ATDs of synthetic *N*-glycan standards (as $[M\text{-}2H]^{2-}$ ions) derivatized with 2AA by reductive amination at the reducing end with 2AA (n=3 measurements). The ATDs were obtained with either a Jet Stream interface with superheated gas or the standard ESI interface.**

#### 5.3 Disialylated *N*-glycans derived from a biological.

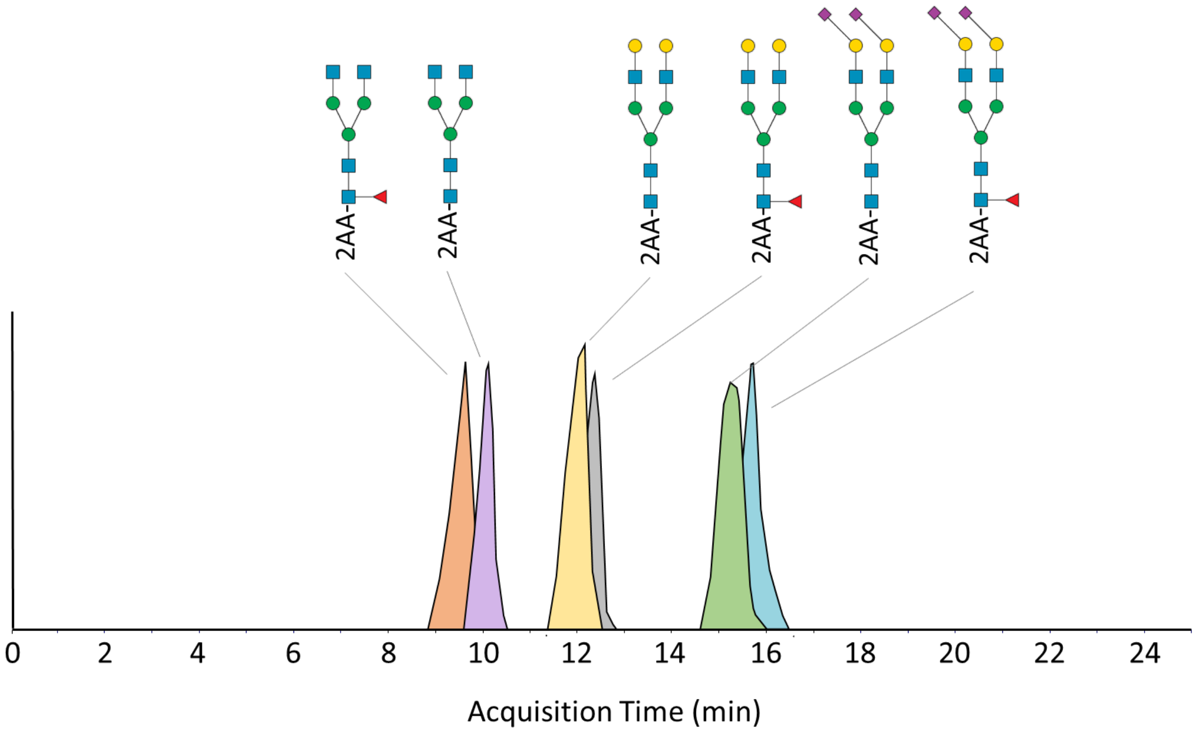

**Supplementary Figure 160. Extracted ion chromatogram of identified *N*-glycans (as $[M\text{-}2H]^{2-}$ ions) derived from the protein Aflibercept (n=1 sample). Glycans were released from proteins by the enzyme PNGaseF, derivatized with 2AA by reductive amination at the reducing end and analyzed with HILIC-IM-MS.**

### 6. Fragmentation spectrum of *N*-glycan standard 60

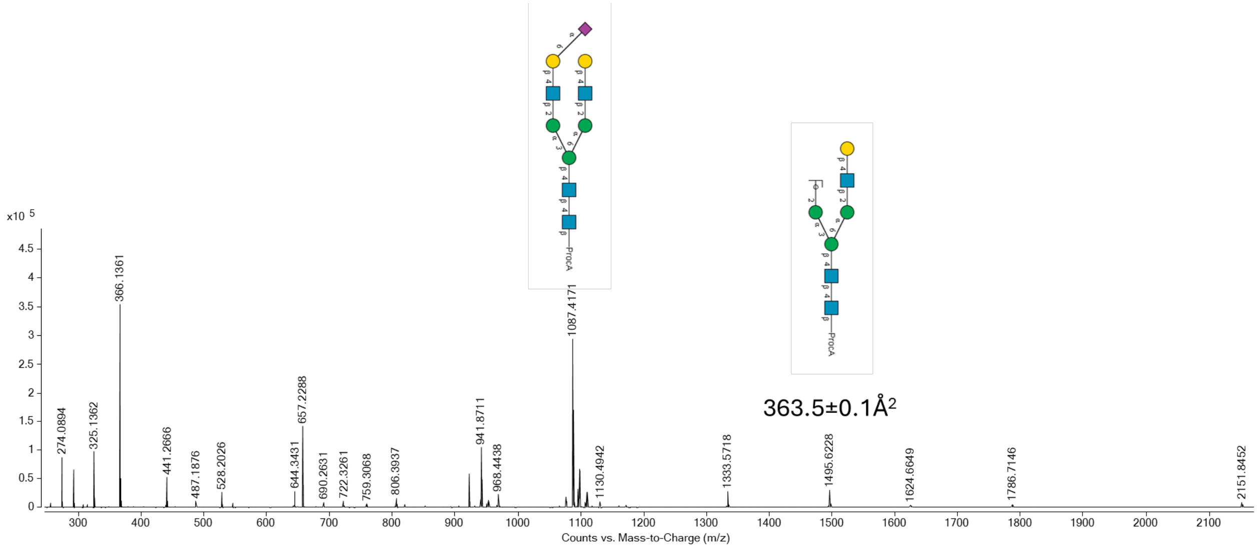

**Supplementary Figure 161. MS fragmentation spectrum of *N*-glycan standard 60.**

### 7. Analysis results of *N*-glycans derived from proteins.

#### 7.1 Chromatogram with *N*-glycans derived from Aflibercept.

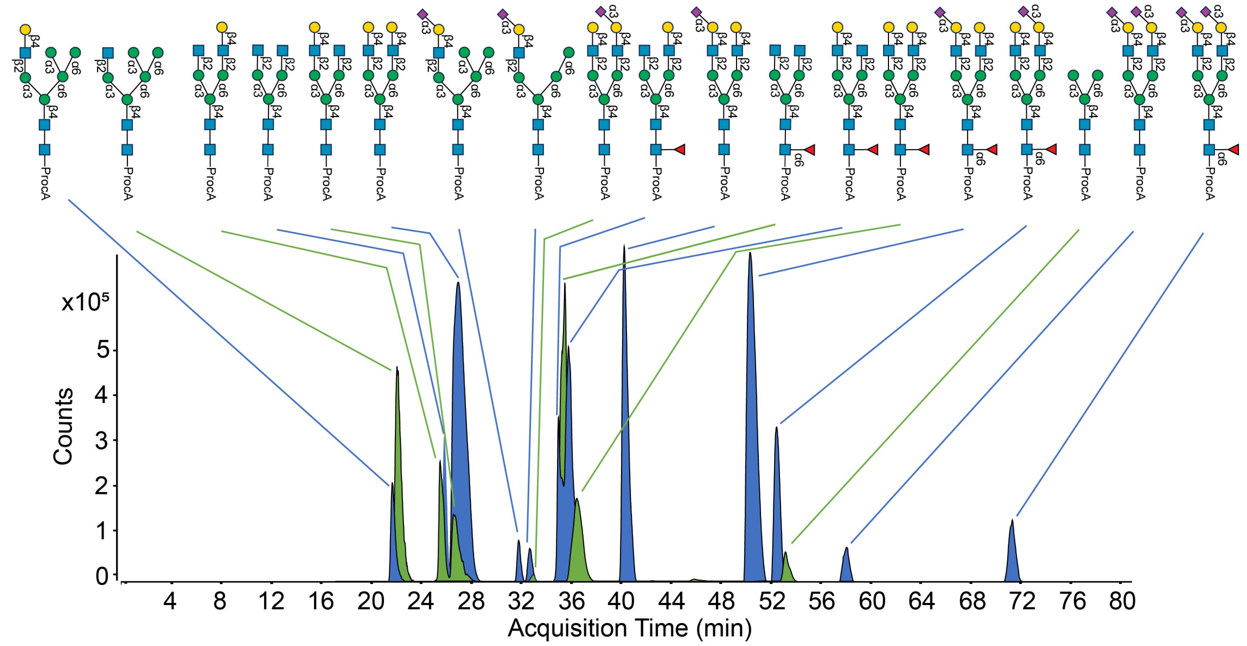

**Supplementary Figure 162 . Extracted ion chromatogram (from [M+2H]$^{2+}$ ions) obtained during PGC LC-IM-MS analysis of sialylated *N*-glycans derived from the protein aflibercept.**

#### 7.2 MS spectra and CCS values of *N*-glycans derived from Aflibercept.

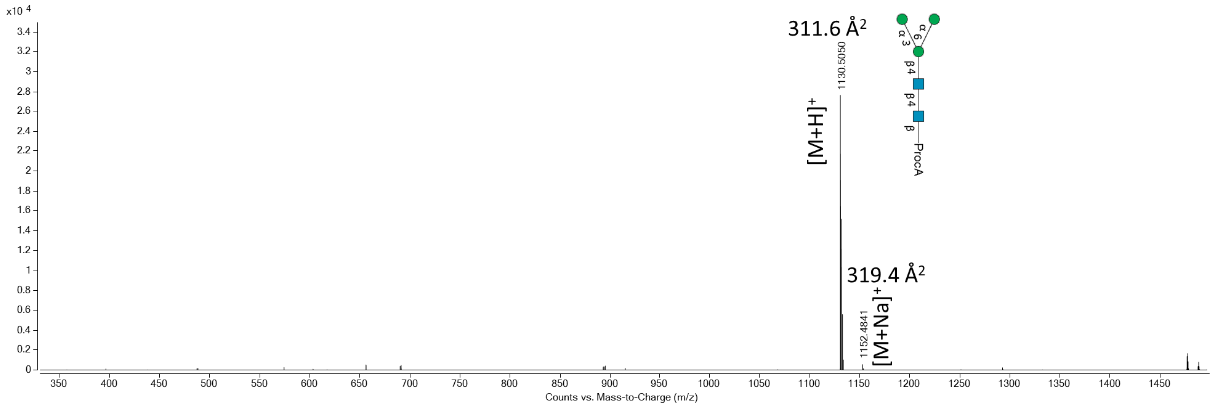

**Supplementary Figure 163. MS spectrum of an *N*-glycan derived from the biological Aflibercept.**

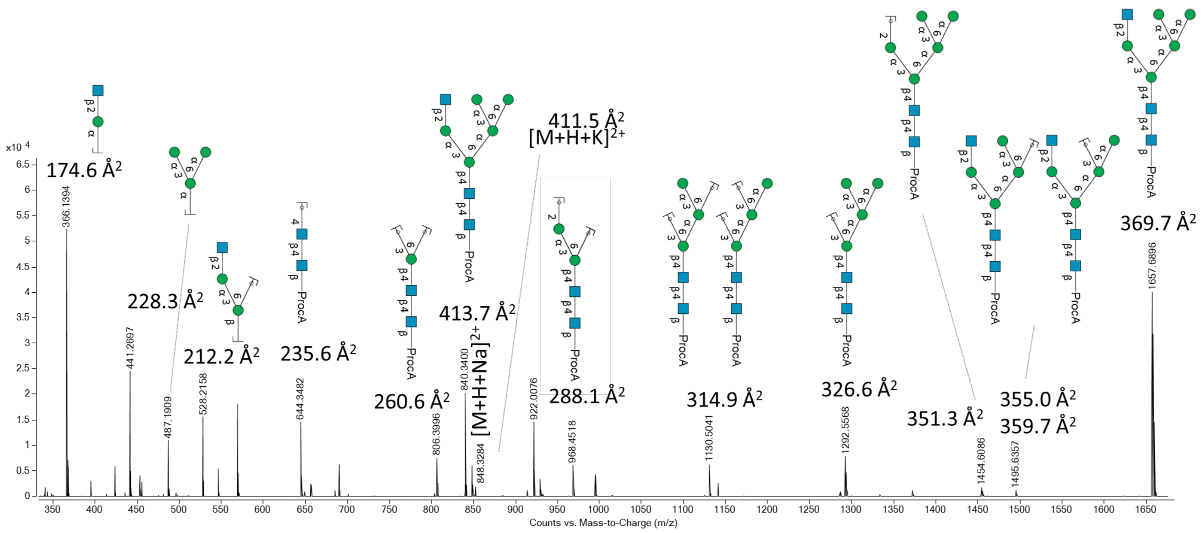

**Supplementary Figure 164. MS spectrum of an *N*-glycan derived from the biological Aflibercept.**

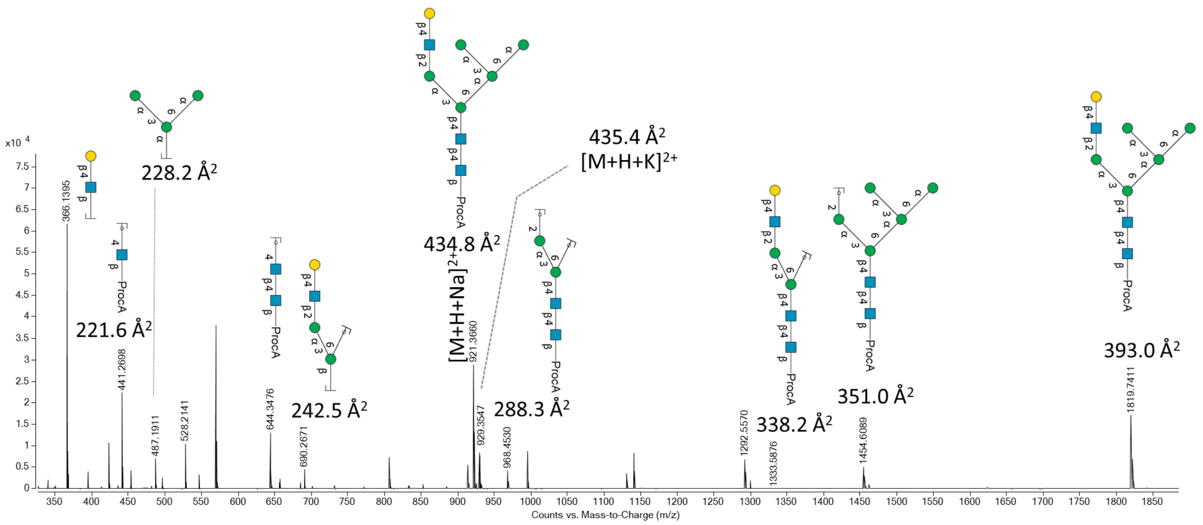

**Supplementary Figure 165. MS spectrum of an *N*-glycan derived from the biological Aflibercept.**

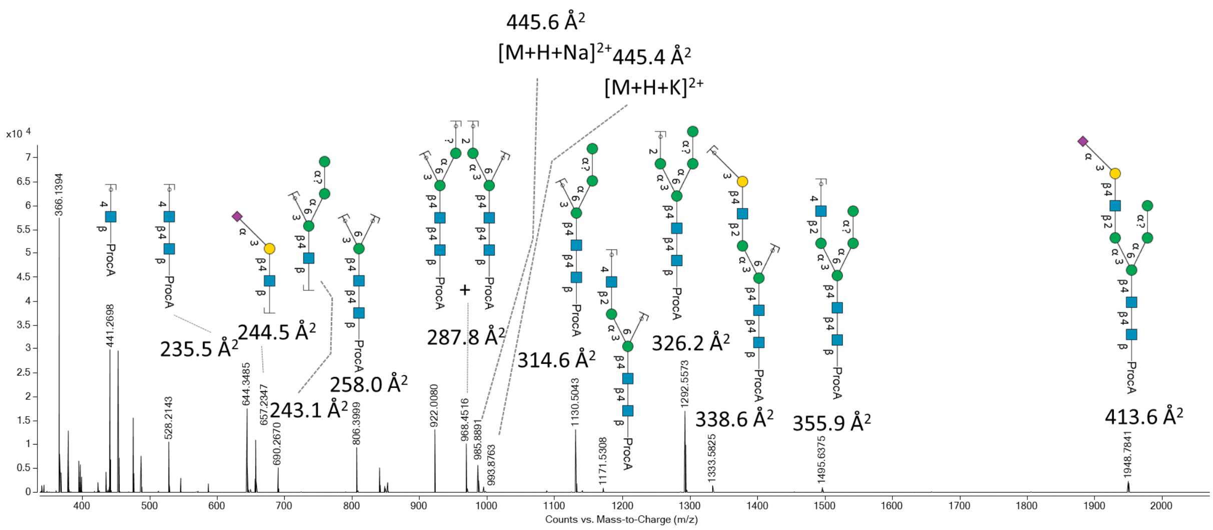

**Supplementary Figure 166. MS spectrum of an *N*-glycan derived from the biological Aflibercept.**

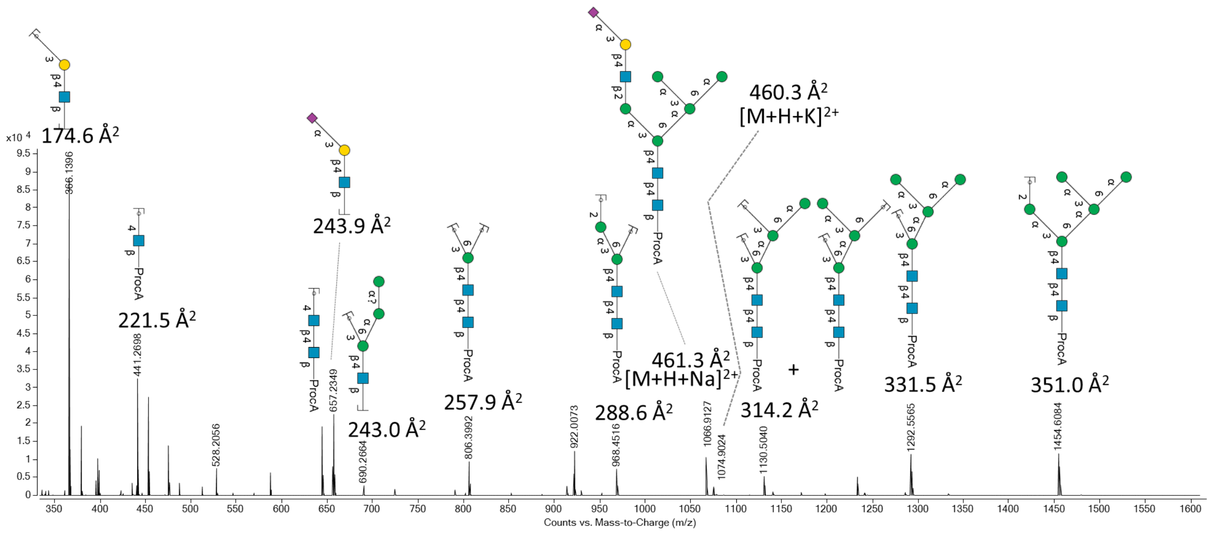

**Supplementary Figure 167. MS spectrum of an *N*-glycan derived from the biological Aflibercept.**

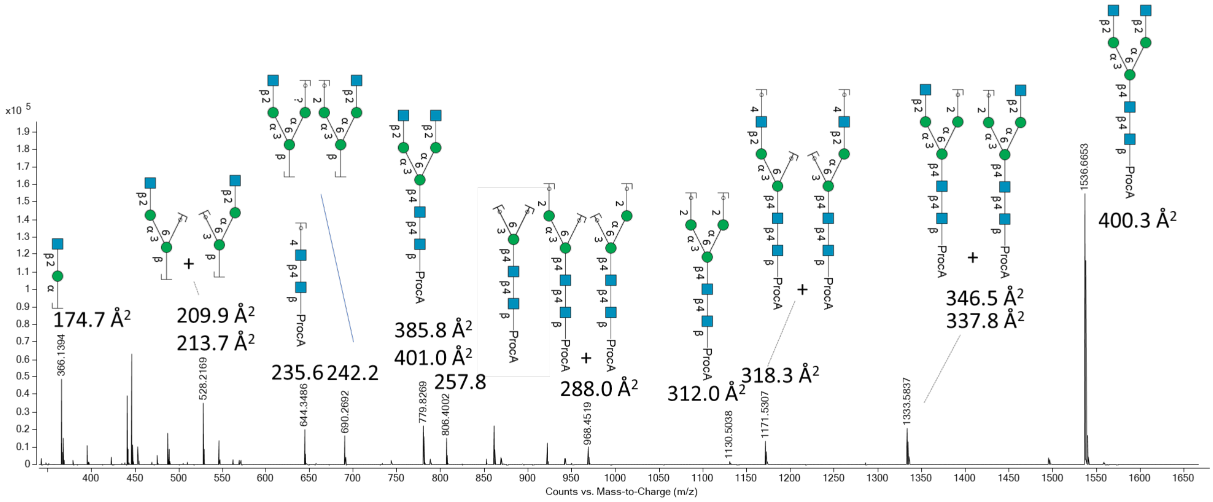

**Supplementary Figure 168. MS spectrum of an *N*-glycan derived from the biological Aflibercept.**

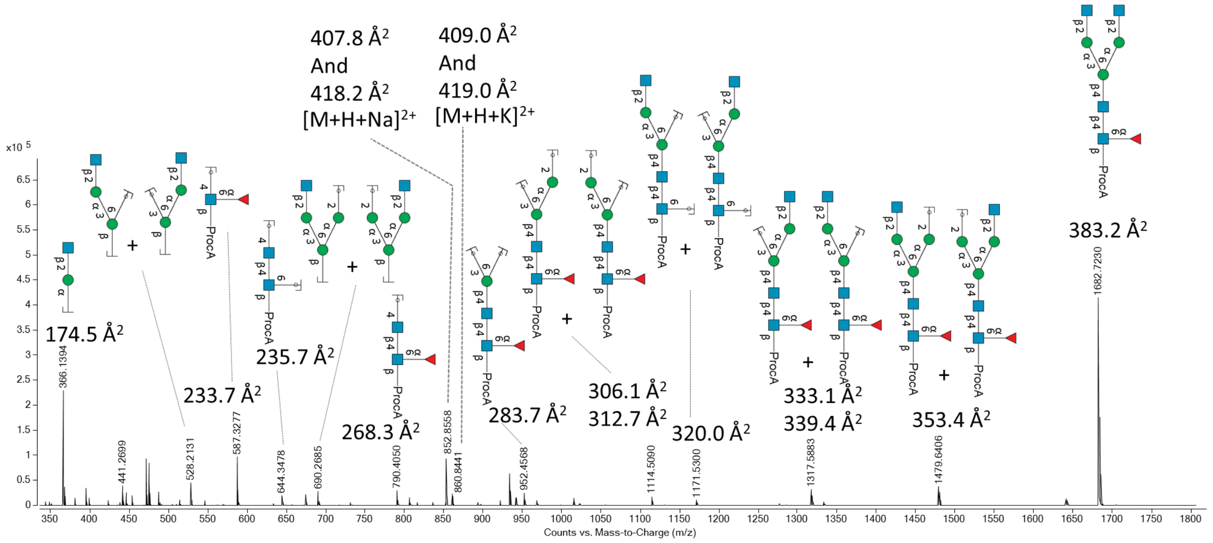

**Supplementary Figure 169. MS spectrum of an *N*-glycan derived from the biological Aflibercept.**

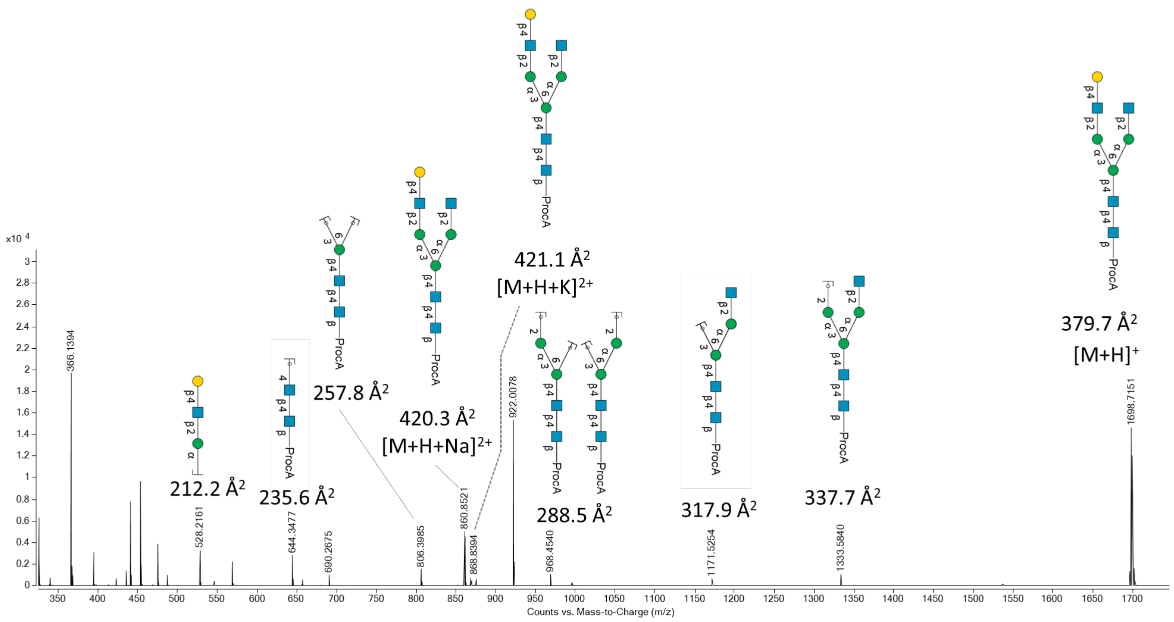

**Supplementary Figure 170. MS spectrum of an *N*-glycan derived from the biological Aflibercept.**

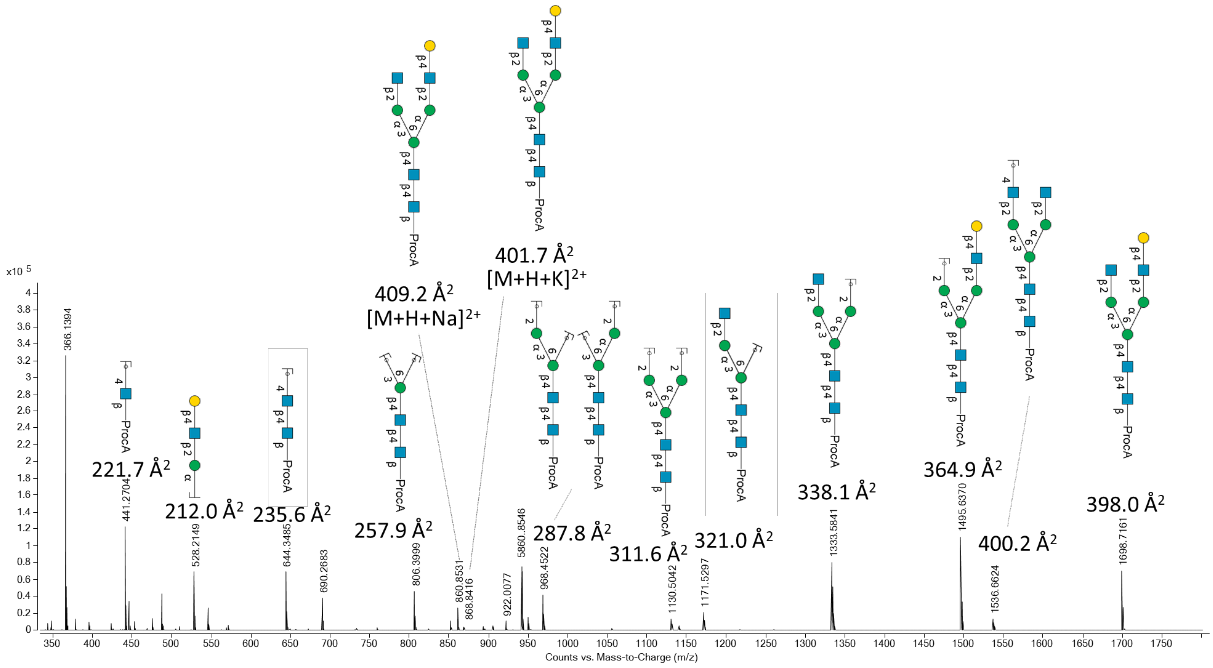

**Supplementary Figure 171. MS spectrum of an *N*-glycan derived from the biological Aflibercept.**

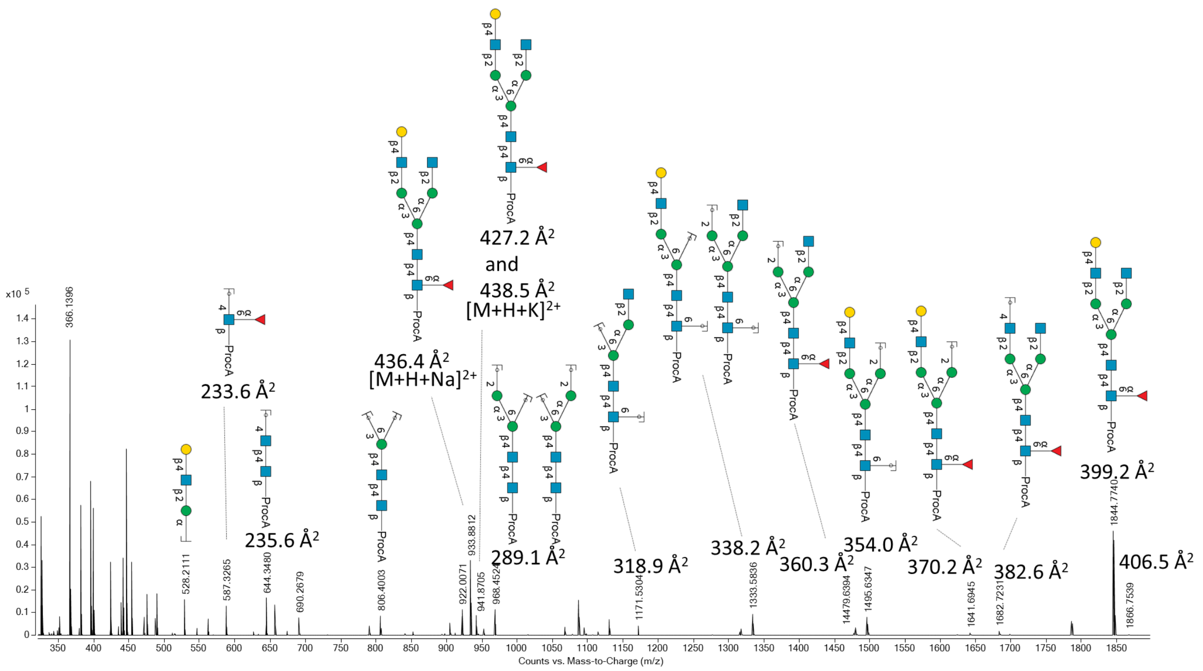

**Supplementary Figure 172. MS spectrum of an *N*-glycan derived from the biological Aflibercept.**

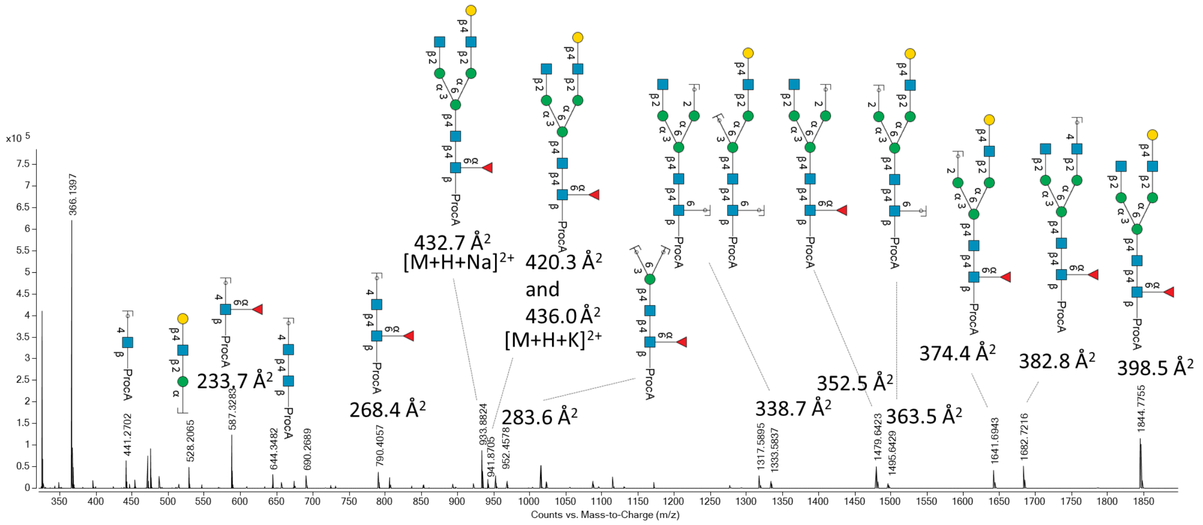

**Supplementary Figure 173. MS spectrum of an *N*-glycan derived from the biological Aflibercept.**

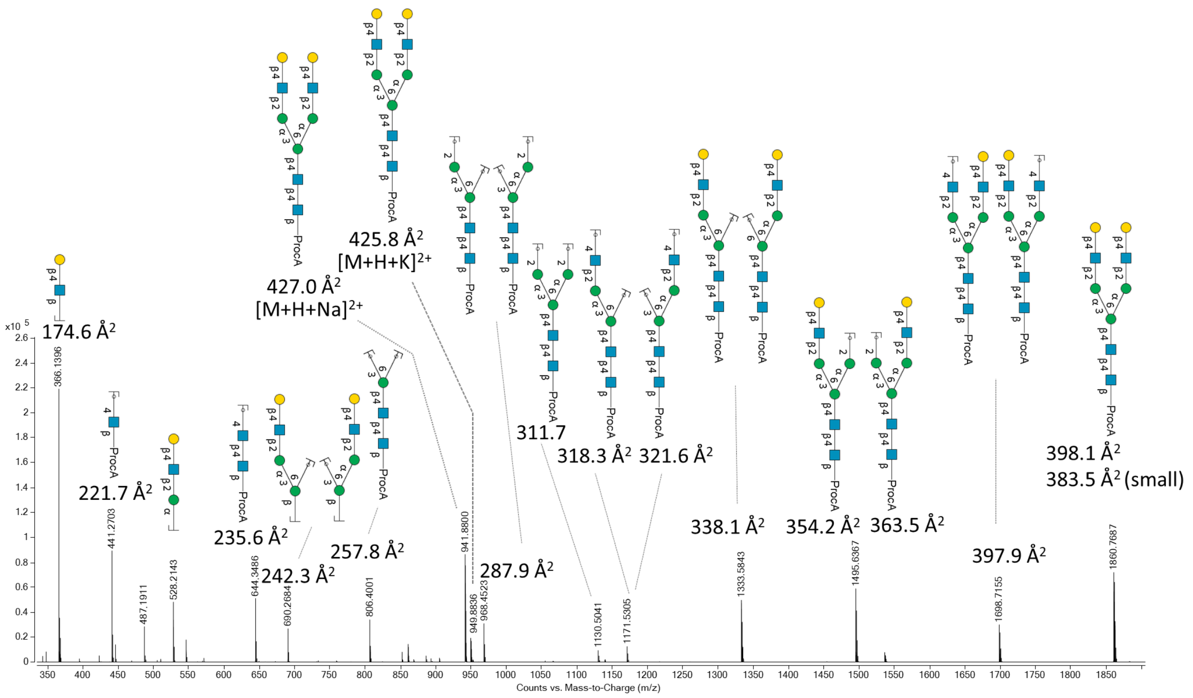

**Supplementary Figure 174. MS spectrum of an *N*-glycan derived from the biological Aflibercept.**

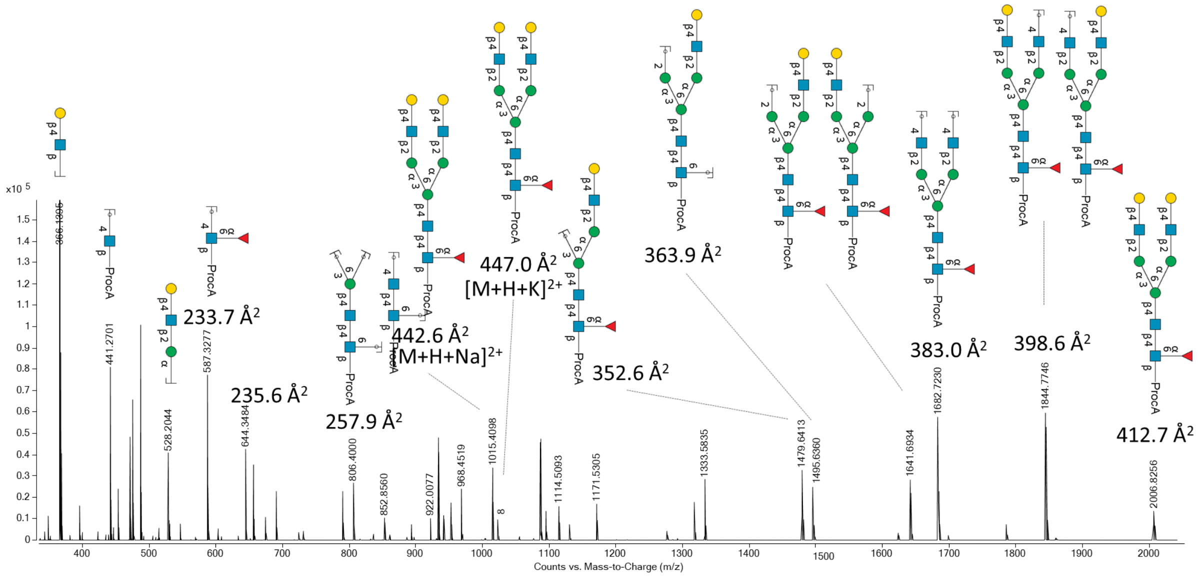

**Supplementary Figure 175. MS spectrum of an *N*-glycan derived from the biological Aflibercept.**

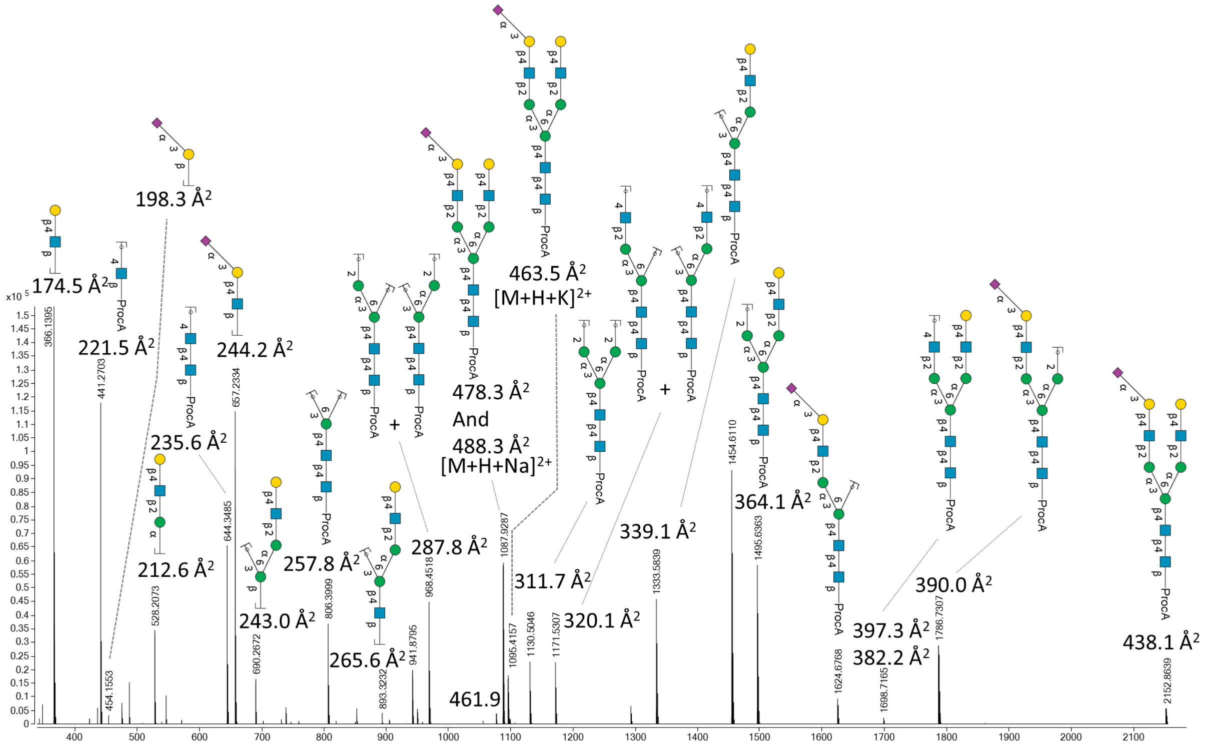

Supplementary Figure 176. MS spectrum of an *N*-glycan derived from the biological Aflibercept.

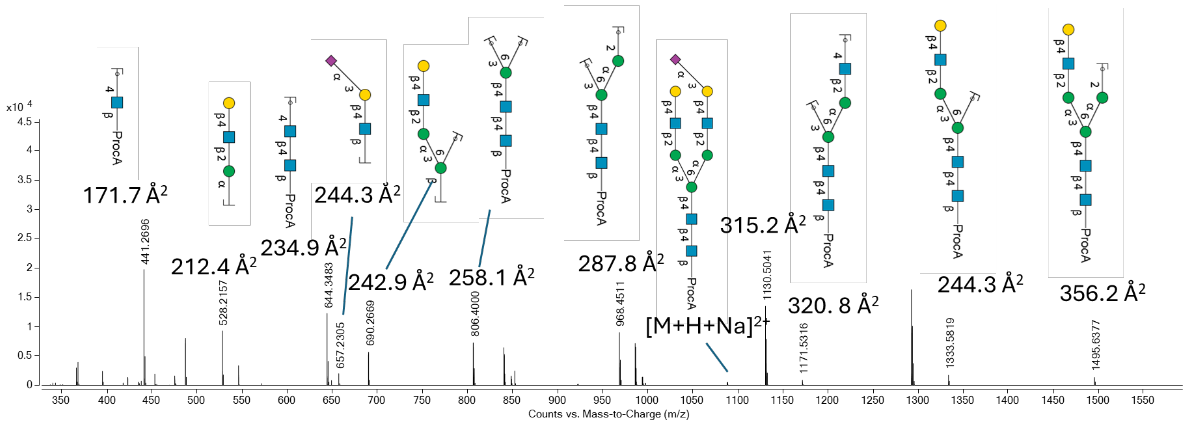

Supplementary Figure 177. MS spectrum of an *N*-glycan derived from the biological Aflibercept.

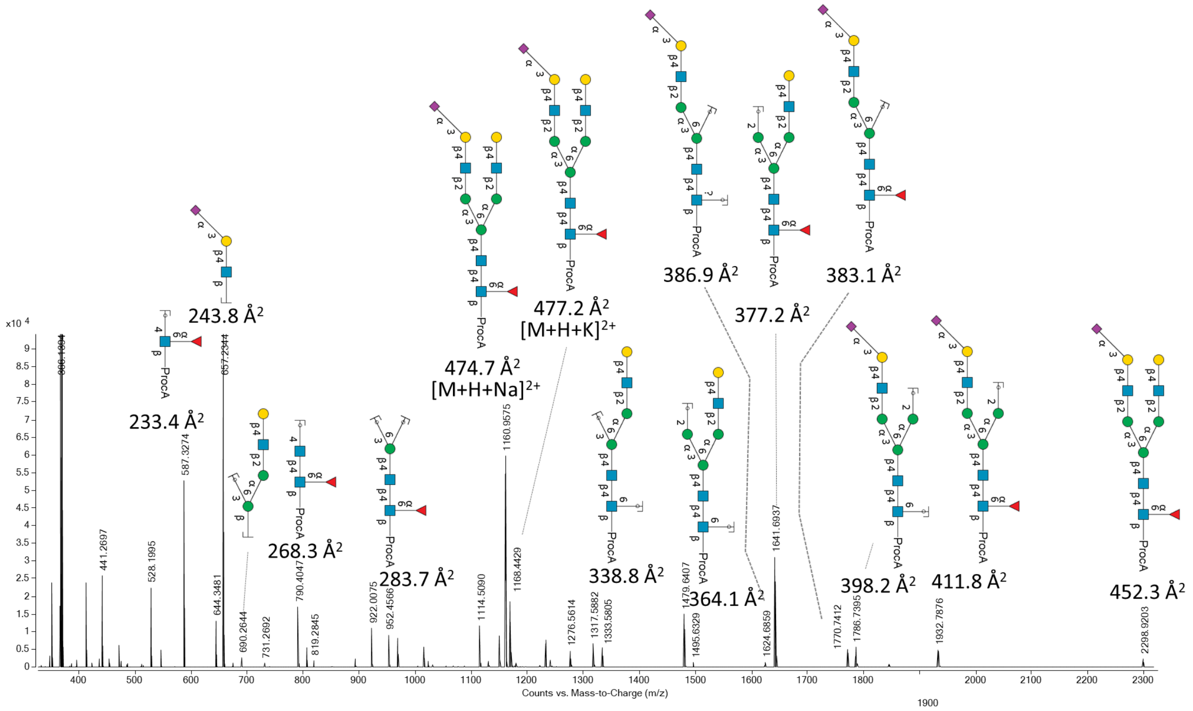

**Supplementary Figure 178. MS spectrum of an *N*-glycan derived from the biological Aflibercept.**

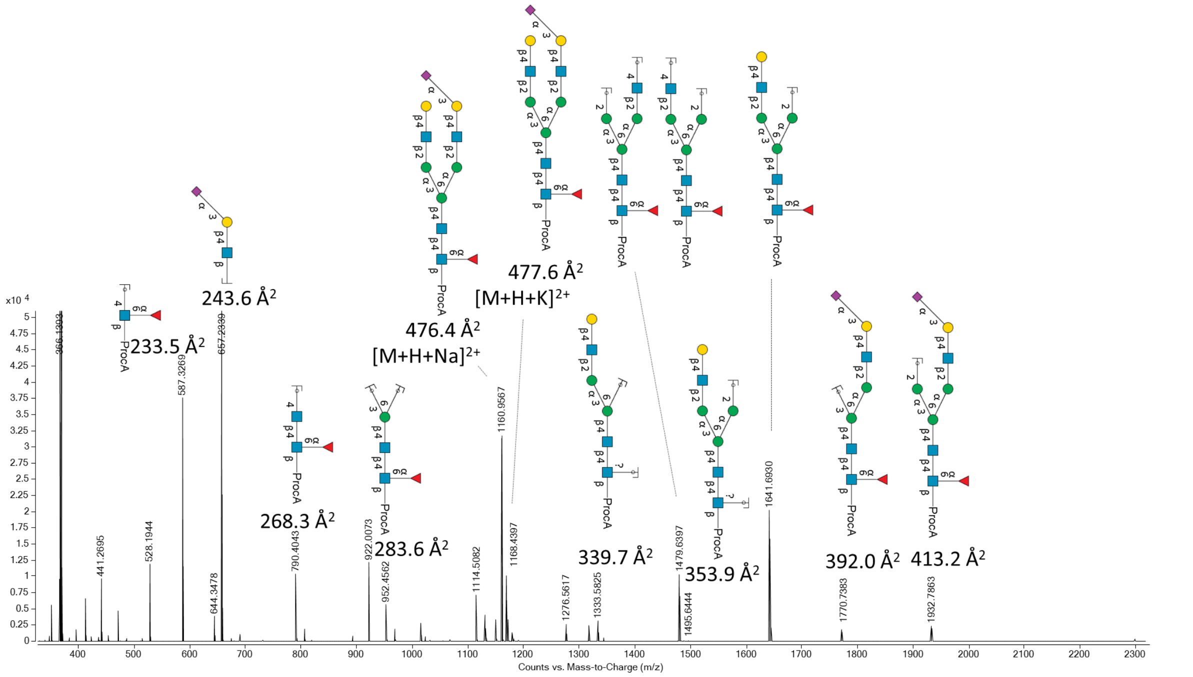

**Supplementary Figure 179. MS spectrum of an *N*-glycan derived from the biological Aflibercept.**

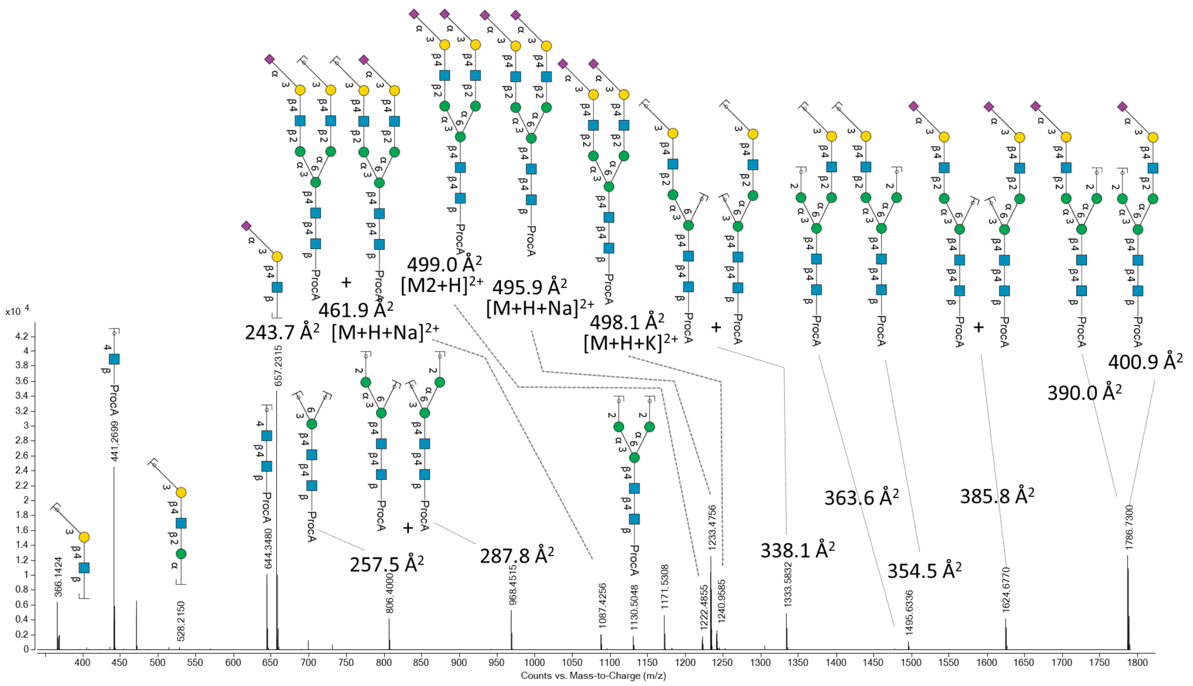

**Supplementary Figure 180. MS spectrum of an *N*-glycan derived from the biological Aflibercept.**

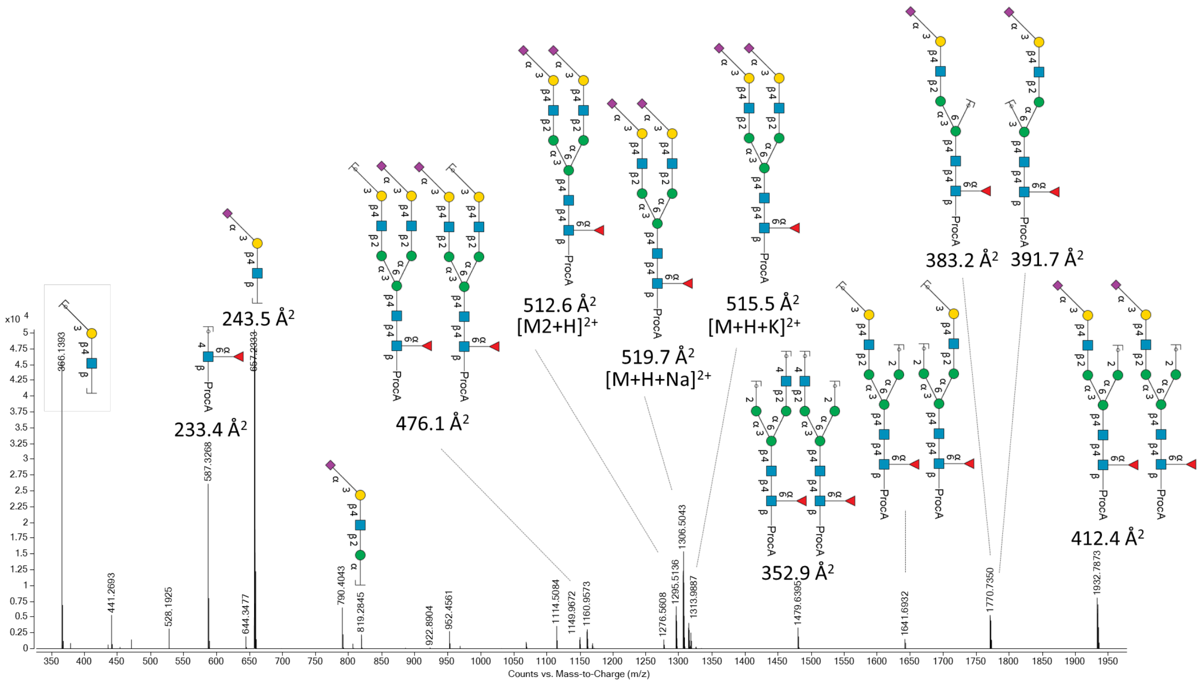

**Supplementary Figure 181. MS spectrum of an *N*-glycan derived from the biological Aflibercept.**

#### 7.3 Chromatogram with *N*-glycans derived from transferrin.

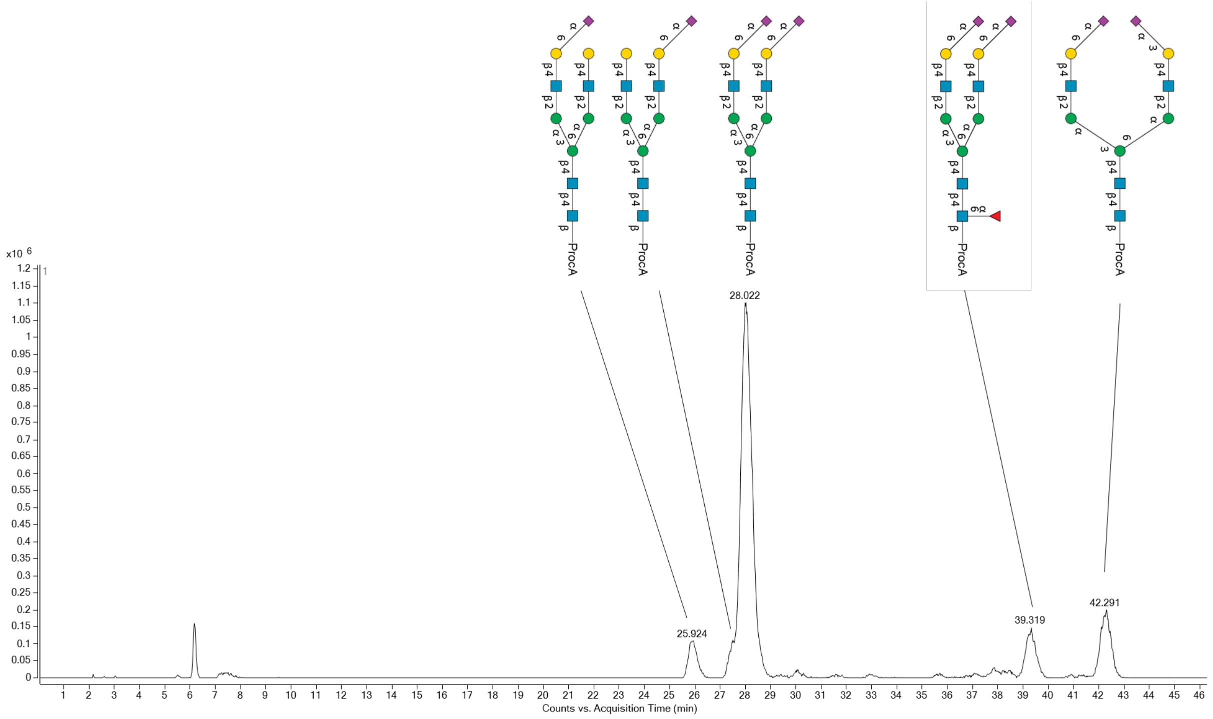

**Supplementary Figure 182 . Extracted ion chromatogram (from oxonium subunit fragment ions at *m/z* 657-659) obtained during PGC LC-IM-MS analysis of sialylated *N*-glycans derived from the protein transferrin.**

#### 7.4 Mass spectra and CCS values of *N*-glycans derived from transferrin.

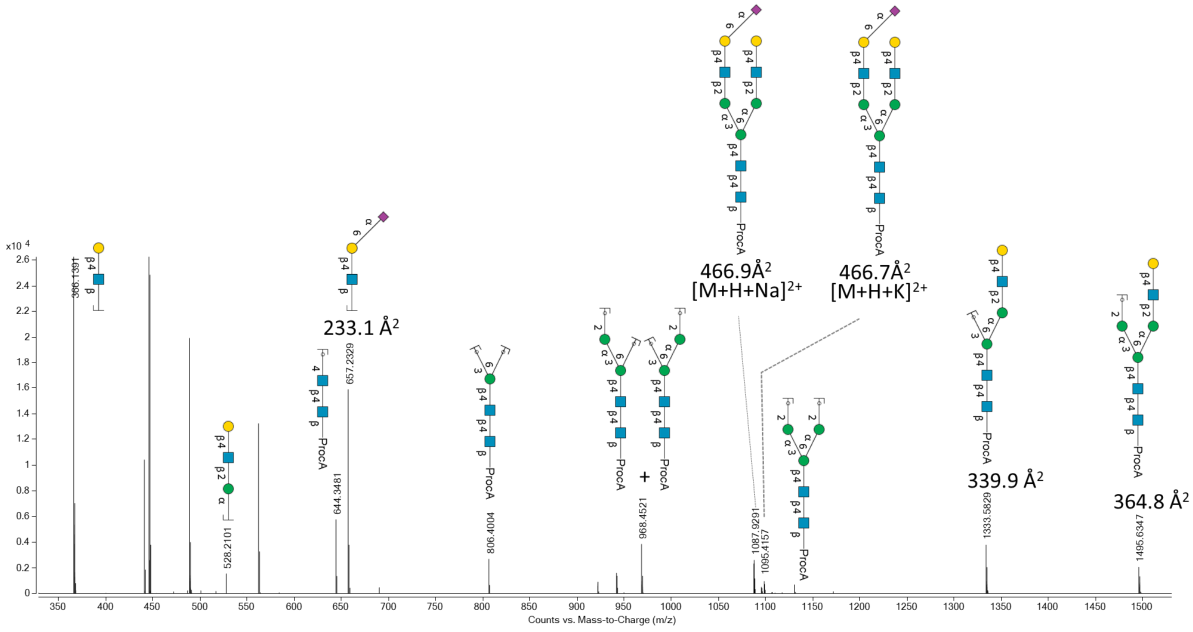

**Supplementary Figure 183. Mass fragmentation spectrum of an *N*-glycan derived from the protein transferrin.**

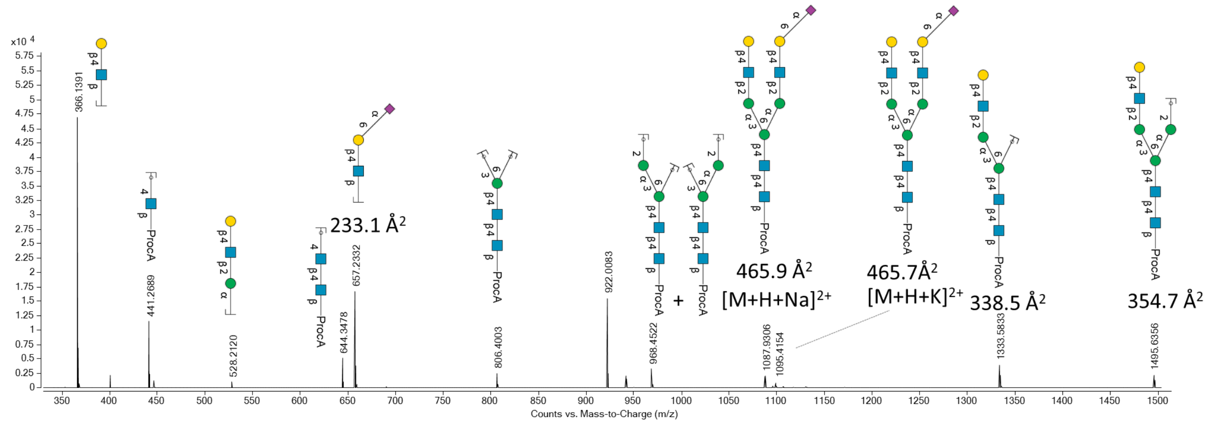

**Supplementary Figure 184. Mass fragmentation spectrum of an *N*-glycan derived from the protein transferrin.**

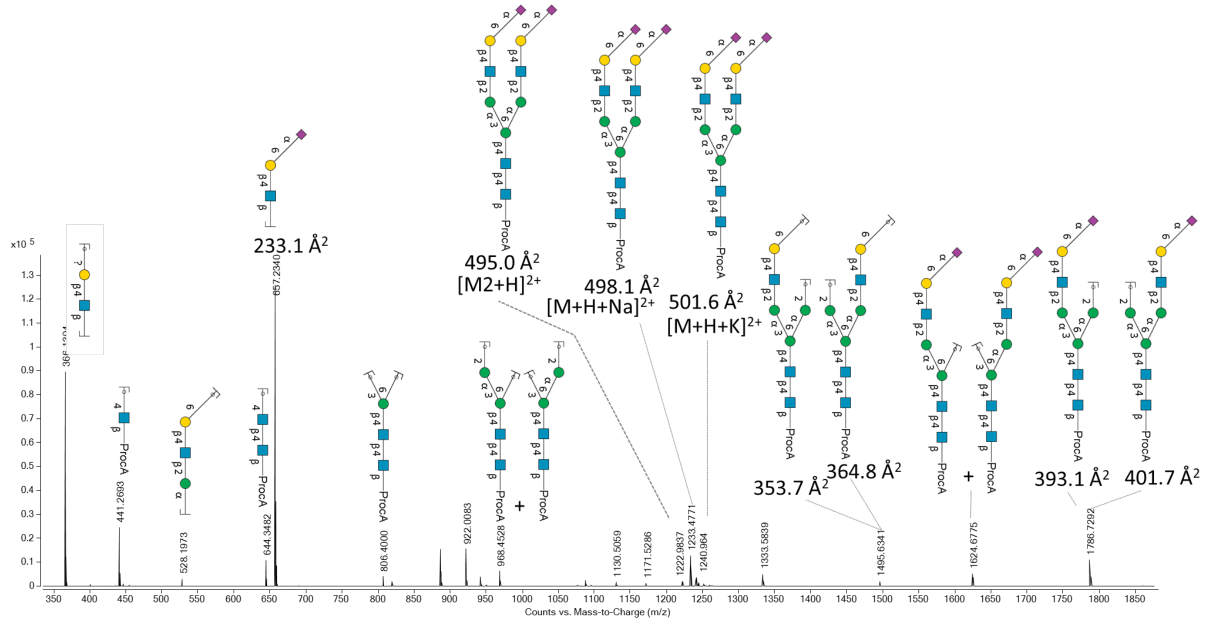

**Supplementary Figure 185. Mass fragmentation spectrum of an *N*-glycan derived from the protein transferrin.**

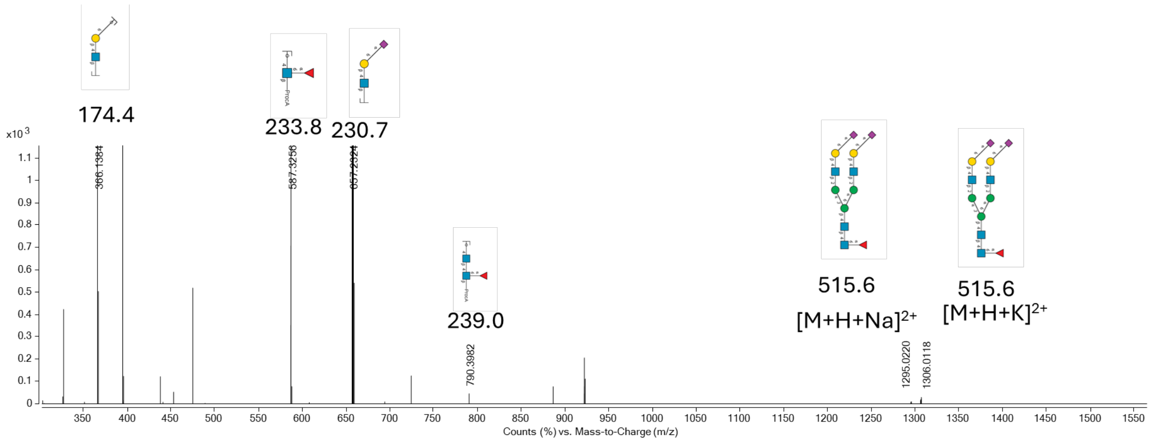

**Supplementary Figure 186. Mass fragmentation spectrum of an *N*-glycan derived from the protein transferrin.**

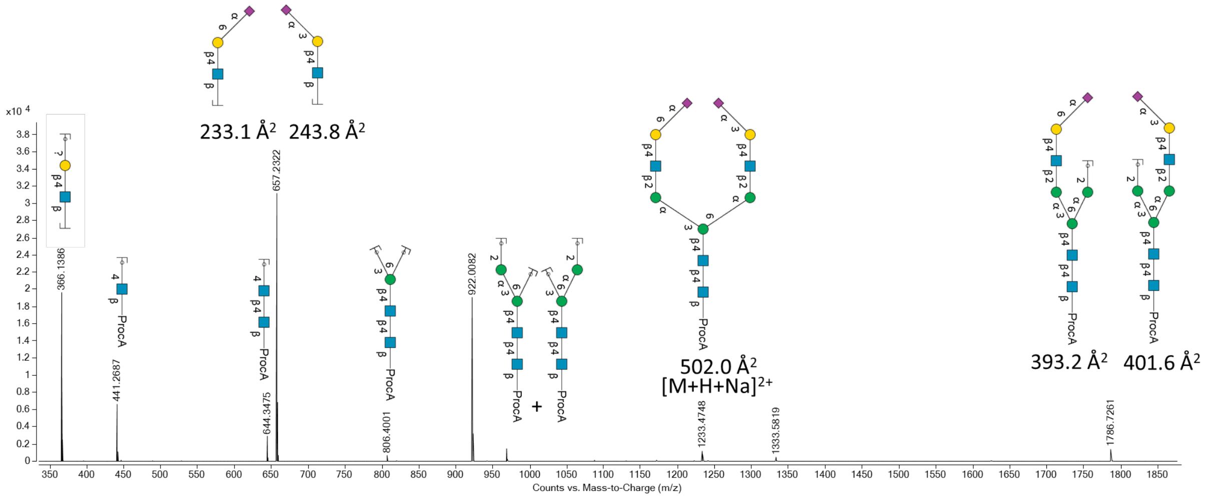

**Supplementary Figure 187. Mass fragmentation spectrum of an *N*-glycan derived from the protein transferrin.**

### 8. ATDs of elucidated *N*-glycan structures and standards for structure assignment validation.

#### 8.1 ATDs from *N*-glycans derived from Aflibercept and synthetic standards for validation.

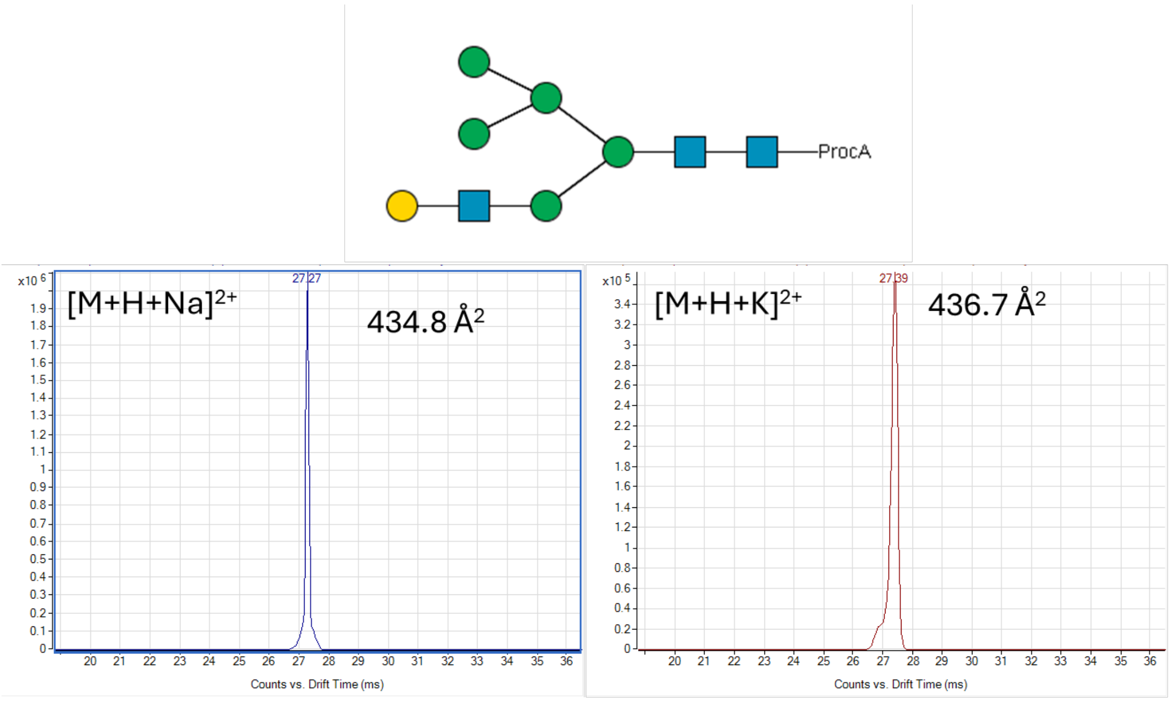

**Supplementary Figure 188. Structure and ATDs of an elucidated *N*-glycan structure derived from Aflibercept. The data was acquired using a fragmentor voltage of 550V (n= 1 measurement).**

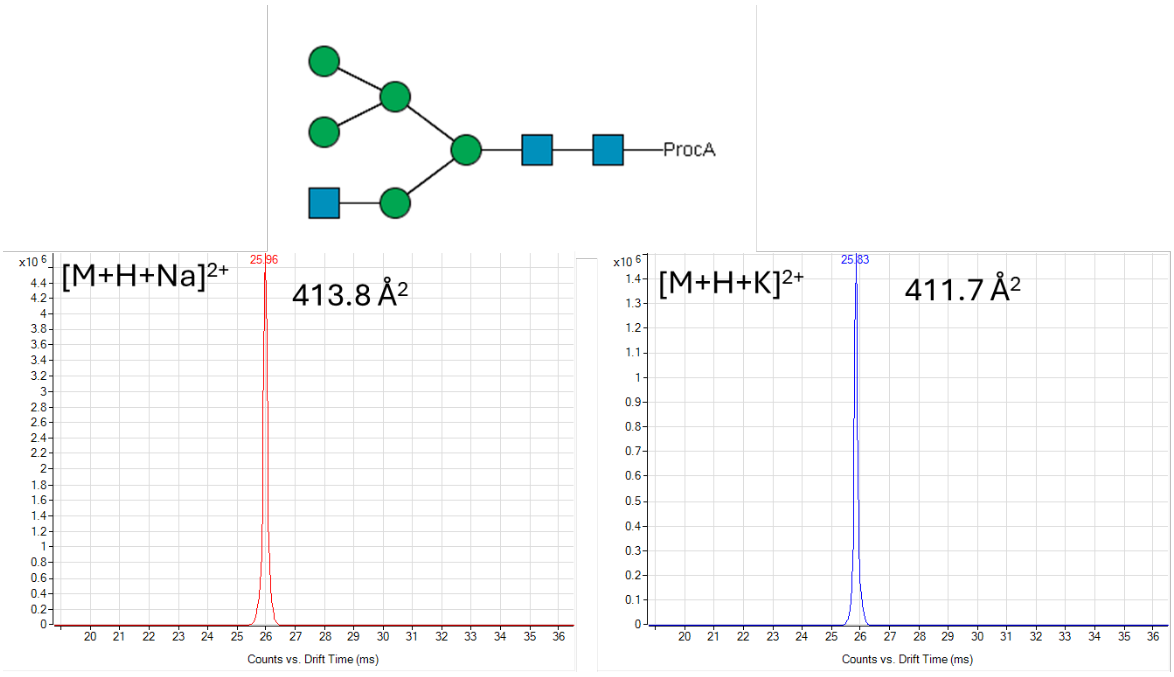

**Supplementary Figure 189. Structure and ATDs of an elucidated *N*-glycan structure derived from Aflibercept. The data was acquired using a fragmentor voltage of 550 V (n= 1 measurement).**

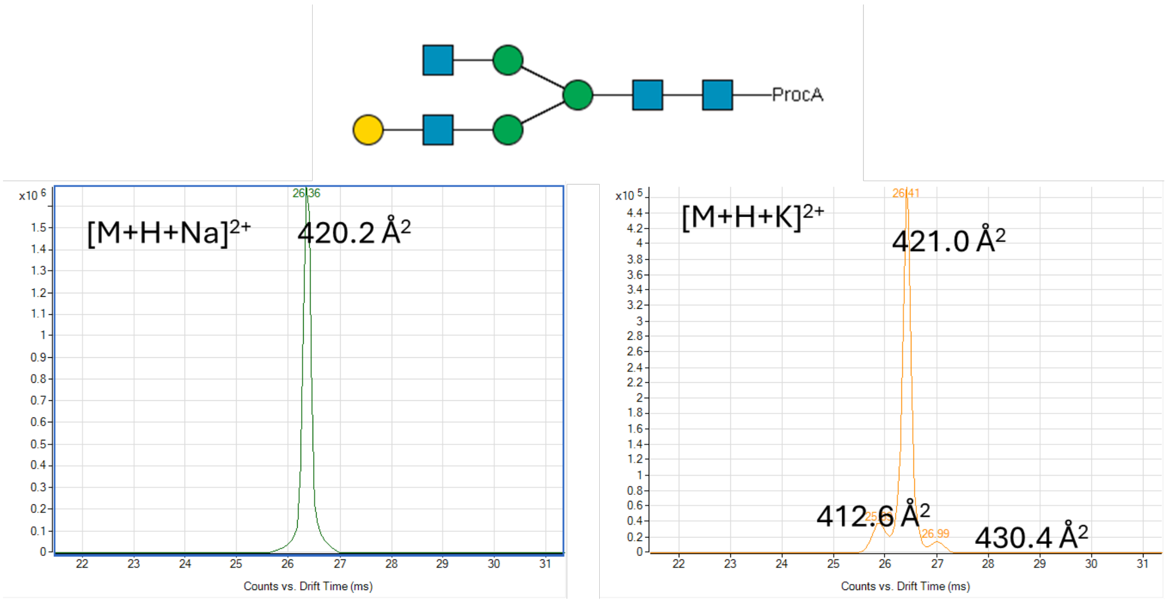

**Supplementary Figure 190. Structure and ATDs of an elucidated *N*-glycan structure derived from Aflibercept. The data was acquired using a fragmentor voltage of 550 V (n= 1 measurement).**

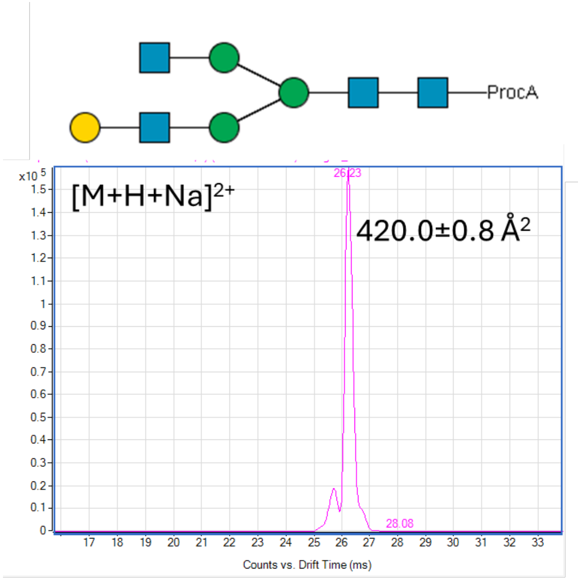

**Supplementary Figure 191. Structure and ATDs of an *N*-glycan standard for validation. The data was acquired using a fragmentor voltage of 360 V (n=3 measurements).**

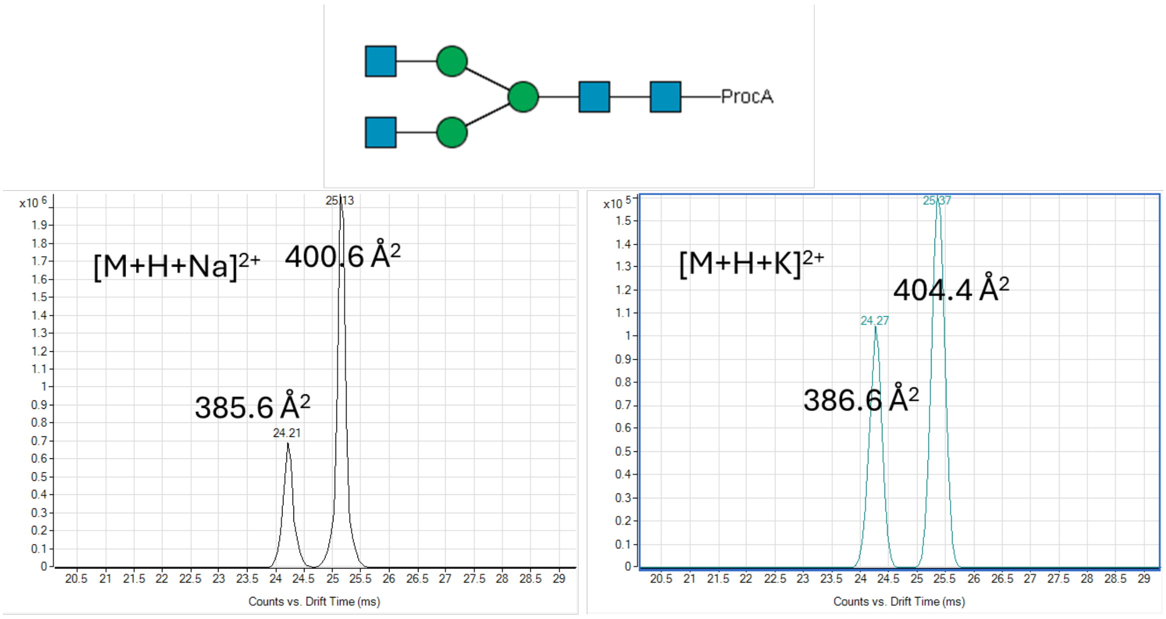

**Supplementary Figure 192. Structure and ATDs of an elucidated *N*-glycan structure derived from Aflibercept. The data was acquired using a fragmentor voltage of 550 V (n= 1 measurement).**

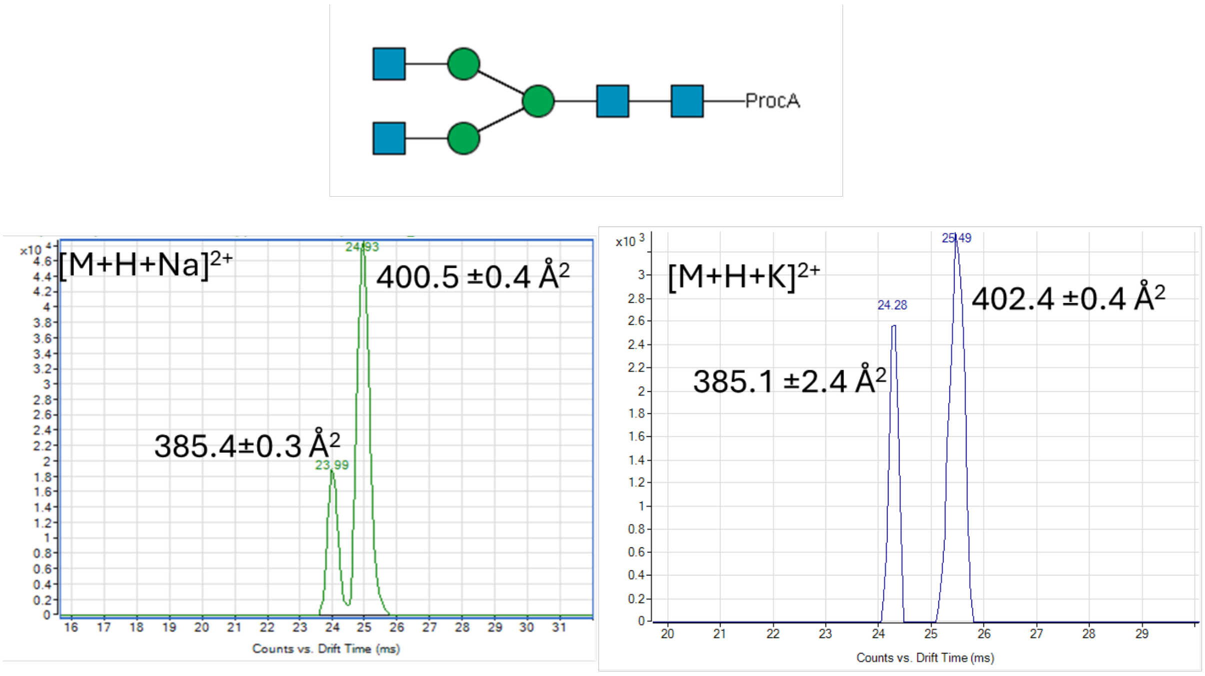

**Supplementary Figure 193. Structure and ATDs of an *N*-glycan standard for validation. The data was acquired using a fragmentor voltage of 360 V (n=3 measurements).**

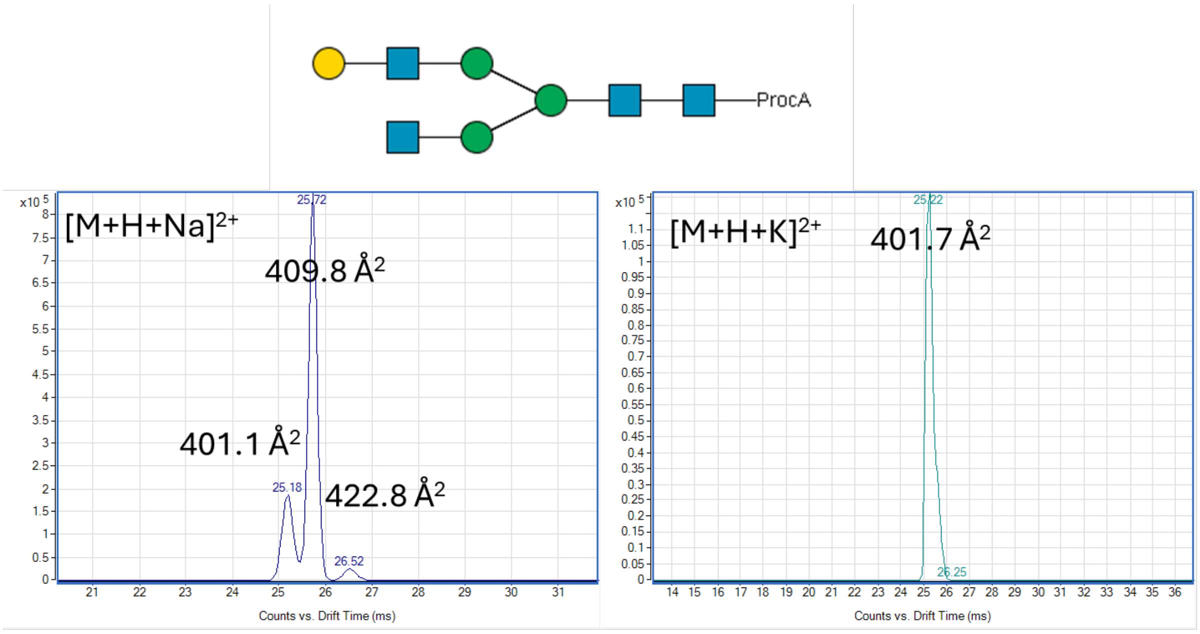

**Supplementary Figure 194. Structure and ATDs of an elucidated *N*-glycan structure derived from Aflibercept. The data was acquired using a fragmentor voltage of 550 V (n= 1 measurement).**

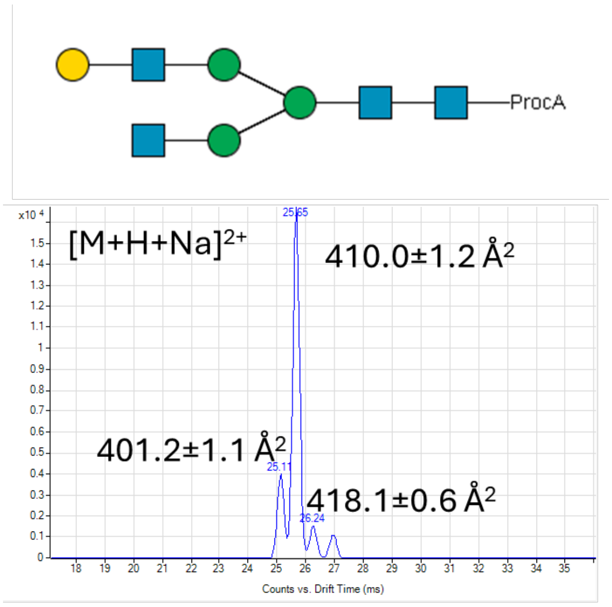

**Supplementary Figure 195. Structure and ATDs of an *N*-glycan standard for validation. The data was acquired using a fragmentor voltage of 360 V (n=3 measurements).**

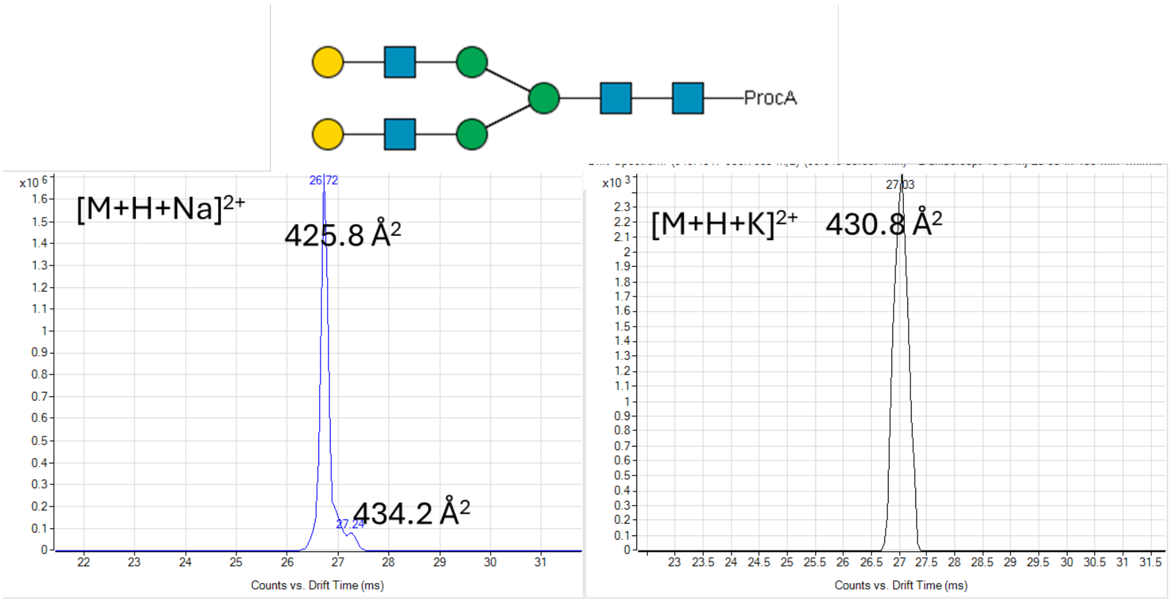

**Supplementary Figure 196. Structure and ATDs of an elucidated *N*-glycan structure derived from Aflibercept. The data was acquired using a fragmentor voltage of 550 V (n= 1 measurement).**

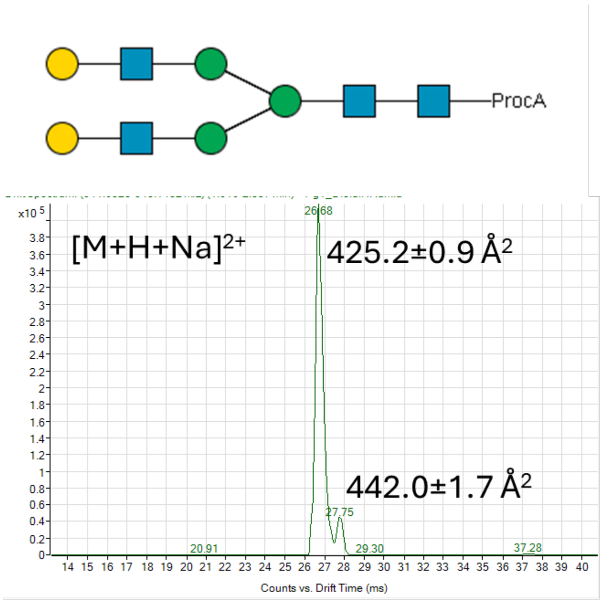

**Supplementary Figure 197. Structure and ATDs of an *N*-glycan standard for validation. The data was acquired using a fragmentor voltage of 360 V (n= 3 measurements).**

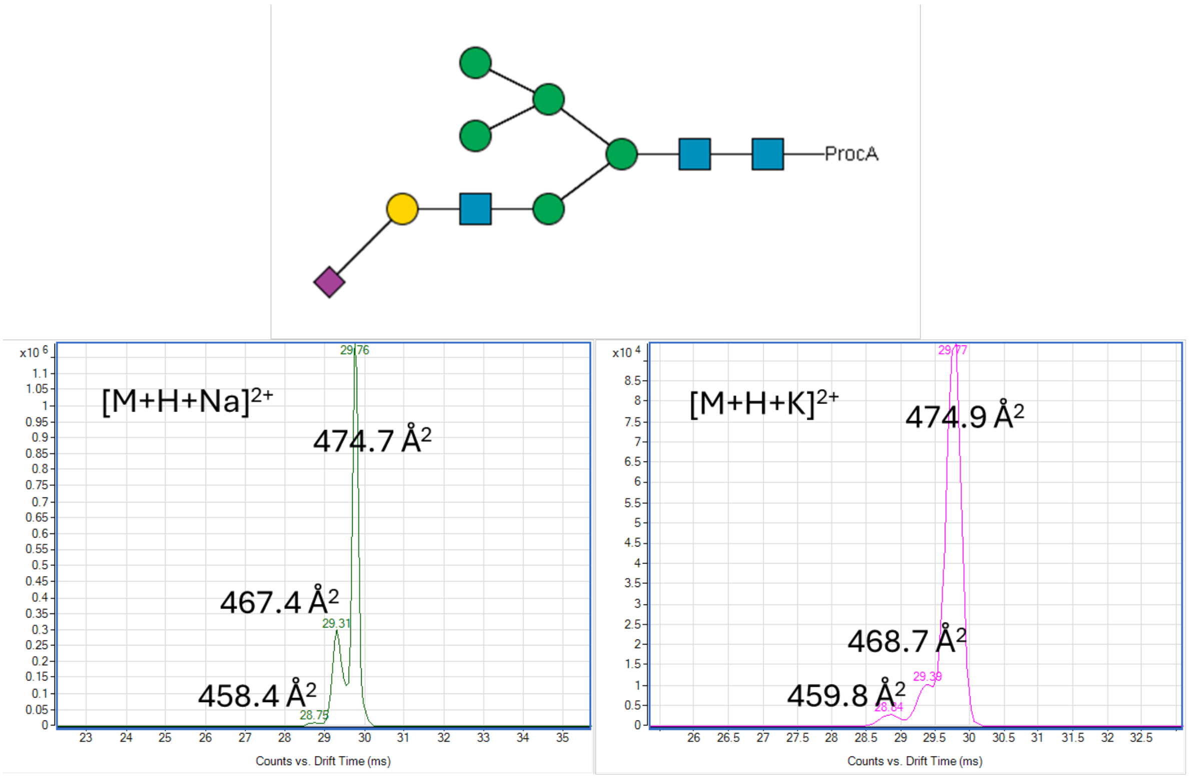

**Supplementary Figure 198. Structure and ATDs of an elucidated *N*-glycan structure derived from Aflibercept. The data was acquired using a fragmentor voltage of 550 V (n= 1 measurement).**

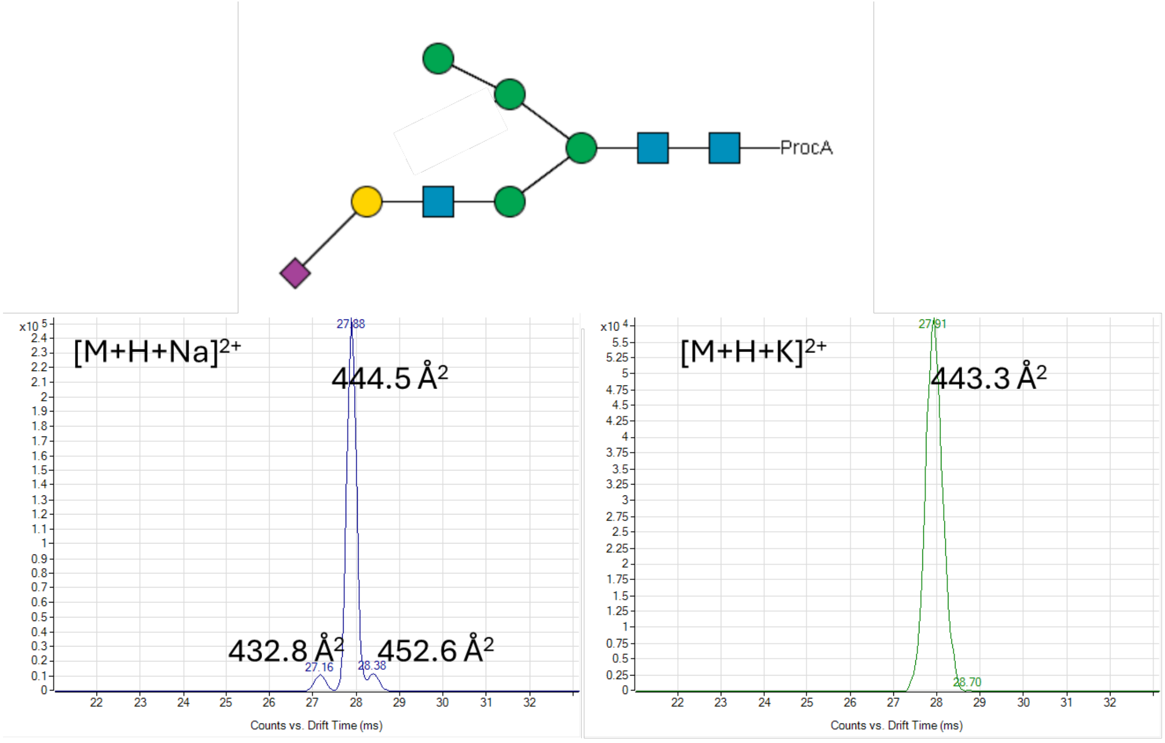

**Supplementary Figure 199. Structure and ATDs of an elucidated *N*-glycan structure derived from Aflibercept. The data was acquired using a fragmentor voltage of 550 V.**

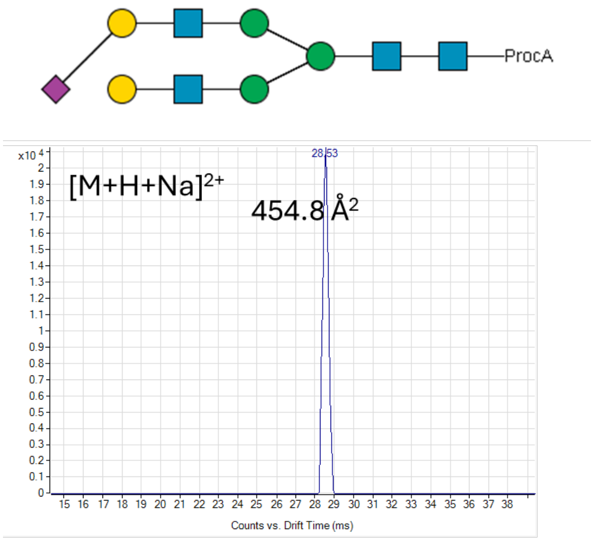

**Supplementary Figure 200. Structure and ATDs of an elucidated *N*-glycan structure derived from Aflibercept. The data was acquired using a fragmentor voltage of 550 V (n= 1 measurement).**

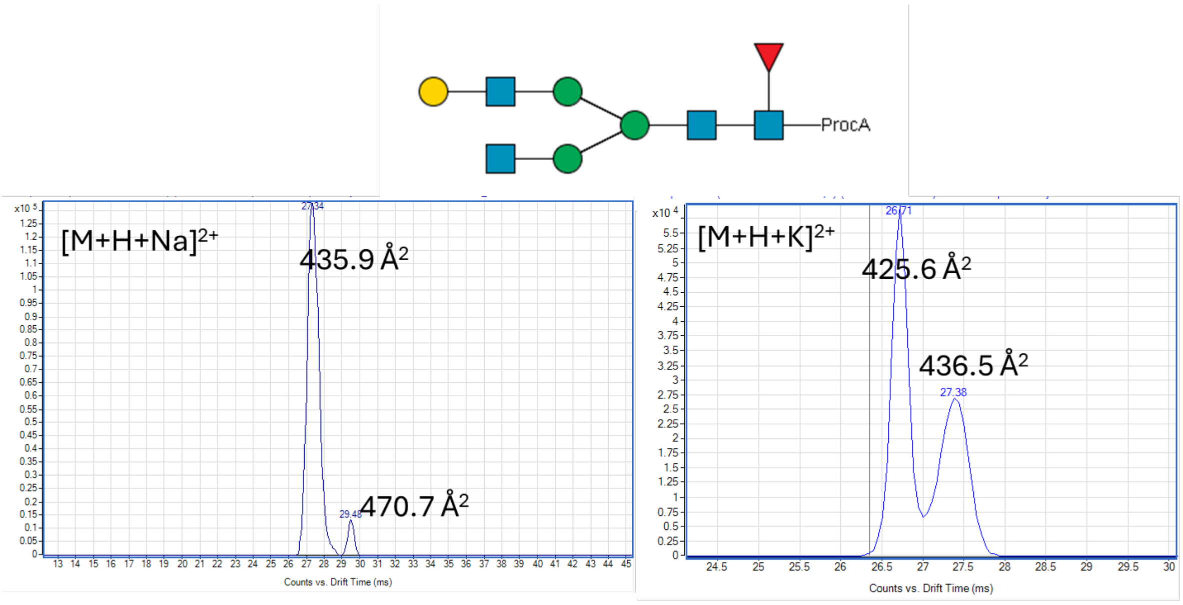

**Supplementary Figure 201. Structure and ATDs of an elucidated *N*-glycan structure derived from Aflibercept. The data was acquired using a fragmentor voltage of 550 V (n= 1 measurement).**

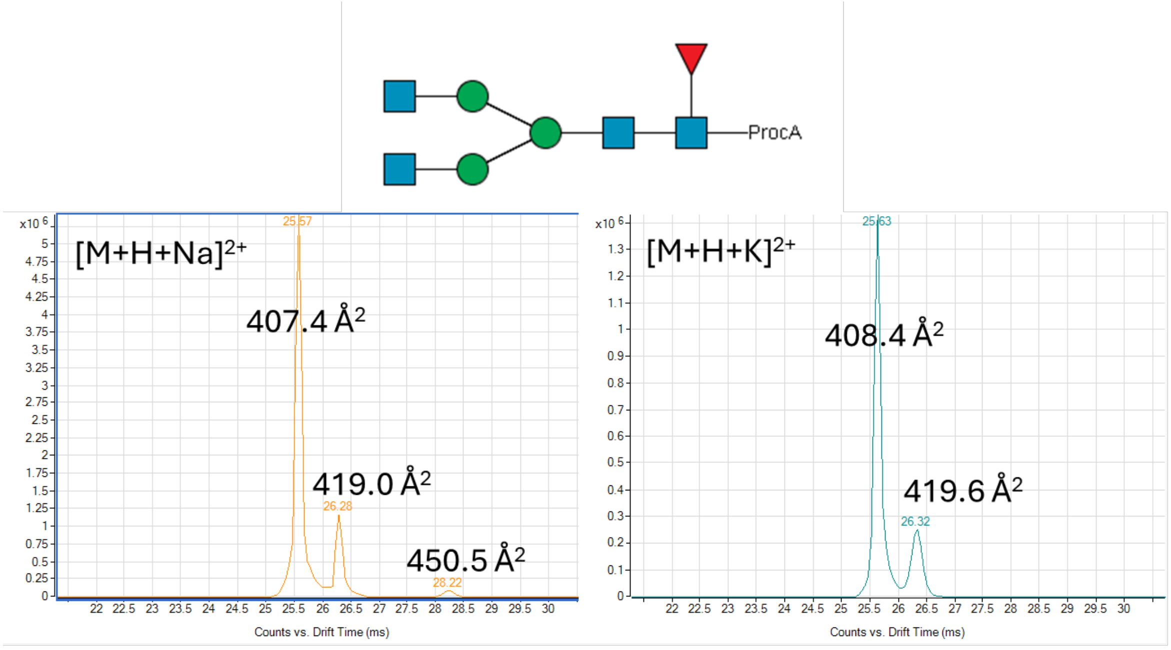

**Supplementary Figure 202. Structure and ATDs of an elucidated *N*-glycan structure derived from Aflibercept. The data was acquired using a fragmentor voltage of 550 V (n= 1 measurement).**

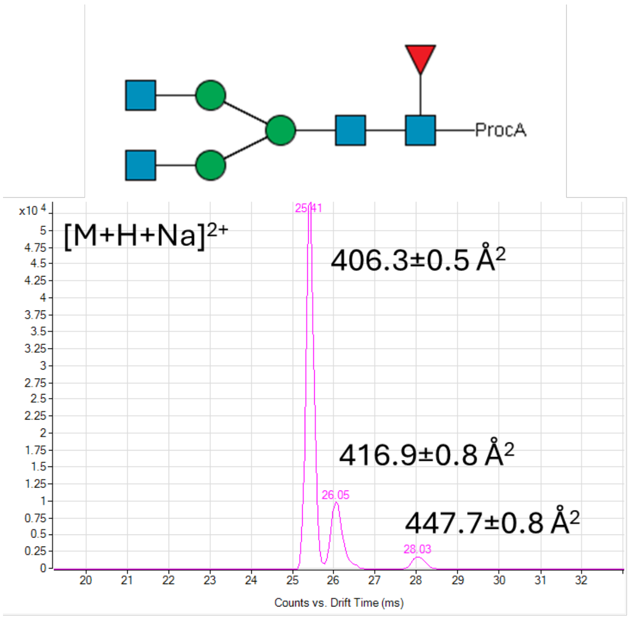

**Supplementary Figure 203. Structure and ATDs of an *N*-glycan standard for validation. The data was acquired using a fragmentor voltage of 360 V (n=3 measurements).**

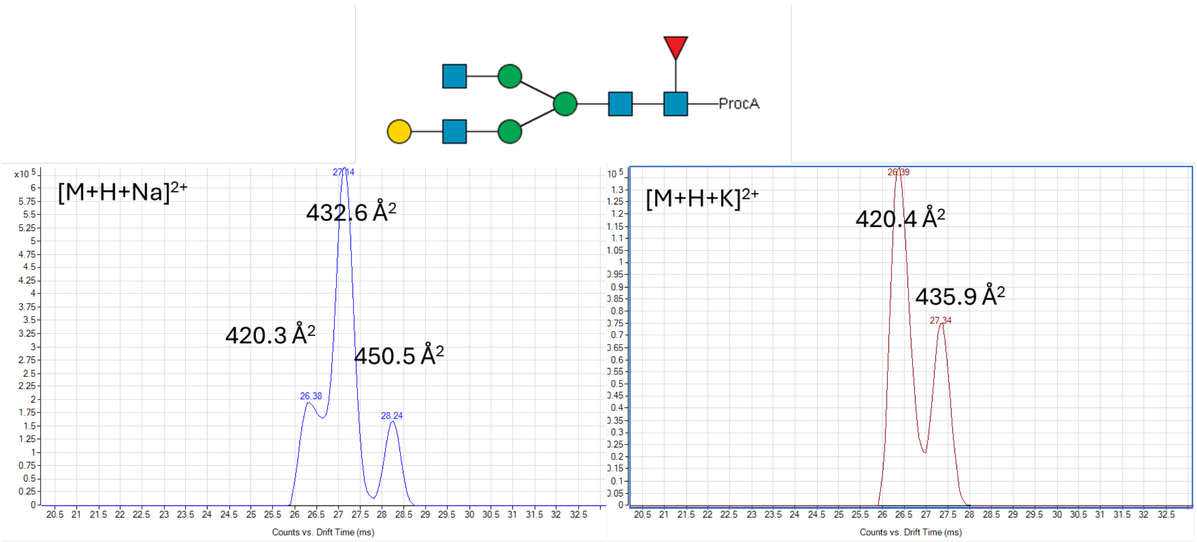

**Supplementary Figure 204. Structure and ATDs of an elucidated *N*-glycan structure derived from Aflibercept. The data was acquired using a fragmentor voltage of 550 V (n= 1 measurement).**

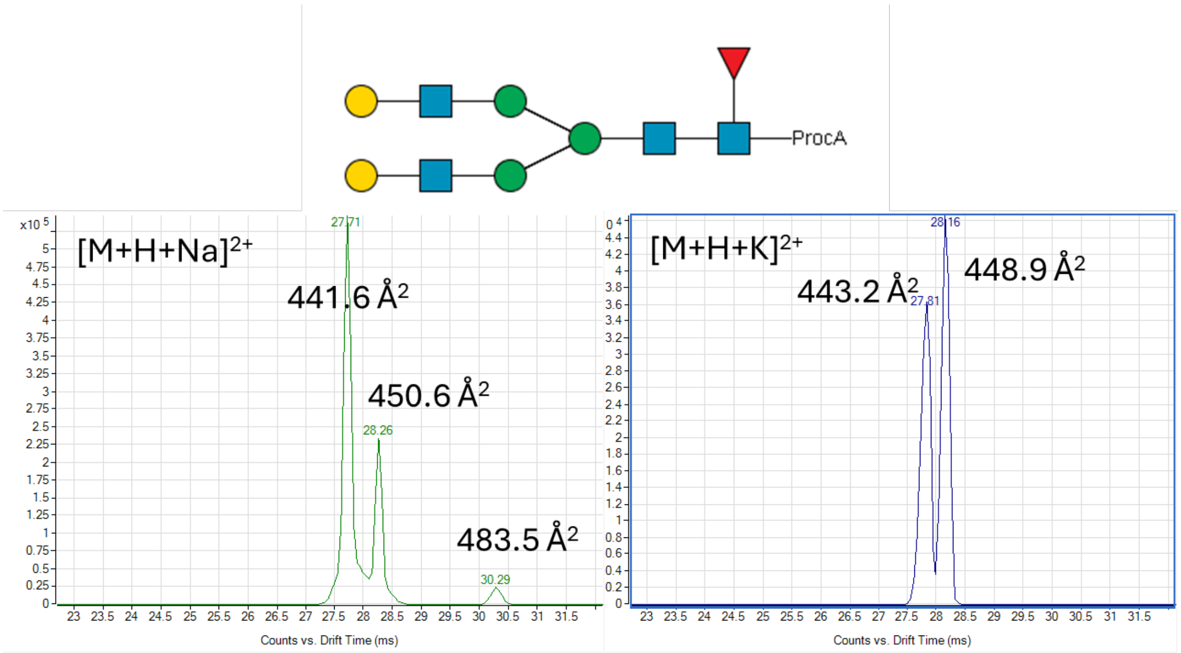

**Supplementary Figure 205. Structure and ATDs of an elucidated *N*-glycan structure derived from Aflibercept. The data was acquired using a fragmentor voltage of 550 V (n= 1 measurement).**

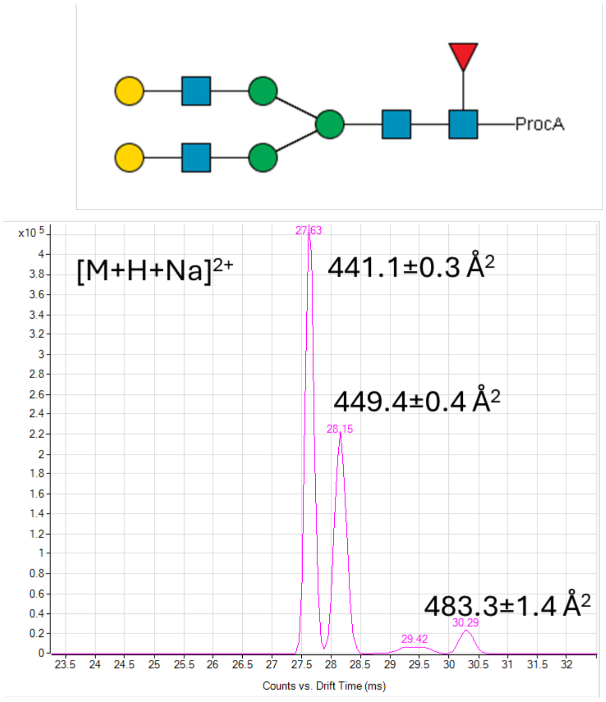

**Supplementary Figure 206. Structure and ATDs of an *N*-glycan standard for validation. The data was acquired using a fragmentor voltage of 360 V (n=3 measurements).**

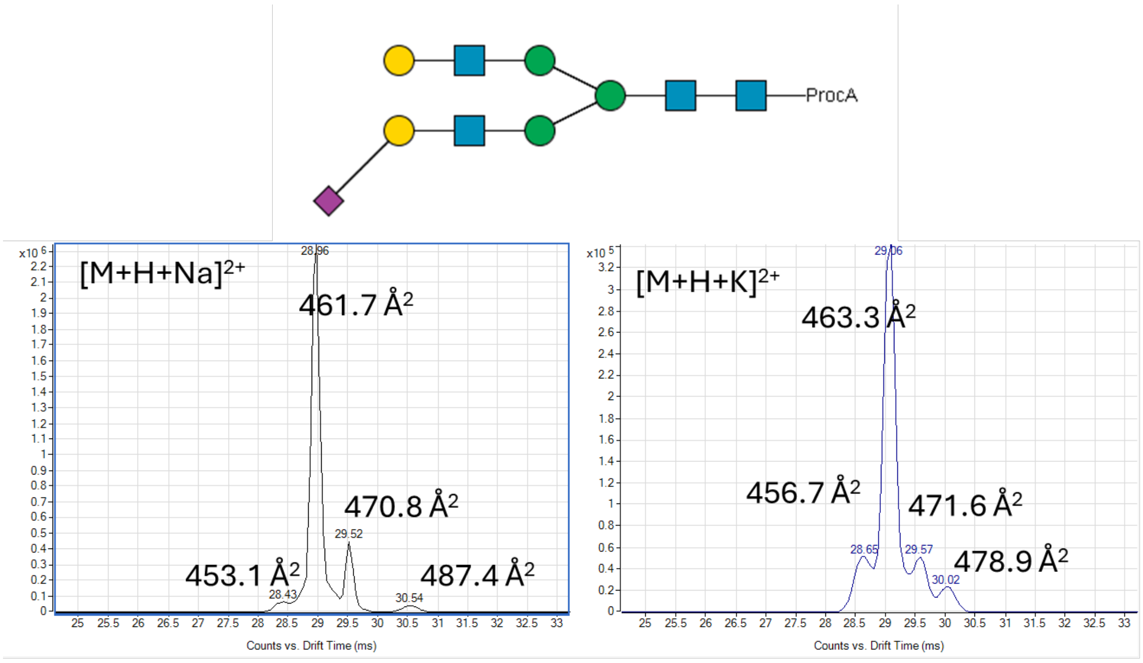

**Supplementary Figure 207. Structure and ATDs of an elucidated *N*-glycan structure derived from Aflibercept. The data was acquired using a fragmentor voltage of 550 V (n= 1 measurement).**

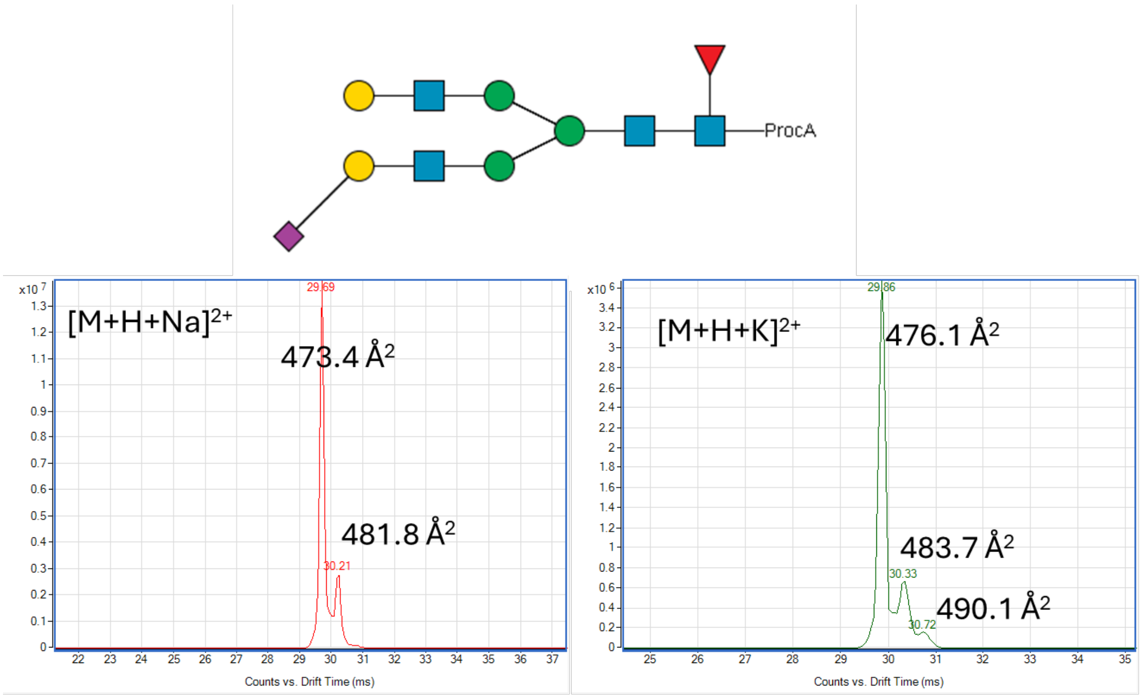

**Supplementary Figure 208. Structure and ATDs of an elucidated *N*-glycan structure derived from Aflibercept. The data was acquired using a fragmentor voltage of 550 V (n= 1 measurement).**

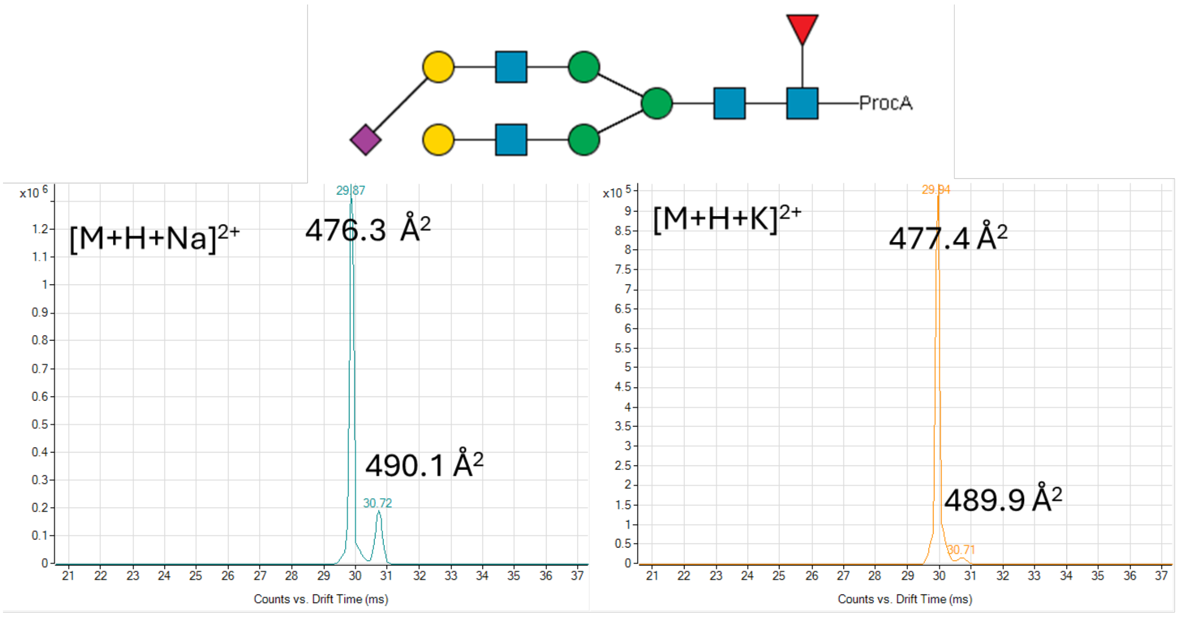

**Supplementary Figure 209. Structure and ATDs of an elucidated *N*-glycan structure derived from Aflibercept. The data was acquired using a fragmentor voltage of 550 V (n= 1 measurement).**

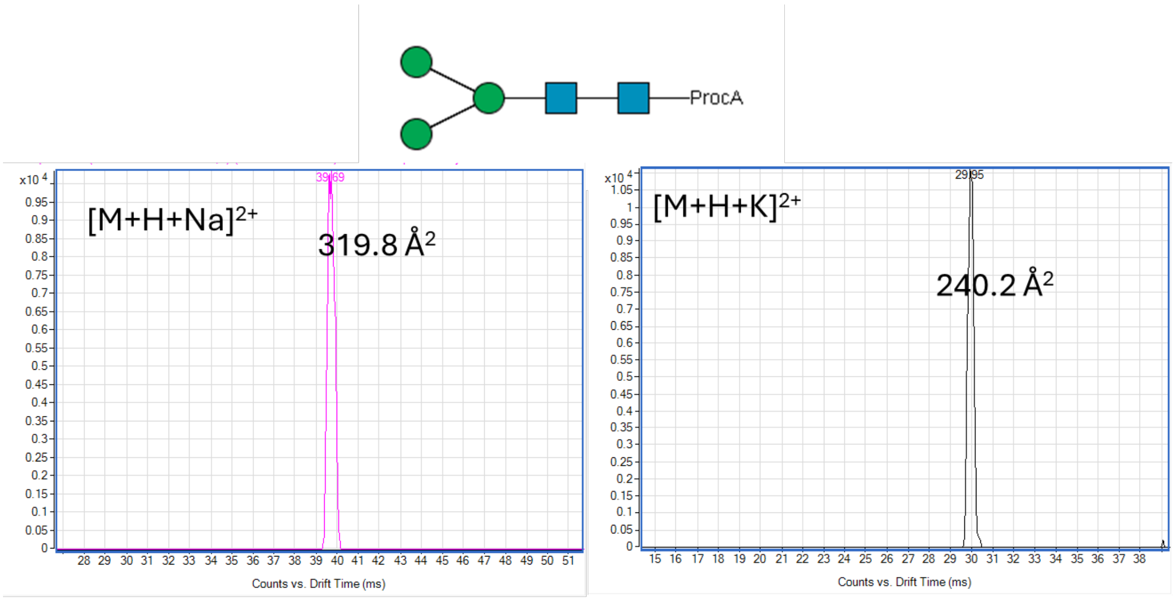

**Supplementary Figure 210. Structure and ATDs of an elucidated *N*-glycan structure derived from Aflibercept. The data was acquired using a fragmentor voltage of 550 V (n=1 measurement).**

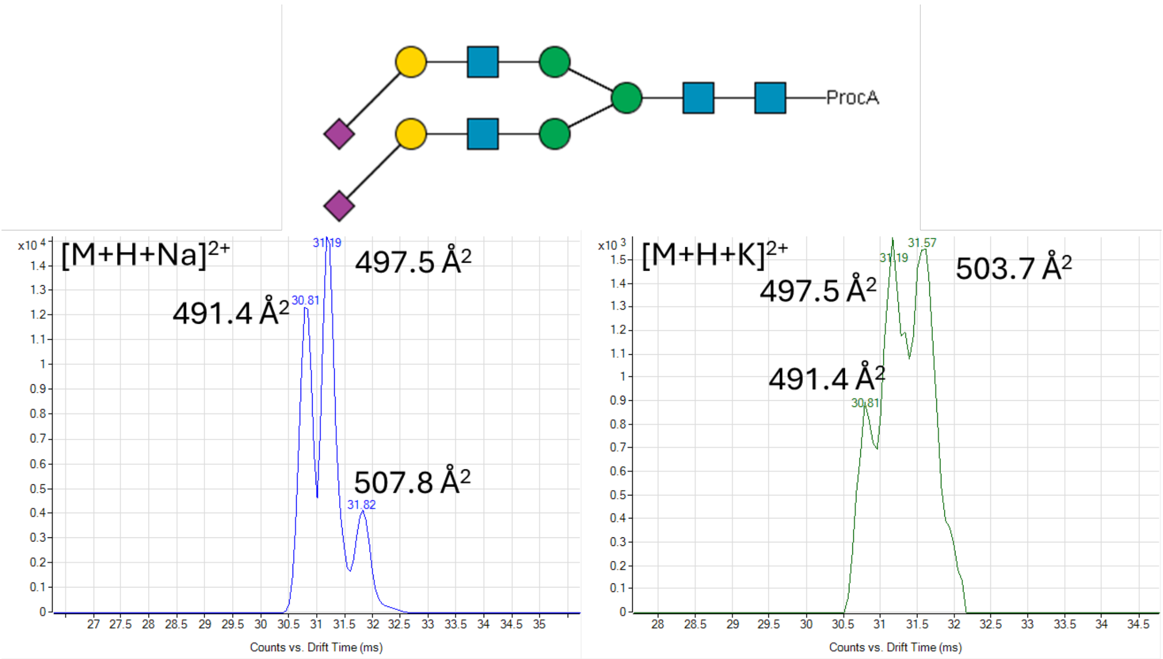

**Supplementary Figure 211. Structure and ATDs of an elucidated *N*-glycan structure derived from Aflibercept. The data was acquired using a fragmentor voltage of 550 V (n= 1 measurement).**

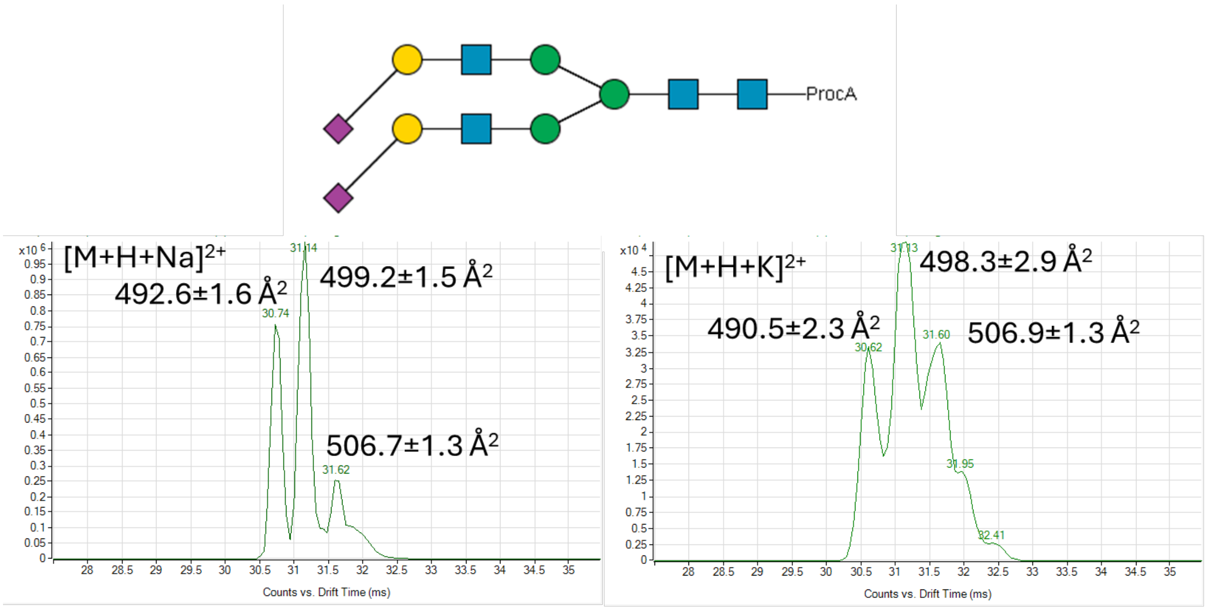

**Supplementary Figure 212. Structure and ATDs of an *N*-glycan standard for validation. The data was acquired using a fragmentor voltage of 360 V (n=3 measurements).**

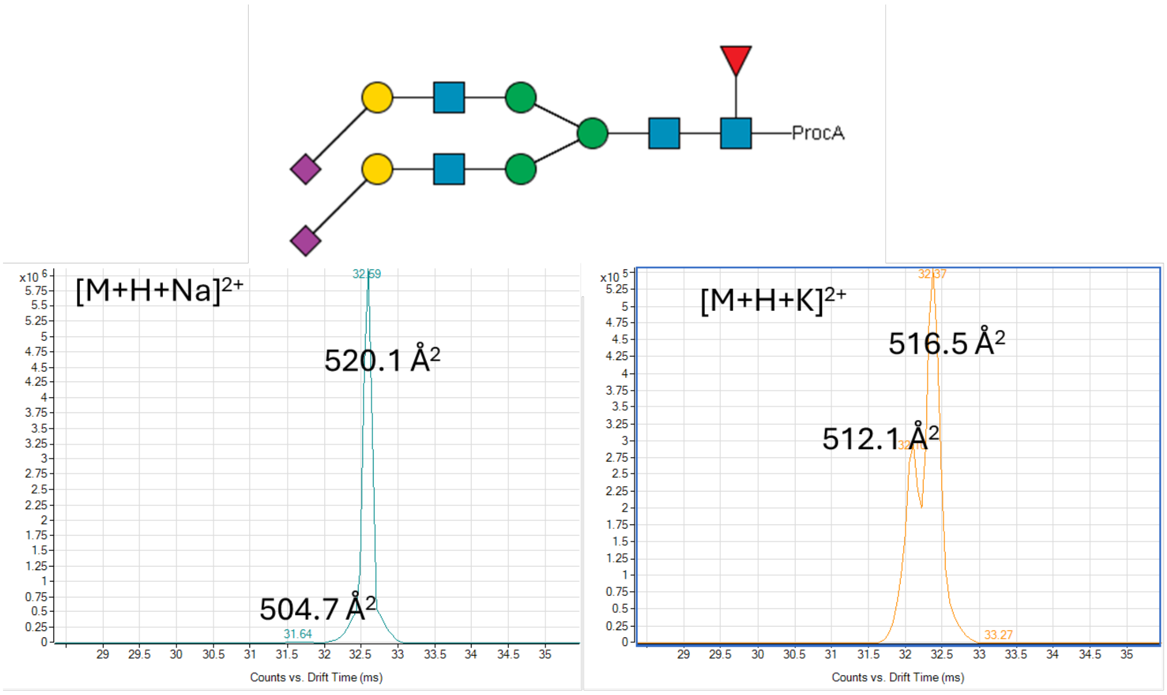

**Supplementary Figure 213. Structure and ATDs of an elucidated *N*-glycan structure derived from Aflibercept. The data was acquired using a fragmentor voltage of 550 V (n= 1 measurement).**

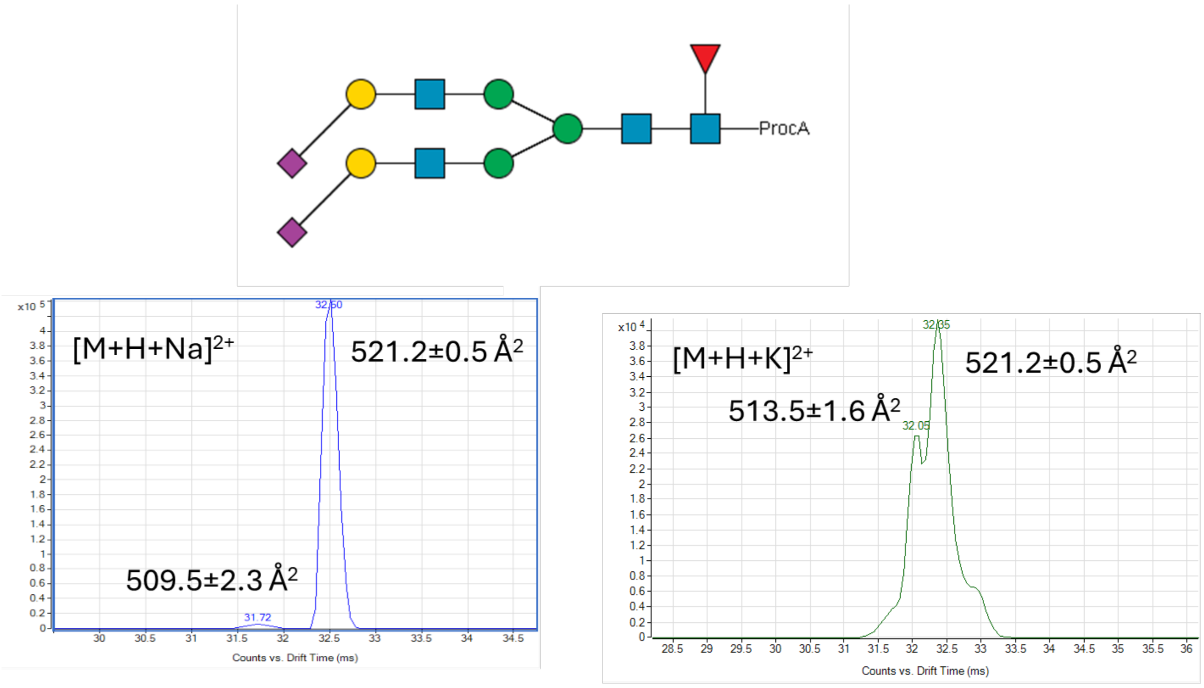

**Supplementary Figure 214. Structure and ATDs of an *N*-glycan standard for validation. The data was acquired using a fragmentor voltage of 360 V (n=3 measurements).**

#### 8.2 ATDs from *N*-glycans derived from transferrin and synthetic standards for validation.

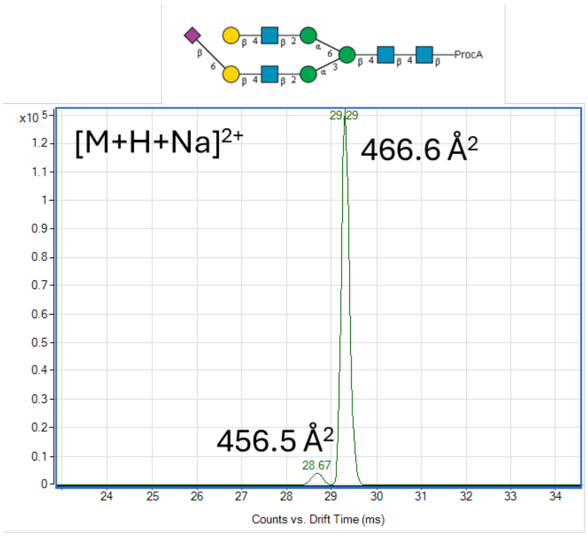

**Supplementary Figure 215. Structure and ATDs of an elucidated N-glycan structure derived from transferrin. The data was acquired using a fragmentor voltage of 550 V (n= 1 measurement).**

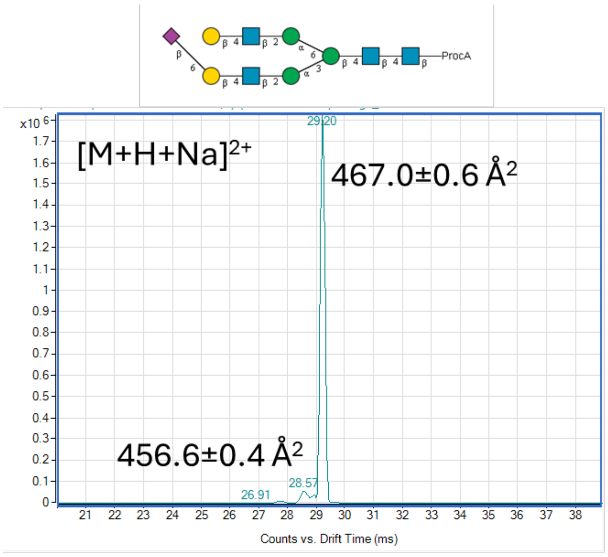

**Supplementary Figure 216. Structure and ATDs of an *N*-glycan standard for validation. The data was acquired using a fragmentor voltage of 360 V (n=3 measurements).**

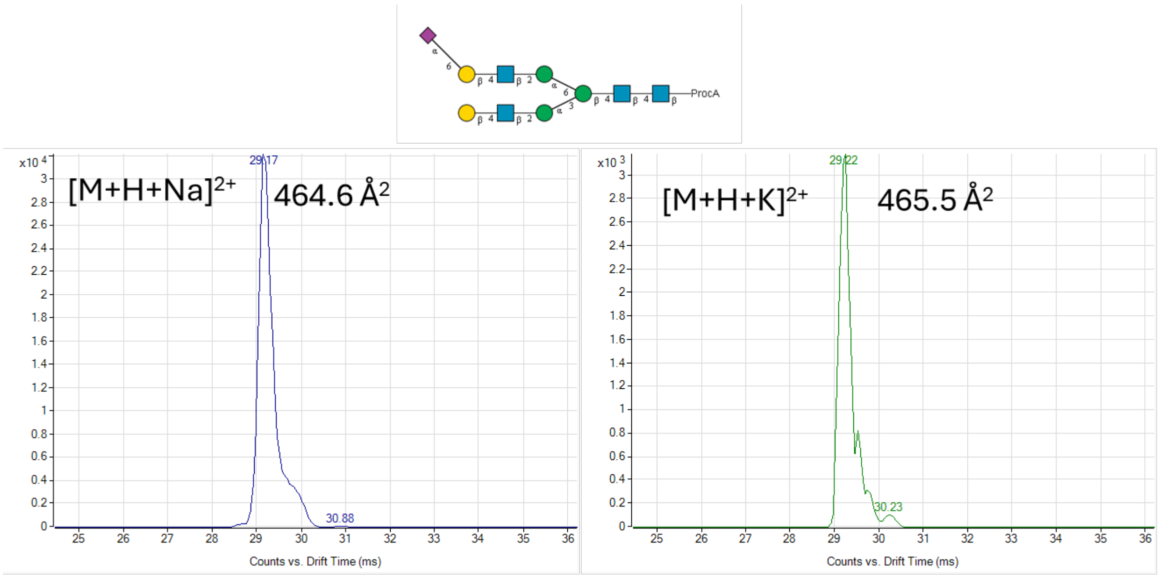

**Supplementary Figure 217. Structure and ATDs of an elucidated N-glycan structure derived from transferrin. The data was acquired using a fragmentor voltage of 550 V (n= 1 measurement).**

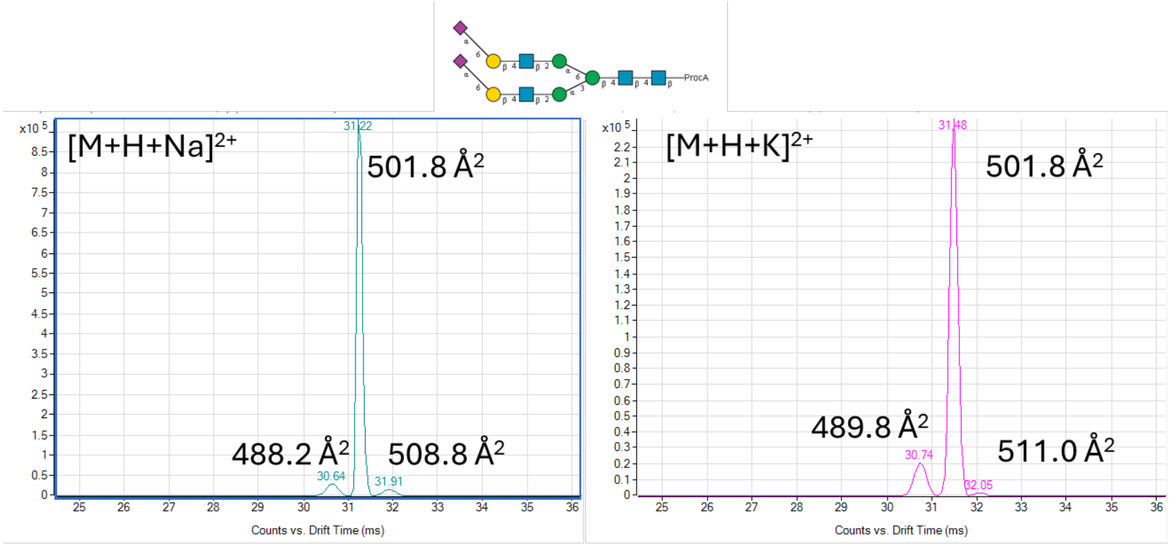

**Supplementary Figure 218. Structure and ATDs of an elucidated N-glycan structure derived from transferrin. The data was acquired using a fragmentor voltage of 550 V (n= 1 measurement).**

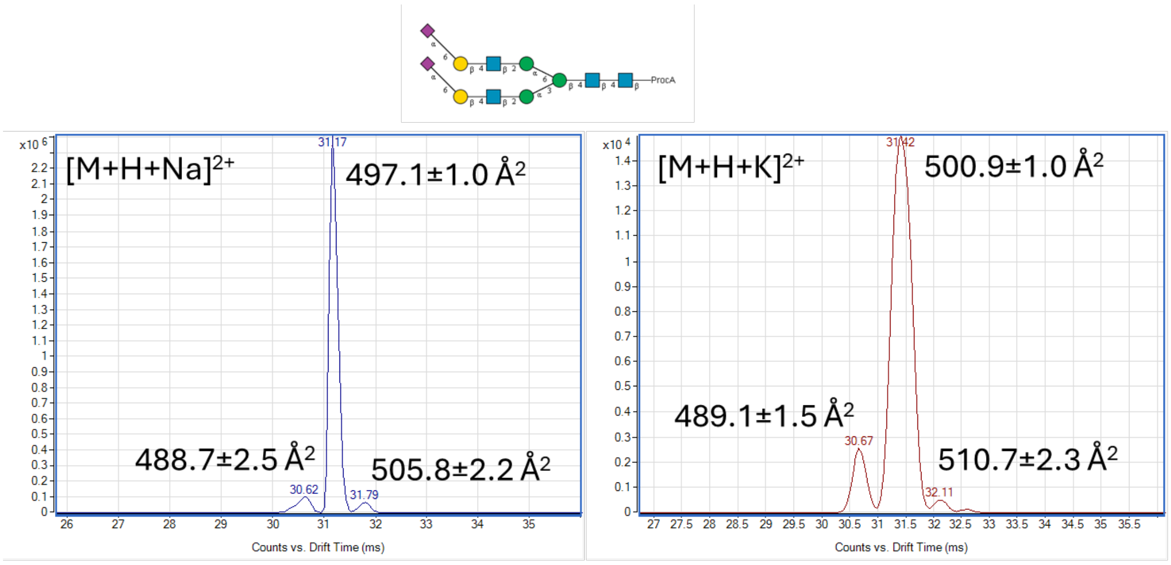

**Supplementary Figure 219. Structure and ATDs of an *N*-glycan standard for validation. The data was acquired using a fragmentor voltage of 360 V (n=3 measurements).**

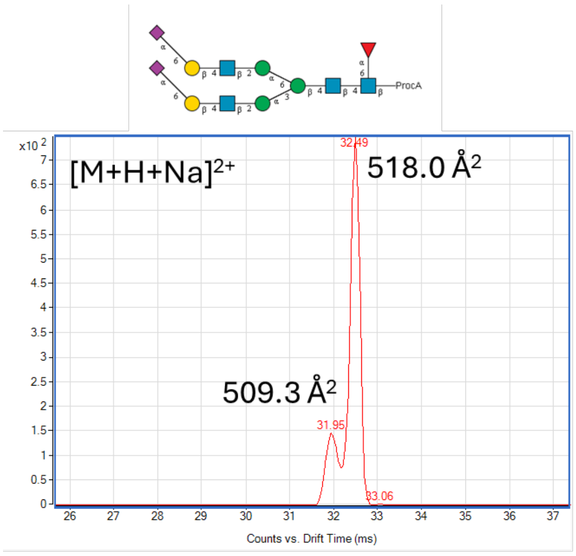

**Supplementary Figure 220. Structure and ATDs of an elucidated N-glycan structure derived from transferrin. The data was acquired using a fragmentor voltage of 550 V (n= 1 measurement).**

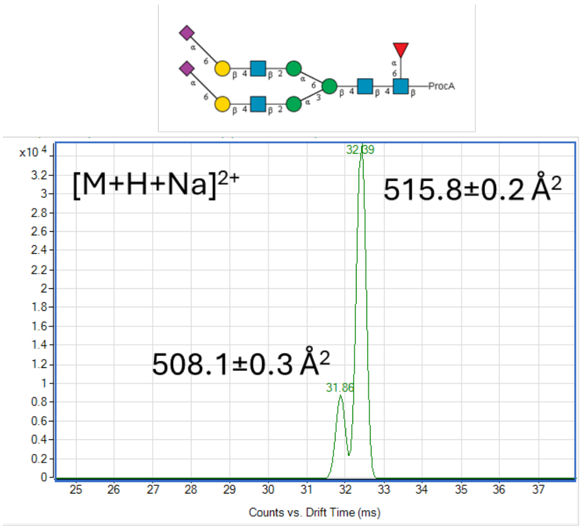

**Supplementary Figure 221. Structure and ATDs of an *N*-glycan standard for validation. The data was acquired using a fragmentor voltage of 360 V (n=3 measurements).**

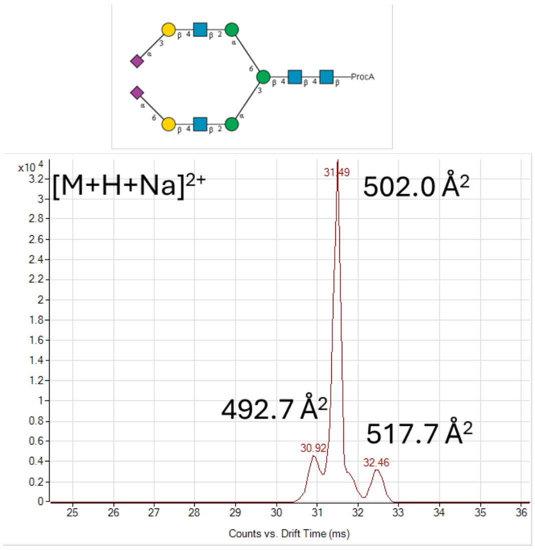

**Supplementary Figure 222. Structure and ATDs of an elucidated N-glycan structure derived from transferrin. The data was acquired using a fragmentor voltage of 550 V (n= 1 measurement).**

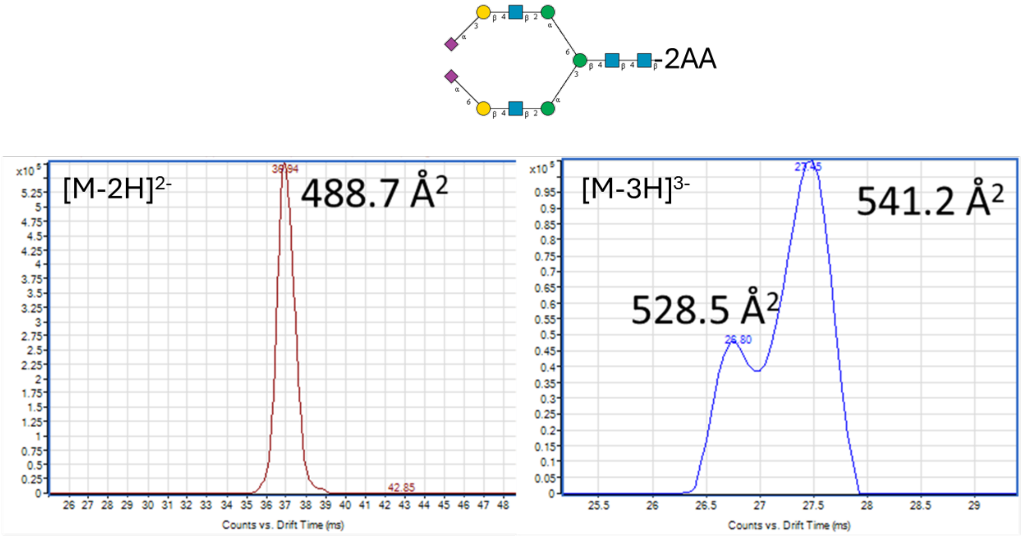

**Supplementary Figure 223. Structure and ATDs of an elucidated *N*-glycan structure derived from transferrin. The data was acquired using a fragmentor voltage of 360 V (n=1 measurement).**
